# Pangenome Discoveries from Clinical Long-Read Genomes in Underrepresented Populations with Neurodevelopmental Disorders

**DOI:** 10.64898/2026.11.07.765034

**Authors:** Mohammad Amiruddin Hashmi, Nasna Nassir, Bipin Balan, Shehzad Hanif, Bassam Jamalalail, Mariam Eldesouky, Richa Tambi, Shuhd BinEshaq, K Dasuki Hansana Perera, Hanan Elsokary, Suhana Shiyas, Binte Zehra, Dia Advani, Solima Sabeel, Omar Bashir, Ahmed Khalifa, Aysha Ibrahim, Maryam Al Obathani, Shaima Alsuwaidi, Mohammed Yousuff, Nesrin Mohamed, Alexander Giddey, Naser Alshamsi, Wen-Wei Liao, Prajna Hebbar, Peter A. Audano, Amirul Islam, Omer S AlKhnbashi, Hosneara Akter, Shaoli Sarker, Reem K. Jan, Munazza Murtaza, Bakhrom Berdiev, Nelson C. Soares, Babacar Cisse, Shola Faniran, Nidheesh Chencheri, Haitham Elbashir, Syed Ali Bokhari, Rasha Buhumaid, Hanan Alsuwaidi, Karen Miga, Erich Jarvis, Mohamed Almarri, Human Pangenome Reference Consortium (HPRC), Stephen W. Scherer, Ammar Albanna, Maahil Arshad, Brett Trost, Egor Dolzhenko, Stefan S Du Plessis, Benedict Paten, Alawi Alsheikh-Ali, Mohammed Uddin

## Abstract

Genetic diagnoses remain elusive for many individuals with neurodevelopmental disorders (NDDs), and the gap is wider in populations poorly represented in genomic reference resources. To demonstrate the equitable advantage of long-read whole genome sequencing (LR-WGS) and pangenomes in terms of clinical utility, we analyzed 211 NDD cases and 107 unaffected relatives from 24 populations, integrating their assemblies, resulting in a total of 1204 haplotypes with the Human Pangenome Reference Consortium Release 2 (HPRC_r2) and the Arab Pangenome Reference (APR) datasets. Clinical interpretation of small variants and structural variants (SVs) revealed pathogenic or likely pathogenic findings in 38.9% of the cases (82/211), and our workflow further detected pathogenic repeat expansions in 3.8% of the cases and runs of homozygosity (ROHs) exceeding 300 Mb in 4.7% of the cases, revealing substantial autozygosity suggestive of strong inbreeding. Beyond these clinical assessments, pangenome analysis revealed 3.9 million population-level novel small variants, 39,782 SVs and an average of 2.17 Mb of novel sequence per individual absent from linear references and global pangenomes. We identified 85.9 kb of case-unique nonreference sequences in exonic segments overlapping with curated NDD gene sets. Graph-based tertiary analysis further identified rare exonic complex homozygous rearrangements in the NDD associated genes for 2.4% of the cases. Together, these findings emphasize how diverse genome assemblies extend variant discovery beyond established linear reference-based clinical interpretation.

## Introduction

Neurodevelopmental disorders (NDDs) comprise a heterogeneous group of conditions affecting brain development and cognitive function, including intellectual disability, autism spectrum disorder, epilepsy, developmental delay, and congenital brain malformations^1^. Collectively, NDDs affect more than 3% of children worldwide and represent a major cause of lifelong disability and healthcare burden^2^. Genetic studies have implicated diverse classes of variation, including single-nucleotide variants (SNVs), insertions/deletions (INDELs), structural variants (SVs), and repeat expansions, which converge on overlapping neurodevelopmental pathways. Despite major advances in genomic sequencing technologies and their integration into clinical diagnostics, a substantial proportion of NDD cases, often approximately 80–90%, remain without a definitive molecular diagnosis, with approximately 1 in 10 children meeting diagnostic criteria for a neurodevelopmental condition^3–6^. This diagnostic gap is particularly pronounced in populations that remain underrepresented in large-scale genomic datasets, where limited population-specific reference data^7^ constrain both variant discovery and interpretation, potentially widening existing inequities in genomic medicine.

A key limitation is the continued reliance on a single linear human reference genome. GRCh38, although fundamental to human genetics, represents a linear reference assembled from a limited number of donors and does not capture the full extent of global genomic diversity^8,9^. Linear reference approaches perform well for many small variants but are less able to represent population-specific sequences, including SVs, gene duplications, and rearrangements within repetitive regions. Such variants are established contributors to rare diseases, including NDDs, but can remain difficult to detect or resolve using short-read sequencing and a single reference framework^10^. In rare disease genomics, pathogenic variants that reside in structurally complex or population-specific regions are often missed^11^ because of the reliance on a single reference genome with little to no representation of some of the most diverse and underrepresented populations.

Recent advances in long-read sequencing (LRS) have enabled highly contiguous diploid genome assemblies and accelerated the development of human pangenomes^12–14^. Unlike a single linear reference, pangenome frameworks represent alternative haplotypes, structural diversity and population-specific sequences within a unified genomic framework. Compared with single reference approaches, the Human Pangenome Reference Consortium (HPRC) has demonstrated improved representation and discovery of structural variations^12,13,15,16^. However, individuals affected by rare genetic disorders remain sparsely represented in current pangenome resources, and the extent to which pangenome analysis improves disease-associated variant discovery remains incompletely defined.

Pangenome references therefore offer a powerful framework for expanding genomic analyses of underrepresented populations, with the aim of improving rare disease diagnostics^17^. Populations across Africa, North Africa, the Middle East, and Central/South Asia encompass distinct demographic histories, including varying degrees of consanguinity, endogamy, founder effects, and population substructure^18,19^. These features can increase genome-wide autozygosity and the probability that recessive disease alleles occur in the homozygous form^20^. Beyond recessive variation, gene duplications, large sequence insertions, tandem-repeat expansions, splice-altering variants and complex haplotype structures may also contribute to neurodevelopmental disease but remain incompletely characterized in these populations^21,22^. The extent to which assembly driven pangenome analyses can resolve these forms of variation and uncover the NDD-associated genomic architecture in underrepresented populations remains poorly defined. A recent study used linear pangenome-derived assemblies as controls to filter variants and to investigate missing autism variants^23^. Building graph pangenomes directly from patient-derived assemblies, particularly from underrepresented populations, has the potential to capture cohort-wide genetic diversity within a single representation. Rather than relying solely on individual assemblies or pangenome-derived linear references, such a framework would harmonize haplotype alleles across samples, enabling systematic characterization of small variants, SVs and complex rearrangements in NDDs.

Here, we applied PacBio HiFi whole genome LRS, haplotype assembly and pangenome-based analyses to a clinical NDD cohort from 24 underrepresented populations representing 2.8 billion people spanning Africa, North Africa, the Middle East and South/Central Asia. To explore the equitable advantage and scalability of the clinical genomics framework for rare disease genomics, we assembled 422 haplotypes from 211 NDD cases with 214 haplotypes from unaffected family members and 568 haplotypes from HPRC_r2 and the Arab Pangenome Reference (APR) resource, generating an analytical framework comprising 1,204 haplotypes. By combining conventional linear-reference analysis with assembly and graph-based approaches, we sought to distinguish population-specific genomic diversity from NDD-associated variation and to quantify the additional variation revealed by pangenome analysis across small-variant SNVs and INDELs, SVs, gene duplication, nonreference sequences and complex rearranged genomic loci.

## Results

### A multiancestry NDD cohort from underrepresented populations

This study integrates 1,204 haplotype assemblies from 602 individuals, one of the largest disease-focused collections of haplotype assemblies, comprising 636 haplotypes from the NDD cohort (from 24 populations) and 568 haplotypes from the Human Pangenome Reference Consortium Release 2^24^ (HPRC_r2, n = 231) (from 27 populations) and the Arab Pangenome Reference^13^ (APR, n = 53) (from 8 Middle Eastern populations) cohorts. The study overview is shown in Supplementary Figure 1. The clinical cohort comprised 318 individuals, 211 affected with NDDs (422 haplotypes) and 107 unaffected family members (214 haplotypes) (Figure 1a). Affected individuals were predominantly male (158 males and 53 females; male-to-female ratio, 2.9:1) and were recruited as 150 singletons or within 61 families comprising parent-offspring trios (258 haplotypes), duos (76 haplotypes) and additional family structures. The cohort spanned four superpopulations, with Middle Eastern ancestry predominating (170/211 cases, 80.6%), followed by Central/South Asian (n = 20, 9.5%), North African (n = 12, 5.7%) and African (n = 9, 4.3%) ancestry (Fig. 1b; Supplementary Table 1). This geographic distribution provides a rare multiancestry NDD cohort from populations that remain poorly represented in current large genomic reference resources.

**Figure 1.**
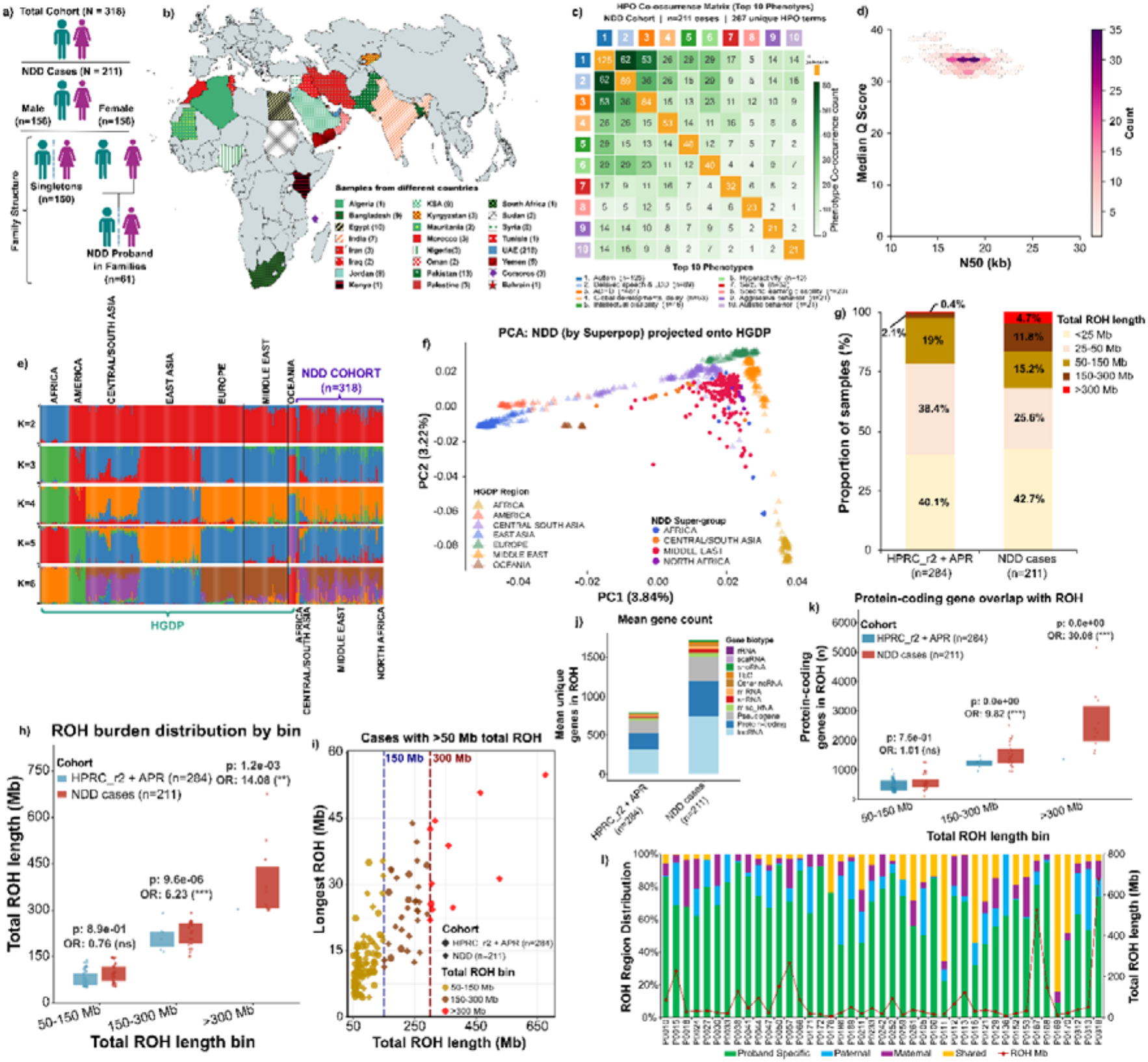
Cohort composition, sequencing quality, population structure and autozygosity in the NDD cohort. **(a)** Composition of the analysis cohort. The cohort comprised 318 individuals, including 211 individuals with NDDs and 107 unaffected family members. Among the affected individuals, 158 were male and 53 were female (blue and pink silhouettes, respectively). NDD cases were recruited either as singletons (n=150) or as probands within families (n=61), including parent– offspring trios, duos and additional family structures. **(b)** Geographic distribution of 318 participants across 24 nationalities spanning the Middle East, North Africa, Africa, and Central/South Asia. Each country is identified by a unique color and/or fill pattern, as defined in the key, with the numbers of participants recruited from each country in parentheses. **(c)** Cooccurrence matrix of the 10 most frequent HPO-encoded phenotypes among the 211 NDD cases, each recorded in ≥5 individuals. The diagonal cells (gold) indicate the number of individuals carrying each term, and the off-diagonal cells (white-to-dark-green gradient) show the pairwise cooccurrence phenotype counts. Autism (n=125), delayed speech and language development (n=89) and ADHD (n=84) were the most frequent terms. The 20-phenotype display and full phenotype matrix are shown in Supplementary Figure 2 and Supplementary Table 2b, respectively. **(d)** Hex-bin density plot of the median HiFi read quality (Q) score against the read N50 across sequenced individuals. The white-to-dark-purple gradient represents the number of samples per bin. The samples clustered tightly at a median Q of 33–35 and N50 of 16–20 kb. **(e)** Unsupervised ADMIXTURE analysis of the NDD cohort cases together with global reference individuals from HGDP at K=2–6 inferred ancestral components (NDD cases shown in Supplementary Figure 5). Each vertical bar represents one individual, and the colored segments indicate the proportional contribution of each inferred ancestry component. Individuals are ordered by population, with HGDP regional groups to the left and the NDD cohort to the right (delimited by red dashed lines and labeled), subdivided by study supergroup. **(f)** Principal component projection of the NDD cohort cases onto the HGDP reference space. HGDP individuals are shown as triangles colored according to their geographic region, and NDD cases are shown as circles colored according to the study superpopulation groups, both of which are defined in the keys. The axis labels indicate the variance explained by each component (PC1 3.84%, PC2 3.22%). **(g)** Distribution of total ROH burden among NDD patients (n=211) and the combined HPRC_r2 and APR cohort (n=284). Individuals were grouped into five total ROH-length categories (<25 Mb, pale yellow; 25–50 Mb, pale pink; 50–150 Mb, gold; 150–300 Mb, brown; and >300 Mb, red). The values within the bars indicate the percentage of individuals within each category. **(h)** Total ROH length per individual in the three highest burden categories for the reference cohort (blue) and the NDD cases (red). The points represent individuals, and the boxes indicate the interquartile range. Above each category are the Fisher’s exact p value and odds ratio (one-tailed, NDD > APR + HPRC_r2) for the proportion of individuals falling in that category [50–150 Mb, p = 0.89, OR = 0.76 (ns); 150–300 Mb, p = 9.6 × 10□□, OR = 6.23 (***); >300 Mb, p = 1.2 × 10□³, OR = 14.08 (**)]. (**i)** Relationships between total ROH length and the longest individual ROH segment, restricted to individuals with >50 Mb total ROH. The circles denote comparisons of individuals, and the diamonds represent NDD cases. The point colors indicate the total ROH category. The vertical dashed lines mark 150 Mb and 300 Mb of total ROH. **(j)** Mean number of unique genes overlapping ROH per individual, stratified by GENCODE (v48) gene biotype (denoted by different colors). **(k)** Protein-coding gene counts within ROH, by total ROH length category, for the comparison cohort (blue) and NDD cases (red). The p values and odds ratios are the same as those in panel h [50–150 Mb, p = 0.76; OR = 1.01 (ns); 150– 300 Mb, OR = 9.82 (***); >300 Mb, OR = 30.08 (***)]. **(l)** Original parent classification of ROH in informative NDD trio probands (n=43). Each stacked bar represents one proband and shows the proportional contribution of ROH for the proband-specific (green), paternally inherited (blue), maternally inherited (purple), and shared by parents (yellow). The overlaid red line gives the total ROH length for each proband, which is plotted against the right y-axis.

Clinical phenotype harmonization across the 211 NDD cases revealed 267 unique Human Phenotype Ontology (HPO) terms. Autism was the most frequent phenotype (59.2%, n = 125), followed by delayed speech and language development (42.2%, n = 89), attention deficit hyperactivity disorder (ADHD) (39.8%, n = 84), global developmental delay (25.1%, n = 53), intellectual disability (21.8%, n = 46), hyperactivity (18.9%, n = 40) and seizures (15.2%, n = 32) (Figure 1c; Supplementary Figure 2; Supplementary Table 2a). Autism frequently cooccurred with delayed speech and language development (n = 62, 49.6% of autism cases) and ADHD (n = 53, 42.4% of autism cases), whereas global developmental delay (GDD) occurred more often alongside other neurological or dysmorphic features, including seizures, abnormal facial shape and hypotonia (Supplementary Table 2b).

PacBio HiFi sequencing generated consistently high-quality long reads across the cohort, with a mean read N50 of 18.0 kb (range 11.0–23.5 kb), a median read quality score of 34 (range Q31–39), and a mean genome coverage of 30.1x (Figure 1d; Supplementary Figure 3a–b; Supplementary Table 3a). Chromosome-level coverage against T2T-CHM13 exceeded 98% for most autosomes and chromosome X, with the expected variability across centromeric, pericentromeric and rDNA regions (Supplementary Figure 4a–b; Supplementary Table 3b–c).

Read-based DeepVariant (v1.9) calls used the linear reference GRCh38 and identified an average of 4,630,018 SNVs/sample (range 4,282,840–5,354,235) and 1,003,013 INDELs/sample (range 926,960–1,154,610) (Supplementary Table 4). Population structure analysis confirmed the broad genomic diversity suggested by geographic recruitment of the cohort. Unsupervised ADMIXTURE analysis across K = 6 (with lowest error) demonstrated heterogeneous ancestry components within the NDD cohort, with substantial Middle Eastern, Central/South Asian representation and African and North African components in a subset of individuals (Figure 1e; Supplementary Figure 5). Consistently, projections onto principal components derived from 929 Human Genome Diversity Project (HGDP) reference individuals placed NDD samples across Middle Eastern, North African, Central/South Asian and African-associated reference clusters (Figure 1f; Supplementary Figure 6). Y chromosome haplogroup distributions supported diverse paternal lineages across the recruited populations (Supplementary Figure 7; Supplementary Table 5a–c).

### NDD cases harbor excess long-range autozygosity

Given the demographic background of the recruited populations, we next examined runs of homozygosity (ROH) as a measure of genome-wide autozygosity. The median total ROH was broadly comparable across groups, with 29.4 Mb (range 18.3–101.1) for the Middle East, 29.2 Mb (range 26.8–33.5) for North Africa, 24.6 Mb (range 17.9–57.9) for Central/South Asia and 19.9 Mb (range 19.0–47.2) for Africa, but the distribution of burden across categories differed significantly (p = 0.036). North African cases were concentrated in the 25–50 Mb category (9/12, 75.0%) relative to Middle Eastern cases (37/170, 21.8%; Fisher’s exact test, BH-adjusted p = 1.3 × 10□³, OR = 10.8, 95% CI 2.8–41.9).

The upper tail of the ROH distribution differed markedly between NDD cases and population controls. Twenty-five NDD patients (11.8%) carried 150–300 Mb of total ROH compared with 2.1% of the controls (OR=6.23, one-sided Fisher’s exact test, p=9.6×10□□), whereas 10 cases (4.7%) carried >300 Mb of the total ROH, compared with 0.4% of the controls (OR=14.08; p=1.2×10□³; Figure 1g–h). In contrast, the lower ROH burden categories showed substantially smaller cohort differences. Ten NDD cases had a total ROH burden >300 Mb (range 300.27–675.67 Mb) (Figure 1i; Supplementary Table 6a).

NDD cases carried a greater mean (3304 genes) number of ROH-overlapping genes than population controls did (2116 genes), with protein-coding genes and lncRNAs accounting for the greatest absolute contributions (Figure 1j; Supplementary Figure 8). For each ROH burden stratum, we counted the unique genes within the ROH, and among the 25 cases carrying 150–300 Mb of the ROH, 81.3% of protein-coding genes (16,664/20,508) and 80.5% of lncRNA genes (28,080/34,882) were within an autozygous interval, compared to 30.6% (6,282) and 32.7% (11,423), respectively, in population reference controls. For >300 Mb of the ROH, the difference was greater between NDD cases and population controls, where 68.2% (13,983) of the genes were protein-coding genes and 69.3% (24,178) were lncRNA genes across ten NDD cases, compared with 6.7% (1,364) and 7.6% (2,636) for single-population controls (Supplementary Table 6b–c). Curated rare and NDD-associated gene sets were significantly enriched (p<0.0001) within the 150–300 and >300 Mb strata (Extended Figure 1; Supplementary Table 6d–e). In the trios with both parents sequenced, most of each proband’s autozygosity was not homozygous for either parent, and an average of 67.6% of the proband’s total ROH length fell within intervals where both parents were heterozygous (Figure 1l; Supplementary Table 6f; Supplementary File 1).

### Haplotype assemblies and pangenome from the NDD cohort

Haplotype assemblies built from the NDD cohort were highly contiguous, with a mean contig N50 of 54.6 Mb (hap1: 54.7 Mb; hap2: 54.5 Mb) (Figure 2a). HMM-Flagger profiling classified a mean of >98% of the assembled sequence as reliable (haploid), with erroneous, duplicated and collapsed assembly states accounting for 1.2% of the total sequence (Figure 2b; Supplementary Figure 9; Supplementary Table 7). The average assembly length was approximately 3.0 Gb per haplotype, and the consensus quality clustered around QV 61 (range 54.4–65.6) (Figure 2c; Supplementary Figure 10a–b; Supplementary Table 8). Liftoff v1.6.3 of GENCODE (v48) annotations recovered a mean of 98.4% (hap1) and 98.0% (hap2) of protein-coding genes per haplotype (Supplementary Figure 10c; Supplementary Table 9). The downstream analysis space was confined to high-confidence ‘easy regions’ (see Methods). A population-scale pangenome graph was constructed from the 636 NDD cohort haplotypes using the Minigraph-Cactus pipeline^25^ (cactus-pangenome) with T2T-CHM13 and GRCh38 as dual reference coordinates. The resulting graph comprises 149,222,686 nodes and 206,562,335 edges, with a mean edge-to-node ratio of 1.38 against the baseline ratio of 1.0 expected for a linear sequence, indicating that most nodes lie along uninterrupted haplotype paths with branching confined to variant sites. Growth curve analysis using Panacus^26^ quantified across 636 NDD assembly haplotypes recovered 567.8 Mb of nonreference sequences in total (Figure 2d; Supplementary Table 10) (details in the Supplementary Results).

**Figure 2.**
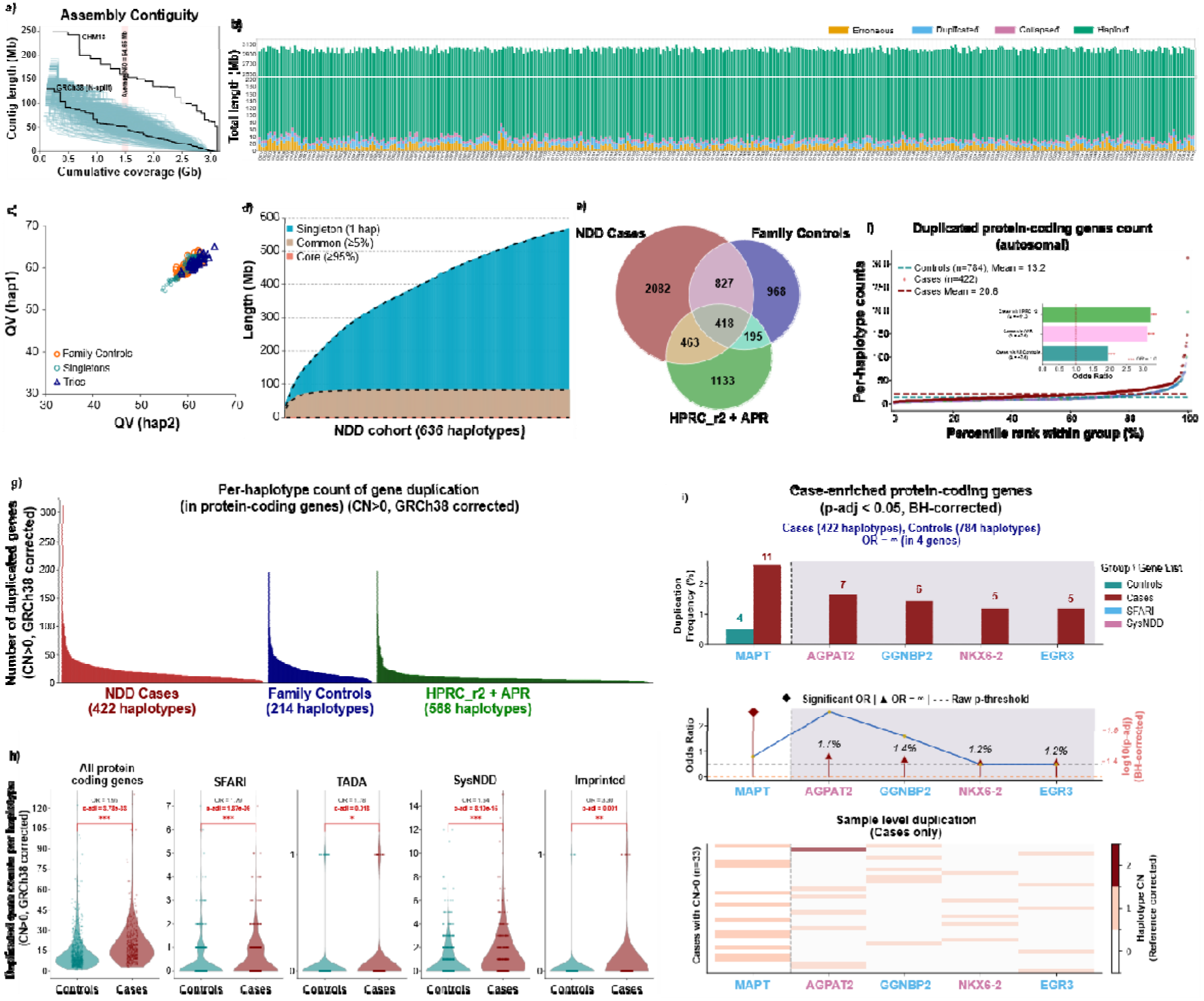
Haplotype assemblies and gene duplication landscape across the study cohorts. **(a)** Contiguity profiles of the NDD cohort haplotype assemblies. Individually assembled haplotypes are shown as pale blue curves, with the contig length plotted against the cumulative assembled sequence. T2T-CHM13 and gap-split GRCh38 are shown as black reference curves. The yellow vertical band indicates the cohort mean assembly N50 of 54.65 Mb. **(b)** HMM-Flagger classification of NDD case haplotype assemblies (422 haplotypes), partitioned into haploid (green), erroneous (dark yellow), duplicated (blue), and collapsed (pink) states. Each vertical bar represents one haplotype assembly. The haploid sequence accounts for ∼2.9–3.1 Gb per haplotype (see Supplementary Figure 9 for family controls). **(c)** Consensus QV of the two haplotype assemblies from each individual plotted against each other. The family controls are shown as orange circles, the singletons are shown as red open circles, and the trio-mode assemblies are shown as light blue triangles. The QV values cluster tightly between 55 and 65 for both haplotypes. **(d)** Pangenome growth curve showing cumulative nonreference sequences as NDD haplotype assemblies are added (636 haplotypes). Blue denotes sequences observed in a single haplotype, brown denotes common sequences present in at least 5% of the haplotypes, and red denotes core sequences present in at least 95% of the haplotypes. The black dashed lines trace the cumulative total of each fraction. **(e)** Venn diagram of duplicated genes detected in the NDD case, family-control and combined HPRC_r2-APR haplotypes (red, blue and green, respectively; blended colors indicate intersections). The numbers indicate the genes unique to each group or shared between groups. **(f)** Ranked distribution of reference-corrected duplicated protein-coding autosomal gene counts per haplotype, with haplotypes ordered by percentile rank within their own group. NDD cases are shown in red, family controls in blue, APR in green and HPRCR_r2 in purple. The red and gray dashed horizontal lines indicate the mean duplication burden in the cases (20.6 genes per haplotype, n=422) and in all the controls combined (13.2, n=782). The two distributions overlap through the lower percentiles and separate in the upper tail. The inset shows odds ratios for case control comparisons (all significant, *** indicates p<0.0001), with the dashed line indicating OR=1. **(g)** Per-haplotype gene duplication burden sorted in descending order within each group [NDD cases (red, n=422 haplotypes), NDD family controls (blue, n = 214) and the combined APR + HPRC panel (green, n=570)]. Each bar represents one haplotype, indicating the number of protein-coding genes with a reference-corrected haplotype copy number of at least 1 additional copy (CN ≥ 1). The distributions are strongly right skewed in every group, with a small number of high-burden haplotypes in each; the extreme is an NDD case haplotype carrying 311 duplicated genes, against group maxima of 196 in family controls and 198 in APR + HPRC_r2. The bulk of the haplotypes in each group fall below ∼30 duplicated genes, with the case distribution shifted upward relative to that in both control sets. **(h)** Distribution of duplicated-gene counts per haplotype in controls (blue, 782 haplotypes) and NDD cases (red, 422 haplotypes) for all protein-coding genes and the SFARI, TADA, SysNDD and imprinted-gene sets. Violins show the density of counts with individual haplotypes overlaid as points. Odds ratios and Benjamini–Hochberg (BH)-adjusted p values are shown above each comparison. **(i)** Protein-coding genes significantly enriched for duplication in NDD case haplotypes after genome-wide BH correction (p_adj < 0.05) that also belong to curated NDD gene sets (labels colored by gene set, blue for SFARI and pink for SysNDD). *Upper panel*, frequency of haplotypes carrying a duplication in controls (blue) and cases (red), with the carrier count above each bar. *Middle panel*, effect size per gene; the red diamonds indicate significant finite ORs, and the red triangles indicate genes duplicated only in cases (OR=∞, gray shaded block, four genes), with the BH-adjusted significance plotted on the right axis and the dashed line at OR=1. *Lower panel*, reference-corrected haplotype copy number for the 33 cases carrying at least one of these duplications; white indicates no additional copy, and progressively darker red indicates higher copy number. *MAPT* is the only one of the five genes that is also duplicated in the controls (11 case haplotypes versus 4 control haplotypes). *AGPAT2*, *GGNBP2*, *NKX6-2* and *EGR3* are duplicated exclusively in cases.

### Gene duplication diversity in NDD haplotypes

We next used the haplotype assemblies to examine gene-copy diversity across the study populations. Gene duplication was widespread, with 415 duplicated protein-coding genes being shared across NDD cases, family controls and HPRC_r2-APR (segmental duplications filtered; see Methods). Each population group in the cases carried a substantial private component, consistent with local genomic diversity, with a mean of 21 protein-coding genes in the Middle East, 15 in North Africa, 15 in Africa, and 22 in South and Central Asia. The duplication burden was distributed across ancestry groups rather than being restricted to a single recruitment population (Supplementary Figure 11). We found 2,082 unique duplicated genes only from NDD cases, compared with 968 among family controls and 1,133 in HPRC_r2-APR (Figure 2e; Supplementary Table 11a). After correcting for population background, compared with the controls, NDD cases presented a greater duplication burden, with the affected haplotypes carrying a mean of 20.6 duplicated protein-coding autosomal genes (OR = 1.367, p < 0.0001) (Figure 2f; Supplementary Table 11b). Protein-coding gene duplications (CN ≥ 1) were strongly right skewed in each group (NDD cases, family controls and HPRC_r2-APR) (Figure 2g). Most haplotypes (∼90%) carried fewer than 30 duplicated genes, but a small tail of high-burden haplotypes was present in all three groups, reaching 312 genes in an NDD case versus 195 and 197 in family controls and APR-HPRC_r2, respectively (Supplementary Table 11c). At the gene-set level, compared with population controls, NDD cases presented a greater duplication burden for protein coding (OR = 1.95). Similar associations were observed for SFARI, TADA, SysNDD and imprinted-gene sets (Figure 2h; Supplementary Table 11d). Genome-wide analysis revealed five protein-coding genes whose duplication frequency was significantly greater among the NDD haplotypes (Figure 2i). *MAPT* duplications were present at a low frequency in the controls but were significantly (p < 0.05, OR = 5.1) enriched in the cases, whereas duplications of *AGPAT2*, *GGNBP2*, *NKX6-2* and *EGR3* were found only in the NDD-affected haplotypes (Supplementary Table 11e). The non-coding gene duplications are shown in Supplementary Figure 12.

### Small pathogenic variants resolve diverse NDD cases

Small variants identified across 211 NDD-affected cases were annotated and classified using two independent tertiary interpretation tools, HORIZON^27^ and ANNOVAR^28^, to assess the consistency of the pathogenicity classification and were reviewed by clinical geneticists according to the ACMG/AMP criteria. Among the 211 NDD cases, 56 had pathogenic (P) or likely pathogenic (LP) findings, yielding an overall molecular resolution rate of 26.54% (56/211) (Figure 3a). Recessive variants accounted for the largest share, and 15 cases (7.11%) were resolved by biallelic variation. This comprises five cases (2.37%) with compound heterozygous variants in the *FBN3*, *NPC1*, *CPA6*, *CYP27A1* and *CC2D2A* (Figure 3b) genes; five cases (2.37%) with autosomal recessive homozygous variants in the *ASL*, *GRIN1*, *KMT2D*, *TRAPPC11* and *PDE2A* genes; and five cases (2.37%) with hemizygous or X-linked homozygous variants in the *HS6ST2*, *AFF2*, *DOCK11*, *GDI1* and *GRIA3* genes. Three additional patients (1.42%) carried heterozygous X-linked variants in *CASK*, *HUWE1* and *SLITRK2*, resulting in eight X-linked findings in total (3.79%). Four cases were molecularly resolved by clinically relevant *de novo* variants, a frameshift deletion in *NRXN2* and *de novo* missense variants in *SPTAN1*, *GRIN2B* and *HECW2* (Supplementary Table 12). The remaining resolved cases carried heterozygous loss-of-function (LoF) variants, including five cases with stop-gain variants affecting the *LMBRD2, TNRC6B*, *SAMD9*, *SCN11A*, and *COL4A2* genes; nine cases with splice-site variants impacting the *GABRB3, SYT1, NAA15, NTRK2, TRIO, CCM2, CDC42BPB, TNRC6A,* and *KDM4B* genes; and six cases with frameshift variants disrupting the reading frame in the *FBXO11, AHDC1, NF1, MYT1L, CAMTA1,* and *ANKRD17* genes. Three samples carried two candidate variants each. All reported variants were confirmed by manual review of IGV alignments and orthogonally validated by Sanger sequencing (Supplementary File 2).

**Figure 3:**
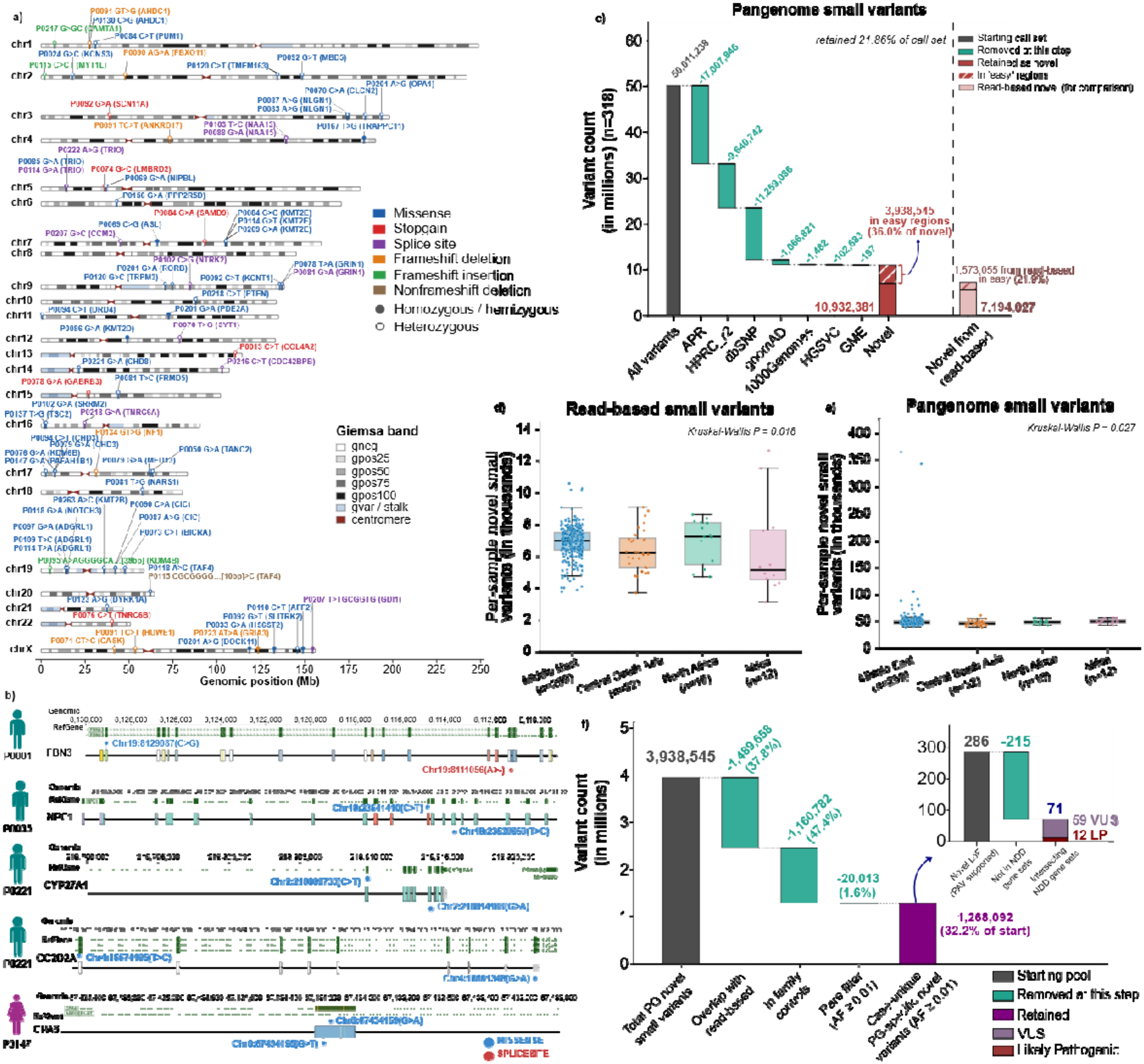
Resolving causal and novel population-specific small variants from read-based and pangenome call sets. **(a)** Ideogram showing the cases with ACMG-guided pathogenic/likely pathogenic variants, plotted at their genomic position across chromosomes 1-22 and X. Each variant is labeled with the respective sample ID, gene symbol, base change and coordinate and colored by sequence (blue, missense; purple, splice site; red, stopgain; green, frameshift insertion; orange, frameshift deletion; brown, nonframeshift deletion). The filled circles denote homozygous or hemizygous individuals, and the open circles denote heterozygous carriers. **(b)** Compound heterozygous variants in NDD candidate genes. The top track shows the RefGene model, and the two tracks below are individual alleles (AL1 and AL2), with the exons drawn to scale. The two variants in each gene are marked on the haplotype carrying them, with coordinates and base changes. In every case, the two variants are *trans*, confirming true compound heterozygosity directly from the phased variant calls. The silhouettes give the sex of each carrier (blue, male; pink, female). **(c, d)** Per-sample counts of population-novel small variants, stratified by superpopulation, for the read-based (c) and pangenome-based (d) call sets. The boxes indicate the median and interquartile range, with the whiskers extending to 1.5 × IQR. Each point represents one sample that is horizontally jittered. The superpopulation sizes are given on the x-axis. Kruskal Wallis tests across all four superpopulations revealed that p < 0.05 (Supplementary Table 13j). **(e)** Waterfall plot showing the sequential database filtering of the pangenome small-variant call set (n = 318 individuals). Starting from 50,011,238 variants (gray), the teal bars indicate the variants removed at each successive database step. The filtering order was APR, HPRC r2, dbSNP, gnomAD, 1000 Genomes, HGSVC and GME, resulting in 10,932,381 novel variants (21.86% of the call set) (dark red), of which 3,938,545 (36.0%) were in high-confidence “easy” regions (hatched). The pale pink bar gives the equivalent novel set from read-based calling for comparison (7,194,027 variants; 1,573,055 easy, 21.9%). **(f)** Case-unique filtering cascade applied to the 3,938,545 pangenome-novel variants at each step, with read-based overlap (1,489,658; 37.8%) in family controls (1,160,782; 47.4%) and rare AF ≥ 0.01 (20,013; 1.6%), resulting in 1,268,092 case-unique pangenome-specific rare novel variants (red; 32.2% of the starting pool). The inset shows 286 novel loss-of-function (LoF) variants; 71 intersect with curated NDD gene sets, comprising 59 variants of uncertain significance (VUS) and 12 classified as likely pathogenic (LP) using HORIZON annotation.

### Pangenome-specific ultrarare small variants

The pangenome calls revealed 50,011,238 distinct small variants across the 318 cohort genomes. After sequential filtering against the APR^13^, HPRC_r2^24^, dbSNP^29^, gnomAD^30^, 1KGP^31^, HGSVC^32^ and GME^33^ databases, 10,932,381 variants (21.9%) were absent from all seven resources (Figure 3c; Supplementary Table 13a–b). The read-based call sets retained 7,194,027 novel variants (14.2%), so graph-based calling recovered 1.52-fold more population-novel variation from the same genomes (Supplementary Figure 13; Supplementary Table 13c). Restricting to ‘easy regions’ resulted in 3,938,545 pangenome-novel variants (36.0% of the novel set) and 1,573,055 read-based novel variants (21.9%) (Supplementary Table 13d–e). Novel small variants grouped across superpopulations differed, with the highest median in African (48,984) and lowest in Central/South Asian (45,465) genomes for both call sets (read-based, p = 0.018; pangenome-based, p = 0.027). The African-ancestry genomes presented the highest median number of novel variants in any group but the lowest proportion of samples (Figure 3d–e; Supplementary Table 13f–i). To identify case-unique pangenome-specific novel variants, novel variants overlapping with read-based calls, family controls and those with a cohort AF ≥ 0.01 were filtered, retaining 1,268,092 case-unique, pangenome-specific rare variants (Figure 3f). Annotation of this set returned 286 PAV-supported LoF variants (splice-site, start-loss and stop-gain). Frameshift events were disregarded as an expected signature of residual assembly error within a homopolymer run or short tandem repeat or a position shift on the left alignment^34^. Among the 286 variants, 71 intersect with the curated NDD gene sets. HORIZON^27^ annotation predicted 59 VUS and 12 LP (Figure 3f, inset; Supplementary Table 13k).

### Pathogenic structural variants (SVs) landscape

SV calls from HiFi reads aligned to GRCh38 using sawfish^35^ (v2.1.1) yielded 41 pathogenic or LP SVs across 35 (16.58%) cases (Supplementary Table 14). The set comprised 31 deletions, nine duplications and one inversion, 37 of which were heterozygous and 4 of which were homozygous or hemizygous SVs. The SV sizes ranged from 57 bp to 5.38 Mb (median 227 kb); 13 events were ≥1 Mb, whereas 6 were <1 kb. Prioritized SVs were distributed across 17 chromosomes and 29 cytobands (Figure 4a). Among the 35 cases with a prioritized SV, nine also carried a P/LP small variant, and 26 (12.32%) of these cases were resolved through SVs. Five cases carried more than one prioritized SV. The X chromosome carried the greatest burden, with 12 SVs across 11 cases. Twenty-three SV events were clustered at nine loci recurrently hit by two or more unrelated individuals. These included 22q11.21 (chr22:18.76-21.33 Mb with two deletions and one 990 kb inversion), the 15q11-q13 interval (chr15:23.13-28.47 Mb with one deletion and one duplication), chr16:21.77-22.45 Mb (1 deletion and 1 duplication), 9q34.3 spanning *GRIN1*-*EHMT1*, and Xq22.3, where three individuals carried deletions of *TBC1D8B*, with two of them having an identical 57 bp homozygous or hemizygous event. Recurrence of both variants in the same locus at Xq22.3 in unrelated individuals is consistent with these intervals, which represent genuine NDD-associated loci rather than sporadic findings. Candidate SVs were reviewed in IGV and orthogonally validated by Oxford Nanopore sequencing.

**Figure 4:**
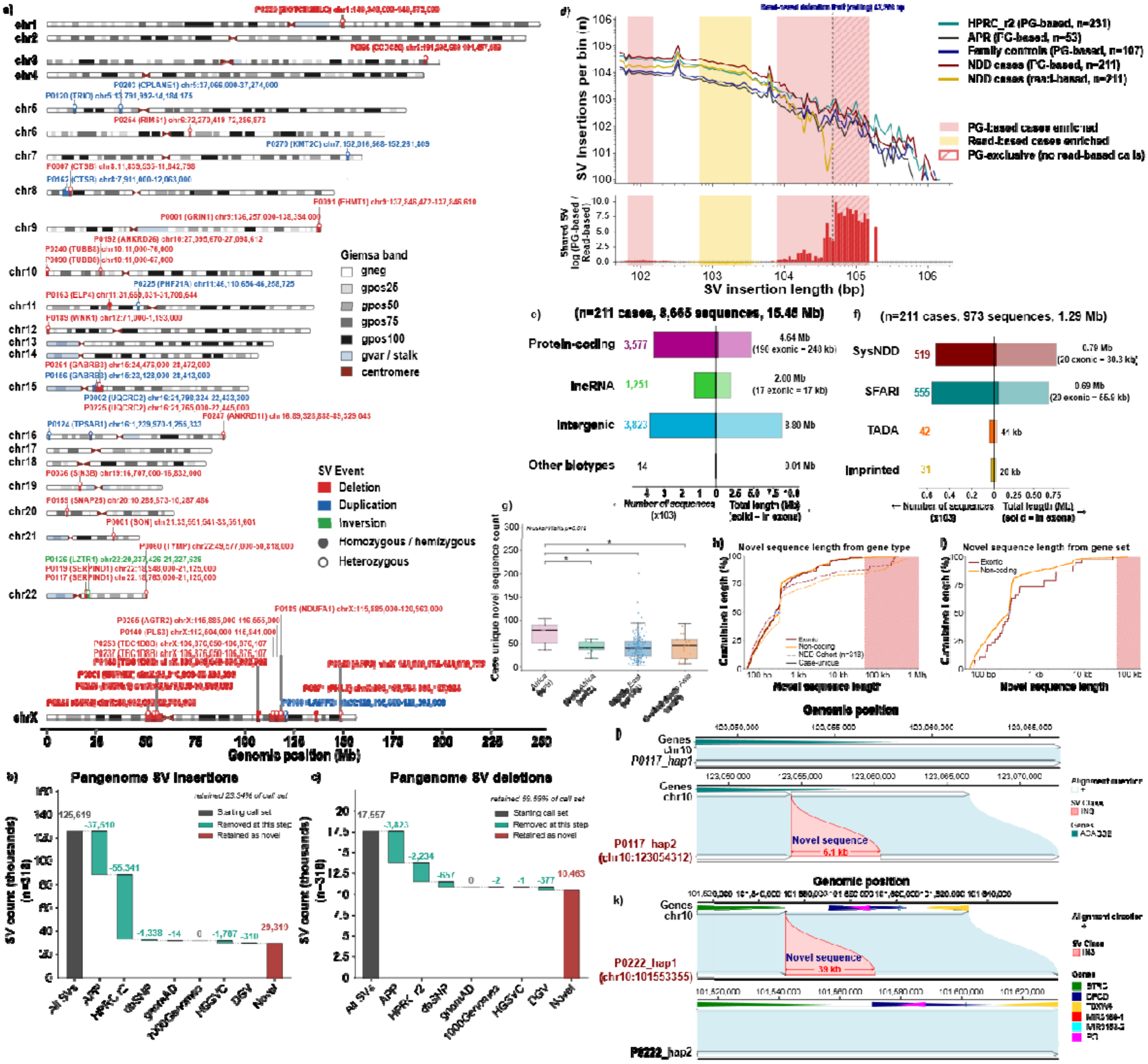
**(a)** Genome-wide distribution of prioritized P/LP SVs. Each of the 41 prioritized SVs is marked on the chromosome at its called interval and annotated above with the sample and coordinates (chr:start-end, GRCh38). The marker and label colors denote the variant class: red for deletion, blue for duplication and green for inversion. The circle immediately above each site denotes zygosity and is filled for homozygosity or hemizygosity and unfilled for heterozygosity. SVs smaller than 1.1 Mb are displayed at a fixed minimum width, and marker width reflects actual SV length only for variants exceeding this threshold. When several variants occupy overlapping intervals, the markers are superimposed and distinguished by their annotations. **(b, c)** Sequential filtering of pangenome-derived SV insertions against reference panels. Waterfall plots showing the attrition of SV insertions and deletions (c) across the cohort (n = 318) as calls were matched, in a fixed order, against APR R1, HPRC release 2, dbSNP, gnomAD, 1000 Genomes, HGSVC and DGV. The gray bars represent the starting call set (125,619 insertions; 17,557 deletions), the teal bars represent the number of SVs removed at each step, and the red bars represent the variants retained as population novel after all seven databases (29,319 insertions, 23.34% of the call set; 10,463 deletions, 59.59%). The filtering is sequential and nonredundant, and counts are plotted in thousands. **(d)** SV insertion length spectra differ systematically between pangenome-based and read-based calling in the same individuals. The number of SV insertions per length (log-scaled) bin for five call sets, HPRC_r2 (n = 231), APR (n = 53) and family controls (n=107), and all pangenome-derived and NDD cases (n = 211) called both from the pangenome and from reads. Bins are of equal width in log_10_ space (25 bins per decade) over 50 bp to 2 Mb, and both axes are logarithmic. Because the PG-based case call set contains 1.95× more calls in total than the read-based call set does, an identical length distribution would place the red curve 1.95× above the solid yellow curve in every bin. The dashed curve is therefore the correct visual null: it is the read-based profile rescaled by 1.95, and PG-based enrichment corresponds to the red curve lying *above the dashed curve*, not above the solid yellow curve. The vertical dashed line represents the read-based detection limit (47,356 bp). The bottom panel shows the log ratio of the yield-normalized frequency (calls per 10,000 calls of the same call set) relative to the read-based case profile. The direction of the bars represents PG-based versus read-based for NDD cases in which the saturated bars pass both the effect-size threshold (|log2FC|≥0.15, dashed horizontal lines) and an FDR < 0.05, and the pale bars fail the effect-size threshold. Shading in both panels indicates contiguous runs of at least two bins meeting both criteria; red indicates enriched PG-based cases (66–151 bp and 7,942–151,540 bp), and yellow indicates enriched read-based cases (660–3,466 bp). Hatching marks the region beyond the read-based ceiling, where the read-based count is exactly zero and the ratio reflects the absence of detection rather than a graded difference. The per-bin counts, frequencies, effect sizes, test statistics and shading calls are listed in Supplementary Table 15d. **(e)** Gene biotype distribution of novel sequences in NDD cases (Supplementary Figure 14 for the NDD cohort, n = 318). The dividing bar represents 8,665 novel sequences totaling 15.45 Mb identified in the cases. The two sides of the axis carry different units and independent scales, so the bar widths are not comparable across the midline. The left side of the y-axis shows the number of sequences per biotype (×10³), and the right side shows the total length (in Mb), with the portion falling in annotated exons (drawn solid and annotated in brackets). **(f)** Case-unique novel sequences intersecting the NDD-associated and imprinted gene sets. Diverging bar chart of the 973 distinct novel sequences (1.29 Mb) in 211 NDD cases that fall within genes from four curated sets. The genes among the sets overlap; thus, the per-set values sum to more than the stated totals. The axis units are the same as those in panel e. **(g)** Case-unique novel sequences grouped by superpopulation. The boxes show the median and interquartile range, whiskers extend to 1.5 × IQR, and individual samples are overlaid as points. Groups were compared by the Kruskal Wallis test (p = 0.019), and the brackets show pairwise Mann Whitney comparisons with BH correction (* q<0.05). **(h)** Cumulative distribution of novel sequence length (log scale) for exonic (red) and noncoding (orange) sequences, shown for the full NDD cohort (dashed, n = 318) and for the case-unique subset (solid, 8,665 sequences). The hatched region indicates lengths above the 48 kb read-based SV detection limit in this cohort (sequences in this range are recovered only by pangenome analysis). **(i)** Cumulative length (log scale) distribution of the 973 novel sequences intersecting the gene sets, split into exonic (red) and noncoding (orange) sequences. None of the novel sequences found in the exonic region for the intersecting gene sets exceeded 48 kb. **(j, k)** Representative example of a case-unique novel insertion at the ACADSB locus in P0117 (j) and BTRC loci in sample P0222 (j) against GRCh38. The black bars represent the reference (top of each subpanel) and the query haplotype contig (bottom), the sky-blue ribbons show collinear alignment blocks in the forward orientation, and the red ribbon marks an unaligned heterozygous insertion of 6.1 kb in ACADSB gene (j) and 39 kb in BTRC gene (k) (both in the exonic region). Gene models are drawn above the reference.

### Pangenome-specific discovery of SVs in NDD cases

Pangenome SV calling across the NDD cohort yielded 125,619 insertions and 17,557 deletions (7.5:1), an insertion bias consistent with long-read assembly-based discovery. Sequential filtering against reference databases (see Methods) retained 29,319 insertions (23.3%) and 10,463 deletions (59.6%) as novel (Figure 4b–c; Supplementary Table 15a–b). Annotation of the novel pangenome-specific SV set and removal of calls also observed in family controls revealed six overlapping curated NDD genes in distinct cases: *GRIN2A* (781 bp, removing 147 bp of a coding exon together with the adjacent splice site), *ARID1B* (322 bp), *RNF135* (160 bp), *BRSK2* (128 bp), *SYN1* (86 bp, X-linked) and *ANK2* (61 bp) (Supplementary Table 15c). Five of these were heterozygous, and the *SYN1* call was hemizygous. These prioritized calls did not overlap with a previously reported disorder-associated interval.

We also compared the length distribution of all insertion SVs called from the pangenome graph (PG-based) with those called from the HiFi reads in the same 211 NDD-affected individuals (Figure 4d; Supplementary Table 15d). The two call sets differ almost twofold in terms of the total detection rate (1,559,528 in PG-based versus 799,379 in read-based insertion calls; ratio 1.951), the raw per-bin counts are not directly comparable, and each bin was tested on each call set share of its own total. Insertion lengths span dynamically, so a single fixed bin width cannot resolve both short and long insertions; therefore, bins were widened in proportion to length, each approximately 10% wider than the previous one, yielding 115 bins across 50 bp to 2 Mb. Of these, 88 contained enough calls to test. Although 83 reached an FDR < 0.05, this reflects the very large number of pooled calls rather than large differences between the call sets, so enrichment was called on effect size (|log_2_FC| ≥ 0.15, a ≥1.11-fold shift in share) with BH-adjusted significance. Insertions of 66–151 bp were overrepresented in the PG-based call set (maximum log_2_FC = 0.23; 335,457 PG-based calls versus 151,617 read-based calls), whereas insertions of the 660–3,466 bp size range were overrepresented in the read-based call set (maximum log_2_FC = −0.31; 374,823 calls versus 225,011 calls). The greatest and most interpretable divergence was above 8 kb, with the percentage of PG-based calls exceeding the read-based share continuously from 7,942 to 151,540 bp (30,834 versus 8,607 calls), with the ratio increasing with increasing insertion length to a hard read-based detection ceiling at 47,356 bp, above which the pangenome retained 4,596 calls (0.29% of the PG-based call set), extending to ∼1.8 Mb. All three PG-based control panels (HPRC_r2, APR and family controls) shifted in the same direction as the PG-based cases in all three regions did, and more strongly (log_2_FC +0.34 to +0.48 at 66– 151 bp, −0.55 to −0.59 at 660–3,466 bp and +1.03 to +1.42 above 8 kb, against +0.18, −0.23 and +0.88 in cases), confirming that these differences are a property of the pangenome representation rather than a feature of the NDD cohort.

### Novel sequences in underrepresented populations missing from current references and pangenomes

Within high-confidence genomic regions (‘easy regions’), we identified 16,943 novel sequences across 636 NDD cohort haplotypes that were absent from the reference pangenomes (HPRC_r2 and APR), resulting in a total of 691.4 Mb (1.09 Mb per haplotype) of novel sequences. Intergenic sequences carried the most novel sequences across the NDD cohort (357.9 Mb, 51.8%), followed by protein-coding loci (187 Mb, 27.1%), lncRNAs (145 Mb, 21.0%) and other gene types (1 Mb, 0.1%). Among these, only 472 novel sequences (10.54 Mb, 1.5%) overlapped with the annotated exons of protein-coding genes (Supplementary Figure 14a; Supplementary Table 16a). Intracohort family controls novel sequence filtering retained 8,665 case-unique novel sequences totaling 15.45 Mb (73,222 bp per case) (Figure 4e; Supplementary Figure 14b; Supplementary Table 16b). Case-unique overlap with the curated NDD gene sets resulted in a total of 973 distinct sequences comprising 1.29 Mb. The genes among the curated sets overlap (one gene might be in different sets); therefore, the per-set values summed to more than the stated totals. Across the gene sets, 3.9% (38/973) of the distinct case-unique novel sequences were found in the exonic region, with a cumulative length of 85,870 bp (Figure 4f). Superpopulation stratification of the case-unique novel sequences revealed that the African origin samples carried significantly more novel sequences than the North African, Middle Eastern or Central/South Asian origin samples did (median 79 vs. 42, 41 and 46; Kruskal Wallis p = 0.019; all pairwise comparisons against North Africa BH-adjusted p = 0.02, Middle East p = 0.002, and Central/South Asia p = 0.019) (Figure 4g; Supplementary Table 16c). The cumulative novel sequence length distribution in the cohort (n = 318) was 15.2% (2,499/16,471) in the noncoding region and 11.6% (55/472) in the exonic region with a length >48 kb, whereas for the NDD cases, unique novel sequences accounted for 0.8% (66/8,475) of the noncoding novel sequence insertions >48 kb, and no novel exonic sequences above 48 kb were case unique (Figure 4h). The cumulative novel sequence length distribution of the case-unique sequences showed only 0.6% (6/935) of the novel sequences were in the noncoding region >48 kb, whereas none of the exonic case-unique novel sequences were found to be above the read-based SV detection limit (Figure 4i). The representative case-unique novel sequences in the exonic domain of the ACADSB and BTRC genes are shown in Figure 4j–k. The mode of inheritance for these genes is autosomal dominant (AD), indicating a clear association with NDD cases.

### Rare exon-impacting complex structural rearrangements in NDD cases

Complex rearrangements were identified on the basis of rare homozygous exons impacting SVs in NDD-associated genes. The frequency of each of the rare homozygous events reported was cross-checked with the APR and HPRC_r2 pangenome haplotype walks and is shown in the panels representing the patterns (all of these were heterozygous in the controls). Homozygous rare exonic complex rearrangement events were observed in a total of 45 SVs in 34 genes (4 SVs associated with NDD) occurring in 181 homozygotes in 170 families, but that total was dominated by common alleles (85 PRR36-, 73 PER3-, 71 OR8G1-, and 65 LNP1-affected homozygotes), and 25 of the 45 also had unaffected homozygotes (Supplementary Table 17a). Among the 45 read-supported homozygous exon-impacting SVs (34 loci), 27 had no homozygote in APR or HPRC_r2, and 20 had no unaffected homozygote in the cohort; 15 SVs in 13 genes met both conditions and were homozygous in 8.1% of the cases (17/211), however, only 2.4% of the cases (5/211) carried rare homozygous complex exonic rearrangements in the NDD associated genes. Representative loci from this set are shown in Figure 5. On chr14, a complex structural region window of 100 kb (Figure 5a) showed a rearrangement in the exon of the *AHNAK2* gene. The zoomed-in graph of the traversal walk associated with the variations in the haplotype is shown in the middle panel. A total of 62 different traversal patterns across the 636 haplotype traversal paths were observed, of which 4 patterns are shown. The gray highlighted segment shows the homozygous rare exonic event in the *AHNAK2* gene. A complex allele replacing 1,622 bp of the first coding exon with 616 bp of the exon paralog-derived sequence was homozygous in 2 affected individuals (both from the same family) and was observed as a heterozygous event in 19 other case haplotypes: 7 family control haplotypes, 3 APR haplotypes and 15 HPRC_r2 haplotypes. The same event pattern showed a homozygous tandem duplication of 495 bp in the same 2 cases (however, the control haplotypes were heterozygous). Another homozygous complex rearrangement was observed in the EPPK1 gene. The graph in Figure 5b shows the different traversal paths of the samples on chr8 (179 kb window). The zoomed section shows the structural rearrangements in the *EPPK1* gene. Among the 636 haplotypes, only 62 haplotypes traversed the same path as the reference (CHM13), and the locus carried 77 alternative traversal patterns. Three distinct complex allele patterns impacting the *EPPK1* exon were observed. One of the patterns (event A) showed a homozygous exonic deletion of 1,688 bp (in 1 case), which was observed as 93 other heterozygous haplotypes (66 case haplotypes and 27 control haplotypes). Another interesting pattern (event B) in a distinct case (n=1) showed a homozygous exonic deletion of 3,275 bp; however, 42 haplotypes displayed the same traversal pattern (24 cases and 18 controls, all heterozygous). A third alternative haplotype walk pattern (event C) in a distinct case was quite similar to the first pattern; however, it differed from the previous pattern by a tandem duplication of 86 bp downstream of the *EPPK1* gene.

**Figure 5:**
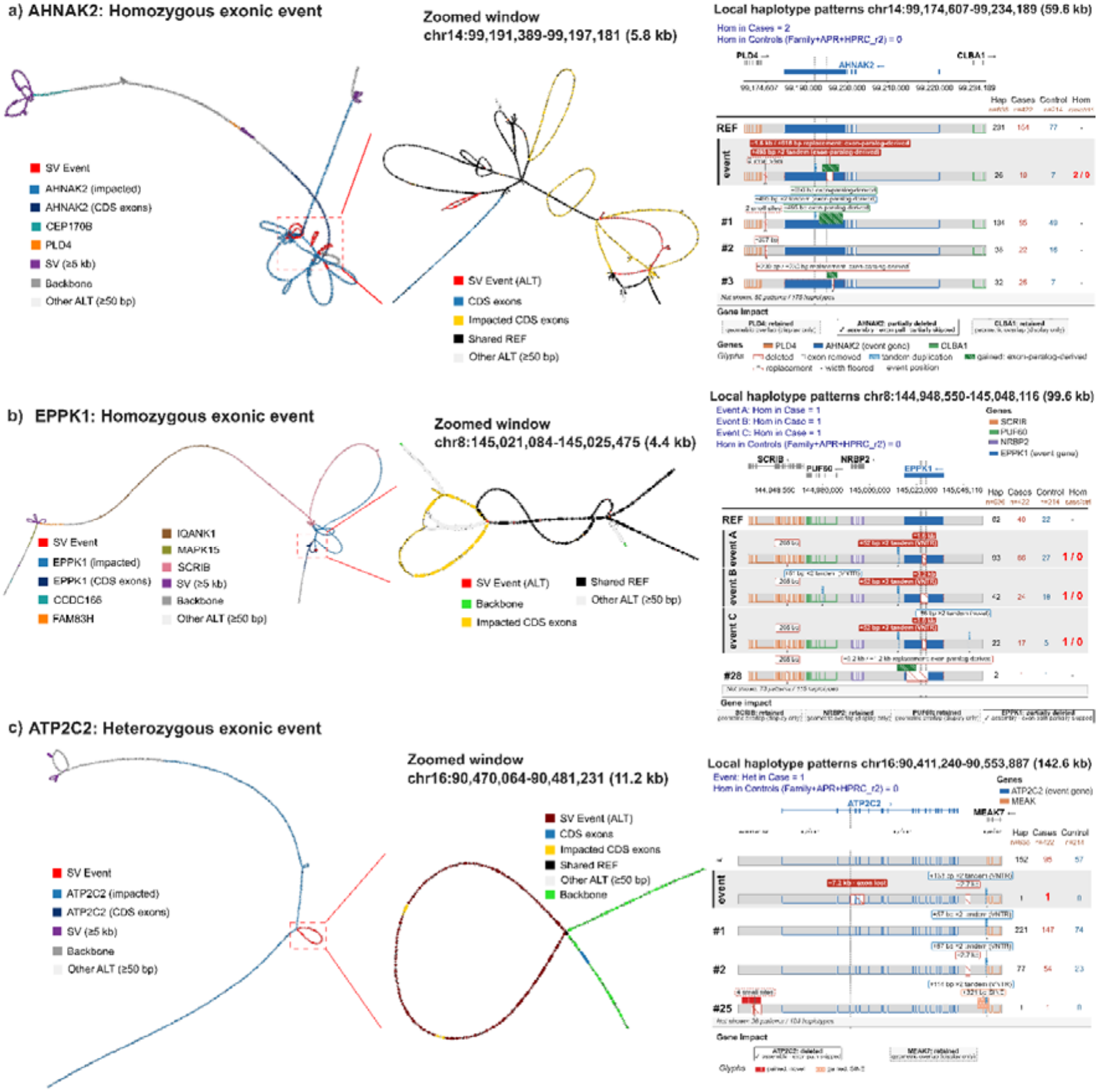
Rare exon-impacting complex structural alleles in NDD cases and their local haplotype context. Each subpanel (horizontal) shows one locus in three views; window, zoom and patterns with complex structural rearrangements. *Left*, the local pangenome graph across the whole window as a topology-preserving contracted graph. Nodes are colored by gene; the impacted genes are shown in light blue with their coding exons shown in dark blue, neighboring genes shown in other hues, intergenic reference backbones shown in gray, structural records ≥5 kb shown in purple, other nonreference nodes ≥50 bp shown in pale gray, and the structural part of the showcased event shown in red. The red box outlines the region expanded in the middle view. *Middle*, zoomed-in view of the event. Red represents alternative-allele nodes; dark red represents the reference interval deleted or replaced; black represents the reference nodes inside the record in which the carrier path also traverses; green represents the reference backbone; blue represents the coding exons of the impacted gene; yellow represents the exon nodes falling inside the event; and pale gray represents other nonreference nodes ≥50 bp. *Right*, local haplotype patterns over the locus, one row per pattern (the branch a haplotype takes through structural sites ≥50 bp), with the reference pattern first and the showcased event second and the gene model above and dashed lines indicating the event position. Glyphs annotate deleted, exon-removing, tandem-duplicated and gained sequences (exon-paralog-derived, novel or SINE), with the widths floored for visibility. The counts at right represent the number of haplotypes carrying each pattern (all, affected, unaffected; n = 636 haplotypes from 211 affected and 107 unaffected individuals), and for homozygous events, the number of individuals homozygous for the exact allele, other total patterns and remaining haplotypes are summarized in the footer row. The text above each panel refers to the exact allele, homozygous and total carrier counts, number of families, and status in the APR (106 haplotypes) and HPRC_r2 (462 haplotypes). The gene-impact boxes below give the assembly-supported consequence per gene, and “geometric overlap (display only)” indicates genes that overlap the window but are not altered by the event. Assembly support, where stated, denotes SV supported by PAV calls. **(a)** *AHNAK2* (chr14), inside a dense structural window. A complex homozygous allele (−1,622/+1,127 bp) replaces part of the first coding exon with an exon paralog-derived sequence, including a tandem-duplicated segment; *AHNAK2* is partially deleted with the exon path partially skipped, whereas *PLD4* and *CLBA1* are retained. It is homozygous in 2 affected individuals from one family and carried in total by 14 affected and 6 unaffected individuals across 15 families; it is not a homozygote in APR or HPRC, with the exact allele on 3 APR and 15 HPRC haplotypes. **(b)** *EPPK1* (chr8), inside a dense structural window. Three complex alleles in a single record (A, −1,688/+179 bp; B, −3,275/+161 bp; C, −1,688/+173 bp), each combining a 205 bp deletion, VNTR tandem duplications and a 1.6–3.2 kb deletion removing coding sequence; each is homozygous in one affected individual (allele A is additionally heterozygous in 3 affected individuals from 2 further families; alleles B and C are Mendelian inconsistent within their families, and allele B is also carried by one unaffected individual). *EPPK1* is partially deleted with the exon path partially skipped, while *SCRIB*, *PUF60* and *NRBP2* are retained. APR or HPRC have no homozygotes, allele A is carried on 2 APR haplotypes, and alleles B and C are carried on no haplotypes. The whole window extends to the two nearest records ≥5 kb (insertions 81 and 97 kb upstream, purple). **(c)** *ATP2C2* (chr16, SysNDD gene), a heterozygous case-only allele inside a dense structural window, a 7,167 bp deletion that removes coding exons in one affected individual from one family, carried on a single cohort haplotype and absent from APR and HPRC. The event haplotype additionally carries a VNTR tandem duplication and a 2.7 kb deletion.

We further explored the heterozygous rare exonic events with stringent criteria such that no other control haplotype (including APR and HPRC_r2) walk would have the same traversal path. Filtering resulted in a total of 55 core heterozygous case-only SVs identified from the initial set of 261 SV loci which consisted of 134 genes (Supplementary Table 17b). As shown in Figure 5c, a rare heterozygous exonic deletion of 7.2 kb was observed in the *ATP2C2* gene, which was traversed by only 1 case haplotype. Similarly, several other such case-only heterozygous events were observed for the *RRBP1*, *PTGER3*, *ZMYM6* and *CLEC16A* genes, as shown in Extended Figure 2.

### Pathogenic tandem repeat expansions (TREs) across clinically relevant loci

Our analysis of the 68 clinically annotated TRE pathogenic loci in the Platinum catalog revealed five distinct loci across 8 (3.8% yield for expansion detection rate) unique individual cases, out of which 10 samples were unresolved through ACMG-guided variant interpretation. The expansions span genes with autosomal dominant (AD, 50.0%), X-linked dominant (XLD, 37.5%), and autosomal recessive (AR, 12.5%) inheritance modes (Figure 6a; Supplementary Table 18a). Expansions at the *FMR1* locus (fragile X syndrome, CGG repeat, and pathogenic threshold >200 repeats) were identified in three probands. One proband carried a full mutation allele of ∼880 CGG repeats, whereas both parents presented normal or premutation-range alleles, suggesting the presence of a maternally transmitted premutation that expanded to full mutation in the proband, a well-established mechanism in fragile X syndrome. Additionally, two siblings with NDDs carried large CGG expansions in FMR1 of 1,844 and 1,950 bp, respectively (Figure 6b). Recurrent pathogenic CAG repeats in *FECD3_TCF4* (AD, threshold >51 repeats) were observed in 2 singleton cases with one of the alleles being approximately 250–275 bp in length (∼85–90 repeats) (Figure 6c). One singleton showing pathogenic CAG repeats (1,327 bp, 409 repeats) was identified in *DM1_DMPK* (AD, threshold >50 repeats), with a single CAA interruption at approximately 1,000 bp from the start of the repeat) (Figure 6d). Similarly, a singleton case showed expansion in *EPM1_CSTB* (CGCGGGGCGGGG, AR; both haplotypes were in the pathogenic range of 456 and 478 bp), which is consistent with biallelic expansion and autosomal recessive Unverricht–Lundborg disease (Figure 6e). One singletons sample carried pathogenic expansions in *DIP2B* (GGC, AD, 1,365 bp with scattered single-G interruptions) (Figure 6f). The pathogenic expansions were orthogonally validated from ONT-LSK runs and visualized through IGV (Supplementary File 3).

**Figure 6:**
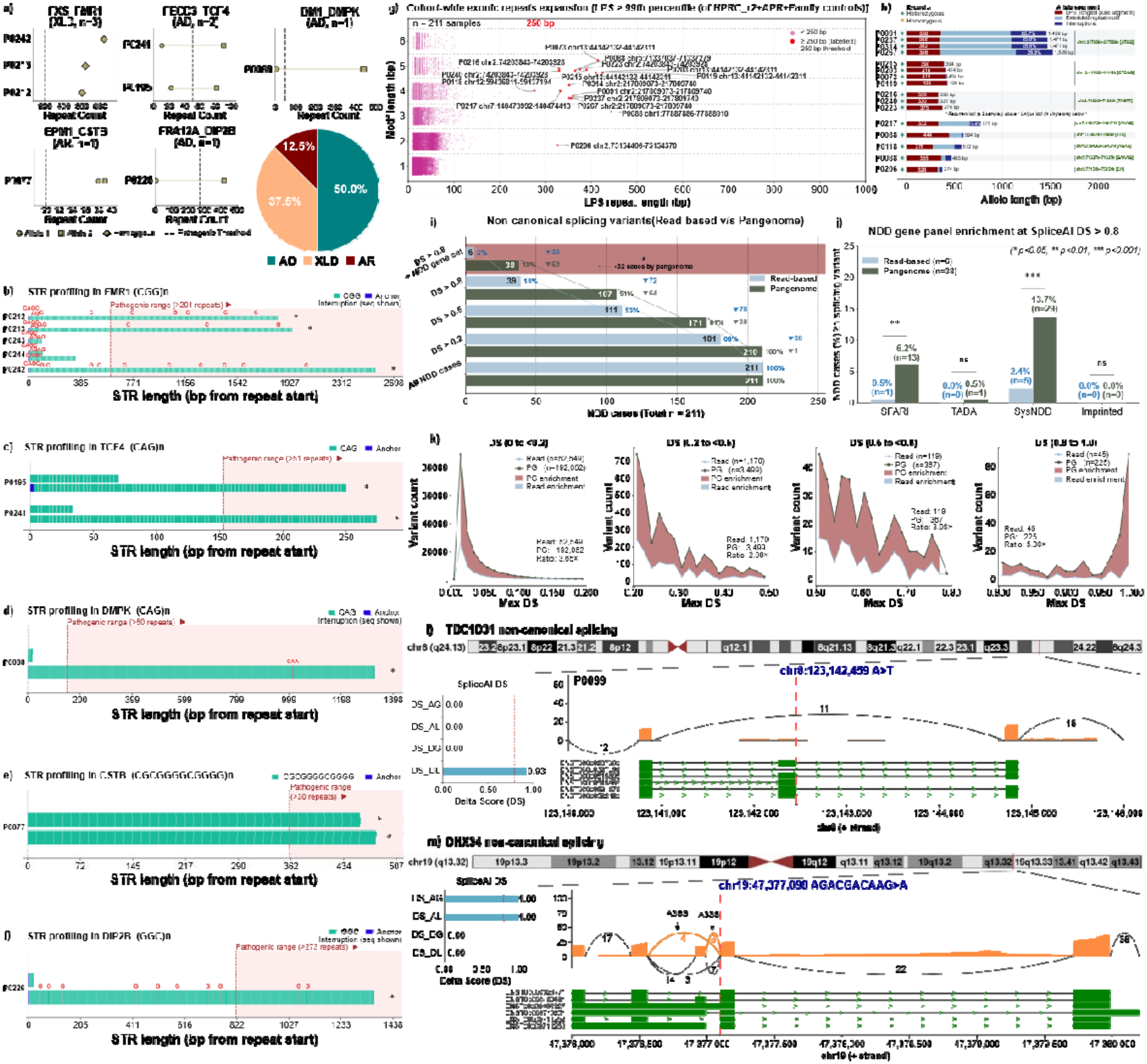
Genome-wide tandem repeat expansion landscape and splicing variant characterization in the NDD cohort. **(a)** Allele-level repeat counts at the seven pathogenic tandem repeat loci identified in this cohort (the circles represent allele 1, the squares represent allele 2, and the diamonds represent hemizygous; dashed line represents the pathogenic threshold). Fourteen expanded loci were identified in 13 distinct samples. The pie chart shows their inheritance mode distribution: 50.0% are autosomal dominant (AD), 37.5% are X-linked dominant (XLD), and 12.5% are autosomal recessive (AR). **(b–f)** Per-allele repeat tract structure at each pathogenic locus (*FXS_FMR1, FECD3_TCF4, DM1_DMPK, EPM1_CSTB* and *FRA12A_DIP2B*). Each horizontal bar is one allele with teal (and orange where the locus has 2 motifs) blocks shown as motif units; blue blocks are flanking anchor sequences, and the interrupting bases are given above the red blocks. The regions are shaded in red are above the pathogenic threshold, and the asterisks (*) indicate alleles exceeding the threshold. **(g)** Longest pure segment (LPS) (bp, x-axis) versus motif length (y-axis) for the global exonic tandem repeat alleles across all 211 NDD cases. Each point represents a single repeat locus; the pink points indicate alleles with an LPS <250 bp, and the red points indicate those with an LPS ≥250 bp, which exceed the expansion threshold (red dashed line). Sample ID and chromosomal coordinates annotated for LPS expansions ≥250 bp. Motif classes of 1–6 bp were assessed, with dense clusters at shorter motif lengths consistent with genome-wide background repeat variation. A similar intronic region LPS expansion plot is shown in Supplementary Figure 15. **(h)** Allele architecture of the expansions identified in panel g. Each bar is a single allele, decomposed into the LPS (red), the surrounding extended motif region (light blue) and interruptions within the tract (dark blue, with the interrupted fraction of the allele shown). All of these are heterozygous (shown by a green diamond marker). Above the dashed line are recurrent loci present in ≥2 samples; below are five loci each observed in a single case, which are annotated with motif coordinates. The ATGG expansions at chr2: 217809k are both the longest alleles observed (∼1.5 kb) and the most heavily interrupted (∼26% of allele length). The list of 230 intronic and 16 exonic allele expansions is provided in Supplementary Table 18b-c. **(i)** Waterfall plot comparing noncanonical variant detection from reads v/s pangenome-based calls under increasing SpliceAI delta score (DS) stringency. Read-based detection falls steeply (211→181→111→39→6 cases), whereas pangenome-based detection is retained more at a stricter DS filter (211→210→171→107→38), with noncanonical splicing variants detected in 32 more cases at the final filter (DS>0.8 intersecting with the NDD gene sets) (see Supplementary Results). **(j)** Percentage of NDD cases carrying ≥1 splicing variant at DS>0.8 within the curated gene panels. Pangenome-based calling exceeds read-based calling in SysNDD (13.7% vs. 2.4%, p<0.001) and SFARI (6.2% vs. 0.5%, p<0.01; Fisher’s exact test), with no significant difference for TADA or imprinted genes. **(k)** Variant count density across four DS bins (0–0.2, 0.2–0.5, 0.5–0.8, and 0.8–1.0). The shading indicates pangenome enrichment over read-based calls, which is greatest in DS 0.8–1.0. **(l, m)** RNA-seq validation of two noncanonical (cryptic) splicing variants. For each gene, the top panel shows the cytogenetic position and the variant within the gene model. The left panel shows the SpliceAI delta scores for acceptor gain (DS_AG), acceptor loss (DS_AL), donor gain (DS_DG) and donor loss (DS_DL); the red dashed line indicates DS=0.8. The right panel is a Sashimi plot of the proband’s long-read RNA-seq data, with exonic coverage and splice junction arcs annotated by junction-spanning read counts; junctions supporting the predicted aberrant event are colored orange. In *TBC1D31* (chr8:123,142,459 A>T, DS_DL = 0.93) **(l)**, the variant disrupts the donor site of the affected exon (exon skipping), whereas in *DHX34* (chr19:47,377,098 AGACGACAAG>A, DS_AG = DS_AL = 1.00) **(m)**, loss of the canonical acceptor is accompanied by the use of two alternative 3′ splice sites (A3SS, orange arcs, 4 and 8 junction reads), confirming the predicted acceptor shift in DHX34.

### Genome-wide TRE profiling reveals expansions beyond known disease loci

Tandem repeats were detected using TRGT across the Platinum Tandem Repeat catalog (∼7.72 million loci), and each allele was summarized by its longest pure segment (LPS) (see Methods). Across 211 cases, 246 expansion calls were identified in 153 distinct cases, 16 of which were exonic expansions and 230 of which were intronic expansions. In the exonic screening, most alleles fell below the 250 bp threshold, as expected for background genome-wide repeat length variation. The 16 exonic calls above the threshold were located within eight distinct loci in 15 (7.1%) unique cases (Figure 6g) and were annotated with sample identifiers and chromosomal coordinates, revealing both recurrent and singleton expansion events distributed across multiple chromosomes. Three exonic loci recurred in two or more unrelated cases. The most prevalent recurrent exonic locus was at chr13:44142132-44142311 (CTCGG, 5-mer), with four cases carrying heterozygous alleles with LPS lengths of 390–425 bp (total allele length 394–429 bp). A second recurrent locus at chr2:217,809k ([ATGG], 4-mer) was also observed in four distinct cases, with LPS lengths of 352–360 bp but substantially longer total allele lengths of 1,459–1,526 bp, reflecting extensive flanking extended repeat motif regions containing 25–26% interruptions in the LPS stretch, which is a structural feature that distinguishes these alleles from simple pure expansions. A third recurrent locus at chr2:74,203k ([TAGTT], 5-mer) was detected in three cases with LPS lengths of 330–375 bp. Five additional unique loci were identified as singletons in individual probands, with LPS lengths ranging from 276 to 444 bp and total allele lengths up to 775 bp, each annotated with chromosomal coordinates and repeat motifs (Figure 6h). Genome-wide analysis of intronic tandem repeats revealed 230 loci with LPS lengths >250 bp in NDD cases, including 10 recurrent loci observed in 4–7 unrelated individuals, with the largest LPS expansions reaching 5.1 kb (Supplementary Figures 16 and 17).

### Noncanonical splicing variant analysis identifies high-confidence splice-disrupting variants in NDD-associated genes

To identify cryptic splicing-disrupting variants, we applied SpliceAI^36^ to ultrarare variants private to 211 NDD cases and restricted them to noncanonical positions more than 2 bp from the canonical donor or acceptor dinucleotide. For each variant, delta scores (DSs) were taken as the maximum across acceptor loss, acceptor gain, donor loss and donor gain, and variants were called high confidence at DS > 0.8 (see Methods). At this threshold, pangenome alignment revealed at least one high-confidence noncanonical splice variant in 107 of 211 cases (50.7%) compared with 39 cases (18.5%) by read-based alignment (McNemar test, p<0.001) (Figure 6i; Supplementary Table 19). These are predictions rather than diagnoses and are restricted to curated NDD gene sets; the difference narrowed to 38 cases (18.0%) (from pangenome calls) versus 6 (2.8%) in read-based calls. Because both methods were applied to the same individuals, the comparison is paired; 34 cases were detected only by the pangenome, and 2 were detected only by read-based alignment (McNemar test, p = 2.38 × 10 ^7^). At the individual panel level at DS > 0.8, the pangenome revealed 13 SFARI cases (6.16%) versus 1 (0.5%) (p < 0.001), 29 SysNDD cases (13.74%) versus 5 (2.37%) (p < 0.001), 1 TADA case (0.47%) versus 0, and 0 imprinted cases in either approach (Figure 6j). Detection was significantly higher for the SFARI and SysNDD genes, whereas the TADA and imprinted comparisons were underpowered given the very small variant count. The variant count distribution across DS bins showed pangenome-to-read ratios of 2.92, 3.01, 3.11, and 5.00-fold for the 0–0.2, 0.2–0.5, 0.5–0.8, and 0.8–1.0 bins, respectively, indicating that the pangenome advantage is pronounced in the highest confidence-predicted variant set (Figure 6k). Two of the validated noncanonical splice-disrupting variants are illustrated, showing the exon skipping events in TBC1D31 and DHX34, both overlapping with the NDD-associated gene set (Figure 6l–m).

### Mitochondrial (MT) genome analysis

Heteroplasmic mtDNA variants were called across all 318 individuals, yielding 69 heteroplasmic genotypes. After filtering for ≥100X depth and a heteroplasmy level between 20% and 80%, 23 calls were retained (13 in cases and 10 in controls). Two case calls were also heteroplasmic in a control and were excluded; in both instances, the control was the proband’s mother (m.14196A>G and m.16192C>T), which is consistent with maternal transmission rather than a somatic or *de novo* event. This filtering retained 5.2% (11/211) of the case-unique heteroplasmies, each in distinct individuals. Heteroplasmy ranged from 21.8– 75.2%, with most variants in the 20–40% band. Five variants were found in the protein-coding OXPHOS subunits, including 3 in Complex I (MT-ND1 m.3524C>T, MT-ND4 m.12013A>G, and MT-ND5 m.13689A>G), one in Complex IV (MT-CO2 m.7743C>T) and one in Complex V (MT-ATP8 m.8557G>A). Four variants were identified in the D-loop control region and 2 in the rRNA genes (Figure 7a).

**Figure 7.**
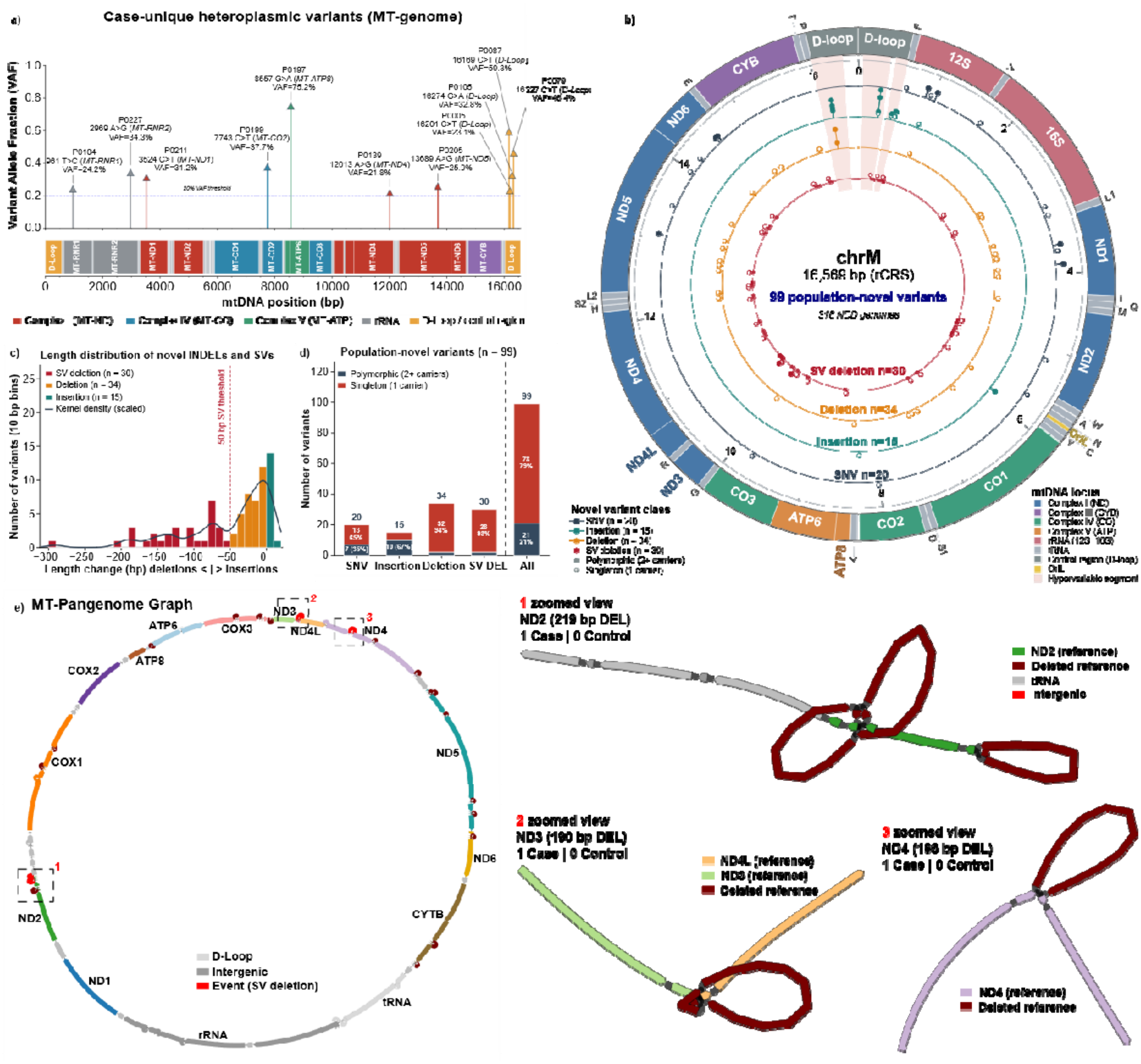
Analysis of mitochondrial heteroplasmy and novel variants. **(a)** Case-unique heteroplasmic variants (n=11) called HiFi alignments using Mitorsaw within the 20%<VAF<80% range. Each lollipop is one variant, positioned at its rCRS coordinate on the x-axis, with the stem height showing the VAF along the y-axis (the dashed line marks the 20% VAF threshold). Labels show the sample ID, position, base change, gene and VAF. Stem and marker color denotes the functional compartment of the affected locus. The track below (x-axis) shows the rCRS locus annotation on the same coordinate scale, using the same color scheme. **(b)** Population-novel variants from the mitochondrial pangenome graph mapped on the rCRS (16,569 bp). The outer ring gives the locus annotation colored by functional class, with tRNAs labeled by single-letter amino acid codes and the OriL marked and hypervariable segments shaded. Concentric tracks, outermost to innermost, show the SNVs (n = 20), insertions (n = 15), deletions (n = 34) and structural deletions ≥50 bp (n = 30). Filled circles show polymorphic (≥2 carriers) sites, whereas the open circles mark the singleton variants. **(c)** Length spectrum of the 79 novel INDELs and structural deletions in 10 bp bins, colored by class. The negative values are deletions and positive insertions, the dashed line indicates the 50 bp SV threshold, and the solid curve is a kernel density estimate scaled to counts. **(d)** Carrier distribution of the 99 population novel variants, by class and in total. Each bar is stacked into polymorphic (≥2 carriers) and singleton (1 carrier) fractions, with counts and percentages labelled. Singletons account for most of the set overall (78/99, 79%) and nearly all deletions (32/34, 94%) and structural deletions (28/30, 93%), whereas insertions are mostly polymorphic (10/15, 67%). **(e)** Mitochondrial pangenome graph of the 318 cohort mitogenomes in a circular layout along with the gene annotations for respective colored segments on the graph. The nodes are colored by the rCRS locus they traverse (key inset), with reference segments deleted by a structural event in red. The red boxes numbered on the graph mark the three representative events, each shown as zoomed view of the alternate path bypassing the deleted reference interval (dark red) [subpanel 2, 291 bp deletion at rCRS 5,211 (*MT-ND2*); subpanel 2, 190 bp deletion at rCRS 10,334 (*MT-ND3*); and subpanel 3, 186 bp deletion at rCRS 11,062 (*MT-ND4*)]. Each was carried out by one affected and no unaffected individual.

Projection of the NDD cohort MT-pangenome graph onto the revised Cambridge Reference Sequence (NC_012920.1) revealed 1,374 variants across 16,569 bp (82.9 variants per kb), comprising 1,258 SNVs (91.6%), 46 small deletions (3.3%), 40 small insertions (2.9%) and 30 SV deletions (2.2%). The individual MT-genomes carried a median of 36 variants (range 12–99). Across the full call set, 826 variants (60.1%) were polymorphic (in ≥2 samples), and 548 (39.9%) were singletons.

After sequential filtering against reference databases, 99 novel variants (7.2% of the call set), comprising 20 SNVs, 49 INDELs and 30 SV deletions, were retained (Figure 7b; Supplementary Table 20). All 30 SVs were deletions, ranging from 57 to 291 bp (Figure 7c), each observed in no more than two individuals. Compared with the full call set (39.9%), the novel set was enriched with singleton variants (78/99, 78.8%), particularly among small deletions (32/34, 94.1%) and SV deletions (28/30, 93.3%) (Figure 7d). After excluding 16 calls (see Supplementary Results), we retained 83 population-novel variants carried by 88 individuals (27.7%), of which 74 (89.2%) were singletons. Novel variants were distributed across all major functional compartments, 38 in complex I subunits, 13 in rRNA genes, 12 in complex IV, 11 in tRNAs, 6 in MT-CYB, one in complex V and 16 in the control region, with 2 intergenic regions. Circular visualization of the MT-pangenome graph constructed from the 318 cohort mitogenomes, with nodes colored according to the rCRS locus traversed, is shown in Figure 7e. Reference segments are deleted in the SV calls, with representative zoomed subpanels showing 3 structural alternate mitogenome traversal walks (MT-ND2, 219 bp DEL; MT-ND3, 190 bp DEL; MT-ND4, 186 bp DEL). These events were singleton and absent in family controls.

## Discussion

In this study, we combined PacBio HiFi sequencing, haplotype assembly and pangenome-graph analysis across an ancestrally diverse NDD cohort and our complementary reference panels. The cohort spans the Middle East, Africa, North Africa and Central/South Asia, representing populations that remain largely unexplored in population genomics and NDD research. We set out to assess what a pangenome built from this study cohort itself adds to rare disease variant discovery, particularly in populations underrepresented in existing genomic references. Within the high-confidence regions, pangenome-based approaches identified additional classes of variation relevant to NDD risk that linear reference-based analyses could miss or characterize incompletely. For example, gene duplication events in terms of haplotype composition, nonreference sequences, rare multiallelic complex loci, large insertions, noncanonical splicing impacts and repeat expansions are not routinely assessed in standard clinical genetics settings. These findings also reveal how ancestry, demographic history and local genomic architecture can shape variant discovery and interpretation in rare diseases.

The combination of ancestral diversity and heterogeneous NDD phenotypes adds to the complexity of variant interpretation and highlights the importance of multipopulation pangenome references for enhancing the understanding of population genomics and the discovery of disease etiology. The sequence absent from the reference altogether is the most direct measure of the reference gap for these populations. All disease associations were reported for GRCh38 because clinical annotations and databases for T2T-CHM13 remain limited despite its more complete sequence. We therefore used CHM13 as the graph backbone and GRCh38 for clinical annotation, which retained compatibility with established interpretation resources while still drawing on multiple haplotypes for variant discovery. The graph embeds 567.8 Mb of nonreference sequence, including 28,459 sequences totaling 49.66 Mb carried only by affected individuals; 90.7% of these sequences occurred in a single case. Although the accumulation curve approached saturation, nonreference sequences continued to accrue, with the addition rate declining 5.5-fold between the first and last 50 haplotypes. Such sequences are inaccessible to conventional linear-reference alignment when no corresponding reference sequences exist.

We identified 704 exonic sequences in 671 protein-coding genes unique to NDD cases, impacting bonafide NDD genes with large insertions in the *BTRC, MEGF8, ACADSB*, and *NDUFA9* genes. Clinical genomics can be utilized to identify additional recessive and loss-of-function variants to resolve complex rare disease cases that might not be readily detected through long-read-based linear genome analyses. From the pangenome graph, we identified 8.1% of cases harboring recessive homozygous exonic complex structural rearrangements. In 2.4% of the cases, these rearrangements affected genes like AHNAK2 and EPPK1 which are associated with NDD^37^. Similarly, in the noncanonical splicing analysis, graph-derived calls revealed high-confidence splice-disrupting variants in NDD-associated genes in 38 of 211 cases compared with 6 from read alignments (6.3-fold difference). We are deliberately careful about the interpretation of these variants. Although these findings are consistent with improved detection in complex genomic contexts, establishing causality for noncoding variants will require functional validation and clinical interpretation.

Genome-wide duplication burden was higher in NDD cases compared with combined family, HPRC_r2 and APR controls within the analyzed segmental-duplication filtered regions. Five genes were enriched in 33 cases. The enrichment of *MAPT* at 17q21.31 was notable given the structural complexity and established neurodevelopmental relevance of this locus^38^. Although HPRC_r2 covers most of the super population, ancestry-matched controls are scarce for some of these populations, and residual population structure may contribute to apparent case control associations. Tandem repeat analysis revealed several potential clinically-relevant findings, including a full FMR1 mutation arising from a maternally transmitted premutation, a biallelic *CSTB* expansion consistent with Unverricht-Lundborg disease, and *DIP2B* findings in one case. These findings illustrate the utility of LRS approaches for resolving large pathogenic expansions that are systematically refractory to short-read approaches. With respect to genome-wide expansions, functional significance has not been established to date, but their size and recurrence across unrelated individuals from diverse ancestral backgrounds warrant further investigation, particularly at loci within or proximal to NDD-associated genes.

There are a few other limitations of these conclusions. Although the cohort captures substantial haplotype diversity from underrepresented populations, it is modest in size for association testing. In particular, the trio subset limits the power for comparisons of *de novo* burden and significance. Owing to underrepresentation, ancestry matching between the cohort and the reference panels is imperfect at the level of individual populations, even when the superpopulation composition is comparable; for instance, the African-ancestry genomes, which show more nonreference sequences than any other superpopulation group does. The route past that boundary is not more depth per genome but more genomes and better graphs. Future work will require larger, more ancestry-representative cohorts and improved graph representation of difficult genomic regions. The populations sampled in this study contribute 636 haplotypes toward that goal, from a disease cohort rather than a healthy reference panel, with the nonreference sequence they carry accumulating further with each genome added. The restriction to ‘easy regions’ represents a deliberate trade-off, prioritizing the feasibility and cost-effectiveness of standard clinical HiFi sequencing over comprehensive coverage of difficult regions. This approach reflects the practical constraints of sequencing clinical samples and the need for methods that can be implemented on a scale. Despite this restriction, the findings remain promising, supporting the equitable advantage and the value of standard clinical HiFi sequencing even at the coverage achieved in this study. Overall, these findings support a broader shift in rare disease genomics from single-reference variant discovery toward multilayered long-read and pangenome-enabled analyses. For NDDs in underrepresented populations, such approaches offer not only improved sensitivity for clinically relevant variation but also a richer view of disease architecture that would remain partially hidden under conventional frameworks. Clinical genomics will rely increasingly on haplotype diversity as population-specific pangenomes grow. Our study provides a framework for more equitable genomic analysis, demonstrating how a single routine clinical long-read genome can support both linear reference-based clinical diagnosis and pangenome-based discovery, advancing the potential for more comprehensive genomic characterization within existing clinical sequencing workflows.

## Methods

### Ethics and study participants

This study was approved by the Dubai Scientific Research Ethics Committee (DSREC-09/2024-01) and the Institutional Review Board of Mohammed Bin Rashid University of Medicine and Health Sciences (MBRU-IRB-2017-004). Written informed consent was obtained from adult participants and from a parent or legal guardian on behalf of minors. Consent covered clinical phenotyping, whole-genome sequencing, and controlled sharing of deidentified genomic data. The sequenced cohort comprised 318 individuals, including 211 individuals with a clinically ascertained NDD and 107 unaffected family members who served as within-cohort controls. Participants were recruited from Al Jalila Children’s Specialty Hospital, Al Amal Hospital, Mediclinic Middle East, and NeuroGen Healthcare, which represent 24 nationalities spanning Africa, North Africa, the Middle East and Central/South Asia. Affected individuals were enrolled either as singletons (n = 150) or within 61 families comprising parent–offspring trios, duos and extended structures so that segregation and parent-of-origin analyses could be performed where family members were available. The inclusion criteria included a clinical diagnosis of ASD, ID, GDD, epilepsy or congenital brain malformation made by a pediatric neurologist or clinical geneticist. Two external population reference panels were used as controls: 231 individuals from the Human Pangenome Reference Consortium release 2 (HPRC_r2)^23^ and 53 individuals from the Arab Pangenome Reference (APR)^13^. Combined with the 107 family controls, this resulted in 391 control individuals (782 haplotypes), against which the 211 cases (422 haplotypes) were compared.

### Clinical phenotyping and HPO harmonization

Clinical information was extracted from medical records and standardized case report forms. Reported clinical features were mapped to Human Phenotype Ontology (HPO) terms (https://hpo.jax.org/). Terms were normalized to their current primary identifier, obsolete identifiers were replaced by their replacement terms, and duplicate terms within an individual were collapsed before analysis. A binary sample-by-phenotype matrix (211 cases × 267 unique HPO terms) was constructed and used to compute term frequencies, sex-stratified counts and pairwise cooccurrences. Cooccurrence between two terms was defined as the number of individuals annotated with both terms; the diagonal of the cooccurrence matrix reports the total number of carriers of each term. The ten most frequent terms are displayed in the main figure, and the twenty most frequent terms are provided in the supplementary figure; the complete matrix is provided in Supplementary Table 2b. Self-reported nationality was used only for descriptive summaries and for stratifying figures. Genetic ancestry was assessed independently by principal component projection and model-based admixture inference as described below.

### DNA extraction, PacBio HiFi library preparation and sequencing

High-molecular-weight genomic DNA was extracted from peripheral blood using a GenFind V3 kit or a Nanobind HMW DNA extraction kit (Pacific Biosciences) according to the manufacturers’ instructions. DNA concentration was quantified by Qubit dsDNA fluorometry, and DNA integrity and fragment size distribution were assessed using the Agilent TapeStation Genomic DNA ScreenTape assay. PacBio HiFi libraries were prepared using the SMRTbell Prep Kit 3.0 and size-selected to enrich DNA fragments of approximately 15–18 kb. Libraries were sequenced on the Pacific Biosciences Revio platform using 24M SMRT Cell with a 24-h movie time. Circular consensus sequencing reads were generated using SMRT Link (v26.1) with default settings and a minimum predicted read accuracy of Q30. For each sample, the sequencing yield, read N50, median read quality and estimated genome coverage were calculated from the HiFi reads. Genome coverage was estimated relative to a 3.1-Gb haploid genome. Chromosome-level coverage was subsequently evaluated against T2T-CHM13 v2.0, including autosomes, chromosome X, centromeric satellite classes and ribosomal DNA intervals. Samples meeting the predefined sequencing-quality criteria of ≥25× coverage, a read N50 ≥10 kb and a median read quality ≥Q30 were retained for downstream analyses.

### Read alignment, small-variant calling and phasing

HiFi reads were aligned to GRCh38 (analysis set, no ALT contigs) with pbmm2 (v1.17.0, https://github.com/PacificBiosciences/pbmm2) using the CCS preset. Small variants (SNVs and INDELs < 50 bp) were called per sample with DeepVariant (v1.9.0). Per-sample gVCFs were jointly genotyped for the trios with GLnexus (v1.4.1) using the DeepVariantWGS configuration^39^. Read-backed phasing was performed with WhatsHap (v2.7)^40^ and alignments were haplotagged *‘whatshap haplotag’* for downstream haplotype-aware analyses. Only variants with PASS filters were retained for downstream analysis. Variants were left-aligned and normalized with *bcftools norm -f GRCh38.fa -m -any* (BCFtools v1.22)^41^.

### Population structure and ancestry inference

Study genomes were projected onto principal components derived from the 929 Human Genome Diversity Project (HGDP)^42^ reference panel. Biallelic autosomal SNPs were allele-matched between our study and HGDP datasets, and monomorphic sites were removed. Reference markers were filtered at minor allele frequency ≥ 0.05 and genotype missingness ≤ 0.10, followed by linkage disequilibrium pruning using a 500 kb sliding window and a r^2^ threshold of 0.20, leaving 128,181 markers. Principal components were computed in the HGDP reference samples using SNPRelate^43^ (v1.32.0), and the study samples were projected onto the fixed reference axes using the corresponding SNP loadings. Unsupervised ancestry inference was performed using ADMIXTURE^44^ (v1.3.0) for K = 2 to 6, with model fit assessed using cross-validation error. Y-chromosome haplogroups were assigned for male participants using Y-LineageTracker^45^. Individuals were assigned to four analysis superpopulations, namely, the Middle East, North Africa, Africa, and Central/South Asia, on the basis of self-reported nationality.

### ROH and parental-origin analysis

ROH were called with Automap^46^ (v1.3) from PASS-filtered autosomal small-variant call sets for NDD cases, HPRC_r2 controls and APR controls, using a minimum ROH length of 1 Mb and otherwise default parameters. Because AutoMap operates on variant call sets rather than array genotypes, all three cohorts were processed from call sets generated with the same variant-calling workflow to avoid technical differences in ROH inference. The per-individual total ROH length was computed as the sum of all called segments, and individuals were stratified into five categories: <25 Mb, 25–50 Mb, 50–150 Mb, 150–300 Mb and >300 Mb. ROH burden was compared between 211 NDD cases and 284 population controls from HPRC_r2 and APR. ROH intervals were intersected with GENCODE^47^ (v48) gene annotations using bedtools^48^ (v2.31.1) intersect, and unique overlapping genes were counted per individual and stratified by gene biotype. ROH intervals were further intersected with curated gene sets [SFARI, SysNDD, TADA, OMIM, imprinted genes and T10p_BEGs (top 10th percentile of brain-expressed genes)] (see “Gene sets” in Supplementary Table). Gene-set enrichment was assessed at the individual level. An individual was scored as a carrier if any ROH segment overlapped at least one gene from the set and compared between cases and controls by Fisher’s exact test. In trio probands, each ROH segment was compared with the parental ROH profiles and classified as paternal, maternal, shared by both parents, or proband-specific (homozygous stretch each from one parent).

### Clinical variant annotation and ACMG interpretation

GRCh38 small variants called using DeepVariant^49^ (v1.9.0) were annotated using HORIZON^27^ and independently with ANNOVAR^28^, and classification concordance between the two tools was assessed. Variants were classified according to the American College of Medical Genetics and Genomics (ACMG) guidelines for variant pathogenicity interpretation^50^, with findings further verified by clinical geneticists. GRCh38 was retained as the coordinate system for clinical annotation because clinical variant databases and gene models are substantially more complete in that reference space. Variant prioritization considered predicted molecular consequences, population allele frequency (<0.001) in gnomAD^30^ (v4.1) and the in-cohort controls, mode of inheritance, family segregation, and concordance between the affected gene and the HPO profile of the samples. Variants were classified under the ACMG/AMP^50^ (2015) framework as pathogenic (P), likely pathogenic (LP) or of uncertain significance (VUS). All variants reported as clinically relevant were reviewed independently by geneticists. All 92 reported variants were inspected in Integrative Genomics Viewer^51^ (IGV) to confirm read support and exclude local alignment artifacts and were orthogonally validated by Sanger sequencing.

### De novo SNV/indel detection and parent-of-origin assignment

*De novo* SNVs and INDELs shorter than 50 bp were identified from 54 parent–offspring trios using the DeepTrio^52^ (v1.9.0) workflow. One affected trio was excluded before burden analysis because of an anomalously high *de novo* count, leaving 53 trios comprising 42 affected probands and 11 unaffected sibling controls. Candidate calls were filtered on the basis of proband genotype quality (GQ ≥ 20), allelic balance and depth (DP ≥ 8), and the absence of alternate-read support in both parents. Candidate *de novo* variants were further required to be absent from gnomAD^30^ (v4.1). The parent-of-origin was assigned using phased inherited heterozygous variants. Joint calling was processed through GLnexus^39^ (v1.4.1) for cohort-level genotyping and phased with WhatsHap^40^ (v2.7). HiFi reads spanning a *de novo* variant and a nearby informative marker were used to assign the event to the paternal or maternal haplotype. Calls with insufficient or discordant read evidence were left unassigned.

### Read-based structural-variant calling and clinical prioritization

SVs were called from HiFi read alignments to GRCh38 using sawfish^35^ (v2.1.1). An upper size bound of 10 Mb was applied and only calls with FILTER = PASS were retained. SVs overlapping the centromeric, pericentromeric and gap regions were subsequently removed. The retained variants were annotated (using HORIZON) for gene overlap (GENCODE v48) and predicted consequences and then filtered for rarity at an allele frequency <0.01 against gnomAD-SV^53^ (v4.1), Database of Genomic Variants^54^ (DGV), HPRC_r2^24^ and APR^13^ and in-cohort family control call sets. Overlaps with ClinVar pathogenic SVs relevant to NDD were also checked for the filtered variants. The filtered SVs were also evaluated for overlap with ClinVar-reported pathogenic SVs associated with NDDs. Prioritized SVs were classified according to the ACMG/ClinGen technical standards for copy-number variant interpretation. Candidate pathogenic SVs were reviewed in IGV^51^ and orthogonally validated by Oxford Nanopore sequencing using the Ligation Sequencing Kit (ONT-LSK). ONT reads were aligned using minimap2 (preset *map-ont*), and the candidate SVs were confirmed by manual inspection.

### Haplotype assembly and assembly quality control

Assembly haplotypes were generated for all 318 genomes with hifiasm^55,56^ (v0.25). Trio mode was used for the 54 parent–offspring trios, with parental k-mers used to partition the proband reads before assembly; the other samples were assembled in standard mode. Assembly contiguity was summarized per haplotype by contig N50 and total assembled length. Consensus quality values (QVs) were estimated with yak^57^ *qv* (v0.1) against a k-mer database built from the sample’s own HiFi reads. Assembly contiguity profiles were compared against those of T2T-CHM13^58^ (v2.0) and against those of gap-split GRCh38^59^. Small-scale assembly errors were evaluated with HMM-Flagger (v1.2). HiFi reads were mapped jointly to both haplotypes with minimap2^60^ (v2.29) using *“-k 25 -ax lr:hqae --cs -- eqx -L -I8g”*. Alignments were filtered with correct_bam using *--primaryOnly --minReadLen 5000 --minAlignment 5000 --maxDiv 0.1*. HMM-Flagger^61^ was run with the *--hifi* preset and centromeric satellite annotations, and the conservative error set was used for downstream summaries. The sequences were partitioned into haploid (Hap), erroneous (Err), duplicated (Dup) and collapsed (Col) states. Gene-space completeness was assessed by lifting GENCODE^47^ (v48) protein-coding annotations to each haplotype with Liftoff^62^ (v1.6.3). In brief, foreign contamination from exogenous DNA, such as EBV and residual adapter sequences, were identified with FCS-GX^63^ (v0.5.5) and removed from the assemblies. Assembly and graph-based discovery were fixed to ‘easy regions’ before any discovery claim was made, using the Genome in a Bottle (GIAB) union stratification of regions falling outside every difficult category on GRCh38 (https://ftp-trace.ncbi.nlm.nih.gov/ReferenceSamples/giab/release/genome-stratifications/v3.6/GRCh38@all/Union/GRCh38_notinalldifficultregions.bed.gz), which excludes segmental duplications, satellite and centromeric sequences, long tandem repeats, low-mappability intervals, GC extremes and the known problematic and false-duplication regions of the reference. Mitochondrial assemblies were not integrated into the remaining genome, as NUMTs were identified with BLAST and removed. The mitochondrial reads were identified and assembled as separate circular contigs using Flye^64^ (v2.9.6).

### Gene annotation and gene duplication analysis

The gene copy number was derived per haplotype from the Liftoff^62^ (v1.6.3) annotations, which were run with extracopy detection enabled (*-copies -sc -p 0.9*). For each haplotype, the number of mapped copies of each GENCODE^47^ (v48) gene was compared with the expected copy count in the GRCh38^59^ reference annotation, and a gene was classified as duplicated when the observed copy number exceeded the reference-expected copy number. This reference correction is necessary because a subset of genes is not single-copy in GRCh38, and without it, those genes generate systematic false-positive duplication calls in every haplotype. The duplication burden per haplotype was restricted to autosomal protein-coding genes for the cohort-level burden comparison to avoid sex-dependent differences in chrX and chrY copy number.

Case control comparisons of per-haplotype burden were performed. For the gene-level case control comparisons, one-sided [NDD cases (422 haplotypes) > all controls (782 haplotypes)] Fisher’s exact tests were used for the carrier counts, with Benjamini–Hochberg (BH) correction applied throughout the genome across all the tested protein-coding genes. Gene-set level comparisons (SFARI, SysNDD, TADA, and imprinted) were performed identically and corrected separately. For trios, each duplicated gene in a proband was classified as paternal, maternal, present in both parental backgrounds, or *de novo* according to the parental haplotype copy profiles. A *de novo* call required the gene to be nonduplicated on all four parental haplotypes.

### Pangenome graph construction and graph QC

A cohort pangenome was constructed from the haplotype assemblies of the full NDD cohort using the Minigraph-Cactus^25^ (v2.9.3) pipeline (cactus-pangenome). T2T-CHM13^58^ (v2.0) was used as the primary reference backbone, and GRCh38 was included as a second reference path, enabling variant projection onto either coordinate system. The pipeline was run on a single 128-core/2.2 TB RAM node. The pipeline produced clipped and unclipped GBZ^65^ and VG graphs^66^, chromosome-level graphs, GFA files and VCFs projected onto both reference coordinate systems. During graph clipping, nodes present in fewer than 9 input haplotypes were removed from the clipped graph. The full unclipped graph was retained for all analyses requiring private and rare sequences. The graph size was summarized by node count, edge count, edge-to-node ratio and total embedded sequence using *vg stats* (v1.61.0). Pangenome growth curves and sequence-frequency classes were computed with Panacus^26^ (v0.4.1). The cumulative nonreference sequence was calculated over the NDD cohort haplotypes as the haplotypes were added in fixed order. The nonreference sequence was classified by the number of cohort samples in which it was observed: core (present in ≥ 95% of haplotypes), common (≥ 5%) and unique (in 1 haplotype). Variants were deconstructed from the graph with *vg deconstruct* (v1.61.0) using *‘-P CHM13’* and *‘-P GRCh38’* to generate call sets in each coordinate system. Postprocessing followed the PanGenome_VCF_PostProcess workflow (https://github.com/Shuhua-Group/PanGenome_VCF_PostProcess). In brief, nested bubbles were flattened with *vcfbub*, alleles at multiallelic structural loci were grouped using a 50 bp length similarity threshold so that alleles differing by less than 50 bp in length at the same locus were treated as one allelic class, records were then split into structural (≥ 50 bp) and small-variant call sets, and call sets were decomposed with *bcftools norm -m -any -f GRCh38.fa* (BCFtools v1.22)^41^ for left alignment and biallelic representation. Normalization was deliberately applied only after allele grouping because decomposing multiallelic records before this step destroyed the alternate signal on which complex-SV identification depended.

### Ultrasrare/unique SNV, indel and SV analysis

Cohort-unique (ultrarare, 0% frequency) variants were defined as variants absent from all interrogated population variant resources. The GRCh38 call set was used to compare unique variants identified through read-based and pangenome-based approaches. Unique variants were identified using PanScan^67^ (v1.6) *“find_uniq_variants”*. For the read-based approach, the SNVs and INDELs were sequentially filtered against our APR and HPRC_v2 DeepVariant calls, followed by successive filtering with dbSNP^29^ (GCF_000001405.40), gnomAD^30^ (v4.1), the 1000 Genomes Project (1KGP)^31^ (Phase 3 integrated SV call set, v8), HGSVC ^31^ (v1.0), and the Greater Middle East variome^32^ (GME). At each filtering step, variants matching the respective resource were removed, retaining only unmatched variants for subsequent filtering. This sequential approach quantified the marginal contribution of each resource to variant filtering and identified the final set of cohort-specific variants. For the pangenome-based variants, initial filtering was performed using our APR and HPRC_v2 pangenome variants as described previously. Variant matching between the call set and reference databases required an exact match in chromosome, genomic position, reference allele, and alternate allele for each SNV and indel after normalization. For SVs, comparisons were performed against dbSNP^28^, gnomAD^65^, 1KGP^30^, HGSVC^32^ and the Database of Genomic Variants (DGV)^54^ (25 February 2020 release, GRCh38). SVs were considered matching if they exhibited ≥80% reciprocal overlap, as determined using PanScan^67^. An SV was classified as unique if it either had no matching entry in any reference SV database or if its reciprocal overlap with any reference SV fell below the 80% threshold. Variants passing all the filters were designated unique to the NDD cohort.

### Novel-sequence identification and annotation

Novel sequences in the NDD cohort were defined as SV insertion alleles of at least 50 bp present in the GRCh38-deconstructed pangenome VCF that were absent from our HPRC r2 and APR pangenome references and were identified using the *“novel_seq”* module of PanScan^67^ (v1.6). Multiallelic records were normalized with *bcftools norm* (BCFtools v1.22)^41^ and decomposed by variant class with RTG Tools^68^ (v3.13). Candidate insertions were compared with the population references (HPRC_r2 and APR) using Truvari^69^ *bench* (v4.0.0) at an 80% reciprocal-overlap threshold. Redundant calls at the same locus were collapsed, and the retained sequences were clustered by coordinate, orientation and sequence similarity using CD-HIT-EST^70,71^ (v4.8.1). Each resulting cluster was represented by a single consensus sequence, which was reported as a nonredundant novel sequence.

For the case-unique analysis, insertions present in any of the 214 family control haplotypes were excluded, retaining only sequences absent from all controls (0% frequency) and specific to affected probands. Retained sequences were intersected with GENCODE annotations (v48) with bedtools^48^ (v2.31.1) and classified as exonic, intronic, lncRNA, misc_RNA or intergenic, with each sequence assigned to a single class. Intersections with the SFARI, SysNDD, TADA, imprinted and T10p_BEG sets were computed at the sequence, gene and unique-carrier levels. Per-individual novel-sequence burden was compared across superpopulations by the Kruskal Wallis test with BH-corrected pairwise Mann Whitney U tests, and effect sizes were reported as rank-biserial correlations. The selected case-unique insertions overlapping exons of the NDD-associated genes were inspected at the sequence level by aligning the insertion sequences to the corresponding local reference windows using minimap2^60^ (-ax asm20) and visualizing the alignment using SVbyEye^72^ (0.99.0) with plotMiro and plotAVA, with the gene annotation tracks overlaid.

### Identifying complex rearrangements and rare exonic events in local haplotype walks

Complex multiallelic loci were identified directly from the pangenome graph rather than from reference coordinates, following our previous approach^12^. These sites were identified as those within a 100 kb window if more than 1 SV of 5 kb was present and if at least 5 alternate structural haplotypes traversed this path. Retained bubbles were intersected with GENCODE v48 to lift over the annotations on the graph haplotype walk. The unit of analysis was the exact allele (one alternative sequence at one record), keyed by chromosome, position and alternative sequence. An allele was considered structural when its reference or alternative sequence was at least 50 bp. Each allele was decomposed at the node level by comparing the ordered node lists of the alternative and reference paths, with node lengths from the graph and an assertion that summed node lengths equal the alternative length. Components of at least 50 bp defined the class (deletion, insertion, tandem duplication by a flank test up to 20 kb, replacement, separated or multiblock complex, inversion), and smaller components were retained but did not define an event. Branches (haplotype paths through 50 bp sites at 50 bp resolution), structural families (same site class, deleted span and inserted length within 2 percent) and site tokens were derived for description only.

For exon consequence and assembly confirmation, structural sites were intersected with the coding exons of ENSEMBL canonical transcripts (332,140 exons). Gene annotations were lifted onto every haplotype assembly with Liftoff, and a per-haplotype exon index was recorded for each exon to determine whether it was intact, altered, divergent, absent or not assessable. An allele gene pair was PAV call supported when at least two carrying haplotypes were assessable, at least one agreed with the prediction, and none disagreed. Alleles whose unique sequence contained a run of at least 50 undetermined bases (stretches of *N*’s) were set aside as scaffold gap artifacts. The layer was checked at known loci (homozygous deletions at TUBA3E and LCE3B/LCE3C, mobile-element insertions at CUL5 and GNPTAB, in-frame expansions at DSPP and MUC17, and a tandem-duplication call at NACAD as a negative example).

Exon-impacting alleles were assigned by their highest zygosity among affected individuals to a homozygous class (at least one affected homozygote) or a heterozygous case-only class (affected heterozygotes only and no unaffected carrier). Unaffected heterozygotes and homozygotes were tabulated for every allele, and an allele with an unaffected homozygote for the same sequence was labeled strongly disfavored under a high-penetrance recessive model. No frequency threshold, score or ranking was applied at any step, and carrier counts, independent-family counts and external-panel frequencies were reported descriptively. Inserted and replaced sequences were annotated with RepeatMasker^73^ (v4.2.4) and Tandem Repeats Finder^74^ (TRF) (4.10.0-rc.2) and assigned one content class per allele, mobile-element insertion (LINE, SINE, SVA or LTR/ERV), tandem repeat (VNTR), exon-paralog-derived, reference-derived, novel unclassified, or no gained sequence for alleles that only remove sequences.

Each homozygous and case-only heterozygous allele was validated in the raw HiFi reads of a class-matching affected carrier (a homozygote for homozygous alleles, any affected carrier for heterozygous alleles), independent of the assemblies and the graph. The complete read set of the carrier was screened with minimap2^60^ (v2.24, *map-hifi*) against sequence models of the reference window, the alternative window and, where relevant, the carrier haplotype, deletion-only and insert-only models; matching reads were extracted with samtools^41^ (v1.21) and realigned to each model. A read was locus linked when it carried at least 300 bp of a reference-unique sequence adjacent to the tested feature, and a junction read when it had 300 bp clean windows on each side of the breakpoint, an identity of at least 0.95, anchoring on every side and an alignment score of at least 10 above the reference model. A call required at least three such reads; negative calls required a flank depth of at least 10x; and a supported homozygous exon loss required 15x depth, 15 anchored reads, junction reads and no locus-linked read containing the intact exon. Windows with less than 50% reference-unique sequences were classified as unresolved and paralog-compatible. Each allele received one verdict (read-supported, contradicted, unresolved paralog-compatible, ambiguous or not assessable), reported beside it; a contradicted or ambiguous verdict did not remove the allele from the catalog. Read support was treated as structural validation only.

Each allele was detected in two CHM13-based pangenome panels, APR^13^ (106 haplotypes) and HPRC_r2^24^ (462 haplotypes). For every external haplotype, the sequence between the two reference anchor nodes flanking the record was reconstructed from the graph walk and compared with the cohort reference and alternative sequences; a haplotype was assessable only when both anchors were crossed once within one contig with reference-identical flanks and was otherwise recorded as not assessable rather than absent. Presence was reported at the exact-allele, structural-family and site-token levels with per-panel numerators and denominators, and external homozygotes were derived as carrier haplotypes minus carrier genomes. The panels served as the population context and were not pooled with the family controls.

Parental and sibling genotypes defined a per-family pedigree state for every allele (Mendelian-consistent, Mendelian-inconsistent, parents genotyped without a homozygous offspring, or no informative relatives), and only Mendelian inconsistency was treated as negative evidence. The identified genes were intersected with the SFARI, SysNDD, TADA, imprinted and T10p_BEG gene sets and with the ClinGen and GenCC entries for disease entity and mode of inheritance, which were recorded as annotation columns with their source. Carrier HPO terms were compared with the annotated disease entity by word-bounded matching and labeled supportive, nonspecific or not informative. Annotation could draw attention to an allele for review, and the absence of annotation did not count against one.

### Tandem repeats analysis

Tandem repeats were genotyped from coordinate-sorted, haplotagged HiFi alignments using TRGT^75^ (v1.5). Genome-wide genotyping was performed using the Platinum Pedigree tandem-repeat catalogue^76^ (v1.0) (https://zenodo.org/records/13178746). Each allele was summarized by its longest pure segment (LPS)^77^, the longest uninterrupted run of the repeat motif. LPS separates the genuine expansion of a pure array from the length that accumulates through interrupted or degenerate repeats; thus, it provides a consistent measure at loci with no prior disease annotation. For motifs of 1–6 bp, a case allele was retained when its LPS exceeded the 99th percentile of the LPS distribution at the same locus across HPRC_r2^24^ (n=231), APR^13^ (n=53) and family controls (n=107) and then exceeded a fixed 250 bp LPS threshold. Potential hits were identified as repeat alleles exceeding the 99^th^ percentile of the control distribution at a given locus. Repeat loci with insufficient read coverage (<5 reads) were excluded from the analysis. The screening covers the whole catalog rather than a curated set of disease loci, so it can recover expansions at unannotated repeats. A clinically focused analysis was performed in parallel at 68 disease-associated loci, with pathogenic thresholds and modes of inheritance curated from STRipy (Database for Short Tandem Repeats)^78^ and STRhicve^79^. For dominant and X-linked loci, an allele exceeding the curated threshold was considered expanded, and recessive loci required the threshold appropriate to the reported disease mechanism. Candidate pathogenic expansions were reviewed through IGV^51^ snapshots. Calls lacking convincing read-level support were not retained as confirmed expansion.

### SpliceAI analysis

Noncanonical splice-altering variants predicted to affect splicing at positions more than 2 bp from canonical donor or acceptor sites were evaluated separately using the same Splice AI delta score (DS) thresholds. SpliceAI (v1.3) delta scores were computed as the maximum across all four splice consequence directions [DS_AL (acceptor loss), DS_AG (acceptor gain), DS_DL (donor loss) and DS_DG (donor gain)] per variant, and the results were evaluated across three increasingly stringent thresholds: DS > 0.2 (any splicing signal), DS > 0.5 (moderate confidence), and DS > 0.8 (high confidence)^36^. Two input call sets were analyzed in parallel, the read-alignment GRCh38 call set and the GRCh38-deconstructed SNV/indel call set from the pangenome VCF. Variants observed in any of the 107 family controls, 53 APR controls or 231 HPRC controls were removed, followed by filtering against dbSNP, gnomAD, the 1000 Genomes Project, HGSVC and the Greater Middle East Variome. The remaining variants were intersected with the SFARI, SysNDD, TADA, imprinted-gene and brain-expressed-gene lists.

### Mitochondrial analysis

Mitochondrial variants were called from the HiFi alignments using Mitorsaw^80^ (v0.2.14) and run per sample against the GRCh38 no-alt analysis set so that reads of nuclear mitochondrial (NUMT) origin could be identified by remapping and removed before variant calling. Calls were made with a minimum of 3 supporting reads, a minimum alternate allele frequency of 0.20, and the default homopolymer filter enabled, which removes length variants at the control-region poly-C tracts (chrM:303–315, chrM:16,180–16,195) and comparable slippage-prone intervals. Heteroplasmy was defined from the mitorsaw genotype field, which reports one entry per resolved mitochondrial haplotype; a call was heteroplasmic when the per-haplotype entries were not identical and homoplasmic otherwise. The allele fraction alone was not used to define heteroplasmy because homoplasmic calls in homopolymer contexts usually show allele fractions of 0.92–0.96 where reads cannot be unambiguously assigned to the reference or alternate allele. The placeholder position chrM:3106–3107, which has an ‘N’ in GRCh38 to preserve historical rCRS numbering, was removed from all the samples. Heteroplasmic calls were retained at a read depth of ≥100 and a heteroplasmy level strictly between 20% and 80%. The lower bound excludes subthreshold calls, and the upper bound separates genuine heteroplasmy from near-homoplasmic haplogroup variation. Variants at positions carrying any heteroplasmic call in a control sample were excluded to remove positions attributable to population-level polymorphism or haplogroup variation rather than potentially pathogenic somatic or germline events.

Mitochondrial reads were identified by mapping against GRCh38, and candidate NUMTs were filtered out. Mitogenomes were assembled as separate circular contigs using Flye^64^ (v2.9.6), retaining a single circularized mitochondrial contig. Assembly headers followed the PanSN naming convention. A mitochondrial pangenome graph was constructed from the 318 cohort mitogenomes using pggb^81^ (v0.7.4) with the revised Cambridge Reference Sequence (rCRS, NC_012920.1) as a reference path. Variants were normalized using *bcftools norm* and split into small-variant and structural (≥50 bp) call sets. The placeholder interval chrM:3106– 3107 was excluded. Variants were classified as polymorphic (≥2 samples) or singleton. Population novelty was assessed by sequential filtering against dbSNP^29^ (GCF_000001405.40) and gnomAD^30^ (v3.1), retaining at each step only variants unmatched by the preceding resource. Calls falling within documented mitochondrial length-variation hotspots (the HVS-II and HVS-I poly-C tracts, the CA dinucleotide repeat at chrM:514–524, and the poly-C tracts at chrM:568–573 and chrM:5895) were excluded from downstream analysis because apparent novelty at these positions is attributable to indel representation mismatch between graph-derived calls and the reference databases. Novel-variant density was compared across functional compartments after the compartment length was normalized.

### Orthogonal validation: Sanger and ONT

Clinically relevant SNVs and small indels identified by PacBio HiFi sequencing were visually inspected in Integrative Genomics Viewer to confirm read-level support and exclude local alignment artifacts. Selected variants were independently validated by Sanger sequencing using genomic DNA from the proband and, where available, relevant family members. Locus-specific primers were designed to amplify the genomic regions spanning each candidate variant, and the resulting PCR products were subjected to bidirectional Sanger sequencing.

Selected candidate variants requiring independent long-read confirmation, including tandem-repeat expansions, were orthogonally evaluated using Oxford Nanopore Technologies (ONT) LRS. High-molecular-weight genomic DNA was extracted using the Nanobind HMW DNA extraction kit (Pacific Biosciences) according to the manufacturer’s instructions and quantified by Qubit fluorometry.

Approximately 3 µg of genomic DNA was used for library preparation with the ONT Ligation Sequencing Kit V14 (SQK-LSK114). DNA repair and end preparation were performed using the NEBNext Companion Module for Oxford Nanopore Technologies Ligation Sequencing (E7180L; New England Biolabs), followed by AMPure XP bead purification. Sequencing adapters were ligated using the reagents supplied with the ONT library-preparation workflow, and libraries were purified using AMPure XP beads with Long Fragment Buffer.

The final libraries were quantified by Qubit fluorometry, and approximately 300 ng of the prepared library in 32 µl was combined with sequencing buffer and library beads and loaded onto R10.4.1 PromethION flow cells (FLO-PRO114M). Sequencing was performed on the ONT PromethION platform using MinKNOW software (v25.09.16).

### RNA extraction and ONT transcriptome sequencing for splice variant validation

To assess the transcript-level consequences of selected variants predicted to alter splicing by SpliceAI, Oxford Nanopore Technologies (ONT) long-read transcriptome sequencing was performed on peripheral blood RNA from NDD cases. Whole blood was collected in Tempus Blood RNA Tubes, and total RNA was extracted using the Tempus Spin RNA Isolation Kit (Applied Biosystems, Thermo Fisher Scientific; Cat. No. 4380204) according to the manufacturer’s instructions. Polyadenylated RNA was enriched from total RNA using Dynabeads Oligo(dT)25 (Thermo Fisher Scientific), followed by magnetic separation, washing and elution.

Approximately 100 ng of poly(A)-enriched RNA was used for cDNA synthesis using the ONT direct-cDNA sequencing workflow. First-strand synthesis and strand switching were performed using Maxima H Minus Reverse Transcriptase (Thermo Fisher Scientific), followed by RNA degradation and second-strand synthesis. The resulting cDNA was subsequently purified using AMPure XP magnetic beads and subjected to end repair and dA tailing. Where applicable, samples were individually barcoded using ONT native barcodes, pooled and ligated to sequencing adapters using the NEBNext Quick Ligation Module (New England Biolabs). Libraries were purified, quantified using a Qubit fluorometer and sequenced on the ONT Promethion P24.

### Statistical analysis

Statistical analyses were performed in R [v4.5.1] and Python [v 3.13.]. Continuous variables were compared using two-sided/one-sided Mann Whitney U tests or Kruskal Wallis tests as specified; categorical variables were compared using Fisher’s exact test. Pairwise p values were adjusted using the Benjamini Hochberg (BH) procedure. Odds ratios (ORs) and 95% confidence intervals (CIs) are reported for categorical comparisons. All tests, sidedness, correction procedures and sample sizes are additionally specified in the corresponding figure legends.

## Data Availability

The sequencing data, assemblies and pangenome graph generated in this study are available for research use through controlled access to the Dubai Health CATG Flamingo server to preserve the IRB policy and clinical patient’s confidentiality. The unique variants and novel sequences are deposited in Zenodo under accession link https://doi.org/10.5281/zenodo.24009781.

The previously published data used in the study are accessible as follows:

APR pangenome assemblies and pangenome graph (<u>The Arab Pangenome Reference -MBRU</u>)

HPRC Release 2 assemblies and pangenome graph (<u>HPRC Data Explorer</u>)

## Code Availability

The study used previously published analysis tools as described in the Methods section. No custom tools were developed or used in this study.

## Supporting information

Supplementary Results and Figures

## Acknowledgments

We would like to thank all the study participants for giving us the consent to incorporate them as part of this clinical research. Yehia Zakaria provided constant support for the CATG Flamingo computation cluster. Avinash Krishnan, and Freeda Pinto provided constant help with logistics at CATG sequencing lab. Our HPRC consortium for their valuable feedback on our study. Dubai Health (DH) funded this work and supported the procurement of reagents, chemicals, and computing hardware. The funders had no role in the design of the study, data collection, analysis, publishing or preparation of the manuscript. We thank Nature Proof-Editing Services for its linguistic assistance, which greatly aided the preparation of this manuscript.

---

**Extended Figure 1.**
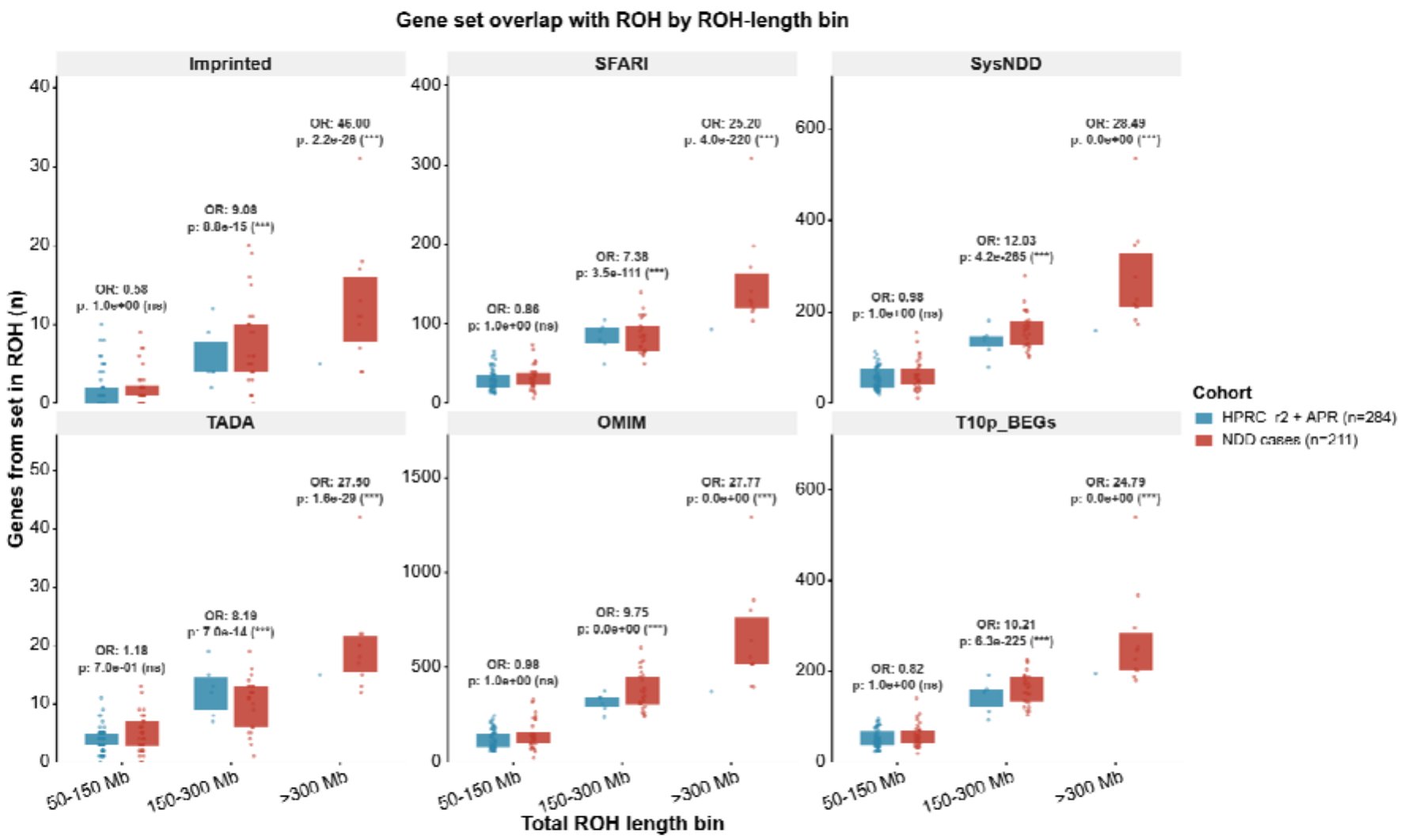
Number of genes from each of six gene sets falling within runs of homozygosity (ROH), shown per individual and stratified by total ROH length (50–150 Mb, 150–300 Mb, > 300 Mb). Panels show imprinted genes, SFARI genes, SysNDD, TADA, OMIM morbid genes and T10p_BEGs (define on first use). Blue denotes the combined HPRC_r2 + APR reference cohort (n = 284) and red the NDD case cohort (n = 211). Each point is one individual, boxes give the median and interquartile range, with whiskers extending to 1.5 × IQR. Odds ratios and p-values above each bin compare cases with references within that ROH-length stratum (test; correction), with significance coded as *** p < 0.001 (BH-corrected) and ns, not significant. The y-axis scale differs between panels and only 1 reference individual carried > 300 Mb of total ROH. In the 50–150 Mb bin the two cohorts are indistinguishable for every gene set (OR 0.58–1.18, all ns). Above 150 Mb, cases show consistent enrichment (OR 7.38–12.03) (except for SFARI and TADA), which strengthens further above 300 Mb (OR 24.79–46.00).

**Extended Figure 2.**
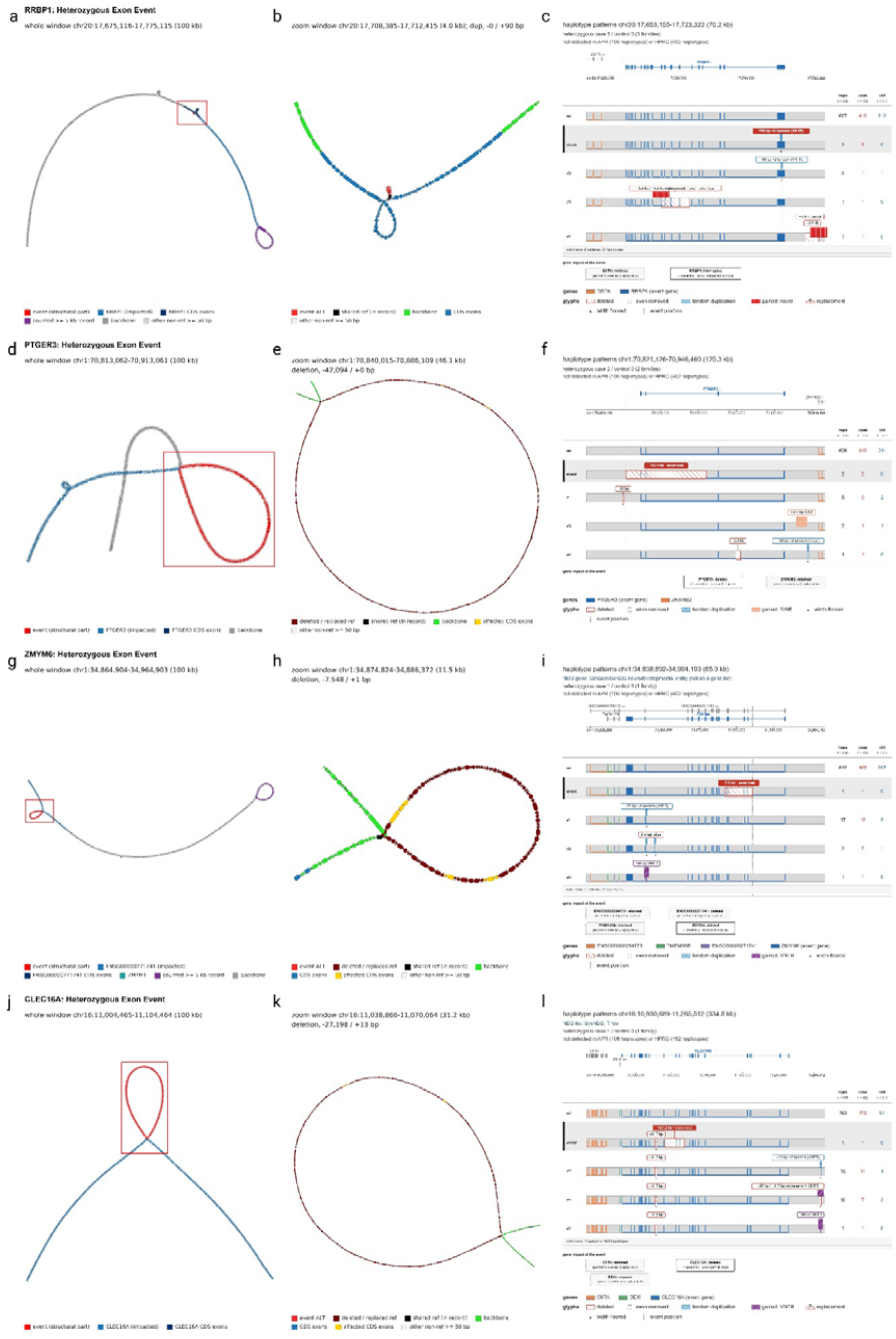
Panel layout, node and glyph colours, and count conventions are as in Figure 5. Each row shows one locus in three views (window, zoom, patterns), with whole window (a, d, g, j) in the left view in all four cases. Counts are n = 636 haplotypes from 211 affected and 107 unaffected individuals, reference panel status is given against APR (106 haplotypes) and HPRC_r2 (462 haplotypes). All four events shown are heterozygous and case-only. The interesting pattern unique to the case is highlighted in gray. (a–c) RRBP1 (chr20). A 90 bp tandem duplication (VNTR) within the gene body interrupting the coding sequence. The allele is carried by 3 affected individuals and by no unaffected individual and is not detected in APR or HPRC. Four further structural patterns are shown, however, either of them are also present in controls or not impacting the exon. (d–f) PTGER3 (chr1). A 42,094 bp deletion removes coding exons, with PTGER3 deleted on the assembly path and the exon path skipped. The allele is carried by 2 affected individuals and no unaffected individuals, and also not detected in APR or HPRC controls. (g–i) ZMYM6 (chr1), a ClinGen/GenCC neurodevelopmental entity not present on the curated gene lists used here. A 7,548 bp deletion removes coding exons, with ZMYM6 deleted and the exon path skipped. The allele is carried by 1 affected individual, and this pattern is not observed in any of the controls. The most recurrent alternative pattern in the cases was found to have an 82 bp VNTR tandem duplication on 17 haplotypes (12 affected, 5 unaffected) (612 haplotypes carry the reference pattern). (j–l) CLEC16A (chr16; SysNDD and T10p_BEGs gene). A 27,198 bp deletion removes coding exons, with CLEC16A deleted and the exon path skipped. The allele is carried by 1 affected individual and no unaffected individual, and also not detected in APR or HPRC. The event haplotype additionally carries a 315 bp deletion shared with the other displayed patterns. In this window a total of 15 patterns were found in which only 183 haplotypes carry the reference pattern.

