## Supplementary Results and Figures for "Pangenome Discoveries from Clinical Long-Read Genomes in Underrepresented Populations with Neurodevelopmental Disorders"

**Table of Contents**

**Supplementary Results**

- *De novo* SNVs and parental phasing
- NDD pangenome graph
- Pangenome nonreference sequence growth
- Global tandem repeat expansions at intronic loci
- Non-canonical splicing variants across delta-score thresholds
- Mitochondrial genome analysis

**Supplementary Figures**

- Supplementary Figure 1
- Supplementary Figure 2
- Supplementary Figure 3
- Supplementary Figure 4
- Supplementary Figure 5
- Supplementary Figure 6
- Supplementary Figure 7
- Supplementary Figure 8
- Supplementary Figure 9
- Supplementary Figure 10
- Supplementary Figure 11
- Supplementary Figure 12
- Supplementary Figure 13
- Supplementary Figure 14a
- Supplementary Figure 14b
- Supplementary Figure 15
- Supplementary Figure 16

**Supplementary Results**

***De Novo* SNVs and parental phasing**

A total of 6,053 *de novo* mutations (DNMs) were identified across 53 trios (42 NDD probands and 11 unaffected siblings), comprising 4,687 single nucleotide variants (SNVs) and 1,366 insertions/deletions (INDELs). The mean number of *de novo* SNVs per proband was 88.4 ± 18.1, with no significant difference between NDD cases (88.6 ± 19.2) and controls (87.7 ± 13.4) (Mann–Whitney U = 228, P = 0.96). *De novo* INDELs were slightly more frequent in controls (30.5 ± 9.5) than in cases (24.5 ± 7.3), though this difference did not reach statistical significance (Mann–Whitney U = 146, P = 0.06). Read-based haplotype phasing using flanking inherited heterozygous SNVs successfully determined parental origin for 90.1% (4,225/4,687) of *de novo* SNVs. Among phased SNVs, 78.0% (3,295/4,225) were of paternal origin and 22.0% (930/4,225) were of maternal origin, yielding a paternal-to-maternal ratio of 3.5:1. This paternal bias was significant in both NDD cases (Wilcoxon signed-rank test, P = 8.23 × 10⁻⁹) and controls (P = 4.88 × 10⁻⁴), with mean paternal fractions of 77.5% and 79.5%, respectively (Figure 1A). The magnitude of paternal bias did not differ significantly between cases and controls (Fisher's exact test, OR = 0.88, P = 0.168). An additional 37 SNVs (0.8%) were classified as post-zygotic mosaic (PZM), and 425 SNVs (9.1%) remained unphased due to insufficient flanking heterozygous markers. For *de novo* INDELs, parental origin was found to be 70.9% (968/1,366) of variants. Among phased INDELs, 58.3% (564/968) were paternal and 41.7% (404/968) were maternal (paternal-to-maternal ratio = 1.40:1). The paternal bias for INDELs was significant in both NDD cases (Wilcoxon signed-rank test, P = 0.004) and controls (P = 0.024), though it was markedly weaker than the bias observed for SNVs (Figure 1B). A total of 336 INDELs (24.6%) were classified as PZM, and 62 (4.5%) remained unphased.

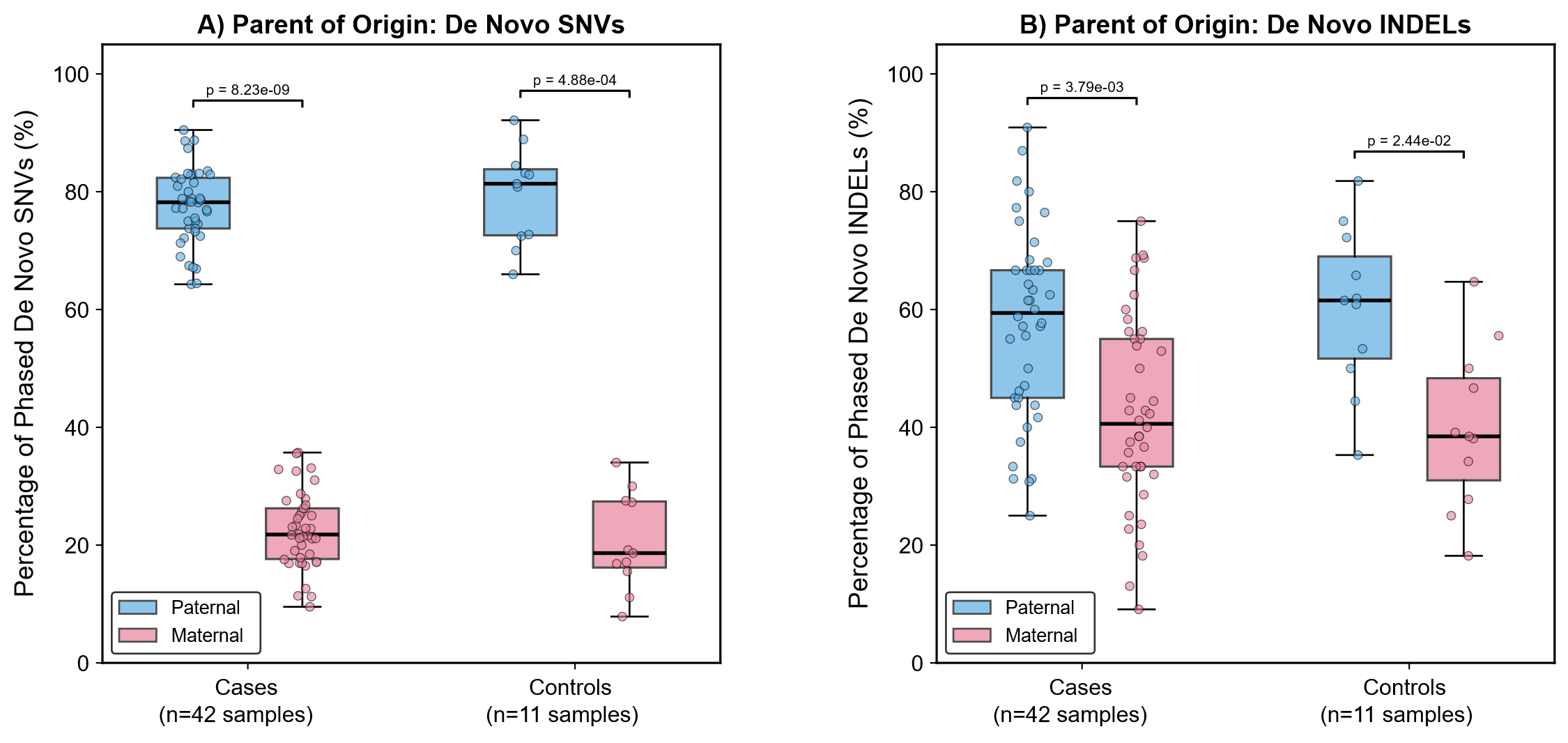

**Figure 1. Parent-of-origin distribution of phased *de novo* SNVs and INDELs in NDD cases and controls. (A)** Per-proband percentage of *de novo* SNVs assigned to paternal (blue) or maternal (pink) origin by read-based haplotype phasing. Among 4,225 phased SNVs, 78.0% were paternal and 22.0% were maternal (paternal-to-maternal ratio = 3.54:1). The paternal bias was significant in both NDD cases (n = 42; Wilcoxon signed-rank test, P = 8.23 × 10⁻⁹) and unaffected sibling controls (n = 11; P = 4.88 × 10⁻⁴). **(B)** Per-proband percentage of *de novo* INDELs assigned to paternal or maternal origin. Among 968 phased INDELs, 58.3% were paternal and 41.7% were maternal (ratio = 1.40:1). The paternal bias was significant in cases (P = 0.004) and controls (P = 0.024) but was substantially weaker than the bias observed for SNVs. Box plots show medians and interquartile ranges; individual data points represent single probands.

Among all DNMs, 81 were located in exonic regions, of which 80 were SNVs and 1 was an INDEL. The number of exonic DNMs was similar between NDD cases (64 variants; 1.52 per proband) and controls (17 variants; 1.55 per proband). Among phased exonic variants in cases, 48 were paternal and 10 were maternal, representing a significant paternal bias (binomial test, P < 0.0001). In controls, 11 were paternal and 5 were maternal, though this difference was not statistically significant (binomial test, P = 0.210) (Figure 2A).

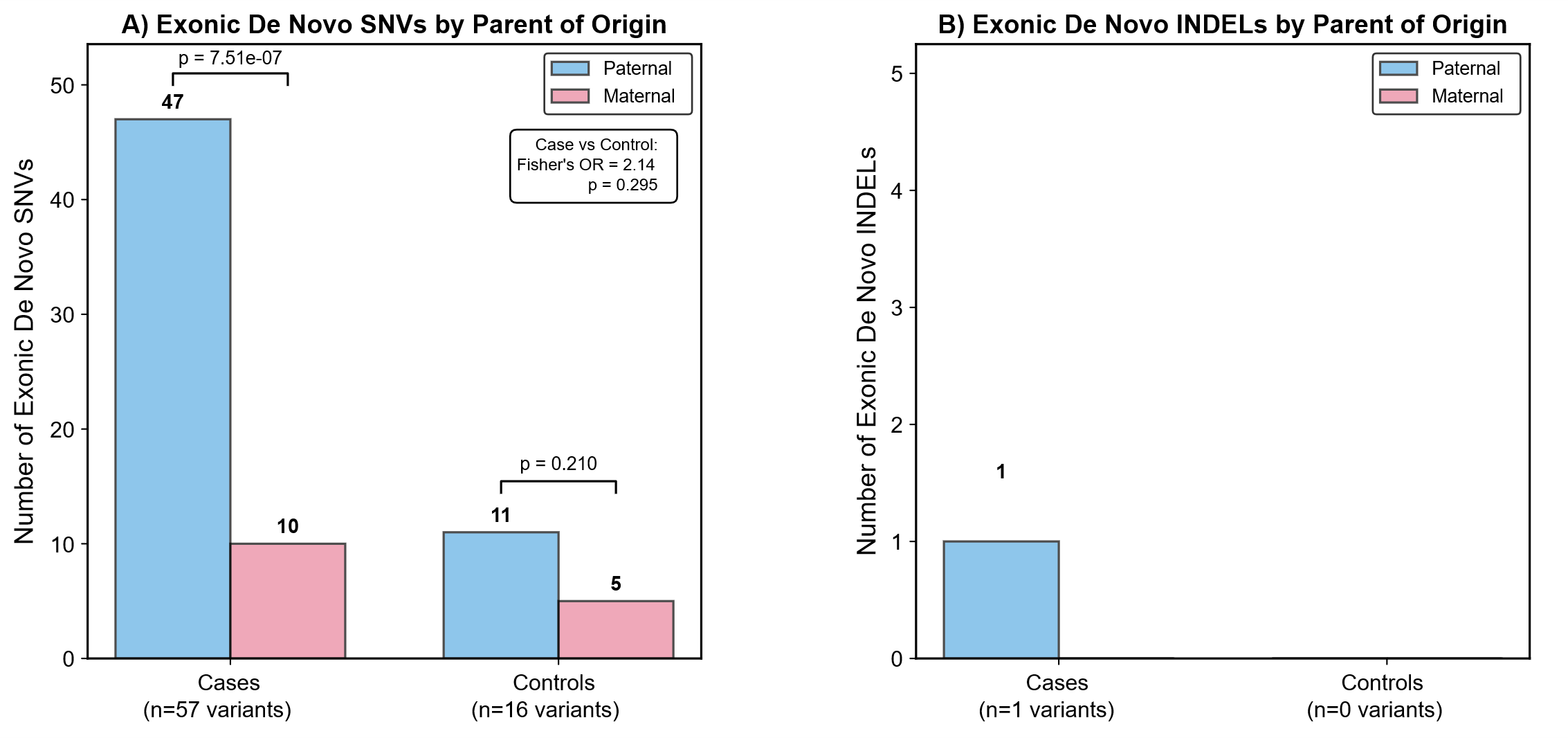

**Figure 2. Parent-of-origin of phased exonic *de novo* variants in NDD cases and controls. (A)** Number of exonic *de novo* SNVs of paternal (blue) and maternal (pink) origin. Among cases, 48 of 58 phased exonic SNVs were paternal (binomial test, P < 0.0001), whereas controls showed 11 paternal and 5 maternal (P = 0.210). **(B)** Number of exonic *de novo* INDELs by parent of origin. Only one exonic INDEL was detected across the cohort. Fisher’s exact test was used to compare the paternal-to-maternal ratio between cases and controls. Numbers above each bar indicate variant counts.

**NDD pangenome graph**

A population-scale pangenome graph as described in the main text was constructed using the Minigraph-Cactus pipeline (cactus-pangenome), integrating haplotype-resolved assemblies from 318 NDD cohort individuals (636 haplotypes) against T2T-CHM13 v2.0 and GRCh38 reference coordinates. The cohort comprised 211 NDD cases and 107 family controls, with the remaining five individuals contributing assemblies to graph construction. Graph clipping applied a minimum node frequency of nine haplotypes, retaining sequence present in at least 1.4% of input haplotypes in the clipped graph; the full unclipped graph was retained for downstream analyses. External population controls comprised 53 APR and 231 HPRC R2 samples. Downstream case–control analyses compared 211 NDD cases (422 haplotypes) with 391 controls (782 haplotypes), including cohort family controls and external population controls.

To our knowledge, this represents one of the largest disease-focused human pangenome graphs constructed to date and integrates long-read haplotype assemblies from NDD-affected individuals across Africa, Central Asia, the Middle East and Central/South Asia alongside population controls. The graph contained 149,222,686 nodes and 206,562,335 edges, with an edge-to-node ratio of 1.38 and 3.699 Gb of embedded sequence, including the reference backbone. Deconstruction of the graph against T2T-CHM13 coordinates yielded 37,445,437 variant records across the 318 cohort samples: 31,603,972 SNPs, 1,289,709 MNPs, 5,414,612 indels and 356,678 complex records, of which 2,868,133 were multiallelic. Applying a size threshold of ≥50 bp identified 173,869 SV-class records, 133,040 of which were multiallelic. Because a multiallelic SV site can carry alleles of different classes, per-allele classification at these sites assigns 157,666 indels, 79,853 SNPs at multiallelic SV sites, 57,792 MNPs and 76,855 complex events. These overlapping record categories reflect the representation of complex and multiallelic variation in the graph-derived call set.

**Pangenome nonreference sequence growth**

Growth-curve analysis using panacus quantified accumulation of 567.8 Mb nonreference sequence across 636 assembly haplotypes (Figure 2d; Supplementary Table 10). The first 100 individuals captured 57.6% (326.8 Mb) of the final total, rising to 81.5% (462.5 Mb) at 200 individuals and 97.8% (555.5 Mb) at 300, the final 18 individuals contributed a further 12.3 Mb (2.2%). The mean rate of new sequence addition declined 5.6-fold over the cohort, from 4.18 Mb per individual across the first 50 individuals to 0.75 Mb per individual across the final 50. Sequence discovery is therefore decelerating but has not plateaued, and the curve shows no indication of an asymptote at this sample size. Stratifying this sequence by the fraction of individuals carrying it shows that the great majority of it is rare. Only 311.6 kb is near-core, present in ≥95% of individuals, and this component is fixed within the first nine individuals sampled. A further 81.8 Mb is shared by 5–95% of individuals, while 485.6 Mb (85.5%) of all nonreference sequence is carried by fewer than 5%. Nonreference content in this cohort is thus overwhelmingly rare or haplotype-private, and the continued rise of the growth curve is driven almost entirely by this low-frequency fraction rather than by the accrual of new common sequence. Together, these results demonstrate substantial nonreference sequence diversity and continued discovery of sequence with increasing haplotype sampling, while highlighting the contribution of rare sequence to the assembled pangenome.

**Global tandem repeat expansions at intronic loci**

A parallel screen of intronic loci in the same genome-wide TRGT catalogue identified 230 expanded alleles with an LPS length ≥250 bp, distributed across 137 distinct loci on all 22 autosomes and chromosome X in 148 individuals (Supplementary Figure 15 and 16; Supplementary Table 18b–c). This is an order of magnitude more than the exonic screen (16 alleles at 8 loci), consistent with the far greater intronic representation in both the genome and the TRGT catalogue. Expansions were predominantly private, 92 of the 137 loci (67.2%) were observed in a single individual, 45 in two or more and 12 in four or more, while 53 of the 148 carriers show expansions at more than one intronic locus. Motifs were AT-biased, with 34 loci carrying purely A/T motifs and a mean motif A/T content of 68.8%.

Ten loci recurrent in 4–7 individuals were examined in detail (Supplementary Figure 16). The longest alleles were observed at chr16:32,805k–32,808k ([TTCAT], 5-mer; n = 7), where total allele lengths reached 6,611–6,647 bp which is roughly twice the longest allele at any other recurrent locus and 36.8–37.1% of each allele consisted of interruptions. Notably, the LPS at this locus was invariant at 280 bp across all seven carriers, so the length difference between individuals resides almost entirely in the interrupted, impure portion of the tract rather than in the pure core. The opposite pattern was seen at chr9:110,701k ([TTTAT], 5-mer; n = 6), which showed the widest absolute spread in pure-tract length, with LPS ranging from 670 to 4,285 bp (6.4-fold); chr18:70,211k ([AAAAT], 5-mer; n = 5) showed a comparable spread in relative terms (420–3,260 bp, 7.8-fold). Across the catalogue, the single largest sssLPS was a 5,130 bp [AAATA] expansion at chr5:9,426k, observed in one individual. The functional significance of these intronic expansions is not established, but their size, their recurrence across unrelated case individuals from diverse ancestral backgrounds, and the contrast between interruption-driven and pure-tract-driven length variation warrant further investigation, particularly at loci within or proximal to NDD-relevant genes.

**Non-canonical splicing variants across delta-score thresholds**

The main results section reports case-level detection at the high-confidence threshold and the distribution of variant counts across DS bins, the intermediate steps plotted in Figure 6i are described here. Extending case-level detection to the two permissive thresholds shows that the advantage of pangenome alignment scales with stringency rather than representing a fixed offset. At DS > 0.2, pangenome alignment identified at least one non-canonical splicing variant in 210 of 211 cases (99.5%) compared with 181 cases (85.8%) by read-based alignment a 1.16-fold difference, with 29 cases detected only by the pangenome and none only by read-based alignment (McNemar with continuity correction, BH-adjusted p = 2.4 × 10⁻⁷). At DS > 0.5 these fell to 171 cases (81.0%) and 111 cases (52.6%), a 1.54-fold difference with 63 versus 3 discordant cases (BH-adjusted p = 1.1 × 10⁻¹²). The corresponding separation at DS > 0.8 is 2.74-fold, with 73 versus 5 discordant cases (BH-adjusted p = 2.0 × 10⁻¹³) (Supplementary Table 19). Read-based detection therefore declines far more steeply across this range: 21.5% of cases positive by read-based alignment at DS > 0.2 remain positive at DS > 0.8, compared with 51.0% of cases positive by pangenome alignment.

Restricting to the curated NDD gene panels gives the same pattern at the permissive thresholds: 197 versus 146 cases at DS > 0.2 (93.4% versus 69.2%; 53 versus 2 discordant) and 78 versus 33 cases at DS > 0.5 (37.0% versus 15.6%; 50 versus 5 discordant), each significant after BH-adjustment across the six comparisons in Supplementary Table 19. A decline in detection rate with increasing DS is expected at non-canonical positions, where deep intronic and exonic splicing effects are predicted with lower confidence than variants at canonical donor or acceptor dinucleotides; what the threshold sweep adds is the difference in the rate of that decline between the two alignment strategies. The main text establishes the directional result at the level of variant counts per DS bin, and the sweep shows that it holds at the level of affected individuals. It also shows that at DS > 0.2 both approaches assign a non-canonical prediction to the majority of cases, so that threshold carries little discriminative value in this cohort.

**Mitochondrial genome analysis**

Sixteen of the 99 candidate novel variants (16.2%) occurred within known mitochondrial length-variation hotspots, including the HVS-II poly-C tract (chrM:303–315) and HVS-I poly-C tract (chrM:16,180–16,195), the CA dinucleotide repeat as chrM:514–524, and poly-C tracts at chrM:568–573 and chrM:5,895. These calls accounted for 268 of 368 carrier observations (72.8%) and 12 of 21 non-singleton candidates, including a single 2-bp insertion at chrM:309 found in 142 individuals (44.7%). Given the known variability of these regions, differences in INDEL representation between the graph-derived call set and reference databases is a more parsimonious explanation for their apparent novelty. These 16 calls were therefore excluded from the population-novel set, removing 12 of the 21 recurrent candidates and raising the singleton fraction from 78.8% (78/99) to 89.2% (74/83).

**Supplementary Figures**

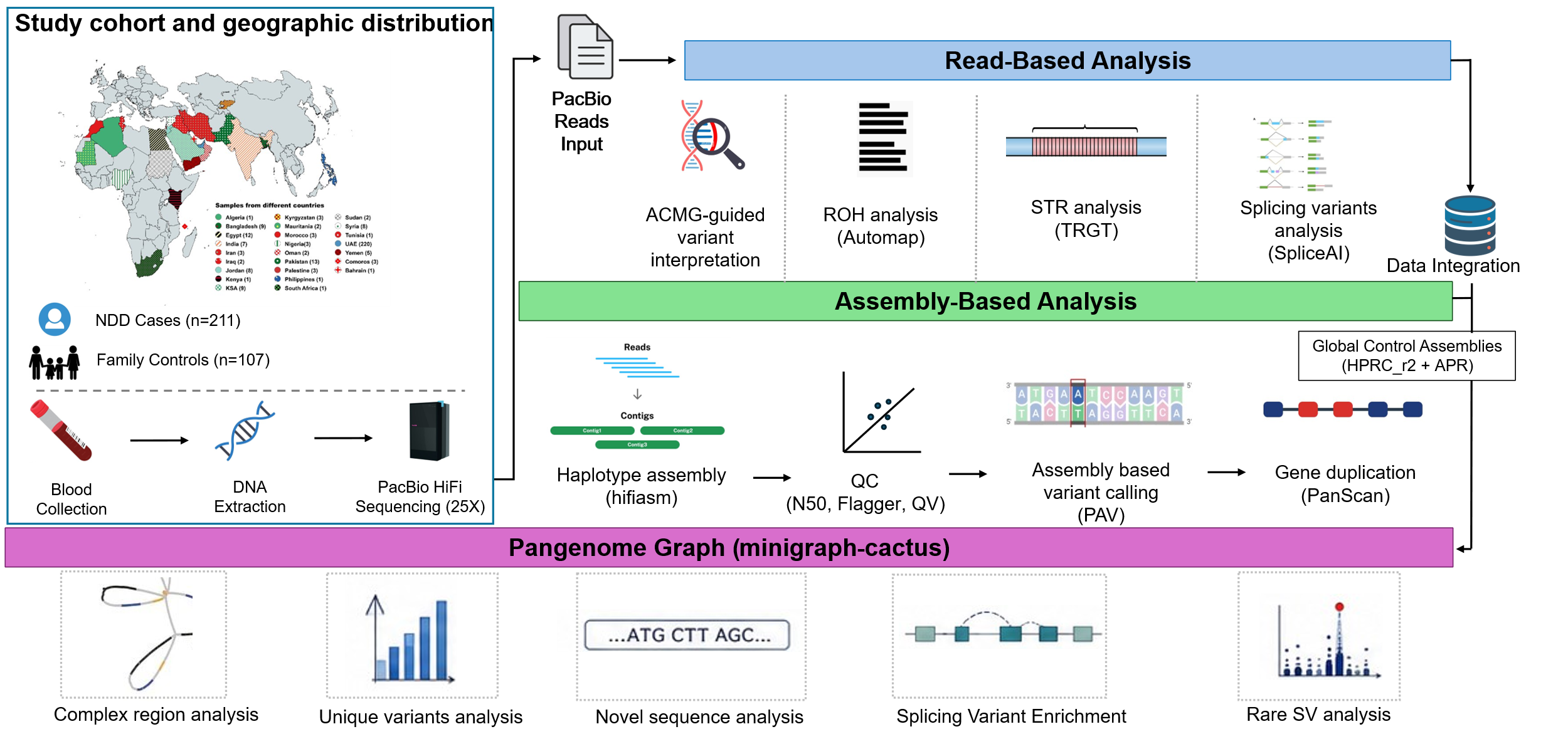

**Supplementary Figure 1.** Study cohort and geographic distribution (*left*). Geographic origin of the study cohort, with countries of recruitment shaded on a world map and the number of individuals contributed by each country given in parentheses in the inset key. The cohort comprised 211 NDD cases and 107 unaffected family members recruited as controls. Blood was collected from all participants, high-molecular-weight DNA extracted, and whole genomes sequenced on the PacBio HiFi platform (LRS). Read-based analysis (*top*). HiFi reads from every individual were processed through four parallel analyses. (i) ACMG-guided interpretation of clinically relevant variants. (ii) Runs-of-homozygosity (ROH) detection with using AutoMap. (iii) Tandem repeats expansion with TRGT. (iv) Identifying non-canonical splice-altering variants with SpliceAI. Assembly-based analysis (*middle*). The same reads were assembled into haplotype genomes using hifiasm. Assemblies passing quality control (contig N50, Flagger misassembly detection, and consensus accuracy QV) were taken forward for assembly-based variant calling with PAV and for gene duplication analysis with PanScan followed by pangenome construction Minigraph-Cactus. Publicly available haplotype-resolved assemblies from the HPRC_r2 and the APR were included as global control assemblies at this stage (*box, right*). Pangenome graph-based analyses (*bottom*). Five analyses were performed on the resulting graph. (i) Characterisation of complex rearrangements. (ii) Identification of cohort specific novel variants relative to public reference panels. (iii) Novel sequence discovery from SV insertions. (iv) Enrichment testing of non-canonical splicing-relevant variation. (v) Detection and visualization of rare structural variants.

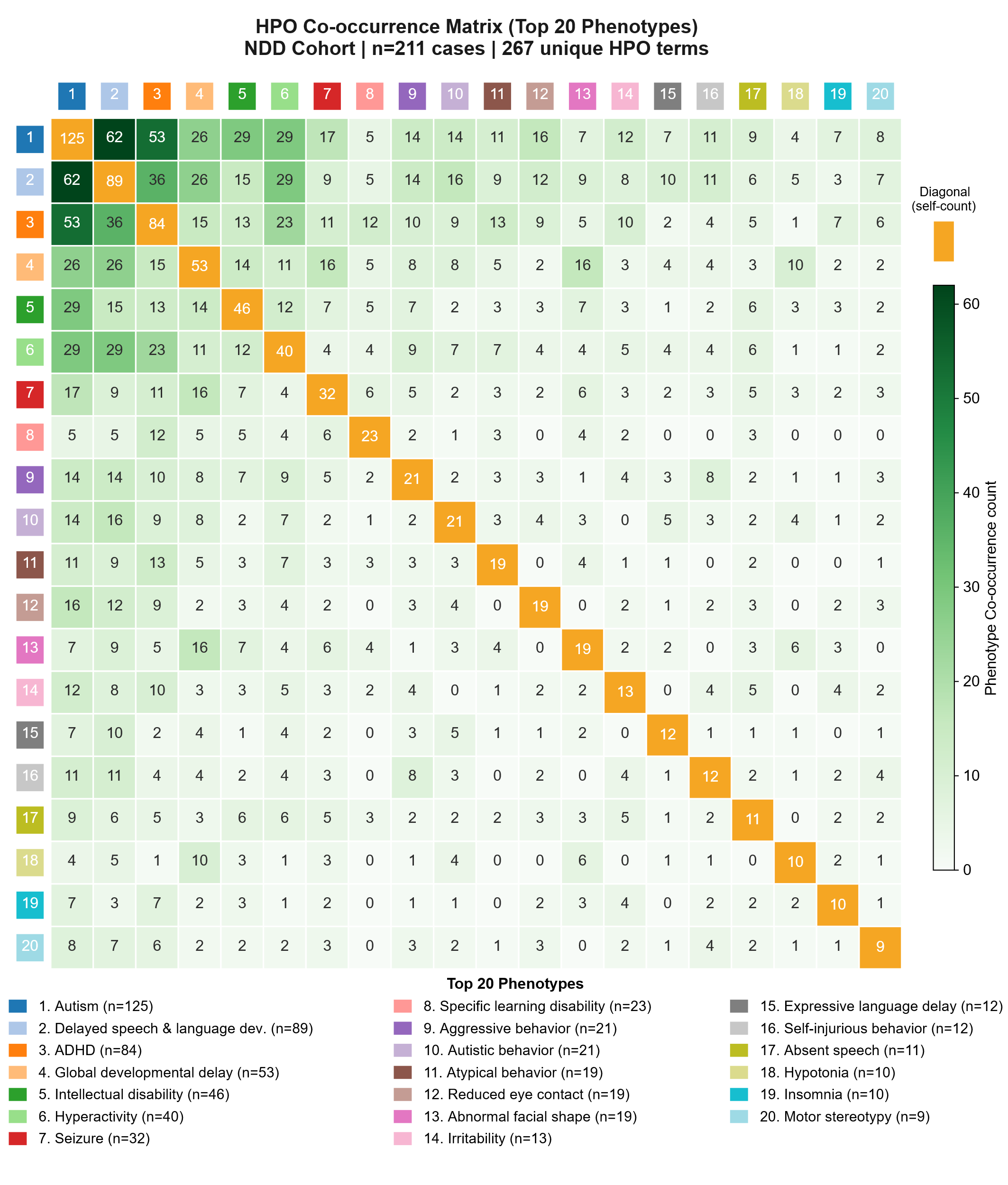

**Supplementary Figure 2.** Co-occurrence of the 20 most frequent HPO terms across the NDD cases, ordered by decreasing cohort frequency and numbered 1-20 in the key below (individual counts in parentheses, range n=125 to n=9). Diagonal cells (orange) give the number of individuals carrying each term, off-diagonal cells give the number carrying both terms of the corresponding pair, shaded on the white-to-green scale at right. HPO identifiers for all 20 terms and the complete phenotype co-occurrence matrix across all recorded terms are provided in Supplementary Table 2b.

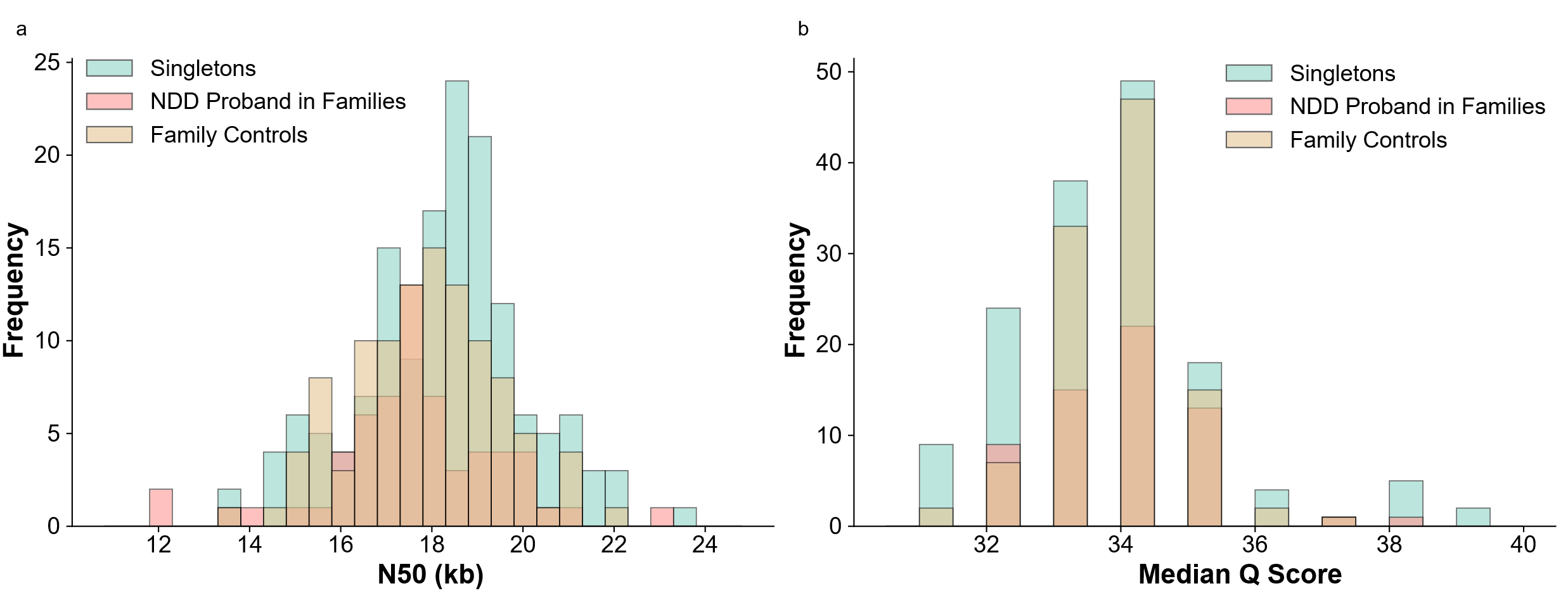

**Supplementary Figure 3. (a)** Distribution of per-sample HiFi read length N50, in kb. **(b)** Distribution of per-sample median HiFi read quality (Q score). In both panels each sample contributes a single value, bars are [stacked/overlaid], and samples are colored by group [singleton cases (teal), NDD probands recruited as part of families (pink), and family controls (tan)]. Read N50 was centered near 18 kb and median read quality near Q34 across all three groups.

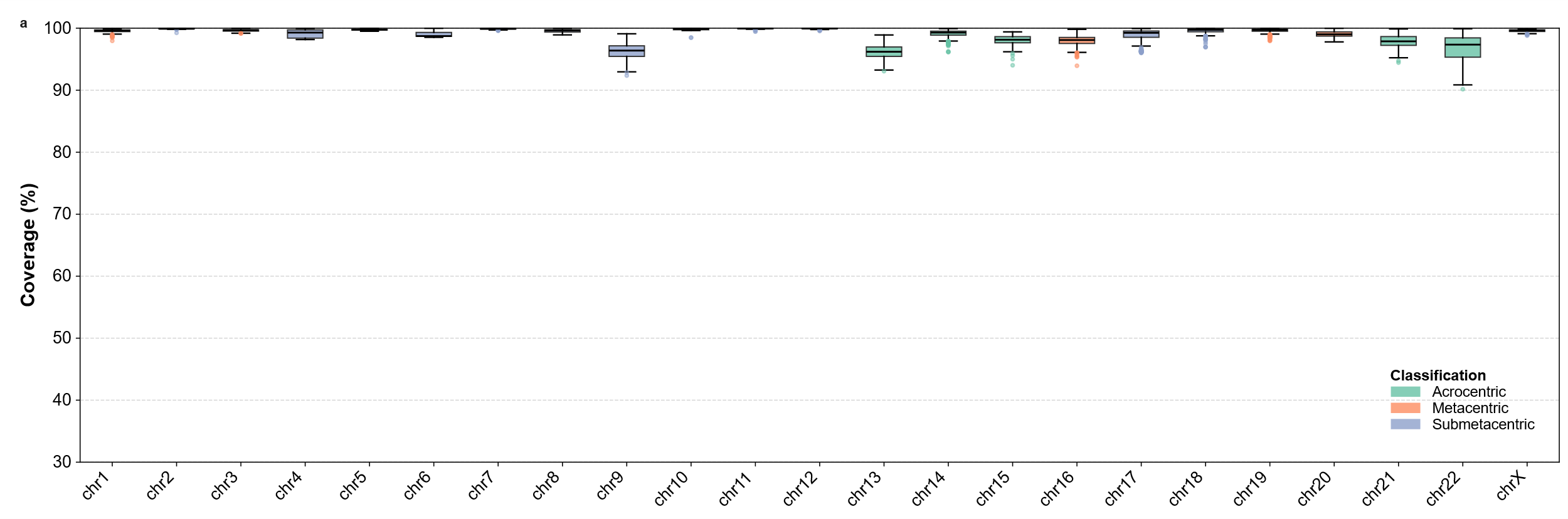

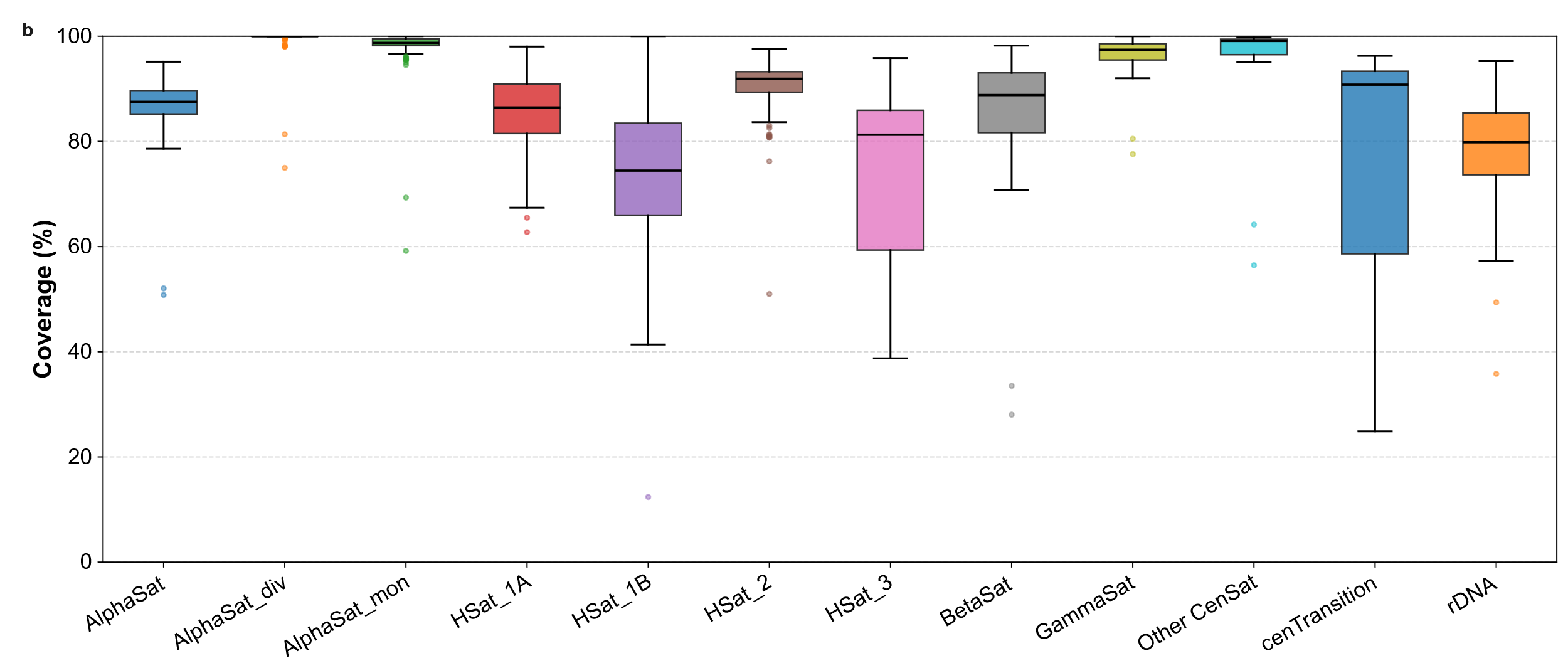

**Supplementary Figure 4. (a)** Per-chromosome coverage of the sequencing reads against T2T-CHM13v2.0 expressed as the percentage of reference bases covered. Chromosomes are colored by centromere morphology [acrocentric (green), metacentric (orange) and submetacentric (purple)]. Boxes span the interquartile range with the median marked, whiskers extend to 1.5× IQR, and points beyond that are plotted individually. Coverage exceeded 95% for most chromosomes. **(b)** Coverage across centromeric and pericentromeric repeat classes annotated in T2T-CHM13, alpha satellite in its higher-order, divergent and monomeric forms (AlphaSat, AlphaSat_div, AlphaSat_mon); human satellite families 1A, 1B, 2 and 3 (HSat_1A, HSat_1B, HSat_2, HSat_3); beta satellite (BetaSat); gamma satellite (GammaSat); remaining centromeric satellite annotations (Other CenSat); the centromeric transition region flanking the higher-order array (cenTransition); and ribosomal DNA arrays (rDNA). Boxes plot drawn as in (a). Coverage was high and tightly distributed for higher-order and monomeric alpha satellite, gamma satellite and other centromeric satellite, and markedly lower and more variable for HSat_1B, HSat_3, the centromeric transition region and rDNA. Note the differing y-axis ranges between panels (a) and (b).

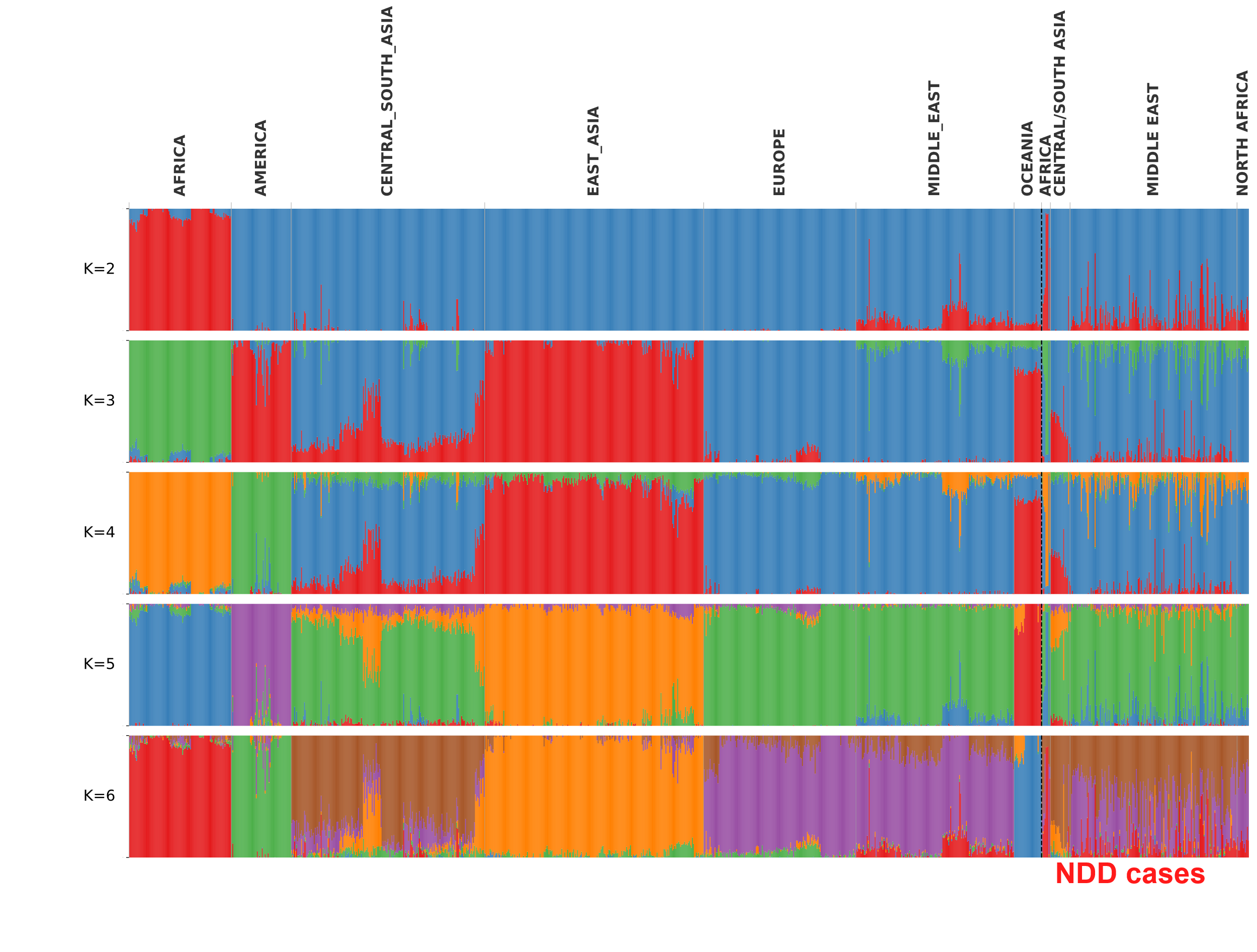

**Supplementary Figure 5.** Genome-wide ancestry inference for the NDD cases against global reference populations. ADMIXTURE ancestry proportions for NDD case individuals analysed jointly with 929 global reference individuals from HGDP, at K=2 through K=6 (top to bottom). Each vertical bar represents one individual and each color one inferred ancestral component, with bar height partitioned according to that individual’s estimated proportion of each component (y axis, 0 to 1). Individuals are ordered by population and grouped as labelled above the plot, reference populations appear to the left of the dashed line and the NDD cases to the right, subdivided into its four constituent regional groups (Africa, Central/South Asia, Middle East, and North Africa).

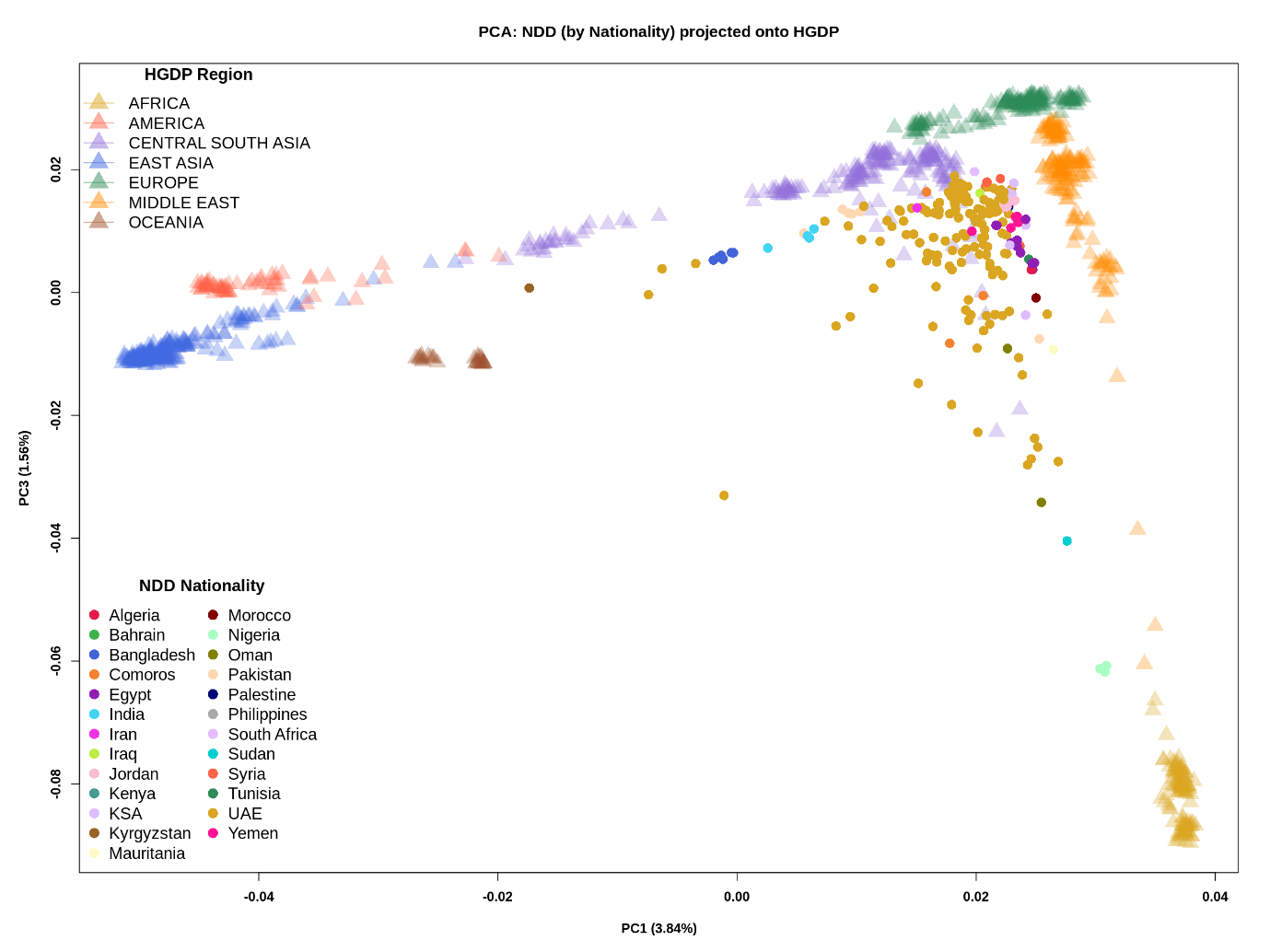

**Supplementary Figure 6:** Projection of samples on HGDP reference principal components, with NDD samples colored by nationality. This higher-resolution view complements the broad superpopulation representation shown in Figure 1f.

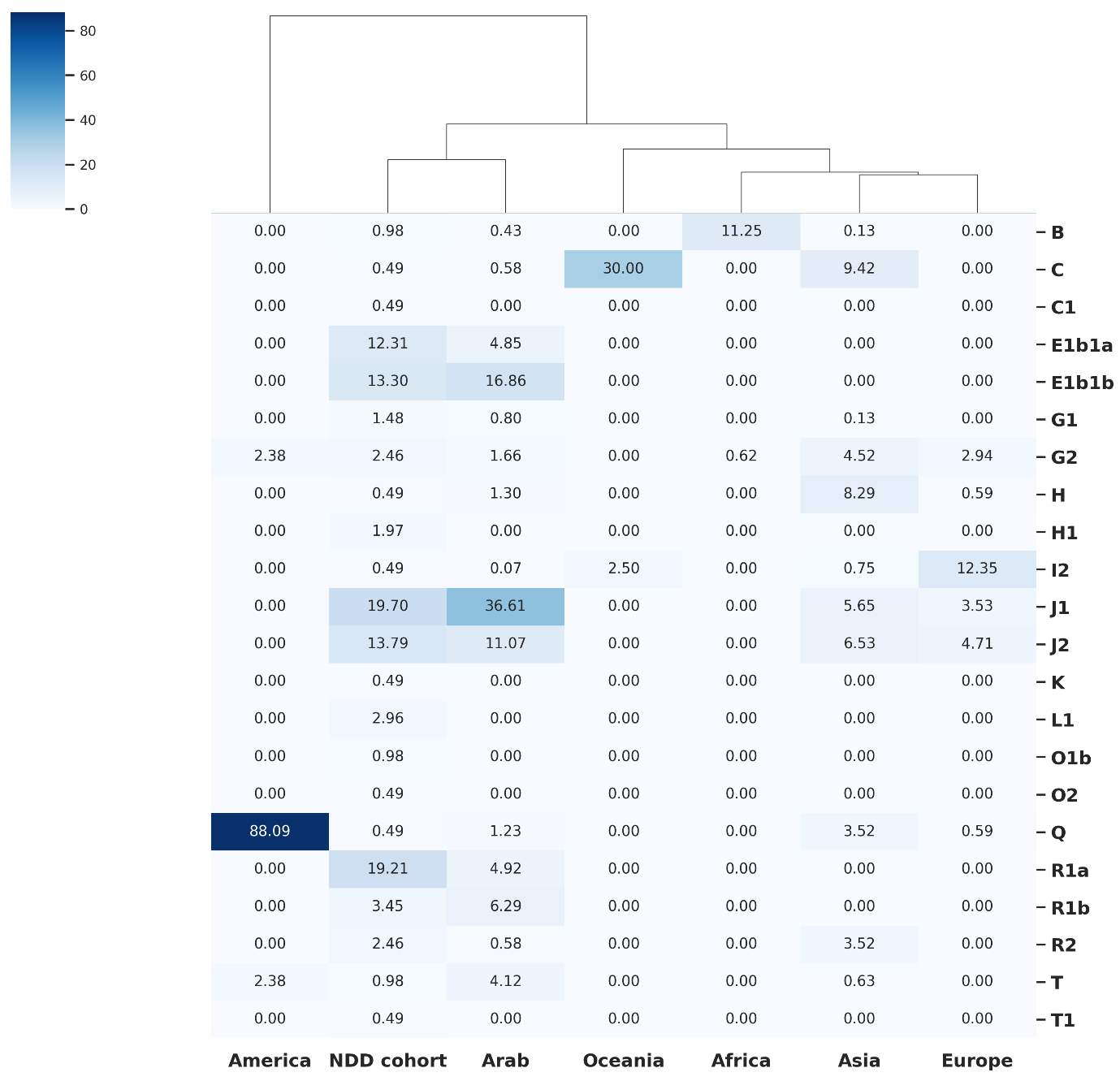

**Supplementary Figure 7.** Y-chromosome haplogroup composition of the NDD cohort and reference population groups. Heatmap of Y-chromosome haplogroup frequencies among male individuals (n=203) in the NDD cohort, grouped by region. Rows are Y haplogroups assigned with Y-LineageTracker where columns are population groups. Cell values give the percentage of males within each group assigned to that haplogroup, so each column sums to 100% (color intensity scales with percentage as shown at *top left*). The dendrogram above the heatmap shows hierarchical clustering of population groups by haplogroup composition. The NDD cohort clustered with the Arab reference group, with J1, J2, E1b1a and E1b1b together accounting for most cohort lineages, consistent with the Middle Eastern and North African origins of most participants.

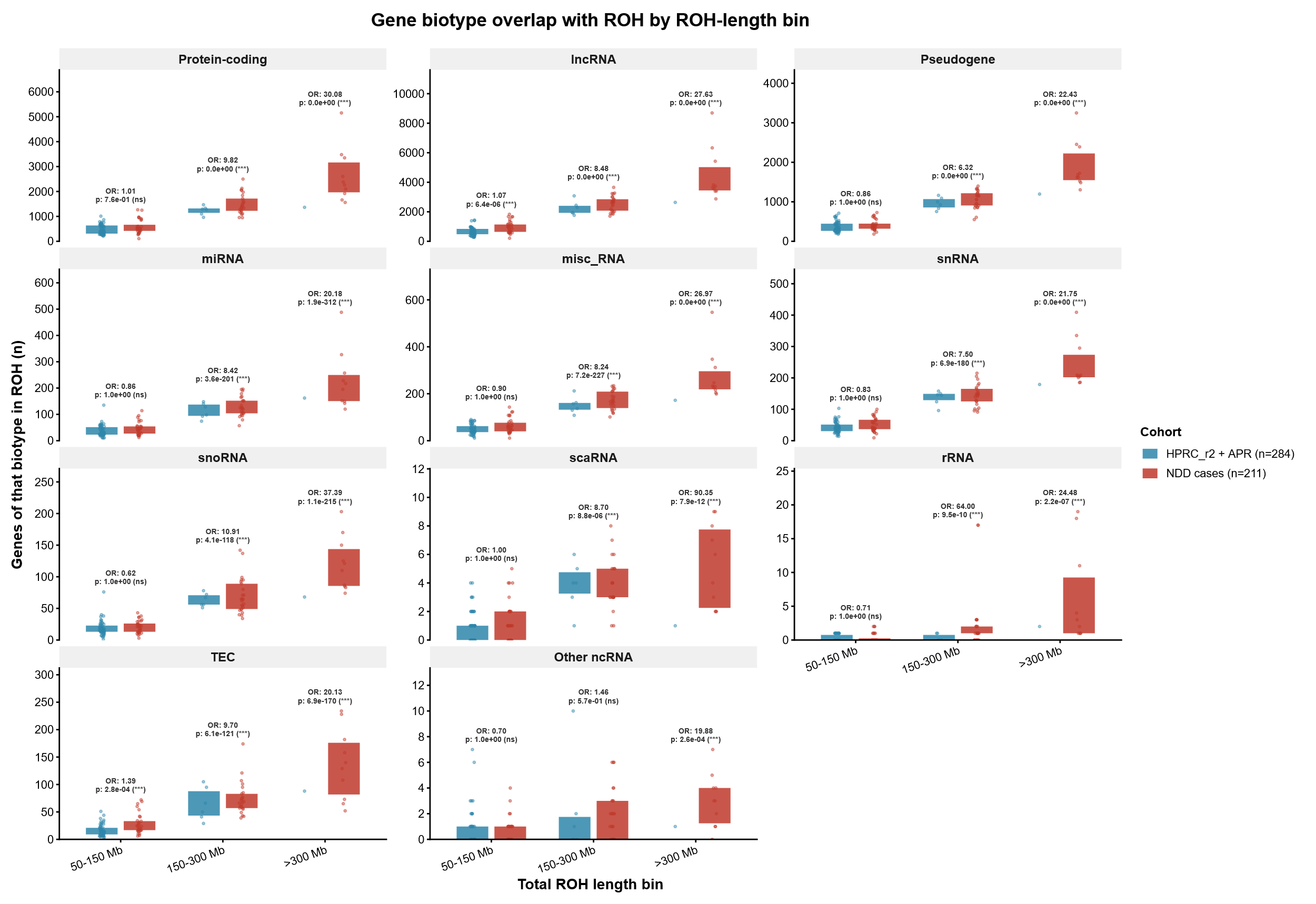

**Supplementary Figure 8.** Gene content within runs of homozygosity, by gene biotype and total ROH burden. Number of distinct genes of each biotype falling within ROH, in 211 NDD cases (red) and 284 combined reference individuals from HPRC_r2 and the APR (blue). Individuals are stratified by total genome-wide ROH burden into three bins (50–150 Mb, 150–300 Mb, >300 Mb). Each panel corresponds to one gene biotype; protein-coding, long non-coding RNA (lncRNA), pseudogene, microRNA (miRNA), miscellaneous RNA (misc_RNA), small nuclear RNA (snRNA), small nucleolar RNA (snoRNA), small Cajal body-specific RNA (scaRNA), ribosomal RNA (rRNA), to-be-experimentally-confirmed loci (TEC), and remaining non-coding RNA classes (Other ncRNA). Overlaid points on the boxplot are individual samples. Odds ratios (OR) and Benjamini-Hochberg (BH) adjusted p-values above each bin are from one-tailed Fisher’s exact tests (NDD > HPRC_r2+APR). Note that y-axis scales differ between panels.

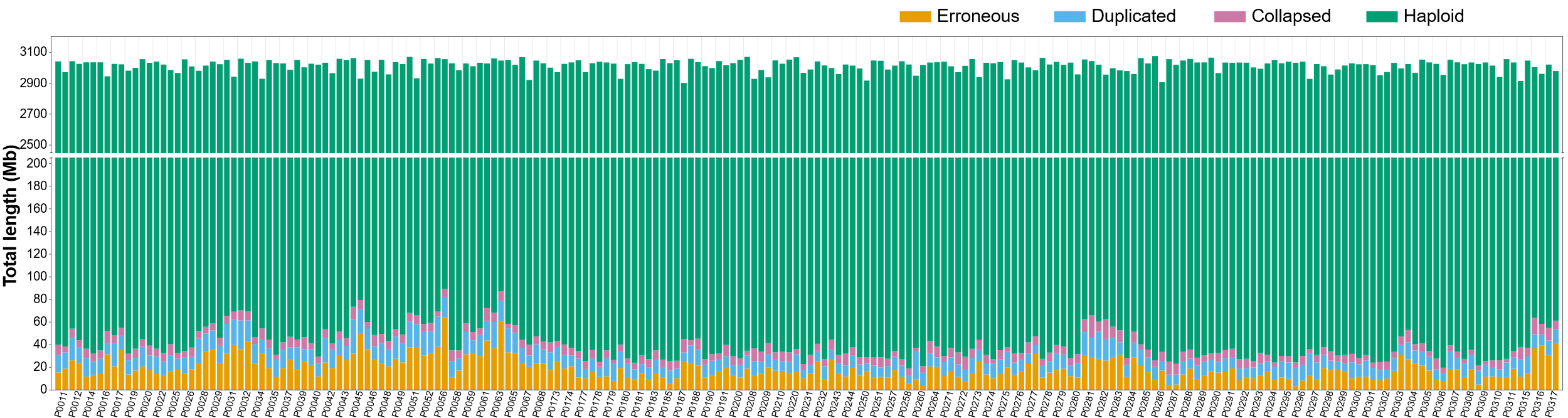

**Supplementary Figure 9.** HMM-Flagger classification of haplotype assemblies from the family controls (214 haplotypes), partitioned into haploid (green), erroneous (dark yellow), duplicated (blue) and collapsed (pink) states. Each vertical bar represents one haplotype assembly, labelled by sample identifier, the two haplotypes of each individual are shown (adjacently). Bar segments give the total length of sequence (Mb) assigned to each state. Haploid sequence accounted for approximately 2.9-3.1 Gb per haplotype.

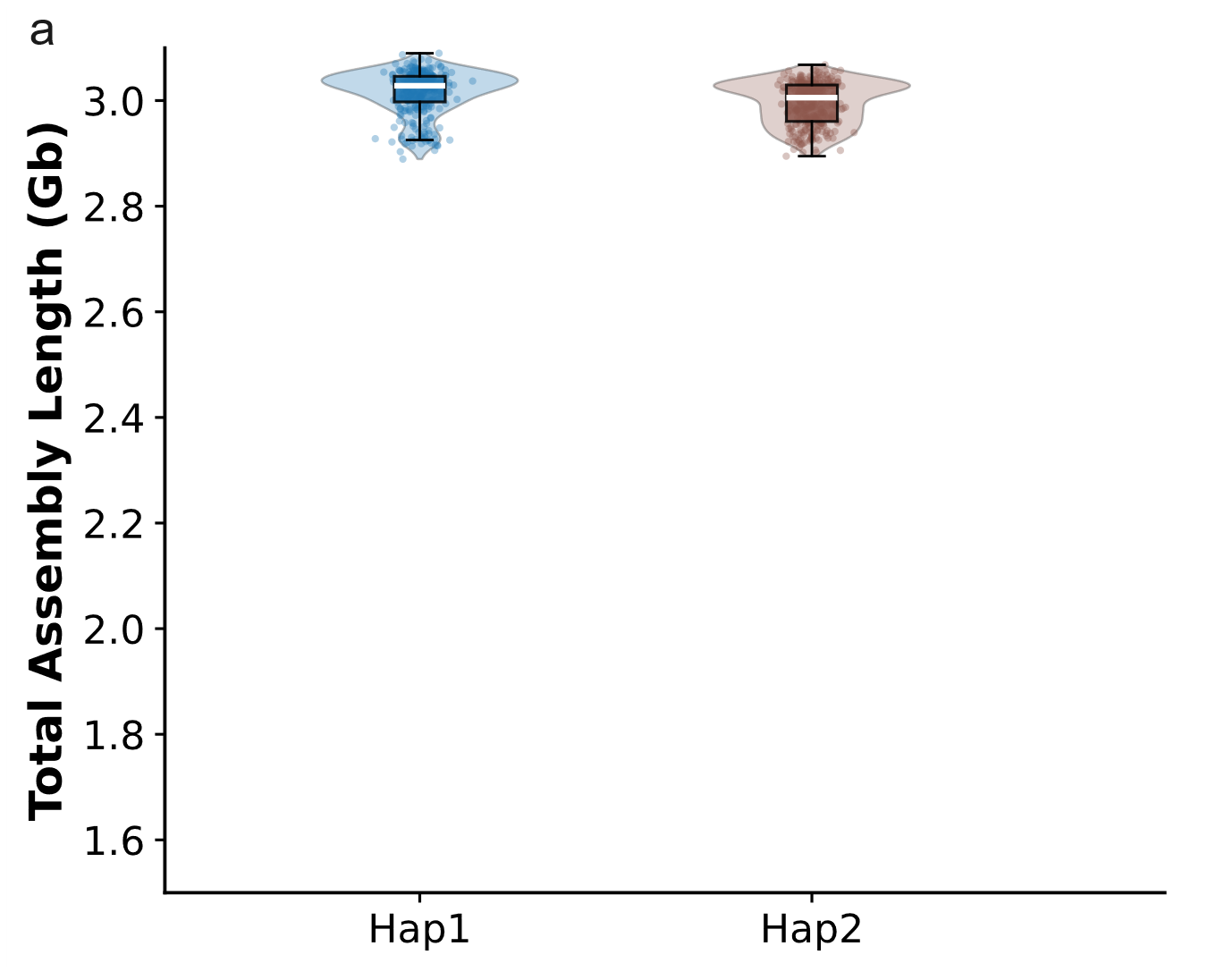

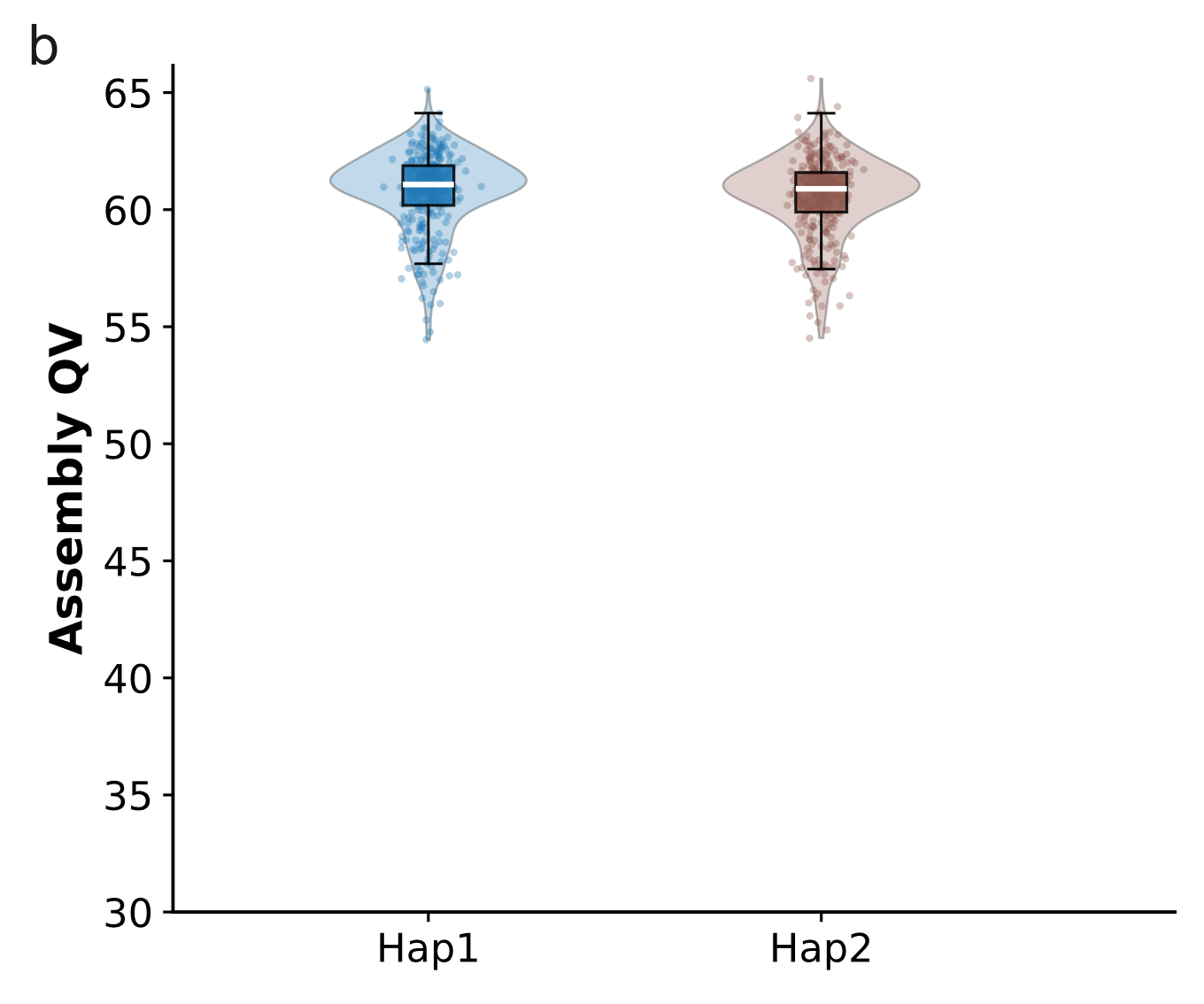

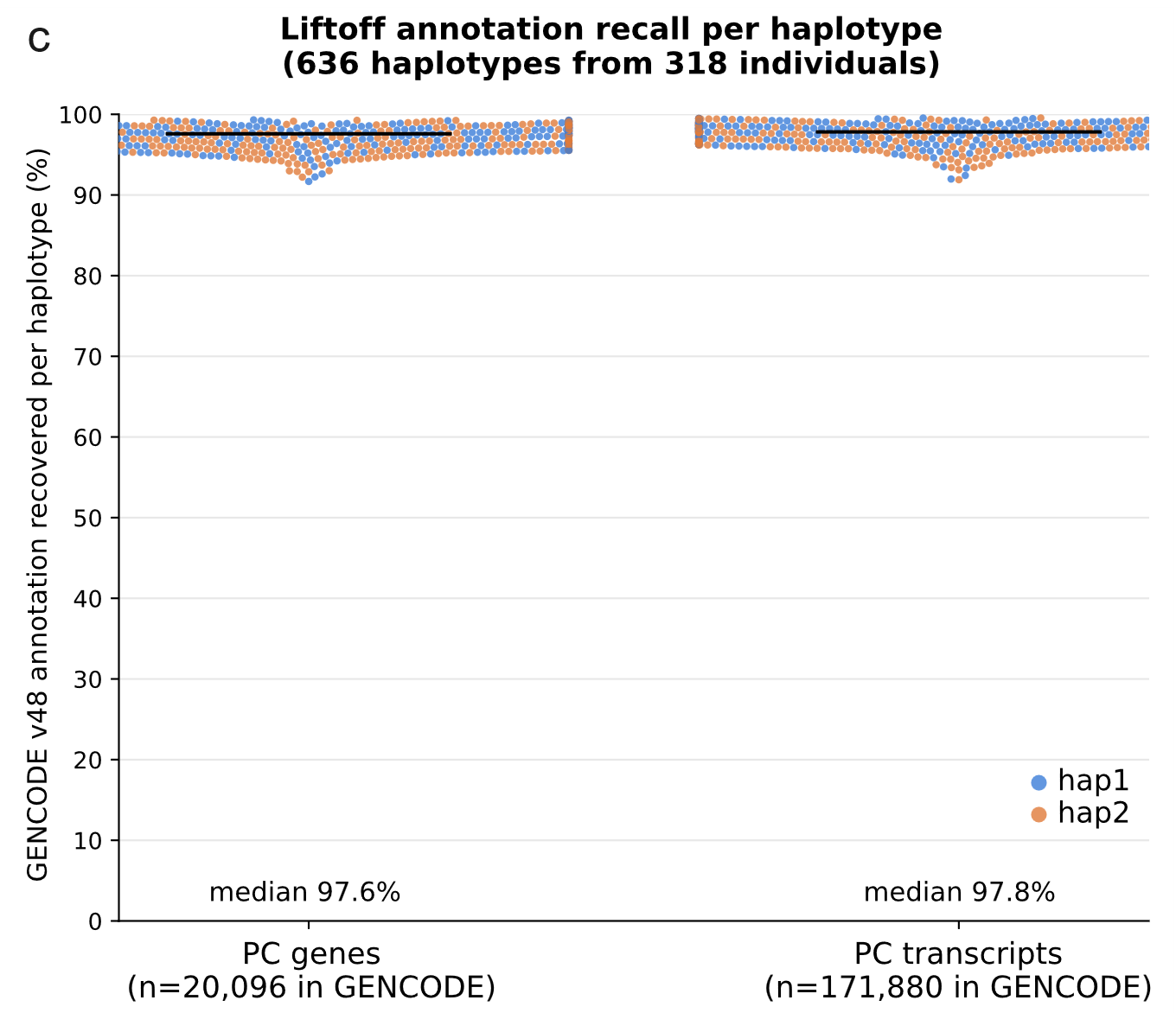

**Supplementary Figure 10. (a)** Haplotype wise total length distribution of the NDD cohort (n=318) assemblies. Violins show the distribution across samples, with overlaid box plots (median, IQR along with whiskers to 1.5× IQR). Hap1 (blue) and Hap2 (brown) average assembly length was approximately 3.0 Gb per haplotype. **(b)** Haplotype wise assembly QV distribution for the NDD cohort (n=318). Violins show the distribution across samples, with overlaid box plots (same as panel a). Hap1 (blue) and Hap2 (brown) assemblies were of comparable quality, with median QV of ~61 for both haplotypes conferring high-quality assemblies. **(c)** Percentage of GENCODE protein-coding annotation in recovered on haplotype assemblies (n=636 haplotypes) using Liftoff [*Left*, protein-coding genes (n=20,096 in the GENCODE reference); *right*, protein-coding transcripts (n=171,880)]. Each point is one haplotype assembly, colored by haplotype of origin (hap1, blue; hap2, orange) and jittered horizontally to reduce overplotting; horizontal black lines mark the median. Median recall was 97.6% for genes and 97.8% for transcripts.

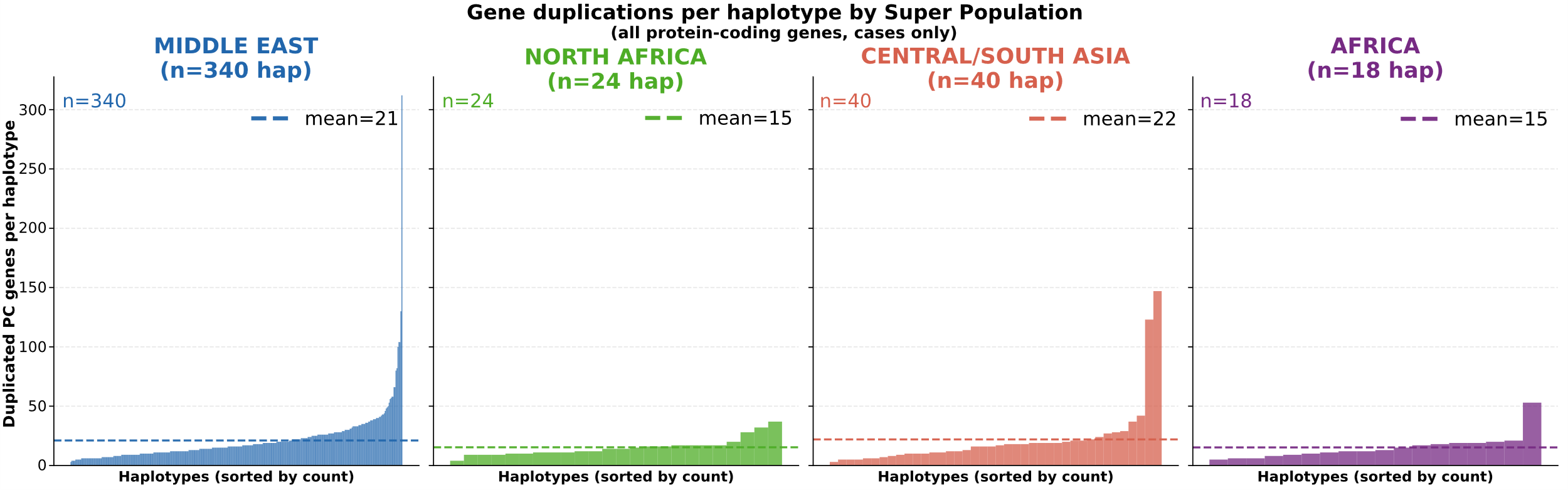

**Supplementary Figure 11.** Reference-corrected count of duplicated protein-coding autosomal genes in each haplotype assembly of the NDD case cohort (422 haplotypes from 211 individuals), shown separately for each of the four superpopulations represented: Middle East (340 haplotypes, blue), North Africa (24, green), Central/South Asia (40, peach) and Africa (sss18, purple). Each bar is one haplotype, ordered within its panel by increasing count; the dashed horizontal line marks the group mean (21, 15, 22 and 15 genes per haplotype respectively). The y axis is shared across all four panels. Most haplotypes carry fewer than 25 duplicated genes, with a small tail of haplotypes carrying considerably more, reaching 312 in the Middle East group and 147 in Central/South Asia.

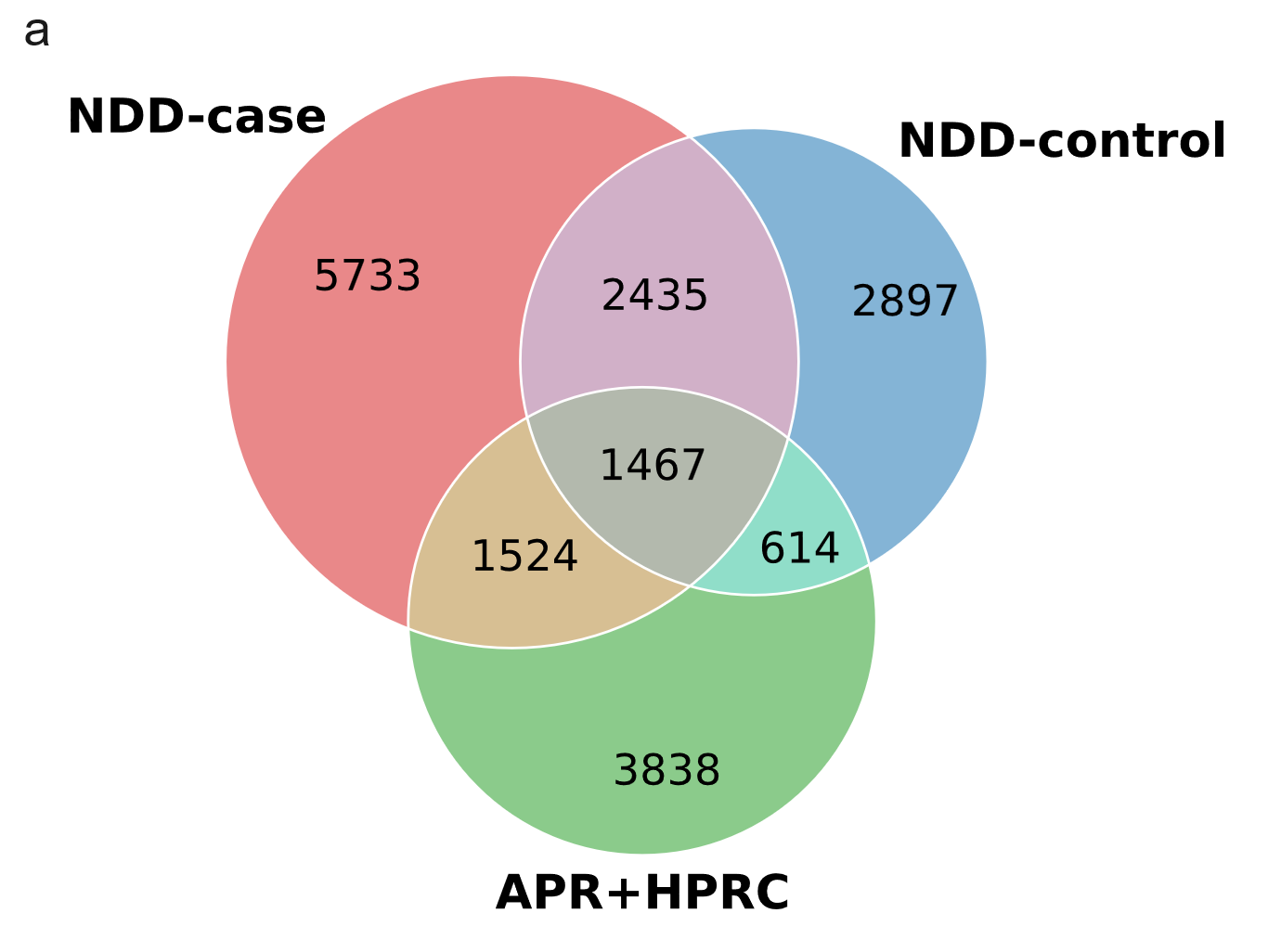

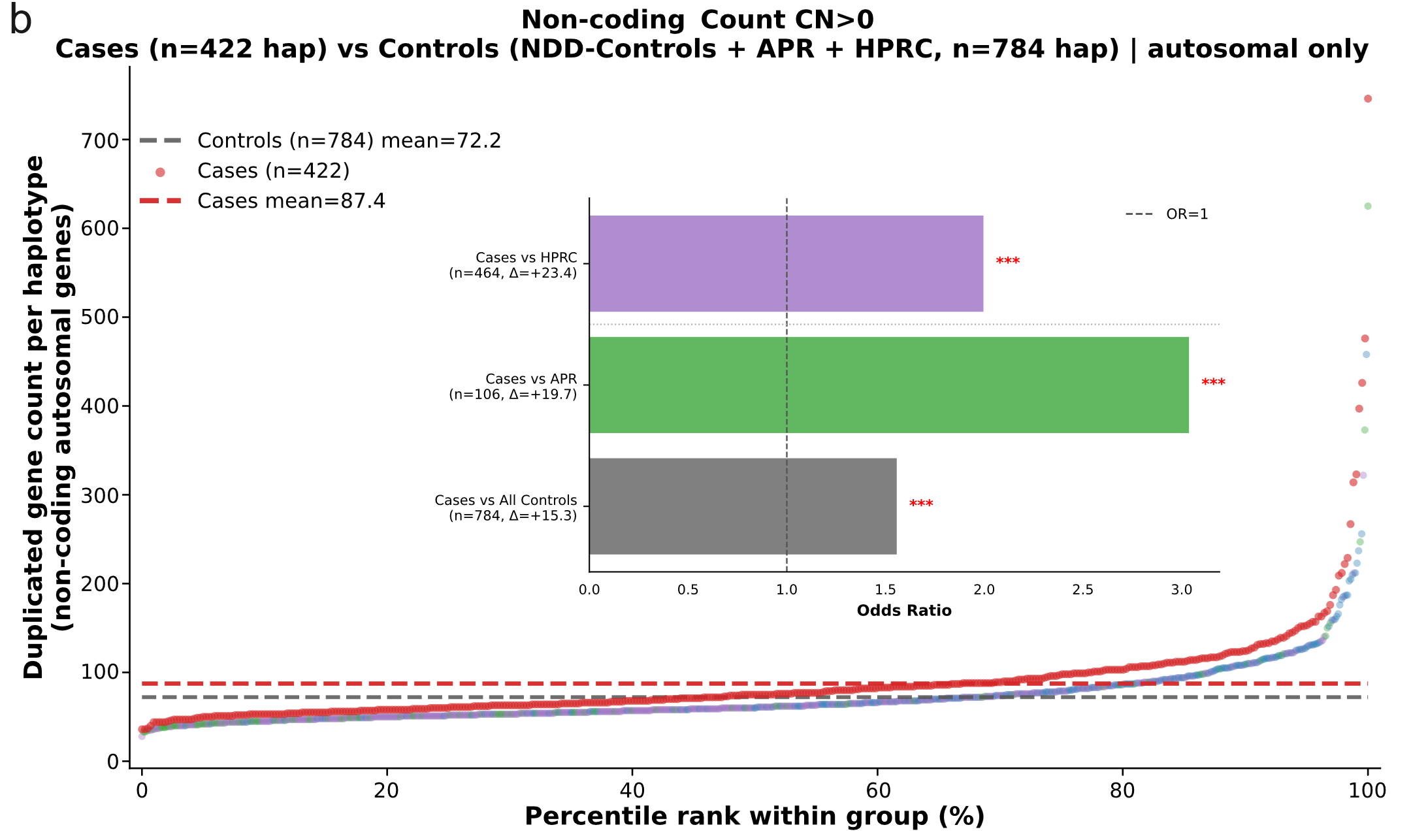

**Supplementary Figure 12.** Duplicated non-coding gene burden in NDD cases relative to control haplotypes. **(a)** Venn diagram of duplicated non-coding genes detected in NDD case, family control and combined HPRC_r2 + APR haplotypes (red, blue and green respectively, blended colors indicate intersections). Numbers give the genes unique to each group or shared between groups (5,733 duplicated non-coding genes were detected only in case haplotypes). **(b)** Ranked distribution of reference-corrected duplicated non-coding autosomal gene counts per haplotype, with haplotypes ordered by percentile rank within their own group. NDD cases are shown in red, family controls in blue, APR in green and HPRC_r2 in purple. The red and grey dashed horizontal lines mark the mean duplication burden in cases (87.4 genes per haplotype, n=422) and in all controls combined (72.2, n=784). The distributions overlap through the lower percentiles and separate in the upper tail. Inset shows odds ratios for case-control comparisons (all significant, *** indicates p<0.0001), the dashed line marks OR=1.

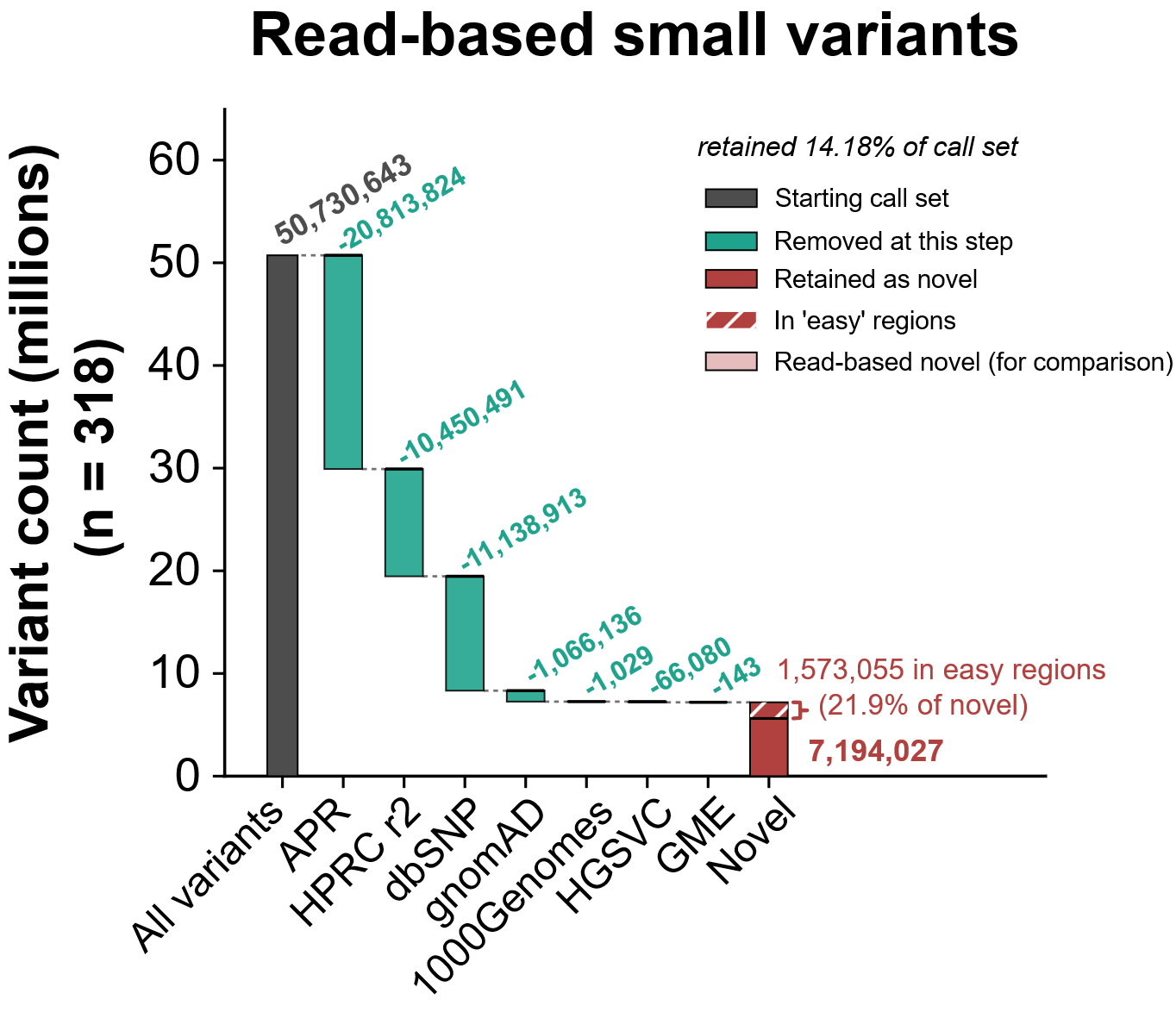

**Supplementary Figure 13.** Waterfall plot of the sequential filtering cascade used to define population-novel small variants (SNVs and indels) called from reads across the cohort (n=318). The dark grey bar gives the starting call set (50,730,643 variants). Teal bars show the number of variants removed at each step as the call set is matched in turn against the Arab Pangenome Reference (APR), HPRC_r2, dbSNP, gnomAD, the 1000 Genomes Project, HGSVC and the Greater Middle East (GME) variome; the number removed is given above each step. The red bar gives the 7,194,027 variants remaining after all seven databases, retained as population-novel (14.18% of the starting call set). Hatching marks the 1,573,055 novel variants (21.9%) falling within high-confidence (“easy”) regions. Note that because filtering is sequential, the number attributed to each database depends on its position in the cascade: a variant present in several databases is counted only at the first one that removes it, so the per-step values are not database-specific counts.

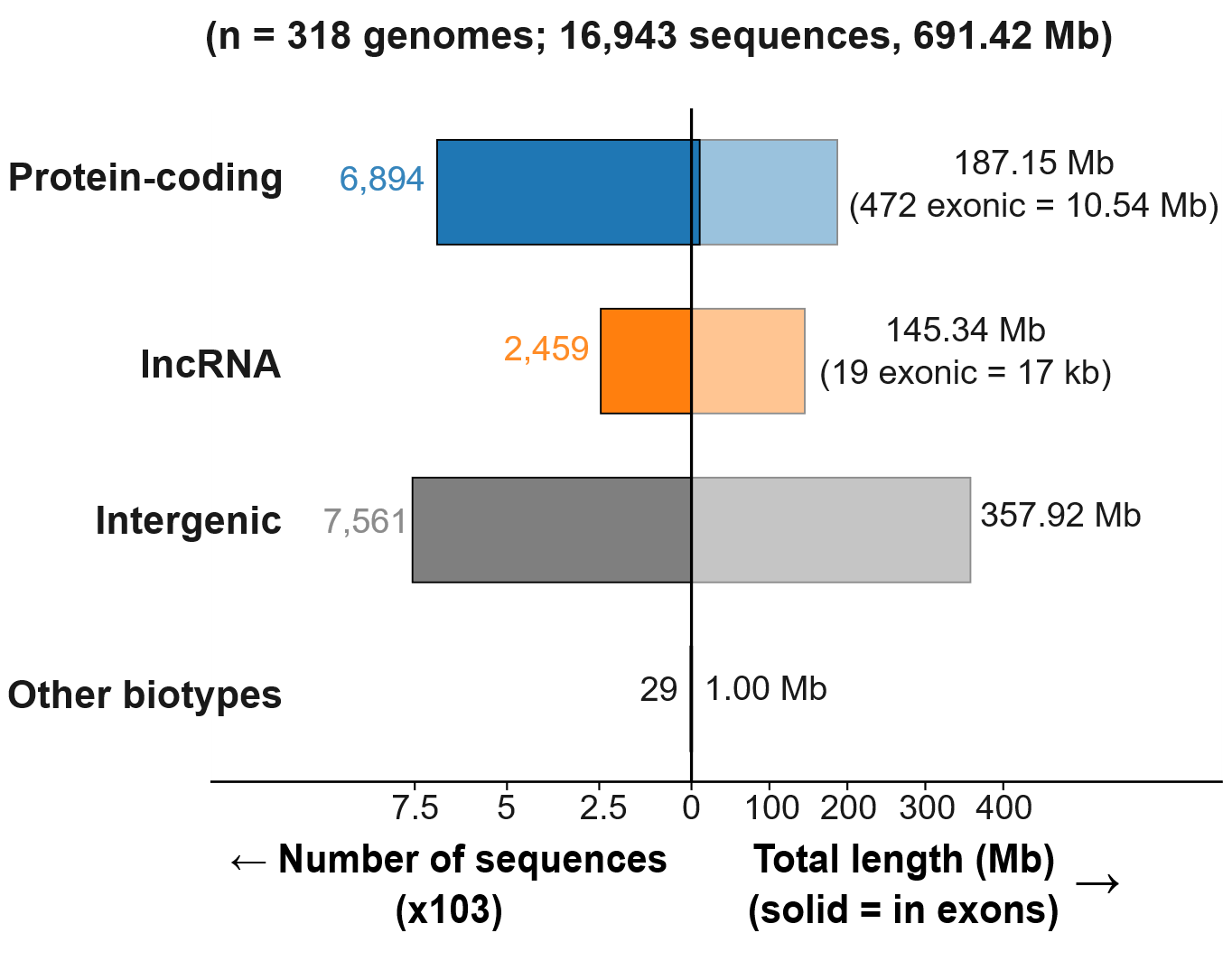

**Supplementary Figure 14a.** Gene biotype distribution of novel sequences in NDD cohort (n = 318). The dividing bar represents 16,943 novel sequences totaling 691.42 Mb identified in the whole cohort. The two sides of the axis carry different units and independent scales, so the bar widths are not comparable across the midline. The left side of the y-axis shows the number of sequences per biotype (×10³), and the right side shows the total length (in Mb), with the portion falling in annotated exons (drawn solid and annotated in brackets).

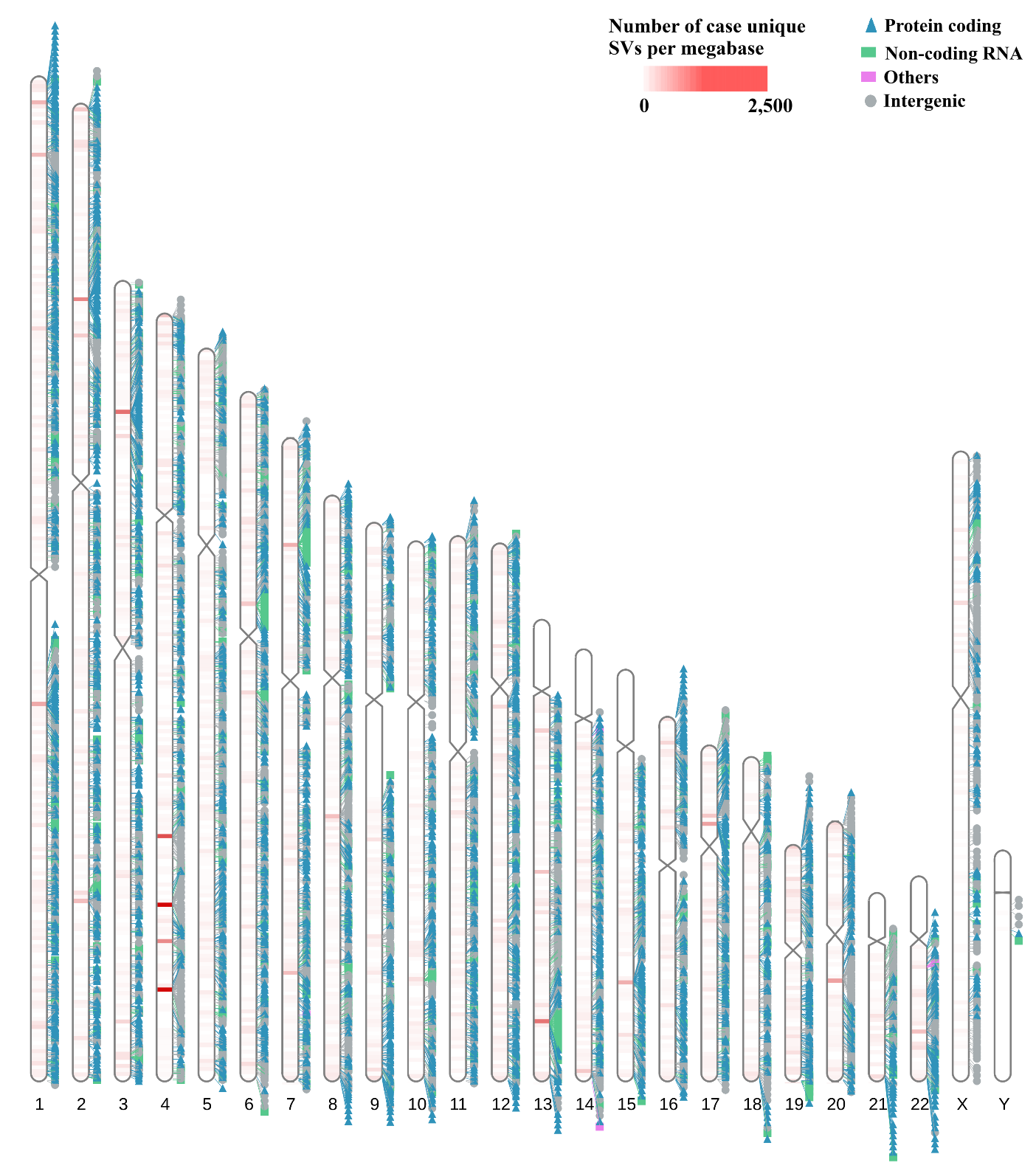

**Supplementary Figure 14b. Genome-wide distribution of case-unique novel sequences in high-confidence (‘easy’) regions.** Ideogram of the GRCh38 reference showing the chromosomal distribution of case-unique novel sequences (SV insertions) identified from the NDD pangenome graph and restricted to high-confidence (“easy”) regions. Each marker denotes one novel sequence, positioned at its insertion site and colored by the annotation class of the overlapping feature: protein-coding genes (blue triangles), non-coding RNA genes (medium sea green boxes), other gene biotypes (orchid boxes) and intergenic positions (grey circles). The heatmap overlaid on each chromosome body gives the density of case-unique novel sequences per megabase, from 0 (white) to 2,500 (red). Case-unique sequences are those absent from all 782 control haplotypes (HPRC r2, APR and family controls) by the sequential filtering cascade.

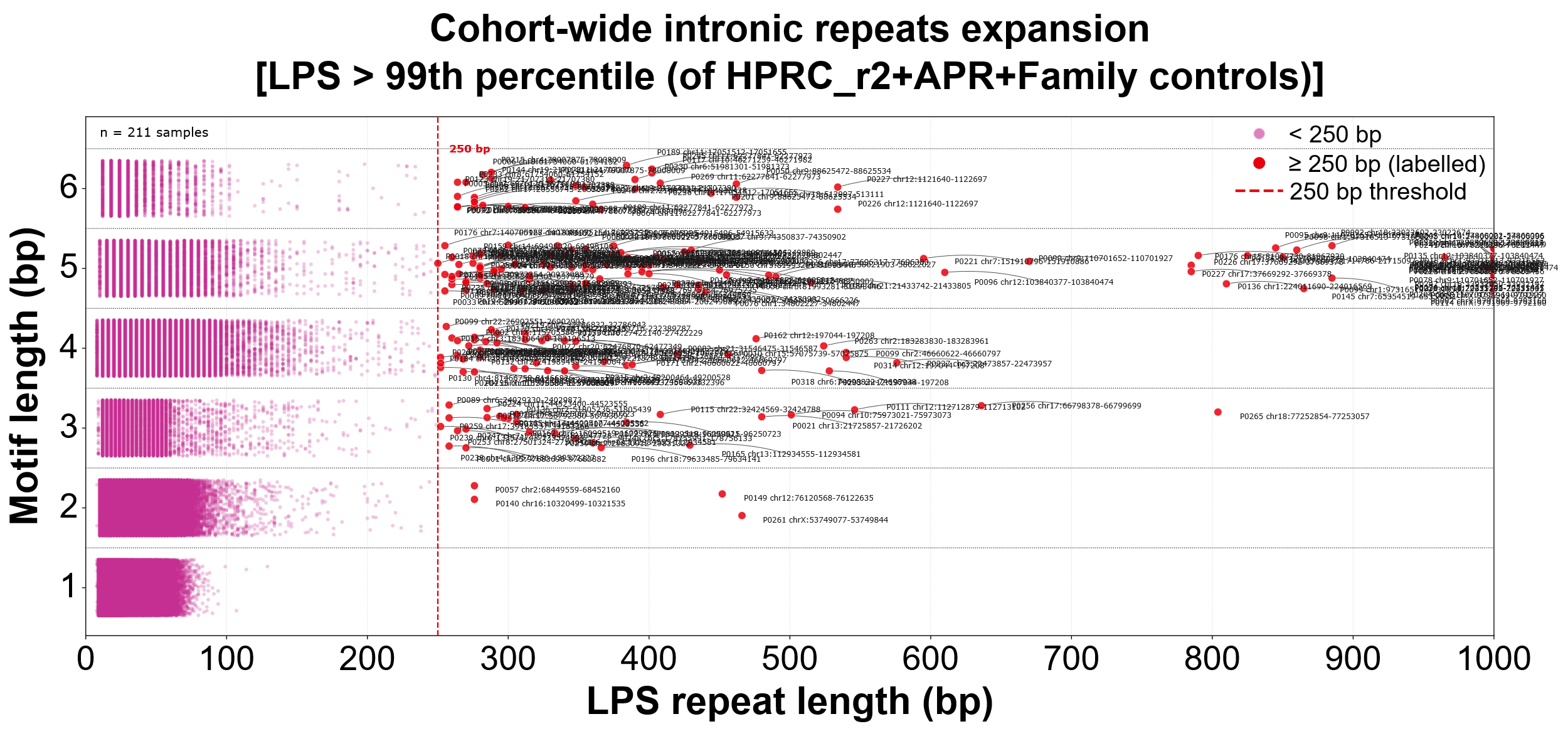

**Supplementary Figure 15.** Longest pure segment (LPS) (bp, x-axis) versus motif length (y-axis) for the global intronic tandem repeat alleles across all 211 NDD cases. Each point represents a single repeat locus; the pink points indicate alleles with an LPS <250 bp, and the red points indicate those with an LPS ≥250 bp, which exceed the expansion threshold (red dashed line). Sample ID and chromosomal coordinates annotated for LPS expansions ≥250 bp. Motif classes of 1–6 bp were assessed, with dense clusters at shorter motif lengths consistent with genome-wide background repeat variation. A total of 230 intronic expansions were observed.

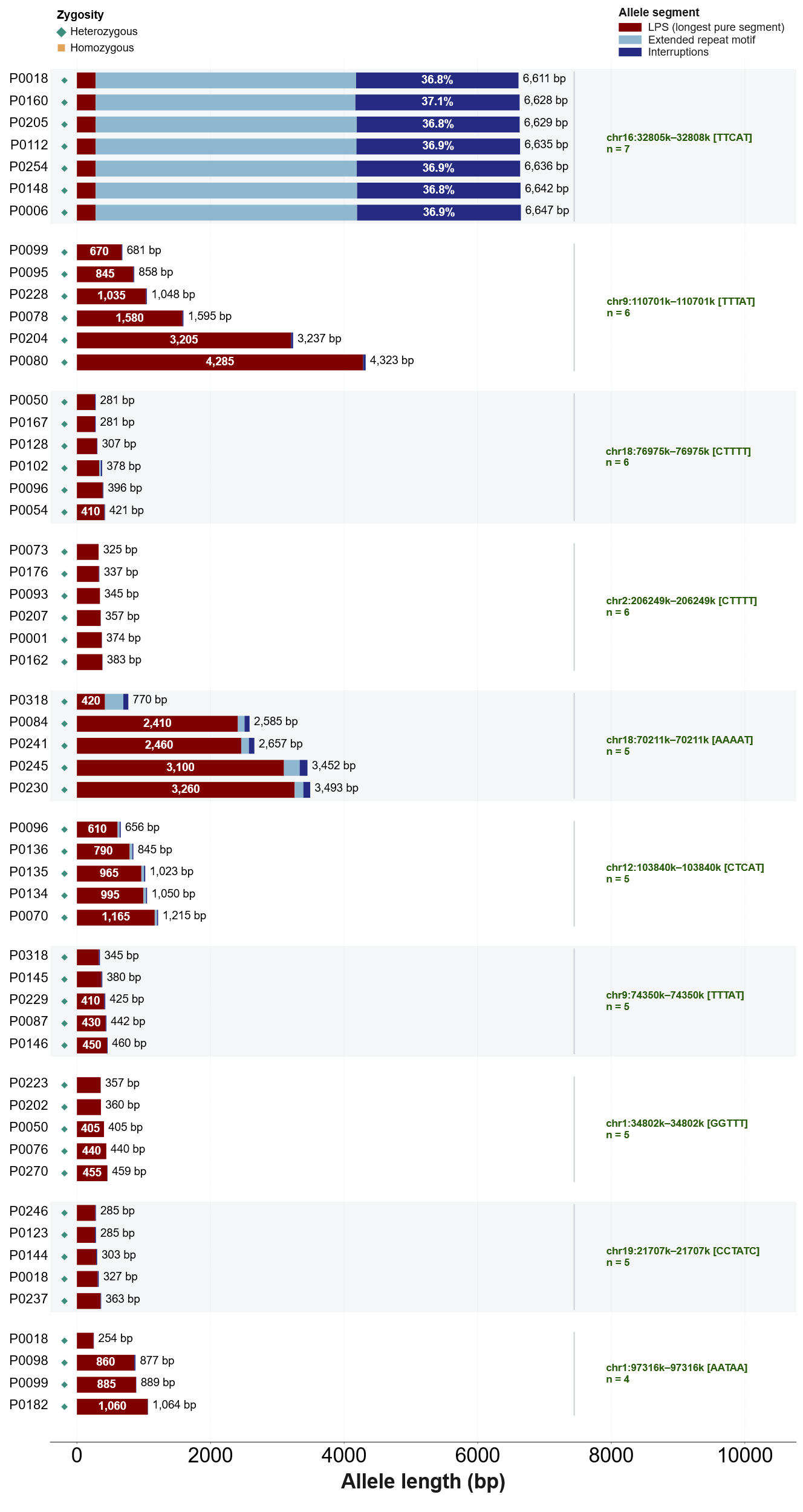

**Supplementary Figure 16.** Allele architecture of the recurrent intronic expansions identified in Supplementary Figure 15. Each bar is a single allele, decomposed into the LPS (dark blue), the surrounding extended motif region (light blue) and interruptions within the tract (red, with the interrupted fraction shown where segment width permits). All of these are heterozygous (shown by a green diamond marker). Total allele length is annotated to the right of each bar and LPS length within the LPS segment where width permits; alleles are grouped by locus, with groups ordered by number of carriers (n) and bars within each group by total allele length. The [TTCAT] expansions at chr16:32,805k–32,808k are both the longest alleles observed (6,611–6,647 bp) and the most heavily interrupted (36.8–37.1% of allele length), yet their LPS is invariant at 280 bp across all seven carriers. The LPS among the recurrent loci was found to be of 4,285 bp expansion in one of the case.
