## Supplementary File 2 for "Pangenome Discoveries from Clinical Long-Read Genomes in Underrepresented Populations with Neurodevelopmental Disorders"

**SNP ANALYSIS** **(CHROMATOGRAM)**

| **Sample** | **Gene** | **Chromosome position** | **Ref.** | **Alt.** | **Result** | **zygosity** |
| --- | --- | --- | --- | --- | --- | --- |
| P0069 | NIPBL | chr5:37000508-37000508 | G | A | G/A | HETERO |

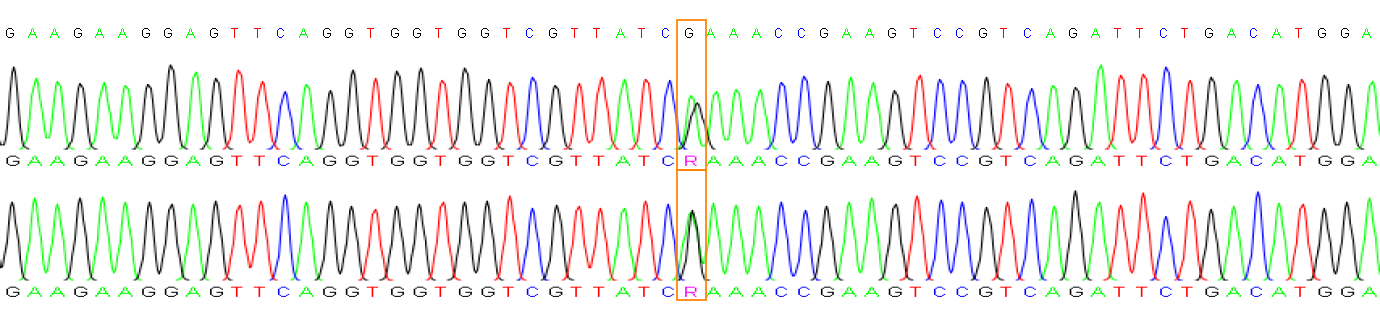

| **Sample** | **Gene** | **Chromosome position** | **Ref.** | **Alt.** | **Result** | **zygosity** |
| --- | --- | --- | --- | --- | --- | --- |
| P0073 | BICRA | chr19:47698693-47698693 | C | T | C/T | HETERO |

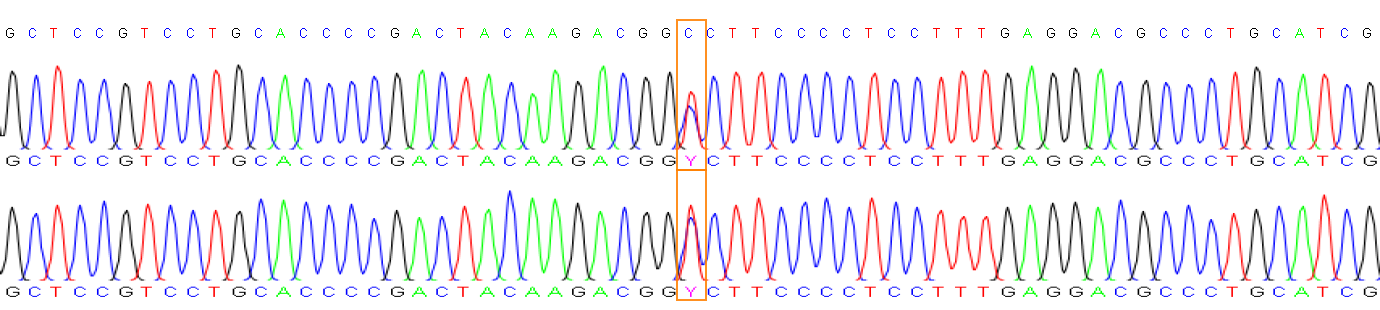

| **Sample** | **Gene** | **Chromosome position** | **Ref.** | **Alt.** | **Result** | **zygosity** |
| --- | --- | --- | --- | --- | --- | --- |
| P0076 | SYT1 | chr12:78864916-78864916 | T | G | T/G | HETERO |

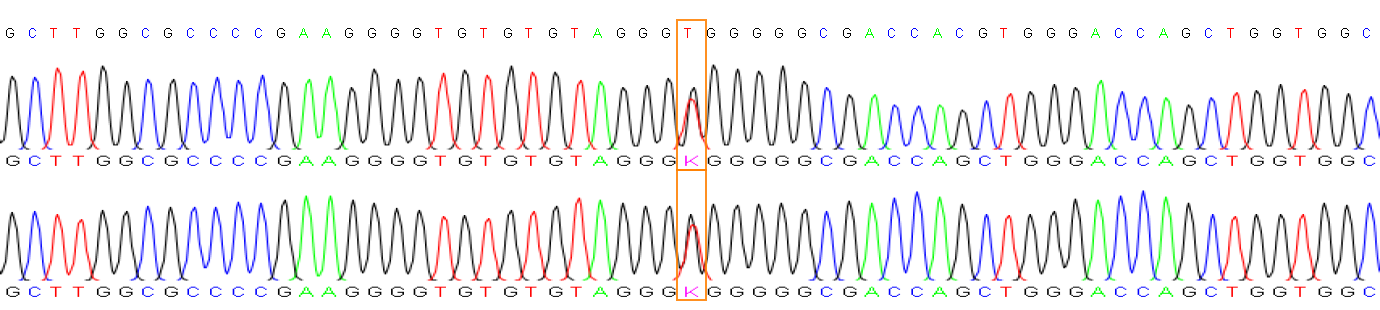

| **Sample** | **Gene** | **Chromosome position** | **Ref.** | **Alt.** | **Result** | **zygosity** |
| --- | --- | --- | --- | --- | --- | --- |
| P0076 | KDM6B | chr17:7849251-7849251 | G | A | G/A | HETERO |

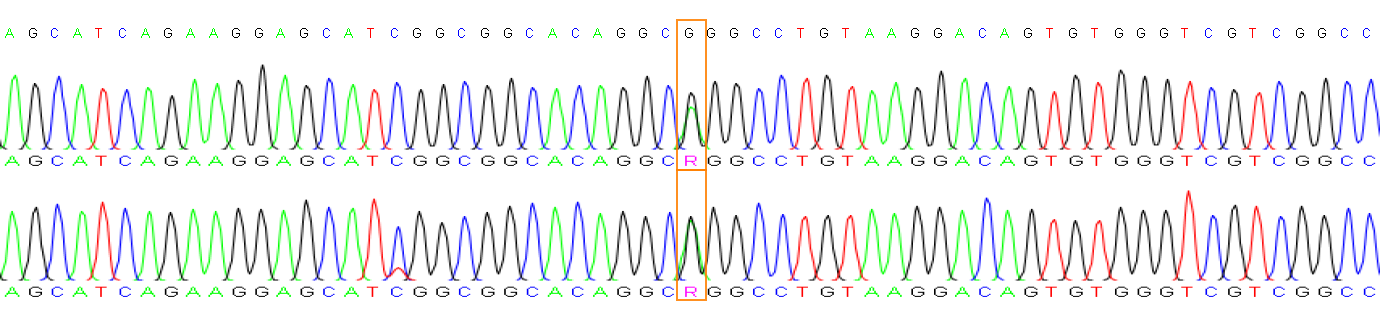

| **Sample** | **Gene** | **Chromosome position** | **Ref.** | **Alt.** | **Result** | **zygosity** |
| --- | --- | --- | --- | --- | --- | --- |
| P0078 | GRIN1-1 | chr9:137157028-137157028 | T | A | T/A | HETERO |

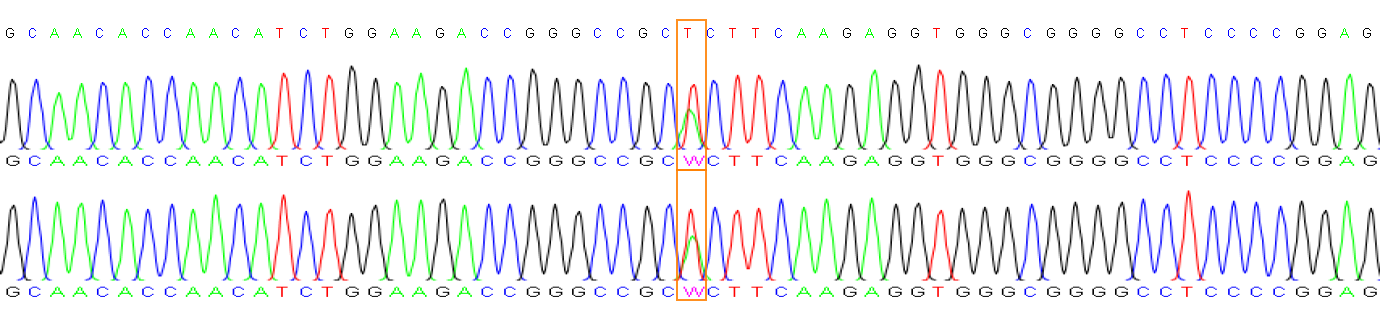

| **Sample** | **Gene** | **Chromosome position** | **Ref.** | **Alt.** | **Result** | **zygosity** |
| --- | --- | --- | --- | --- | --- | --- |
| P0078 | GABRB3 | chr15:26939289-26939289 | G | A | G/A | HETERO |

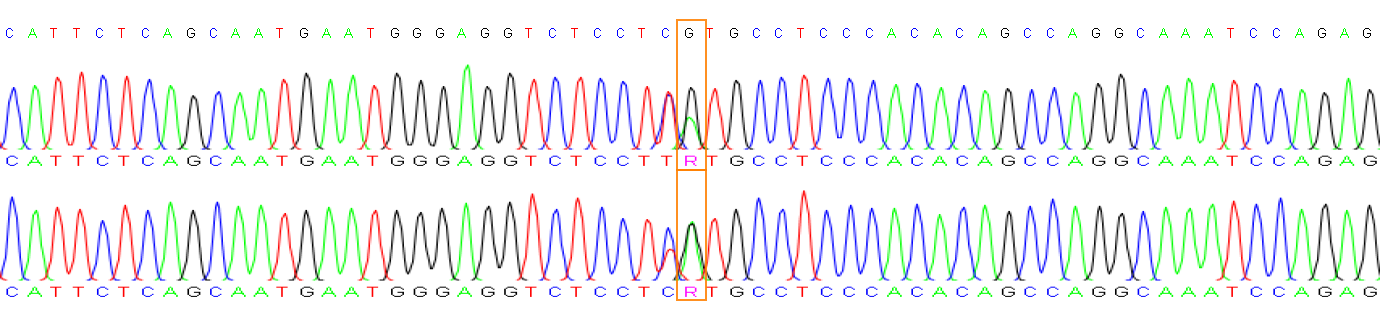

| **Sample** | **Gene** | **Chromosome position** | **Ref.** | **Alt.** | **Result** | **zygosity** |
| --- | --- | --- | --- | --- | --- | --- |
| P0079 | CHD3-1 | chr17:7899056-7899056 | G | A | G/A | HETERO |

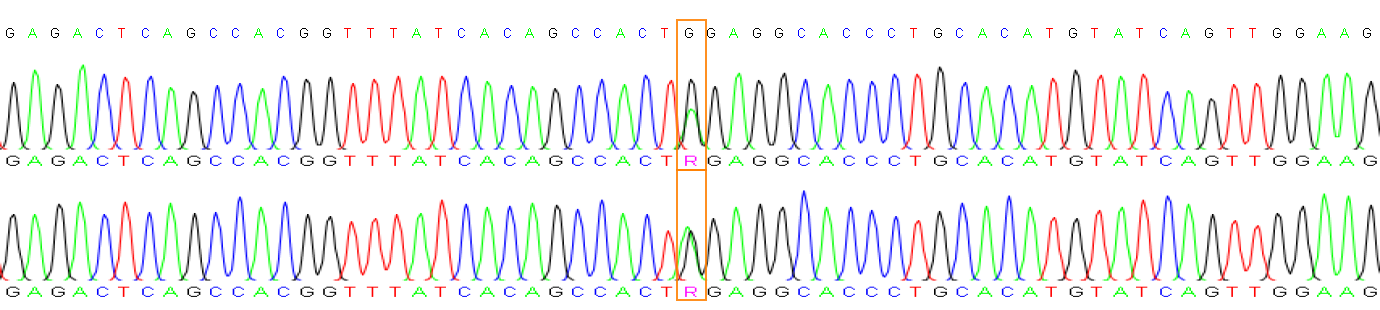

| **Sample** | **Gene** | **Chromosome position** | **Ref.** | **Alt.** | **Result** | **zygosity** |
| --- | --- | --- | --- | --- | --- | --- |
| P0079 | MED13 | chr17:62052622-62052622 | G | A | G/A | HETERO |

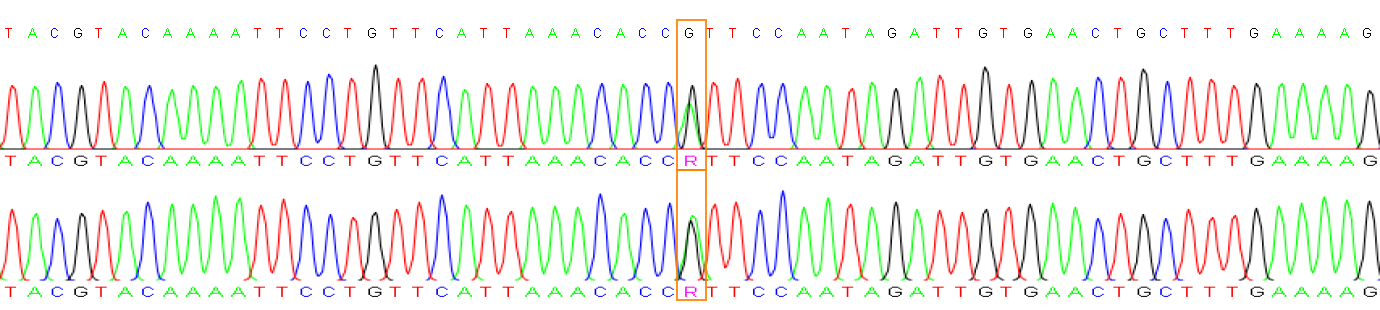

| **Sample** | **Gene** | **Chromosome position** | **Ref.** | **Alt.** | **Result** | **zygosity** |
| --- | --- | --- | --- | --- | --- | --- |
| P0081 | FRMD5 | chr15:43874249-43874249 | T | C | T/C | HETERO |

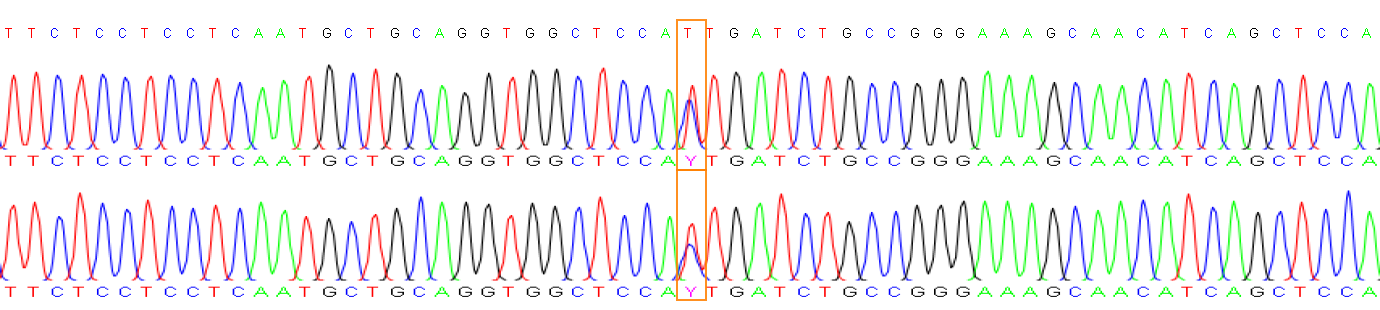

| **Sample** | **Gene** | **Chromosome position** | **Ref.** | **Alt.** | **Result** | **zygosity** |
| --- | --- | --- | --- | --- | --- | --- |
| P0084 | SAMD9 | chr7:93103628-93103628 | G | A | G/A | HETERO |

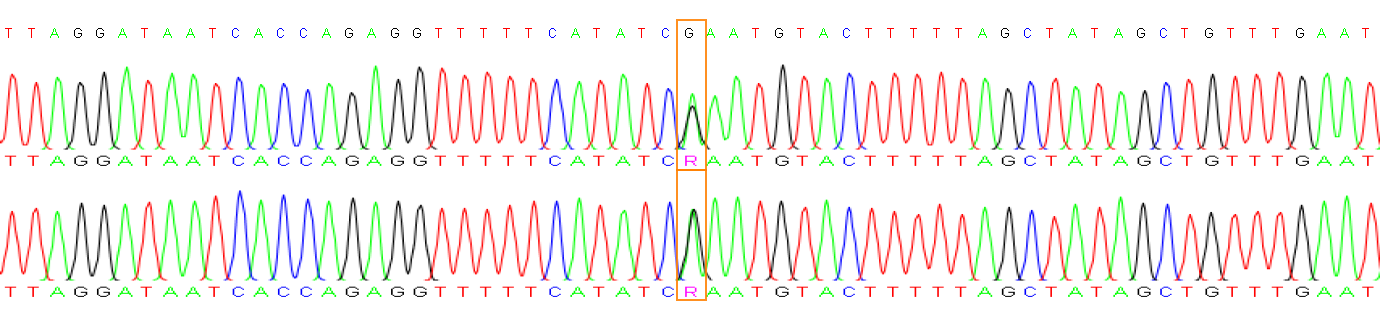

| **Sample** | **Gene** | **Chromosome position** | **Ref.** | **Alt.** | **Result** | **zygosity** |
| --- | --- | --- | --- | --- | --- | --- |
| P0084 | PUM1 | chr1:30936681-30936681 | C | T | C/T | HETERO |

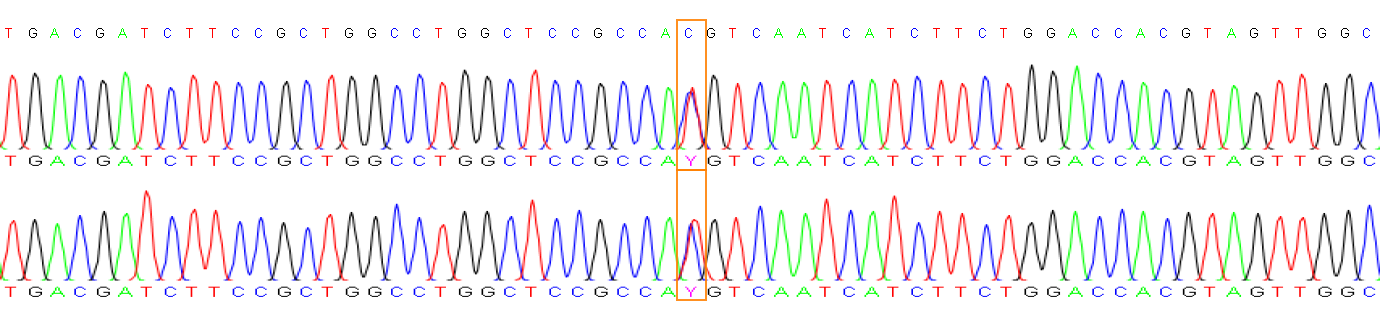

| **Sample** | **Gene** | **Chromosome position** | **Ref.** | **Alt.** | **Result** | **zygosity** |
| --- | --- | --- | --- | --- | --- | --- |
| P0084 | KMT2E-1 | chr7:105107220-105107220 | G | C | G/C | HETERO |

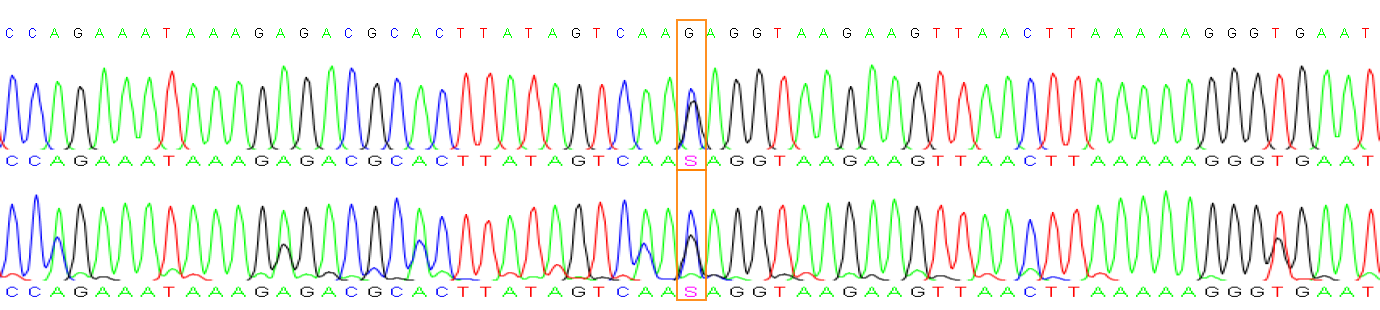

| **Sample** | **Gene** | **Chromosome position** | **Ref.** | **Alt.** | **Result** | **zygosity** |
| --- | --- | --- | --- | --- | --- | --- |
| P0085 | NLGN1 | chr3:174280522-174280522 | A | G | A/G | HETERO |

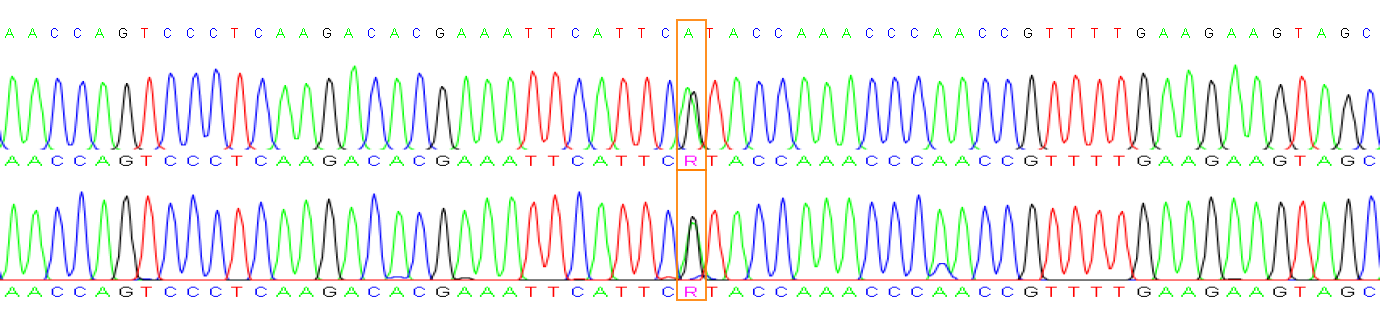

| **Sample** | **Gene** | **Chromosome position** | **Ref.** | **Alt.** | **Result** | **zygosity** |
| --- | --- | --- | --- | --- | --- | --- |
| P0085 | TRIO-1 | chr5:14280367-14280367 | G | A | G/A | HETERO |

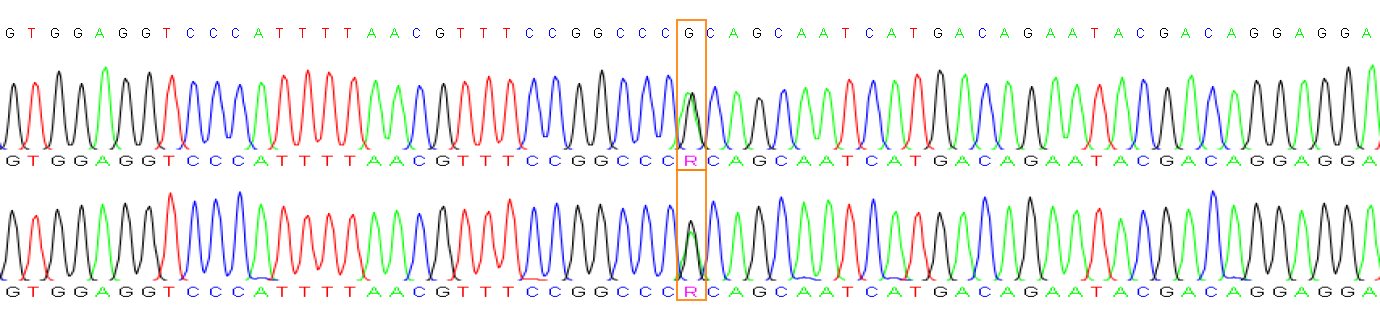

| **Sample** | **Gene** | **Chromosome position** | **Ref.** | **Alt.** | **Result** | **zygosity** |
| --- | --- | --- | --- | --- | --- | --- |
| NDD21 | CIC-1 | chr19:42292730-42292731 | GC | G | GC | HOMO |

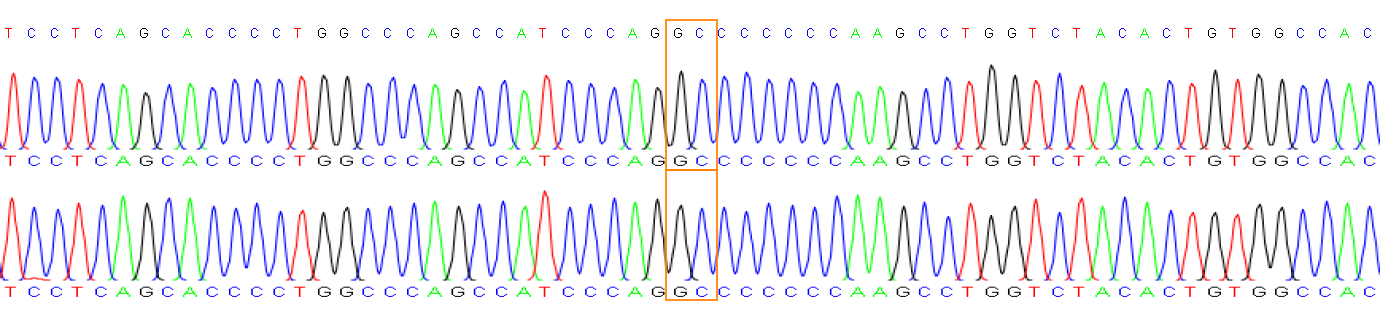

| **Sample** | **Gene** | **Chromosome position** | **Ref.** | **Alt.** | **Result** | **zygosity** |
| --- | --- | --- | --- | --- | --- | --- |
| P0087 | NLGN1 | chr3:174280522-174280522 | A | G | A/G | HETERO |

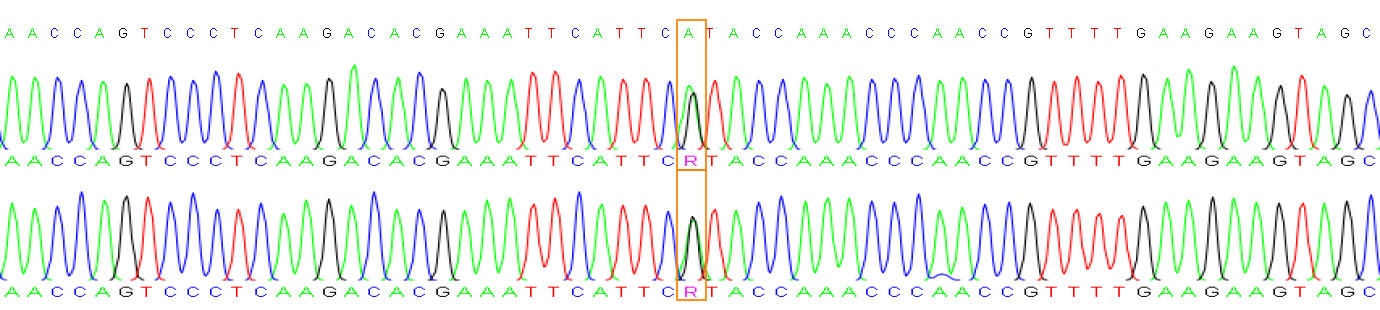

| **Sample** | **Gene** | **Chromosome position** | **Ref.** | **Alt.** | **Result** | **zygosity** |
| --- | --- | --- | --- | --- | --- | --- |
| P0087 | CIC-2 | chr19:42272064-42272064 | A | G | A/G | HETERO |

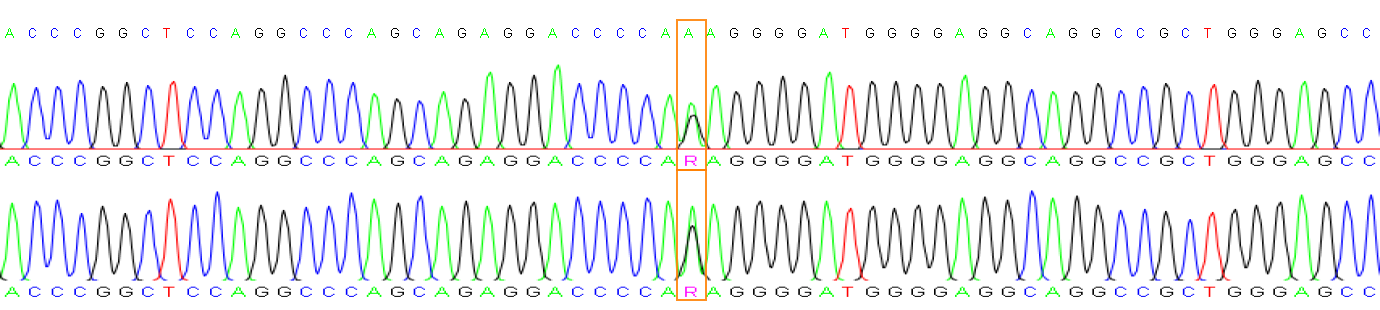

| **Sample** | **Gene** | **Chromosome position** | **Ref.** | **Alt.** | **Result** | **zygosity** |
| --- | --- | --- | --- | --- | --- | --- |
| P0090 | CIC-3 | chr19:42287328-42287328 | C | A | C/A | HETERO |

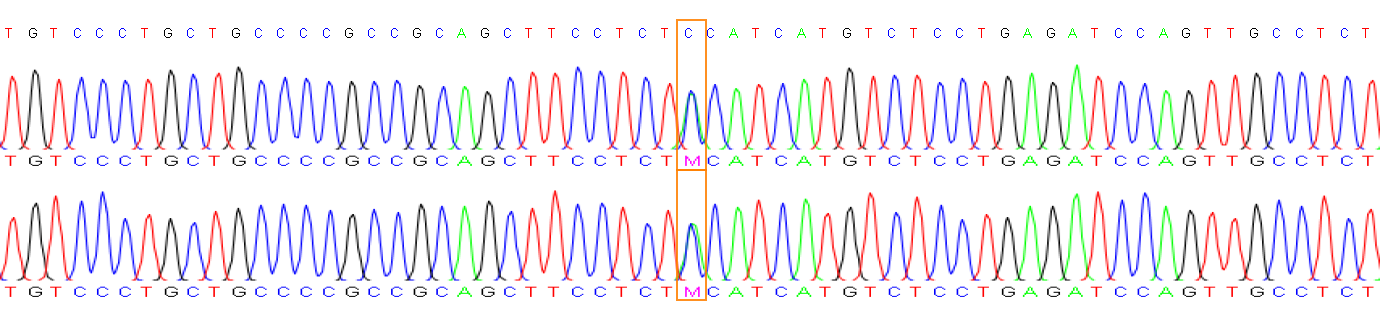

| **Sample** | **Gene** | **Chromosome position** | **Ref.** | **Alt.** | **Result** | **zygosity** |
| --- | --- | --- | --- | --- | --- | --- |
| P0090 | FBXO11 | chr2:47808283-47808284 | AG | A | AG | HOMO |

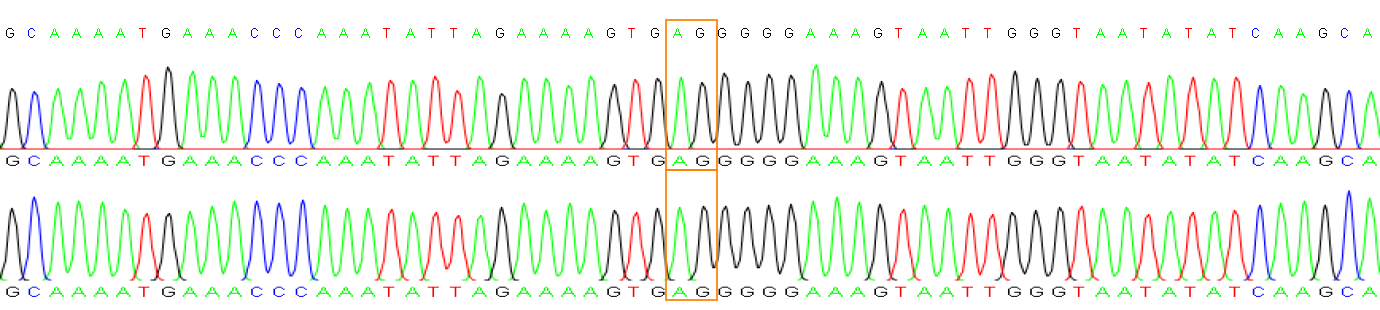

| **Sample** | **Gene** | **Chromosome position** | **Ref.** | **Alt.** | **Result** | **zygosity** |
| --- | --- | --- | --- | --- | --- | --- |
| P0091 | AHDC1-1 | chr1:27550357-27550358 | GT | G | GT | HOMO |

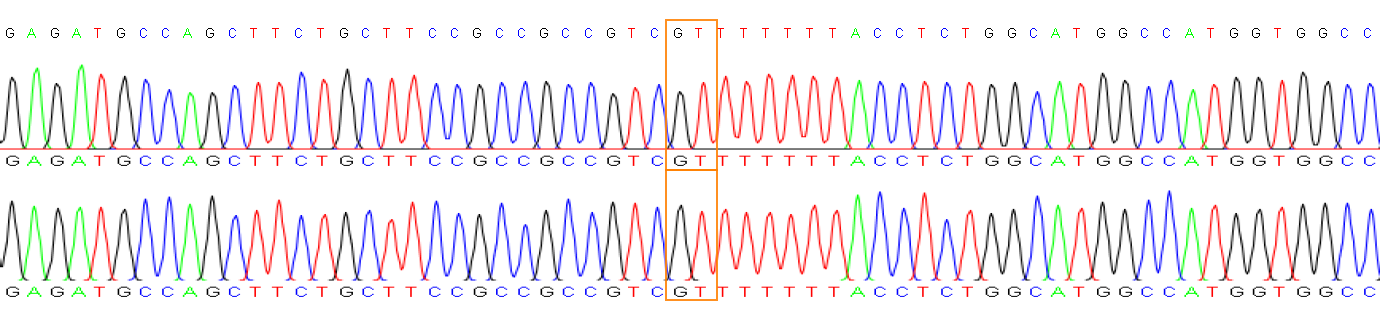

| **Sample** | **Gene** | **Chromosome position** | **Ref.** | **Alt.** | **Result** | **zygosity** |
| --- | --- | --- | --- | --- | --- | --- |
| P0091 | ANKRD17 | chr4:73124922-73124923 | TC | T | TC | HOMO |

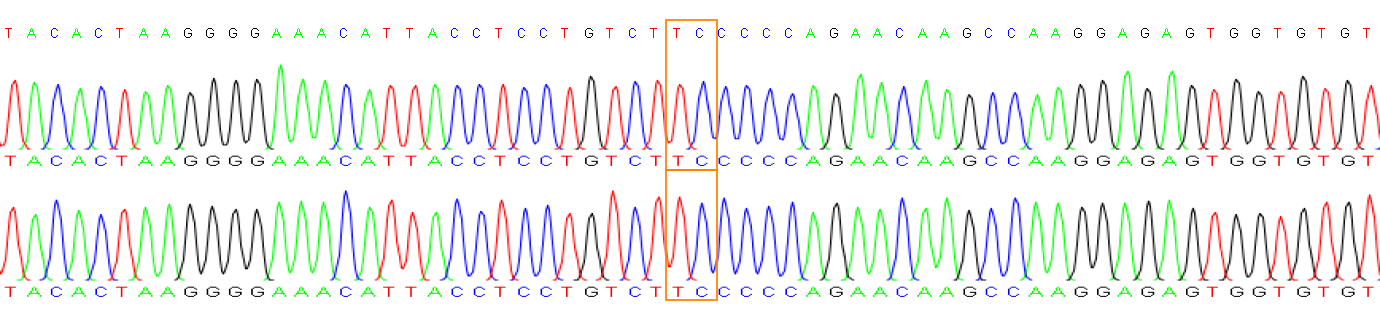

| **Sample** | **Gene** | **Chromosome position** | **Ref.** | **Alt.** | **Result** | **zygosity** |
| --- | --- | --- | --- | --- | --- | --- |
| P0091 | HUWE1 | chrX:53546516-53546517 | TC | T | TC | HOMO |

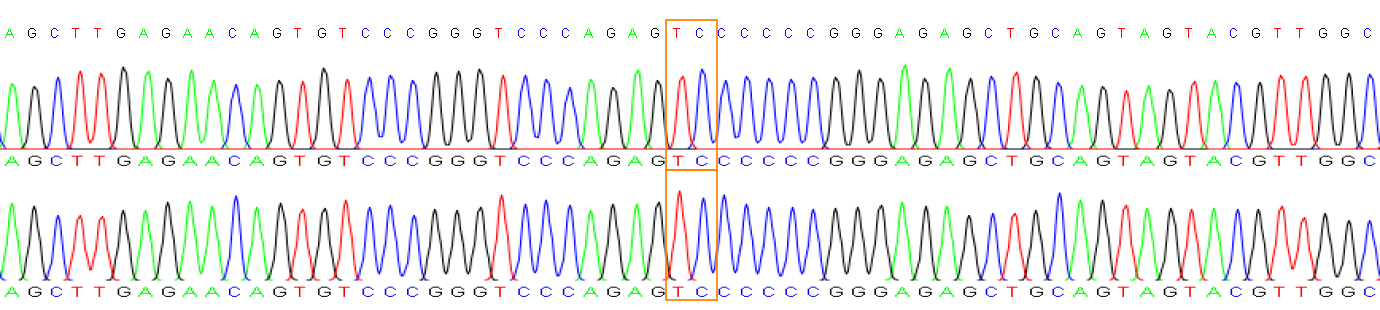

| **Sample** | **Gene** | **Chromosome position** | **Ref.** | **Alt.** | **Result** | **zygosity** |
| --- | --- | --- | --- | --- | --- | --- |
| P0092 | SLITRK2 | chrX:145823965-145823965 | G | T | G/T | HETERO |

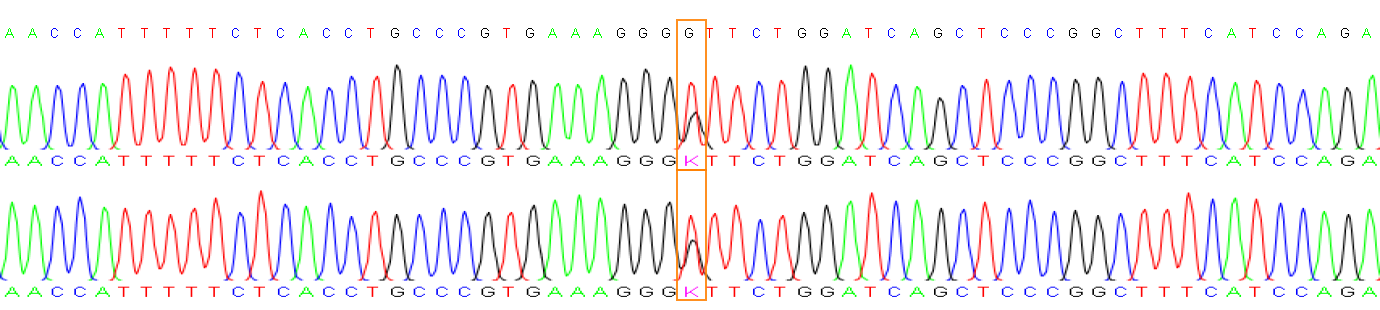

| **Sample** | **Gene** | **Chromosome position** | **Ref.** | **Alt.** | **Result** | **zygosity** |
| --- | --- | --- | --- | --- | --- | --- |
| P0092 | KCNT1 | chr9:135765151-135765151 | C | T | C | HOMO |

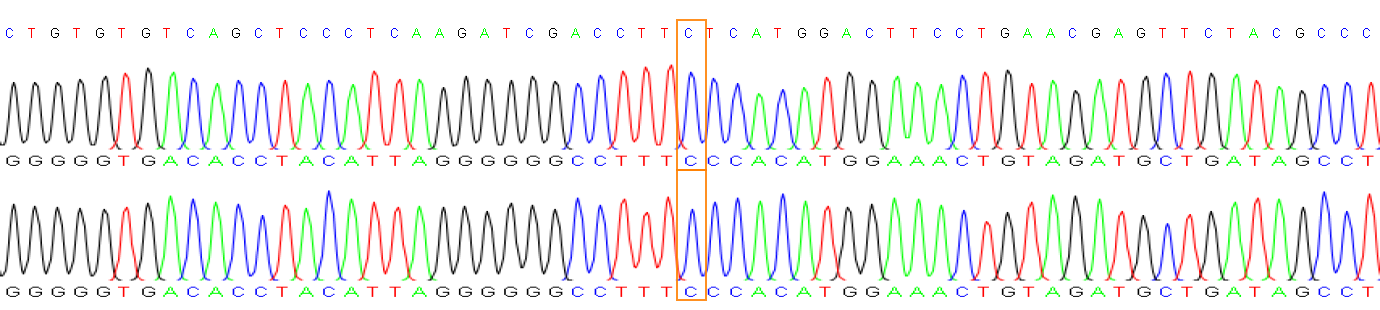

| **Sample** | **Gene** | **Chromosome position** | **Ref.** | **Alt.** | **Result** | **zygosity** |
| --- | --- | --- | --- | --- | --- | --- |
| P0094 | CHD3-2 | chr17:7908447-7908447 | C | T | C/T | HETERO |

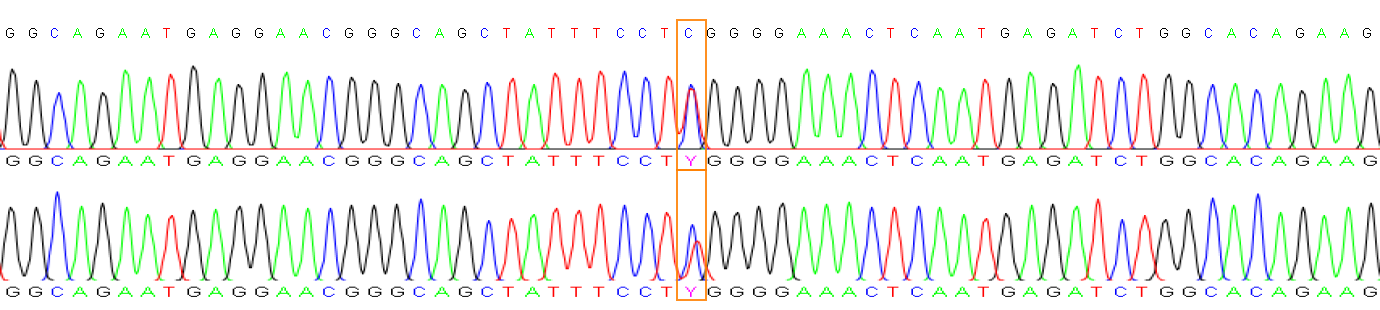

| **Sample** | **Gene** | **Chromosome position** | **Ref.** | **Alt.** | **Result** | **zygosity** |
| --- | --- | --- | --- | --- | --- | --- |
| P0097 | ADGRL1-1 | chr19:14161441-14161441 | G | A | G/A | HETERO |

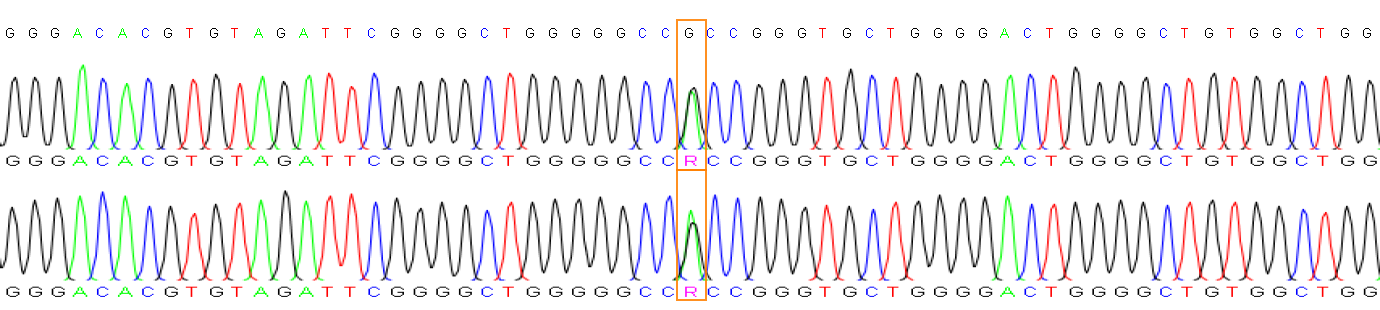

| **Sample** | **Gene** | **Chromosome position** | **Ref.** | **Alt.** | **Result** | **zygosity** |
| --- | --- | --- | --- | --- | --- | --- |
| P0102 | NTRK2 | chr9:84702232-84702232 | C | G | C/G | HETERO |

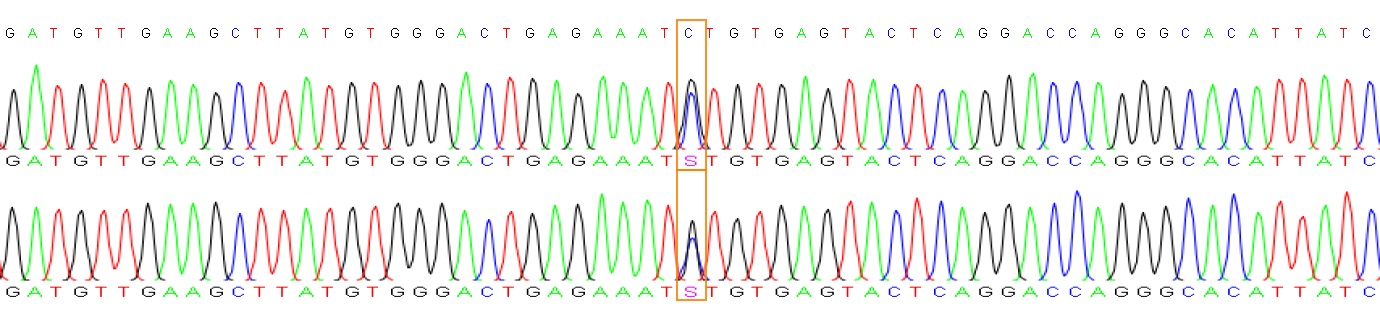

| **Sample** | **Gene** | **Chromosome position** | **Ref.** | **Alt.** | **Result** | **zygosity** |
| --- | --- | --- | --- | --- | --- | --- |
| P0102 | SRRM2 | chr16:2765966-2765966 | G | A | G/A | HETERO |

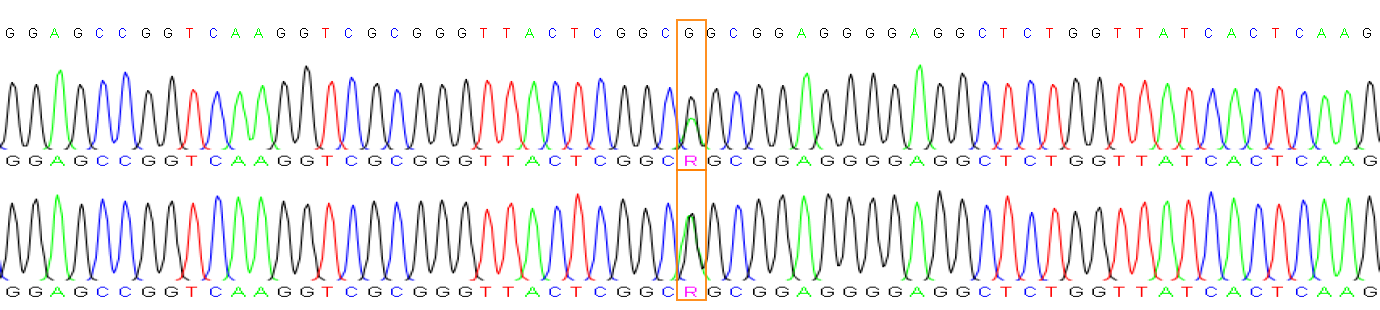

| **Sample** | **Gene** | **Chromosome position** | **Ref.** | **Alt.** | **Result** | **zygosity** |
| --- | --- | --- | --- | --- | --- | --- |
| P0109 | ADGRL1-2 | chr19:14151552-14151552 | T | C | T/C | HETERO |

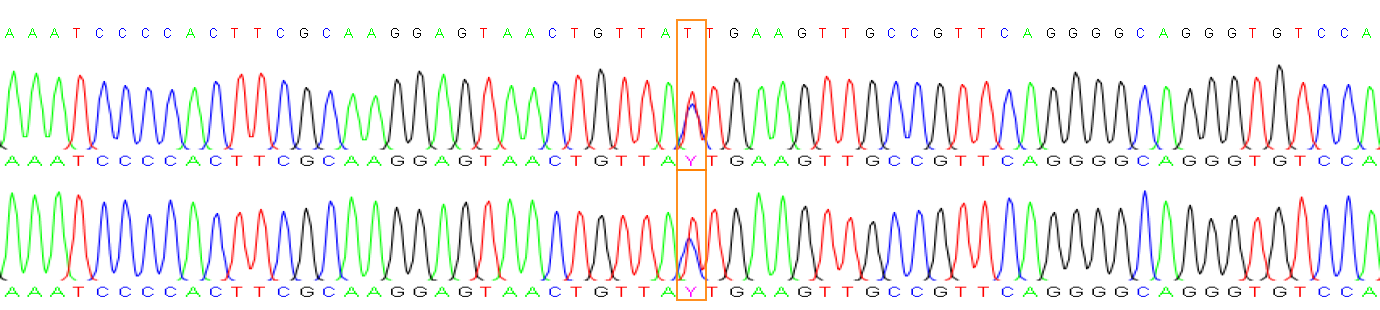

| **Sample** | **Gene** | **Chromosome position** | **Ref.** | **Alt.** | **Result** | **zygosity** |
| --- | --- | --- | --- | --- | --- | --- |
| P0114 | TRIO-2 | chr5:14498640-14498640 | G | A | G/A | HETERO |

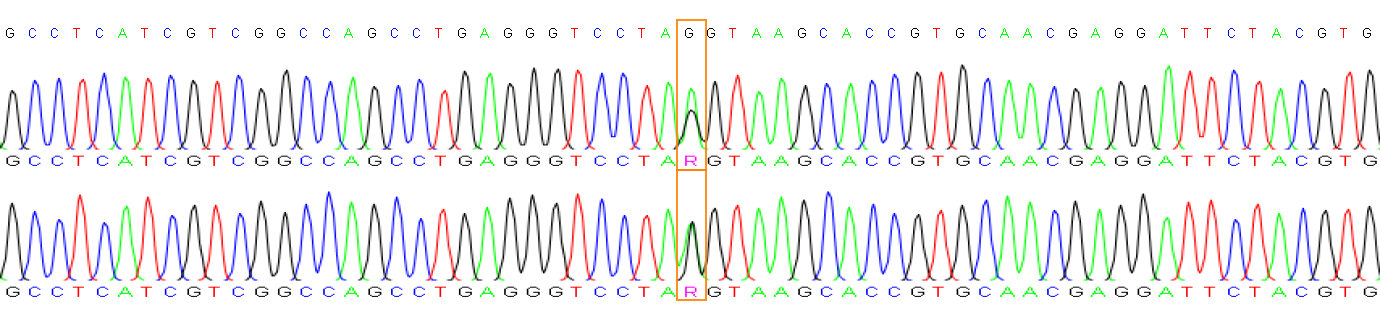

| **Sample** | **Gene** | **Chromosome position** | **Ref.** | **Alt.** | **Result** | **zygosity** |
| --- | --- | --- | --- | --- | --- | --- |
| P0114 | KMT2E-2 | chr7:105090100-105090100 | G | T | G/T | HETERO |

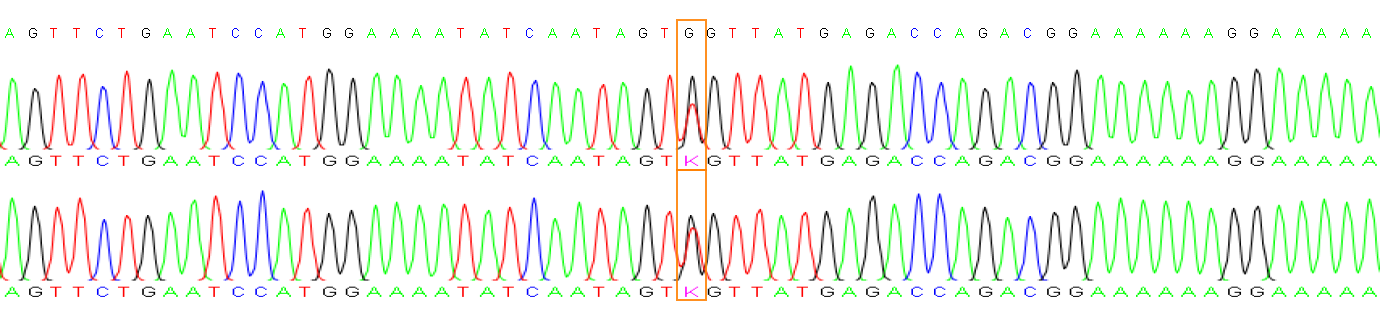

| **Sample** | **Gene** | **Chromosome position** | **Ref.** | **Alt.** | **Result** | **zygosity** |
| --- | --- | --- | --- | --- | --- | --- |
| P0115 | MYT1L | chr2:1943305-1943305 | C | CT | C | HOMO |

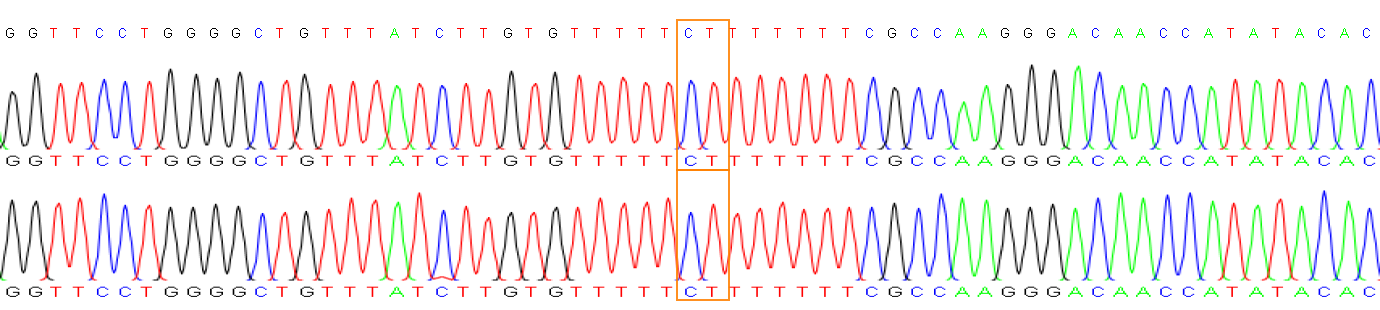

| **Sample** | **Gene** | **Chromosome position** | **Ref.** | **Alt.** | **Result** | **zygosity** |
| --- | --- | --- | --- | --- | --- | --- |
| P0118 | NOTCH3 | chr19:15187315-15187315 | G | A | G/A | HETERO |

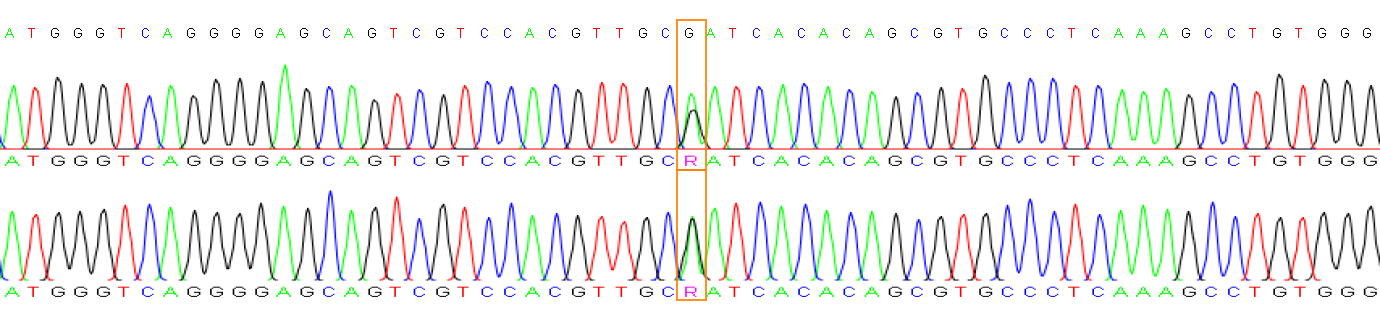

| **Sample** | **Gene** | **Chromosome position** | **Ref.** | **Alt.** | **Result** | **zygosity** |
| --- | --- | --- | --- | --- | --- | --- |
| P0120 | TRPM3 | chr9:70535987-70535987 | G | C | G/C | HETERO |

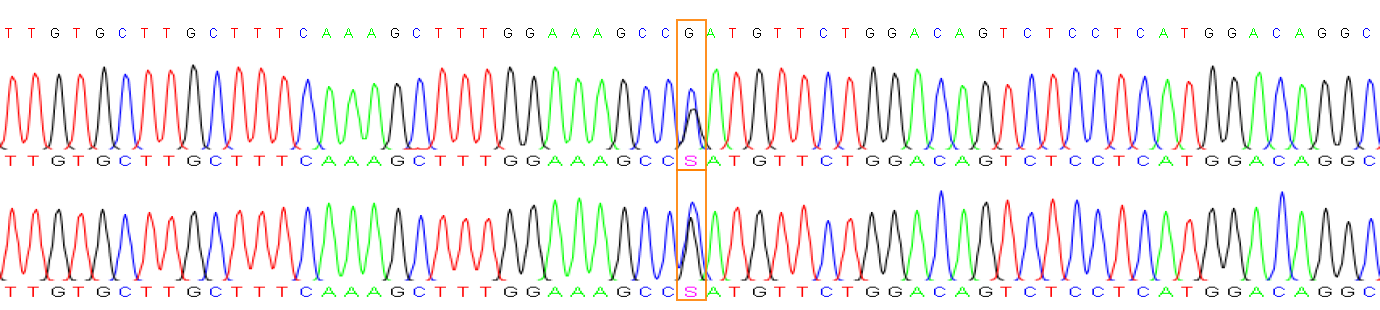

| **Sample** | **Gene** | **Chromosome position** | **Ref.** | **Alt.** | **Result** | **zygosity** |
| --- | --- | --- | --- | --- | --- | --- |
| P0120 | TMEM163 | chr2:134456738-134456738 | C | T | C/T | HETERO |

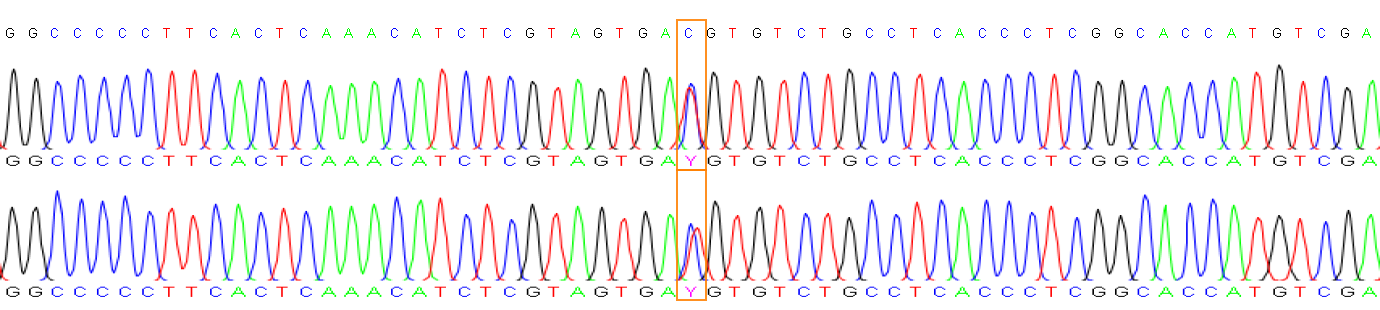

| **Sample** | **Gene** | **Chromosome position** | **Ref.** | **Alt.** | **Result** | **zygosity** |
| --- | --- | --- | --- | --- | --- | --- |
| P0123 | DYRK1A | chr21:37512470-37512470 | A | G | A/G | HETERO |

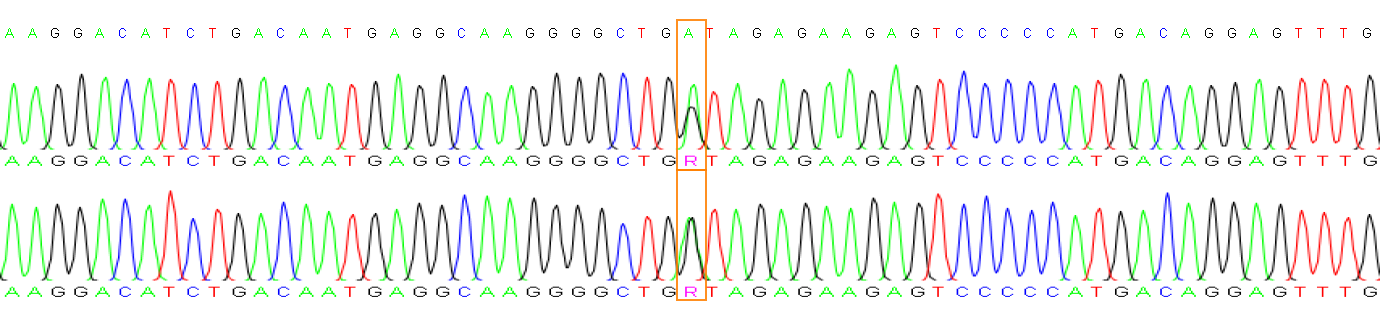

| **Sample** | **Gene** | **Chromosome position** | **Ref.** | **Alt.** | **Result** | **zygosity** |
| --- | --- | --- | --- | --- | --- | --- |
| P0130 | AHDC1-2 | chr1:27549697-27549697 | C | G | C/G | HETERO |

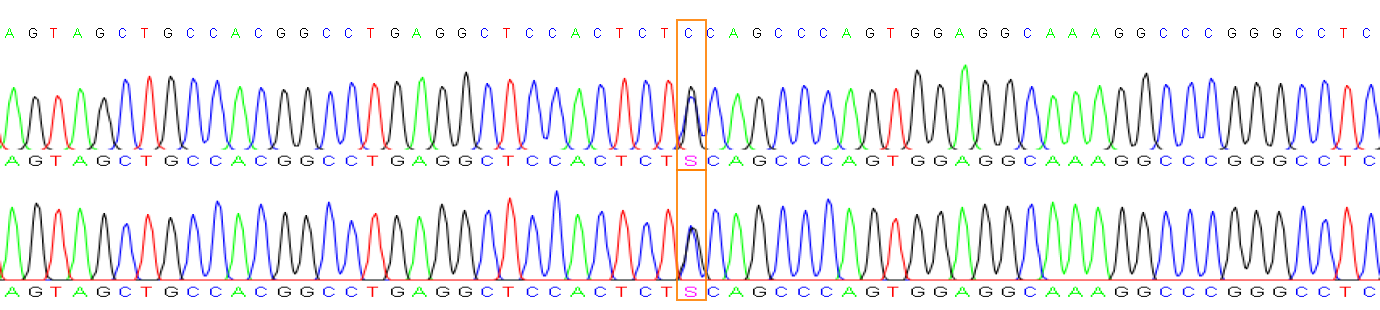

| **Sample** | **Gene** | **Chromosome position** | **Ref.** | **Alt.** | **Result** | **zygosity** |
| --- | --- | --- | --- | --- | --- | --- |
| P0137 | TSC2 | chr16:2085261-2085261 | T | G | T/G | HETERO |

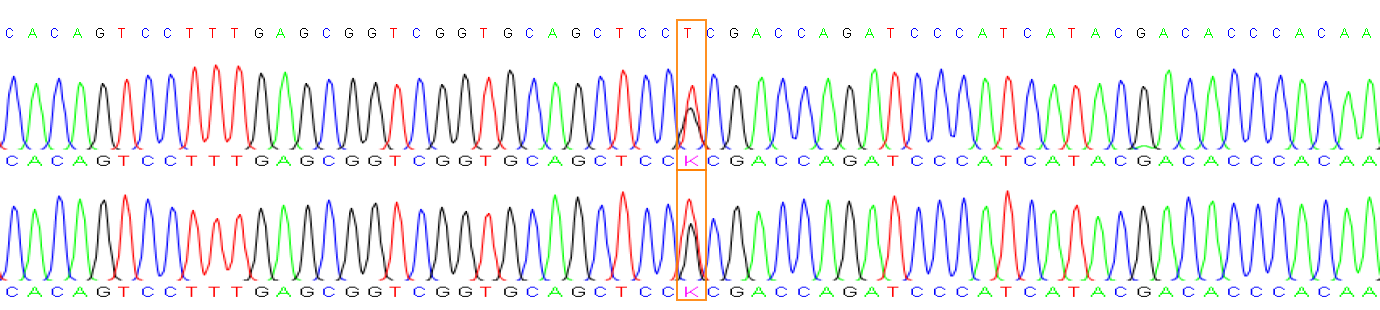

| **Sample** | **Gene** | **Chromosome position** | **Ref.** | **Alt.** | **Result** | **zygosity** |
| --- | --- | --- | --- | --- | --- | --- |
| P0201 | PDE2A | chr11:72579374-72579374 | G | A | A | HOMO |

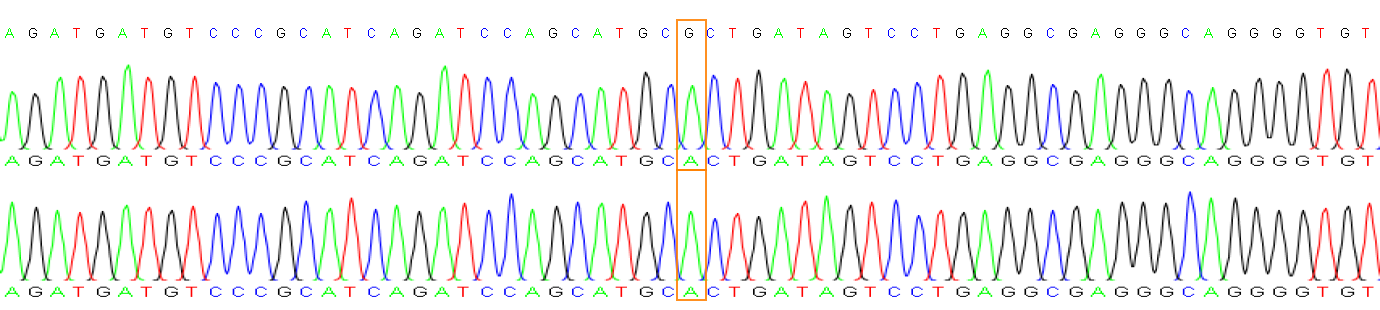

| **Sample** | **Gene** | **Chromosome position** | **Ref.** | **Alt.** | **Result** | **zygosity** |
| --- | --- | --- | --- | --- | --- | --- |
| P0201 | DOCK11 | chrX:118543010-118543010 | A | G | G | HOMO |

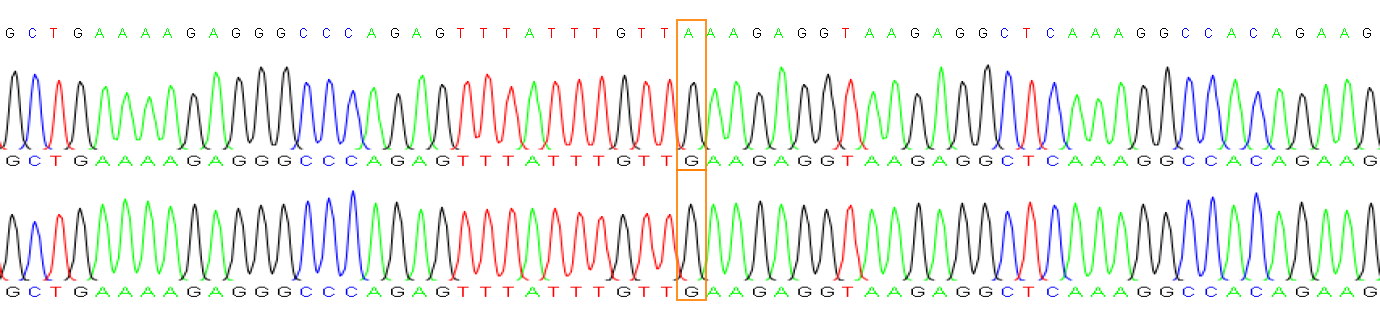

| **Sample** | **Gene** | **Chromosome position** | **Ref.** | **Alt.** | **Result** | **zygosity** |
| --- | --- | --- | --- | --- | --- | --- |
| P0201 | RORB | chr9:74634734-74634734 | G | A | G/A | HETERO |

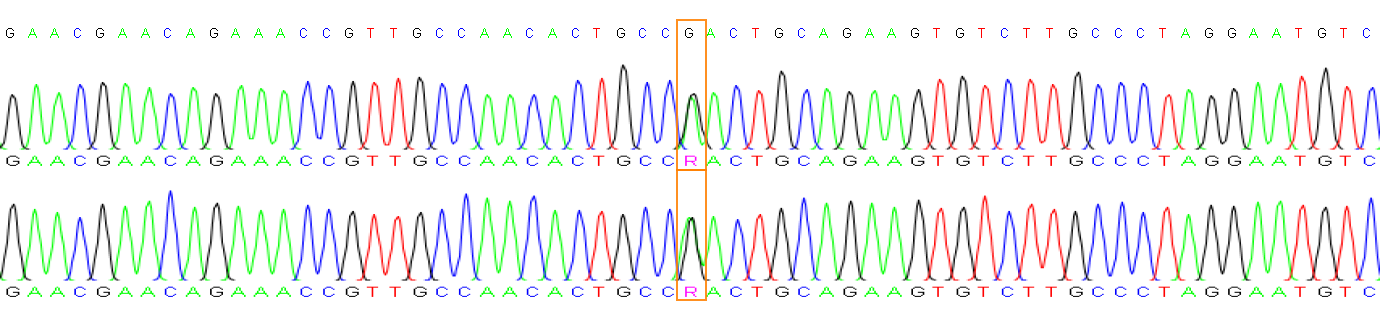

| **Sample** | **Gene** | **Chromosome position** | **Ref.** | **Alt.** | **Result** | **zygosity** |
| --- | --- | --- | --- | --- | --- | --- |
| P0201 | OPA1 | chr3:193643378-193643378 | A | G | A/G | HETERO |

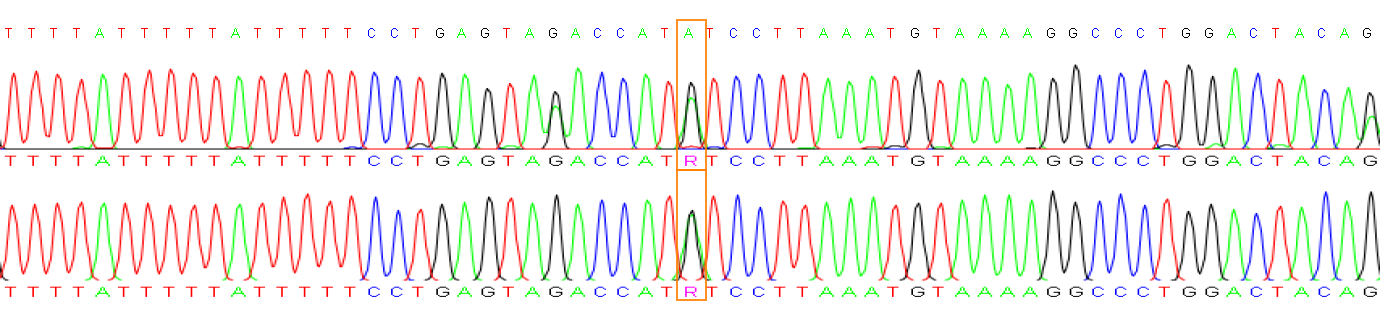

| **Sample** | **Gene** | **Chromosome position** | **Ref.** | **Alt.** | **Result** | **zygosity** |
| --- | --- | --- | --- | --- | --- | --- |
| P0207 | CCM2 | chr7:45064647-45064647 | G | C | G/C | HETERO |

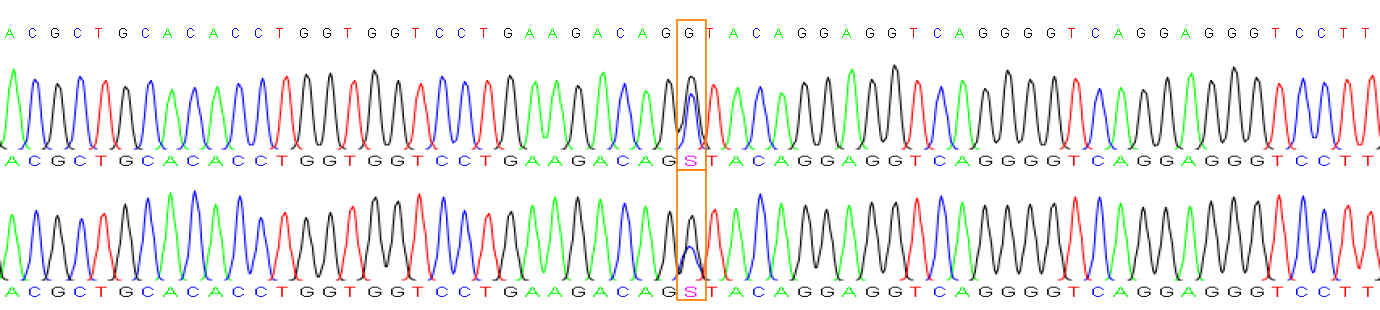

| **Sample** | **Gene** | **Chromosome position** | **Ref.** | **Alt.** | **Result** | **zygosity** |
| --- | --- | --- | --- | --- | --- | --- |
| P0263 | KMT2B | chr19:35726350-35726350 | A | C | A/C | HETERO |

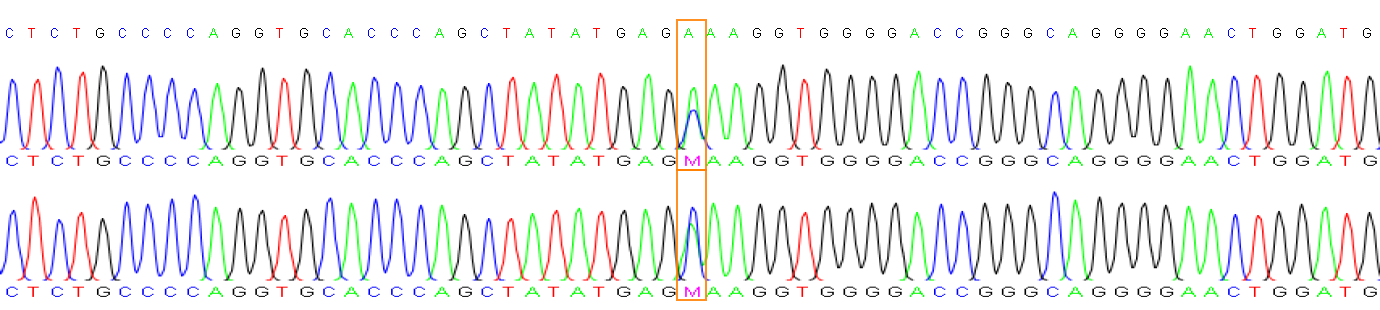

| **Sample** | **Gene** | **Chromosome position** | **Ref.** | **Alt.** | **Result** | **zygosity** |
| --- | --- | --- | --- | --- | --- | --- |
| P0269 | KMT2E-3 | chr7:105081692-105081692 | G | A | G/A | HETERO |

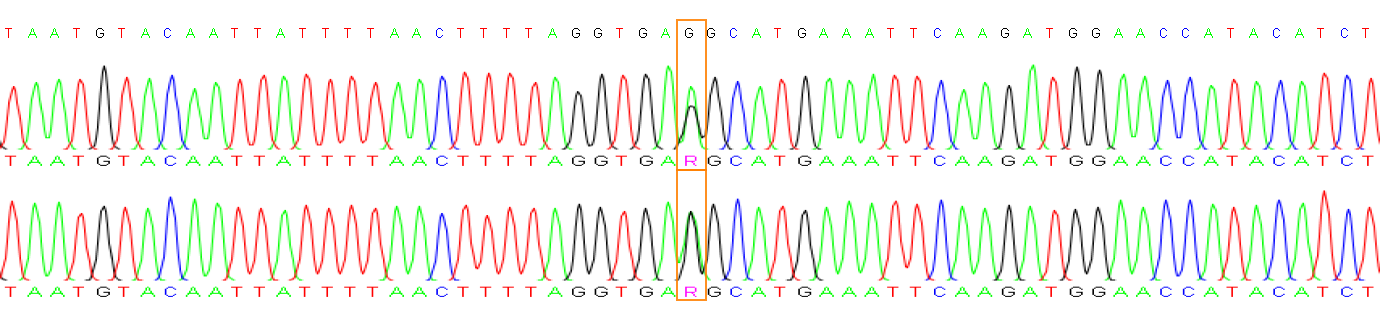

| **Sample** | **Gene** | **Chromosome position** | **Ref.** | **Alt.** | **Result** | **zygosity** |
| --- | --- | --- | --- | --- | --- | --- |
| P0260 | PLA2G6-1 | chr22:38132952-38132952 | G | A | G/A | HETERO |

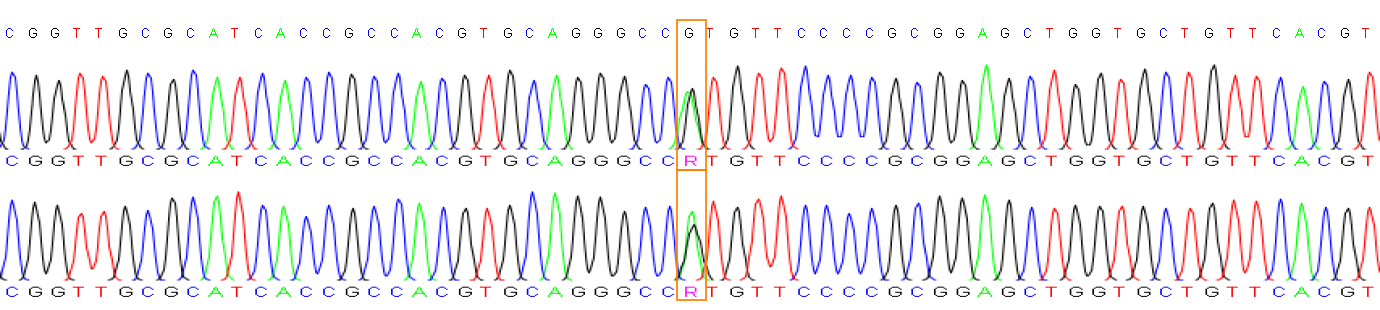

| **Sample** | **Gene** | **Chromosome position** | **Ref.** | **Alt.** | **Result** | **zygosity** |
| --- | --- | --- | --- | --- | --- | --- |
| P0260 | PLA2G6-2 | chr22:38143253-38143253 | G | A | G/A | HETERO |

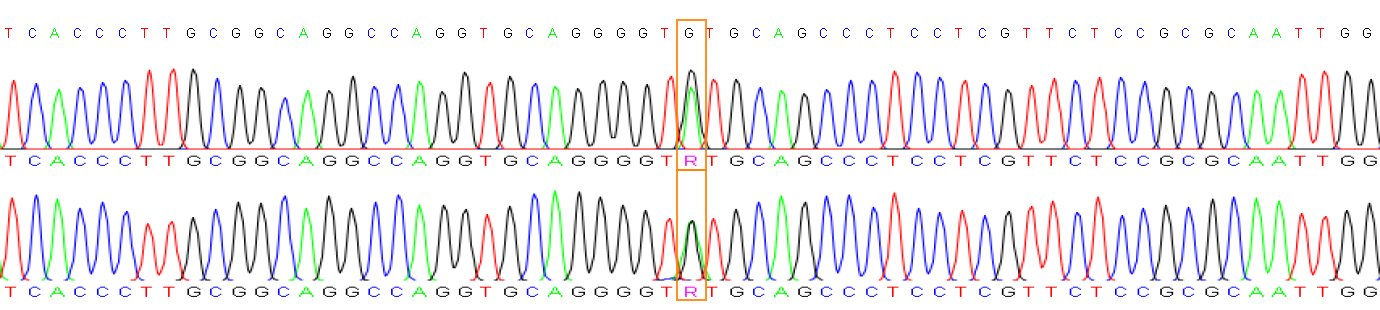

| **Sample** | **Gene** | **Chromosome position** | **Ref.** | **Alt.** | **Result** | **zygosity** |
| --- | --- | --- | --- | --- | --- | --- |
| P0255 | CC2D2A-1 | chr4:15574195-15574195 | T | C | T/C | HETERO |

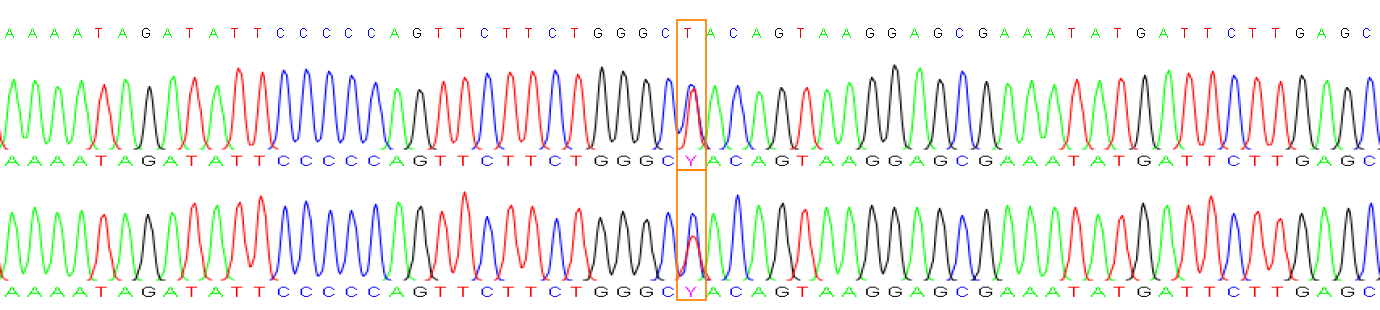

| **Sample** | **Gene** | **Chromosome position** | **Ref.** | **Alt.** | **Result** | **zygosity** |
| --- | --- | --- | --- | --- | --- | --- |
| P0255 | CC2D2A-2 | chr4:15601348-15601348 | G | A | G/A | HETERO |

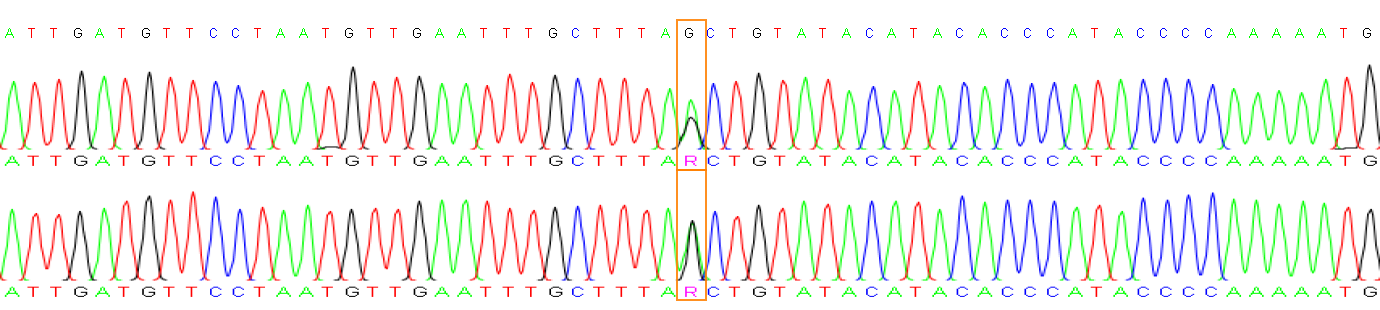

**IGV Screenshots**

Variants in 56 samples across 92 loci (74 genes)

**P0001 (chr19:8,111,006-8,111,106) FBN3**

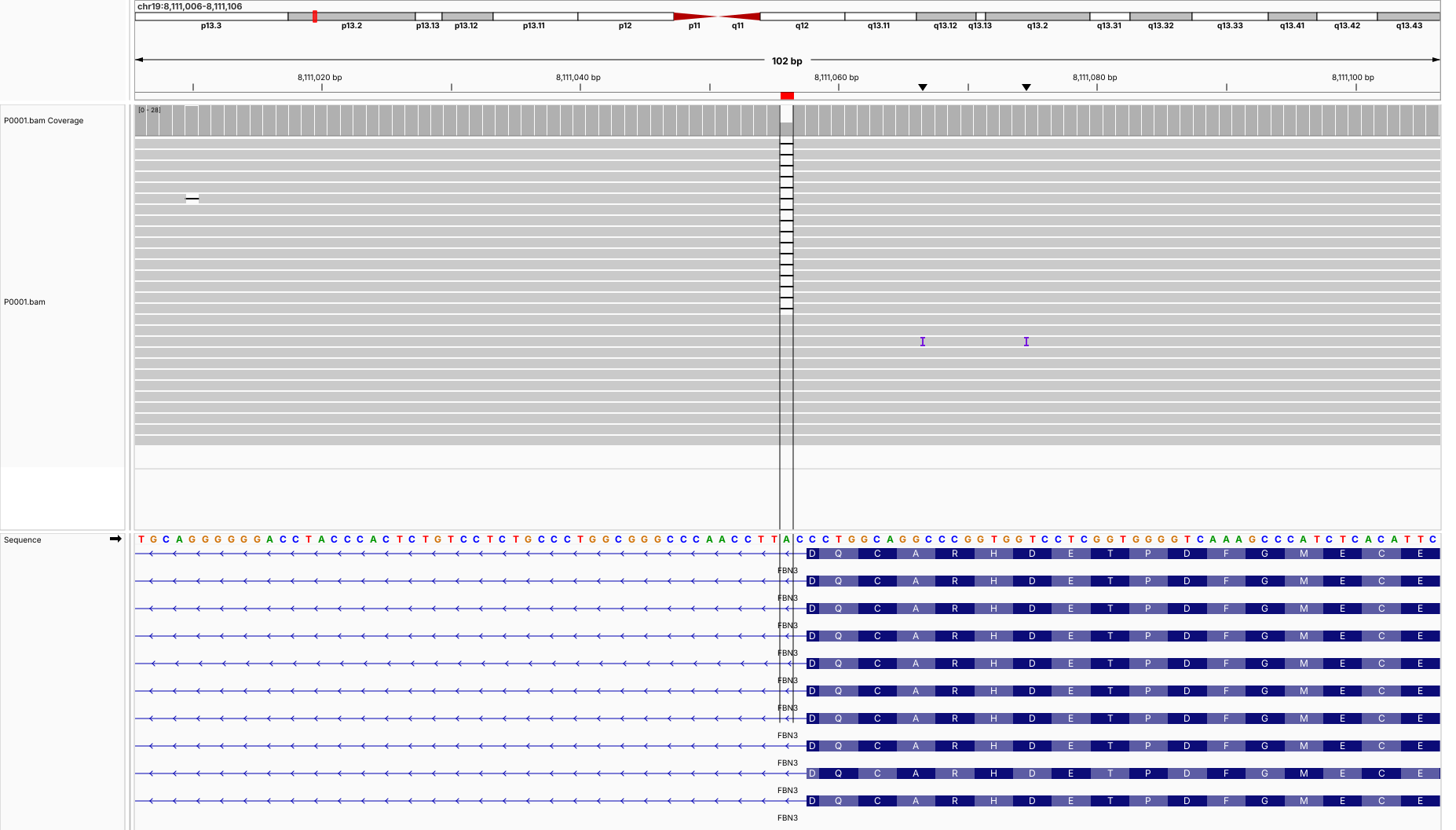

**P0001 (chr19:8,129,037-8,129,137) FBN3**

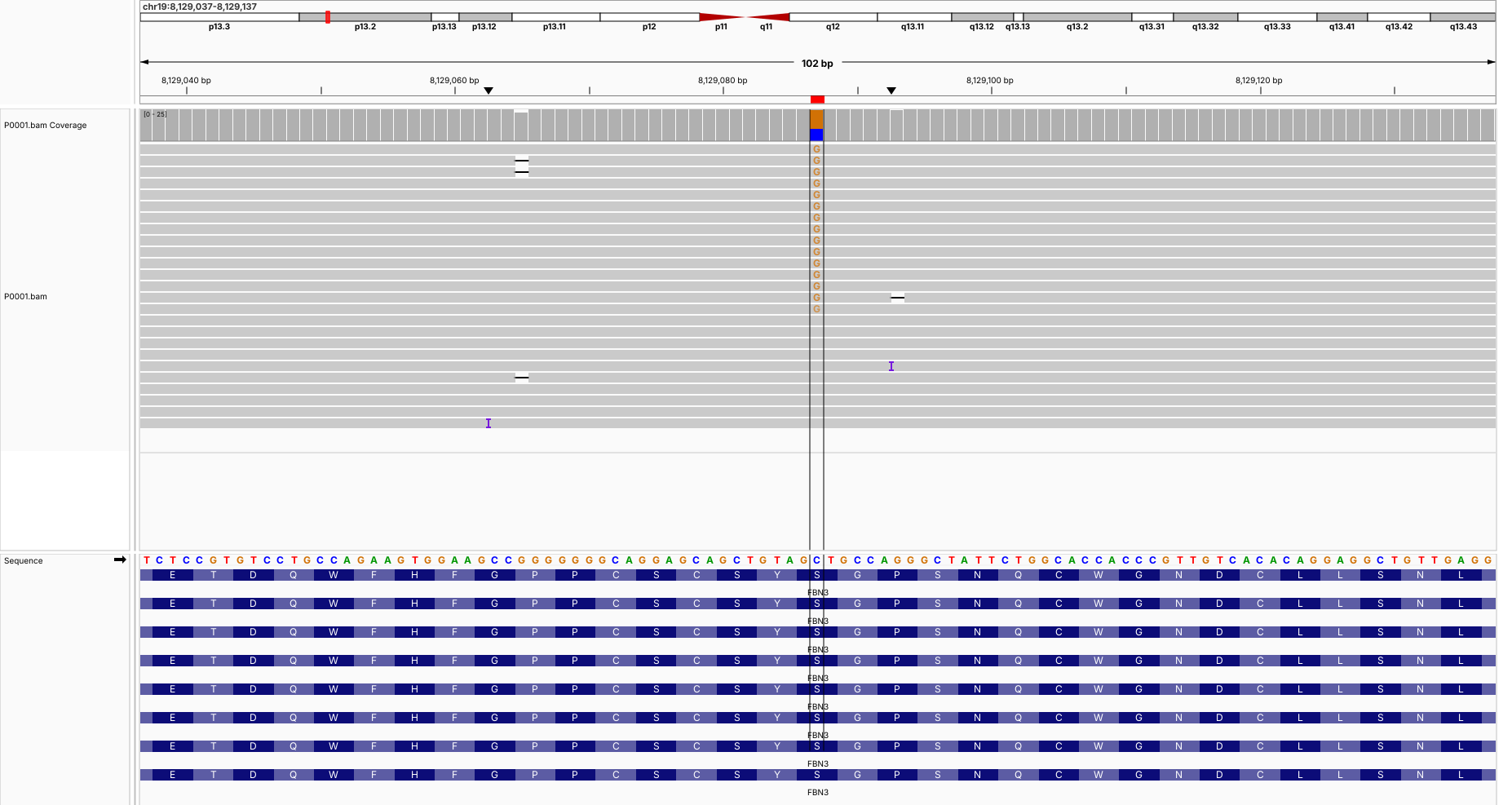

**P0010 (chr11:64,690,396-64,690,496) NRXN2**

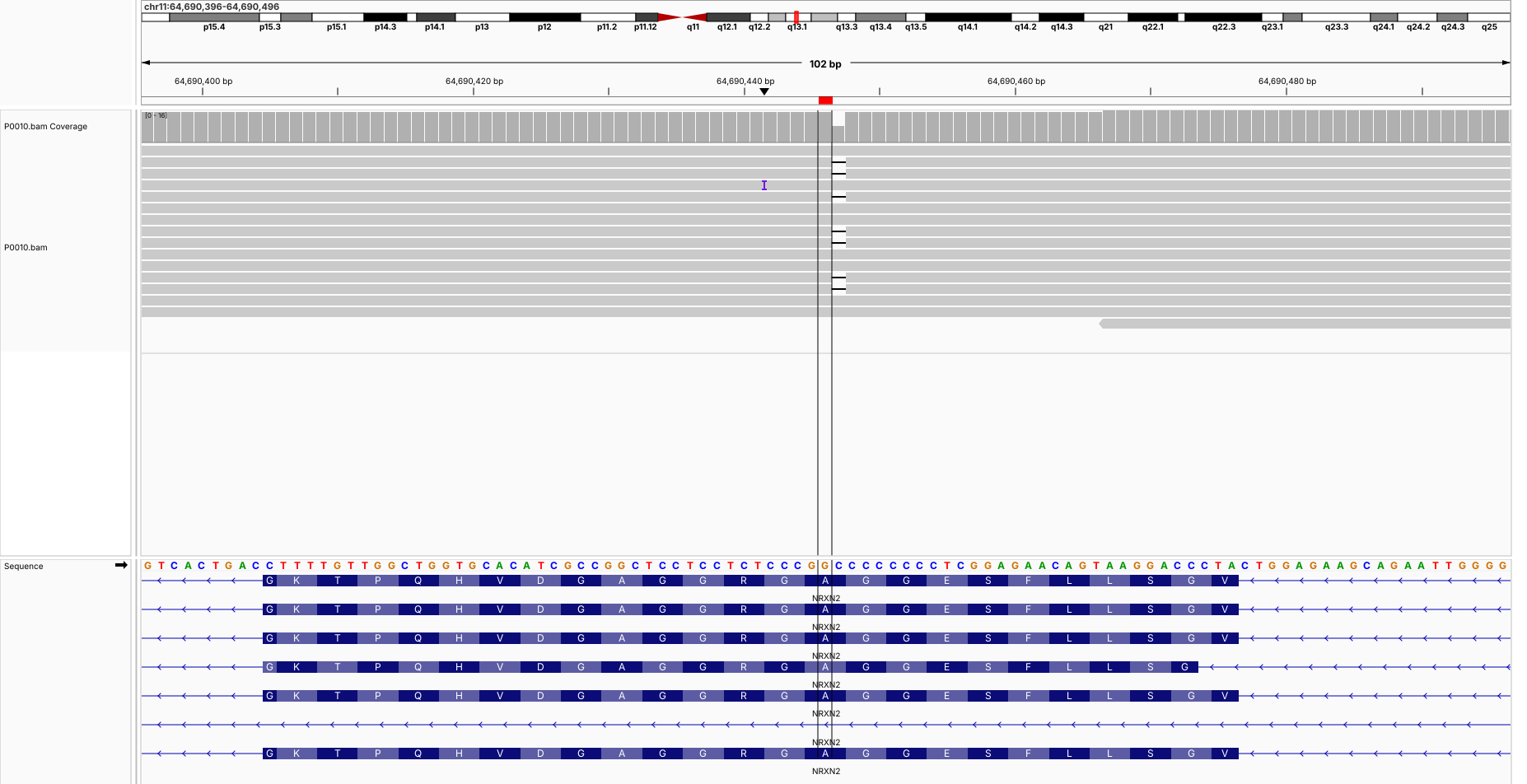

**P0013 (chr13:110,466,006-110,466,106) COL4A2**

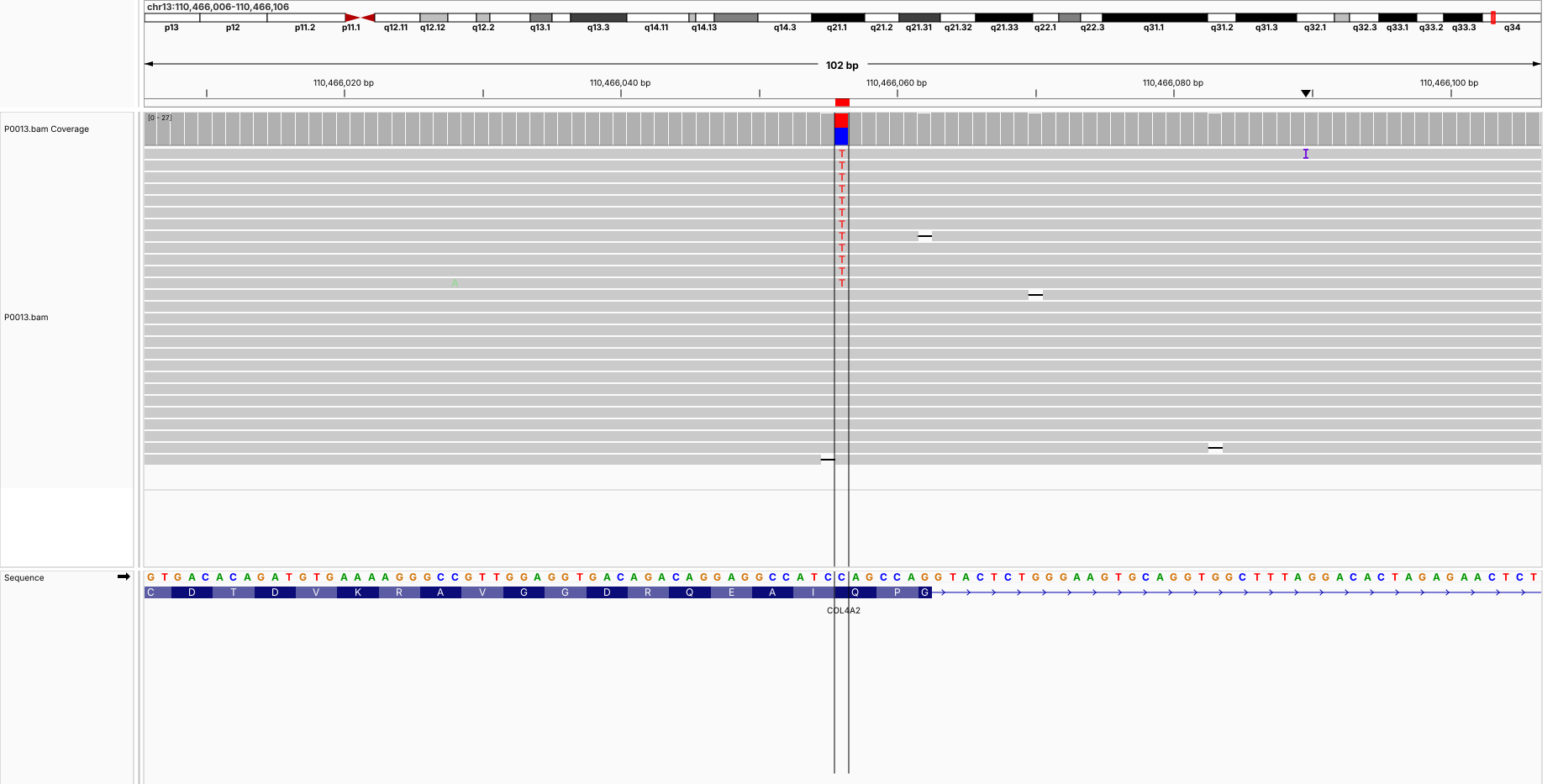

**P0021 (chr9:128,632,236-128,632,336) SPTAN1**

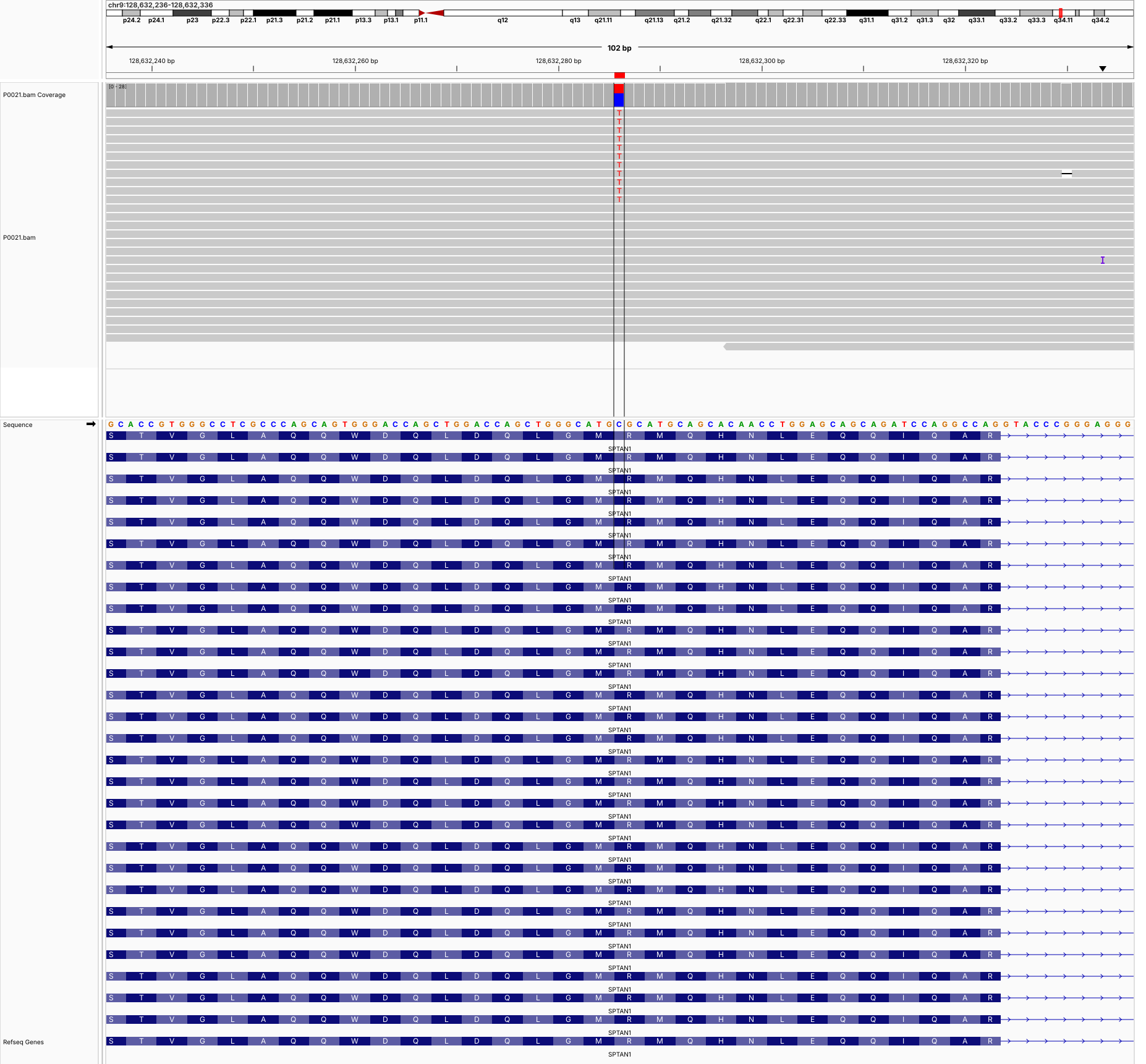

**P0024 (chr2:17,931,991-17,932,091) KCNS3**

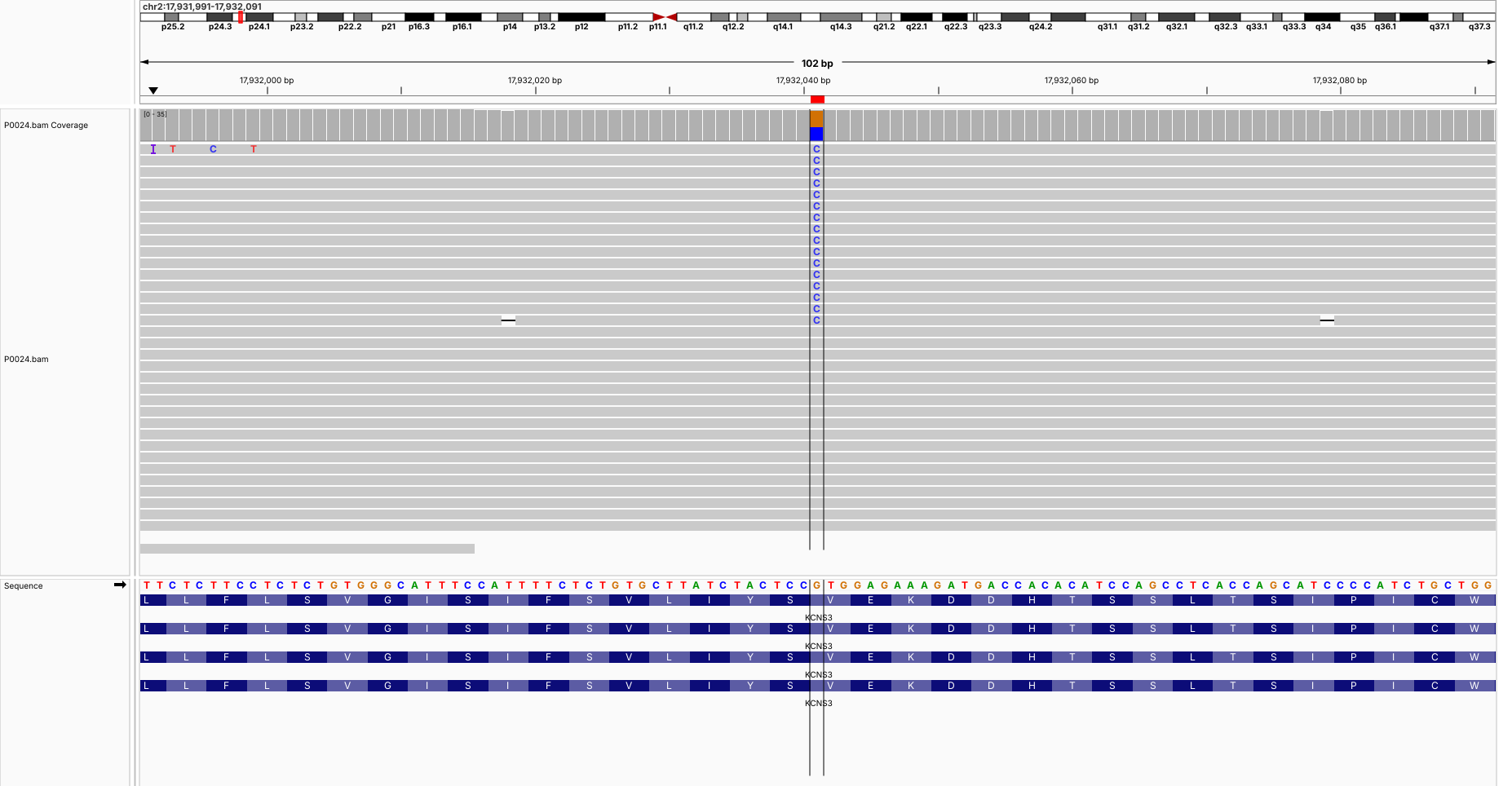

**P0033 (chrX:132,629,012-132,629,112) HS6ST2**

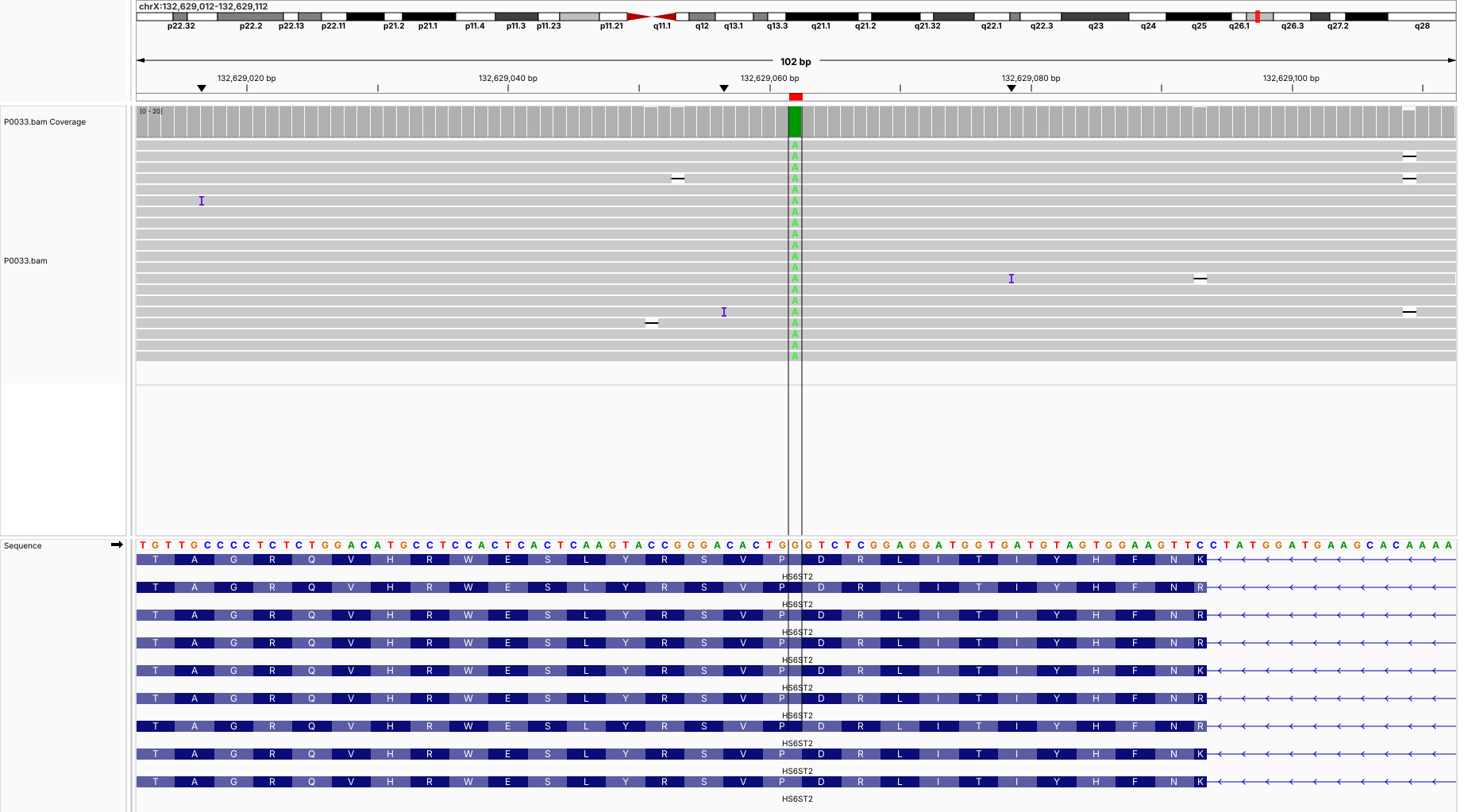

**P0036 (chr18:23,539,809-23,539,909) NPC1**

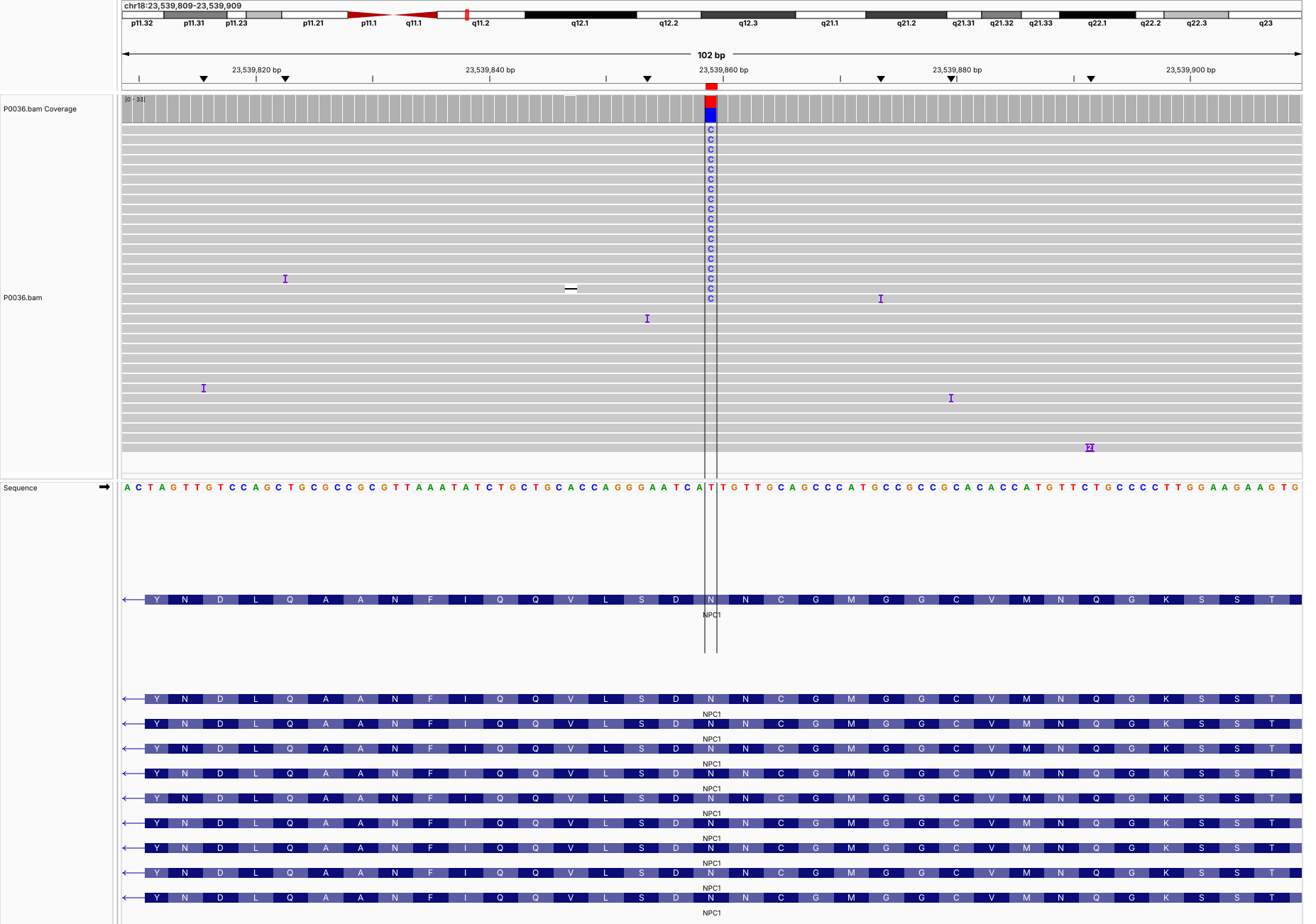

**P0036 (chr18:23,541,360-23,541,460) NPC1**

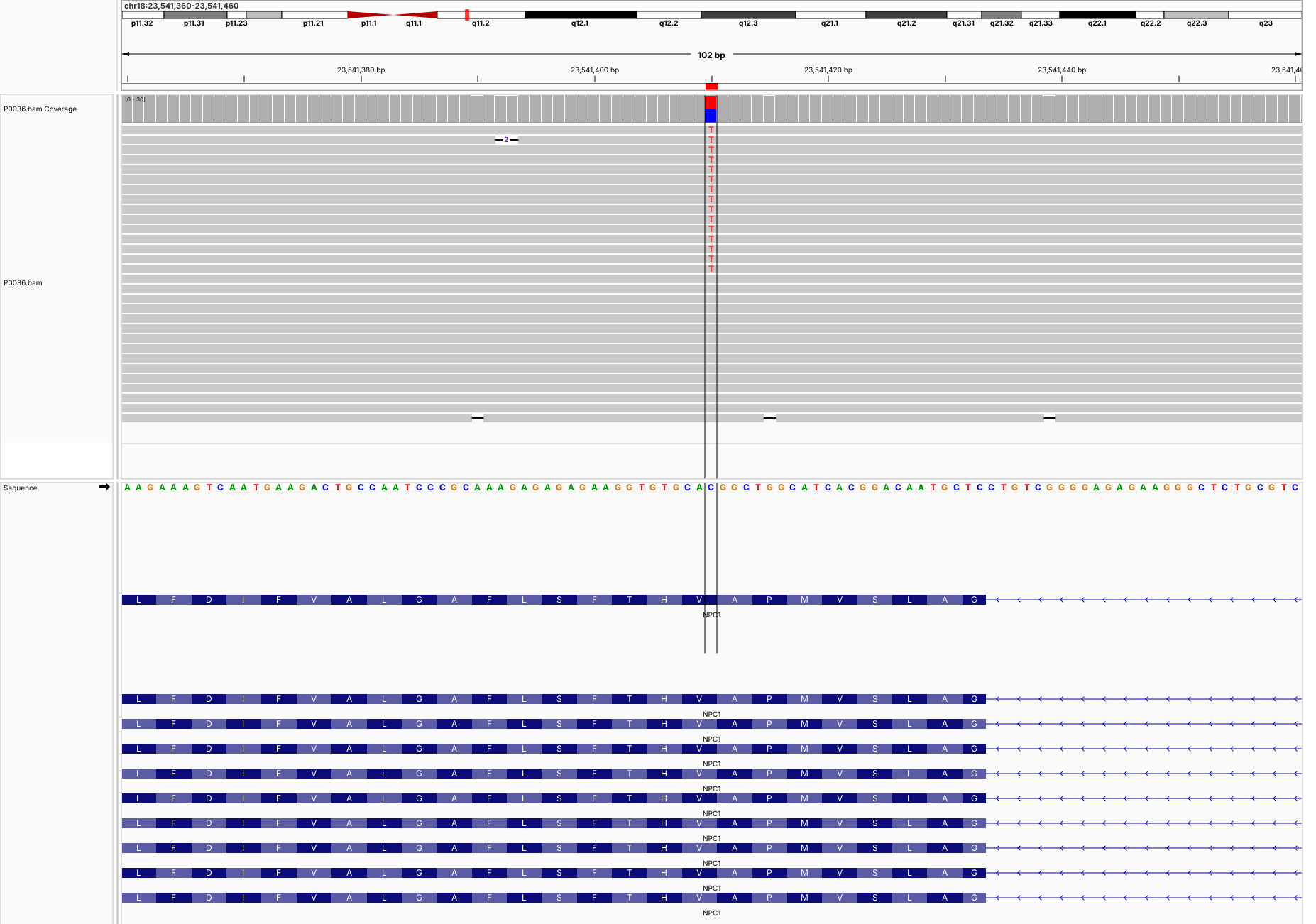

**P0050 (chr17:63,411,590-63,411,690) TANC2**

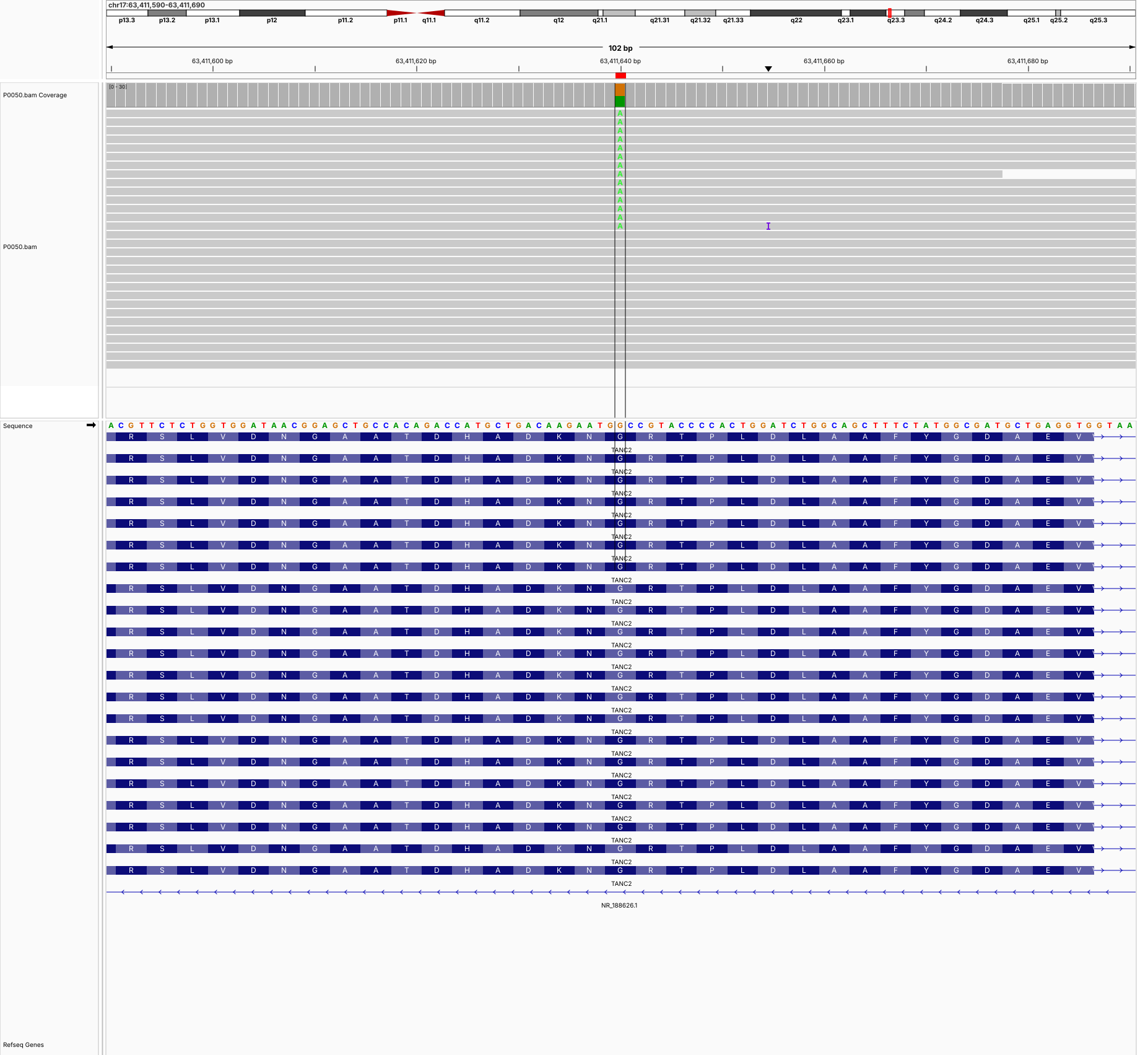

**P0069 (chr5:37,000,458-37,000,558) NIPBL**

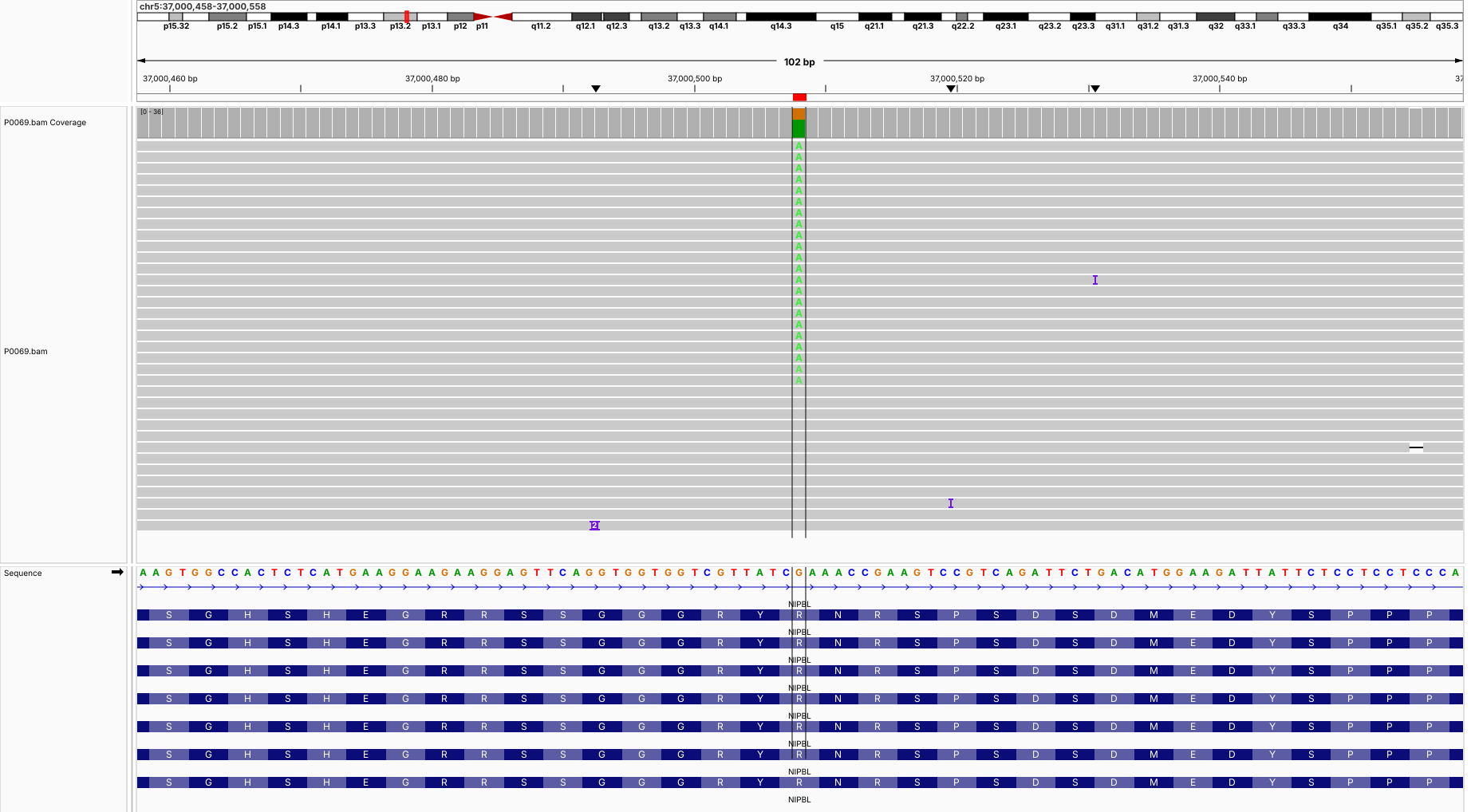

**P0069 (chr7:66,092,028-66,092,128) ASL**

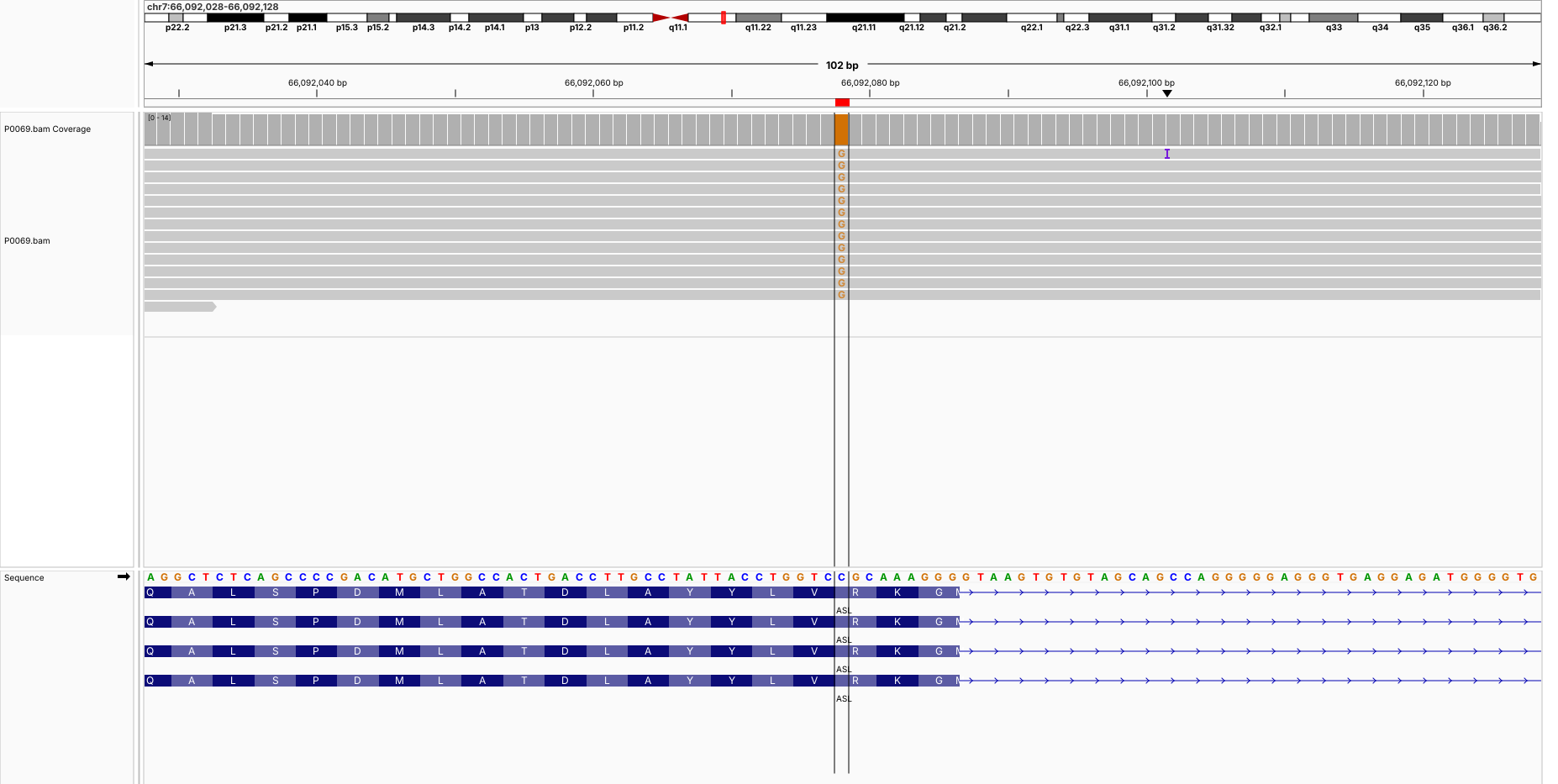

**P0070 (chr3:184,346,702-184,346,802) CLCN2**

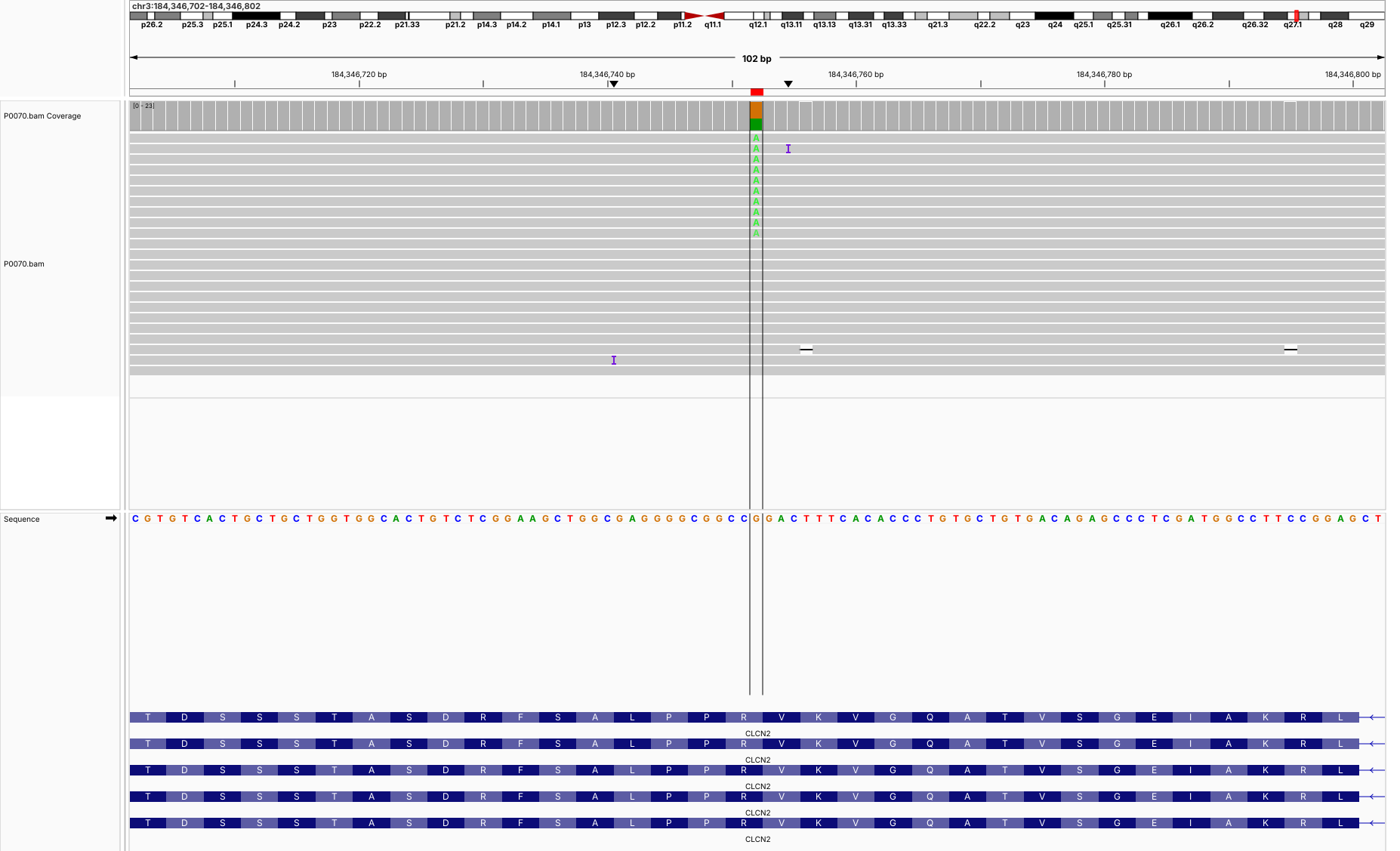

**P0071 (chrX:41,635,657-41,635,757) CASK**

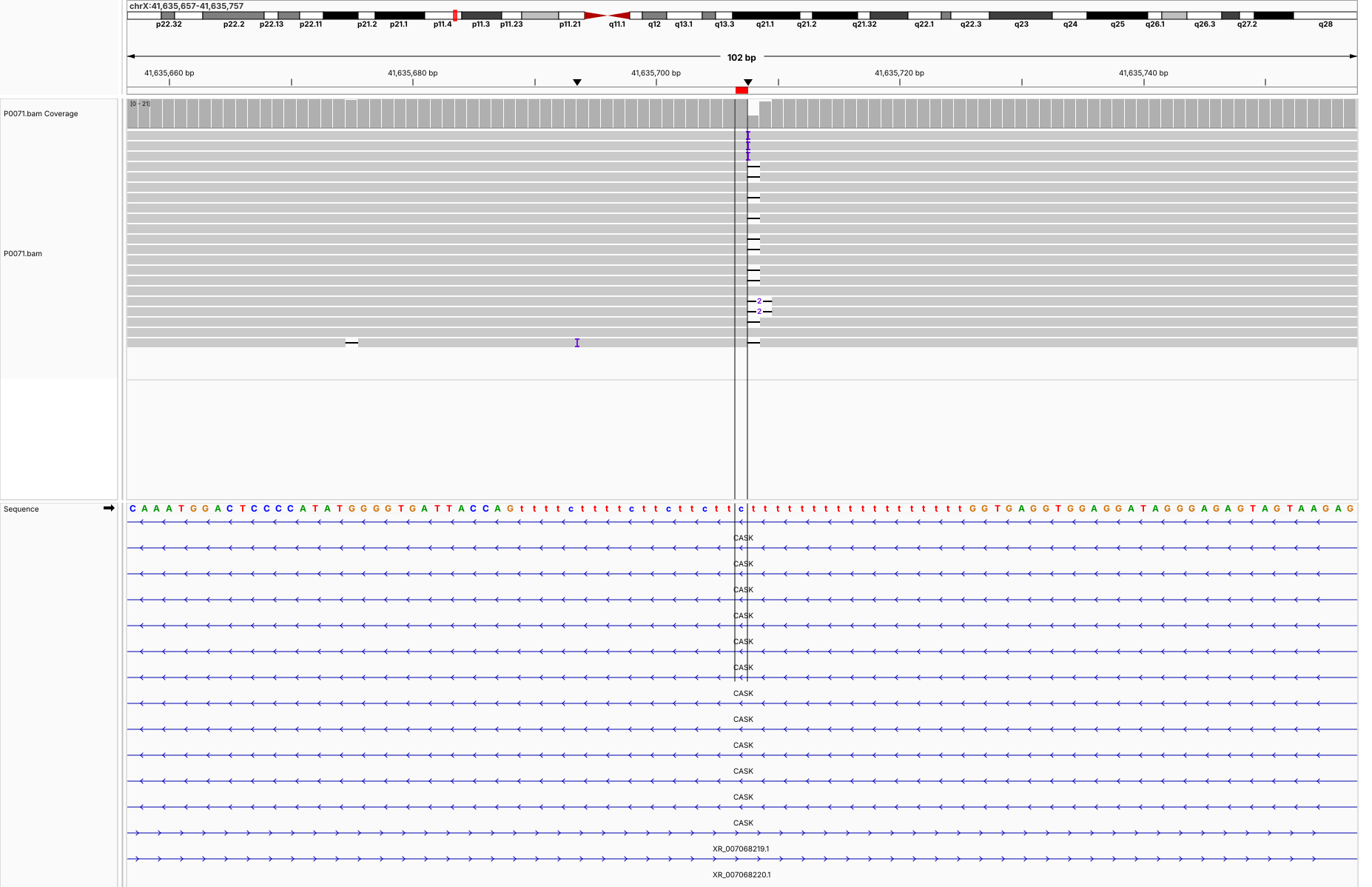

**P0073 (chr19:47,698,643-47,698,743) BICRA**

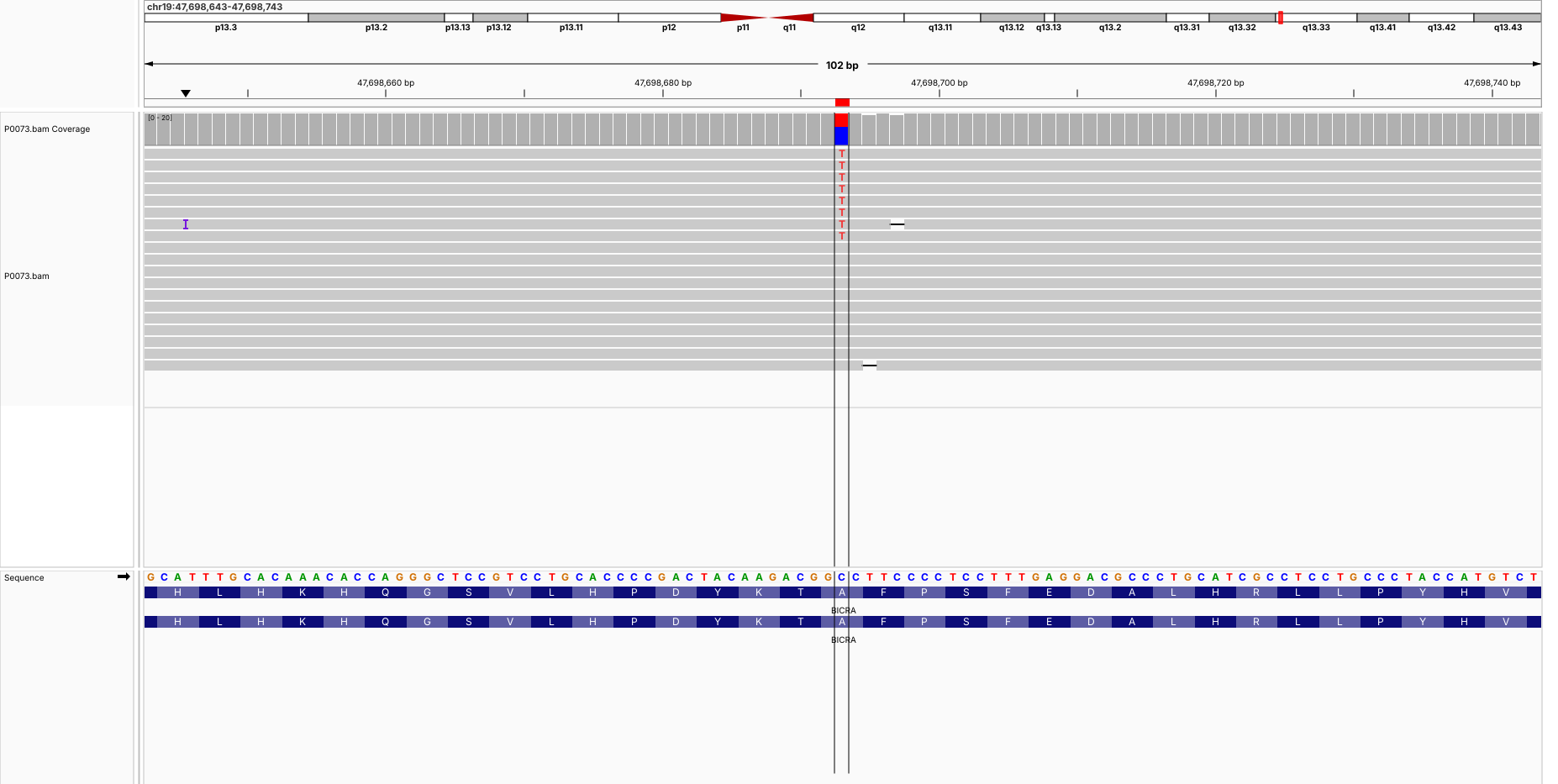

**P0074 (chr5:36,116,452-36,116,552) LMBRD2**

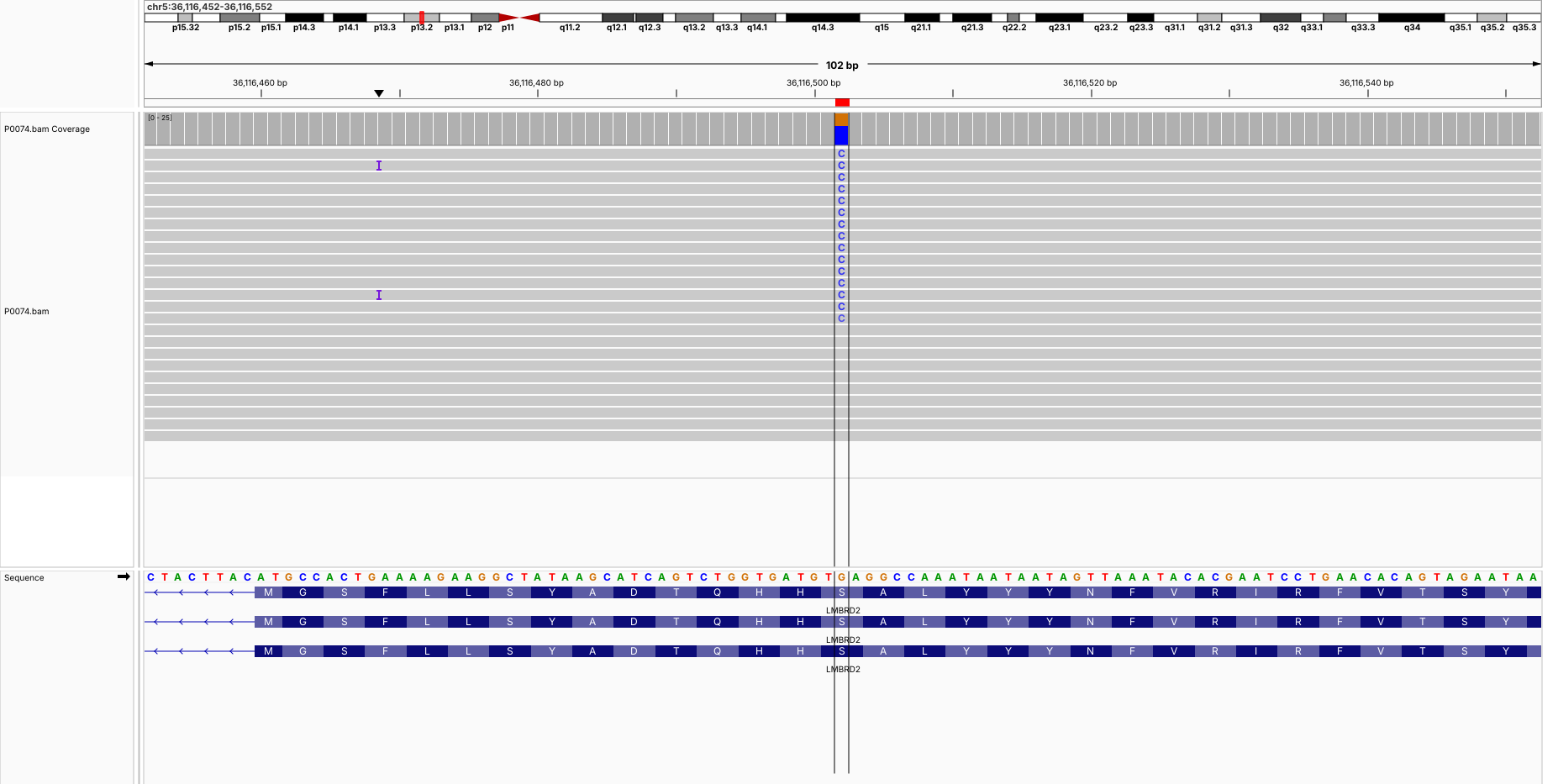

**P0076 (chr12:78,864,866-78,864,966) SYT1**

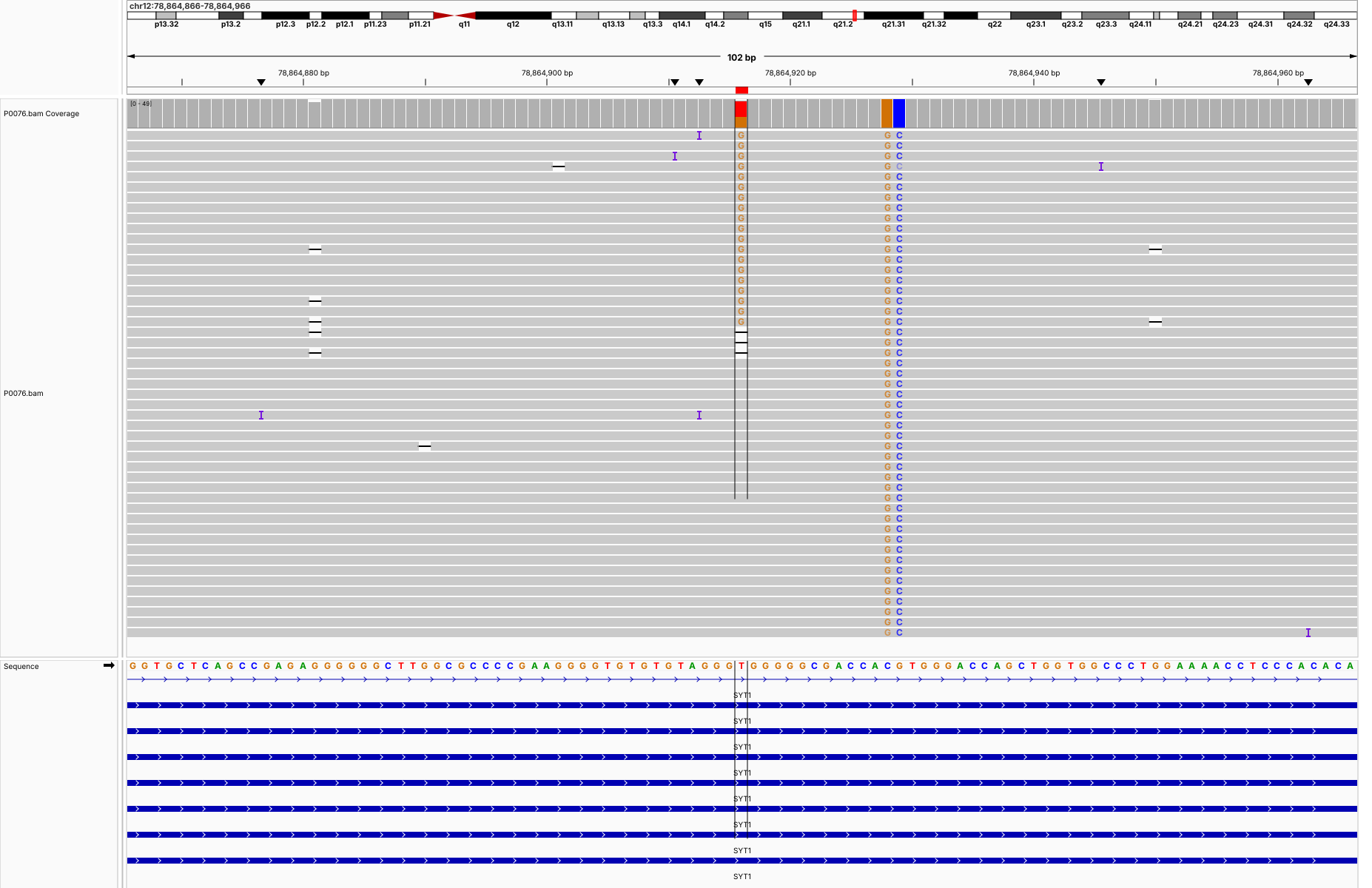

**P0076 (chr17:7,849,201-7,849,301) KDM6B**

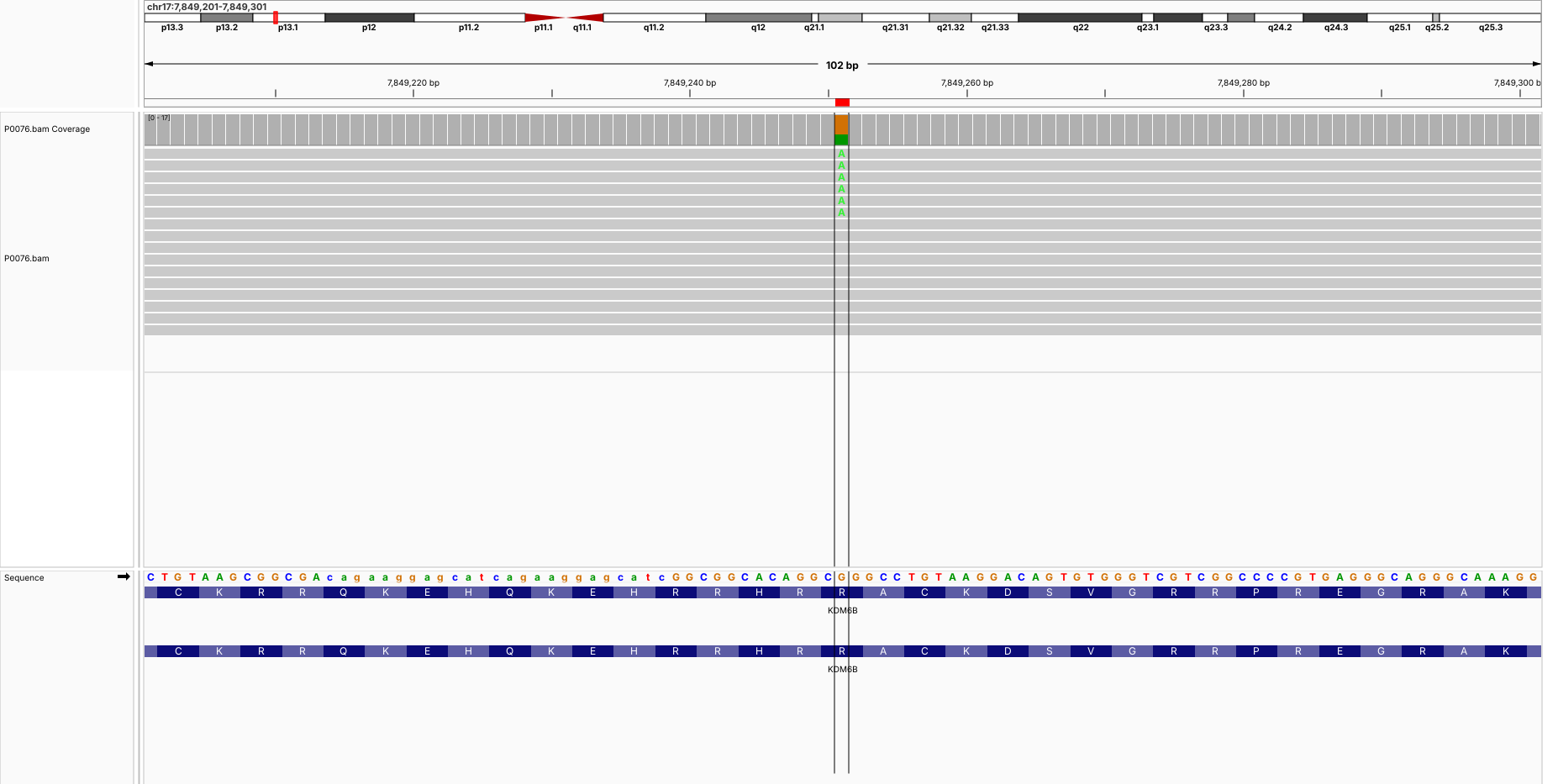

**P0076 (chr22:40,312,545-40,312,645) TNRC6B**

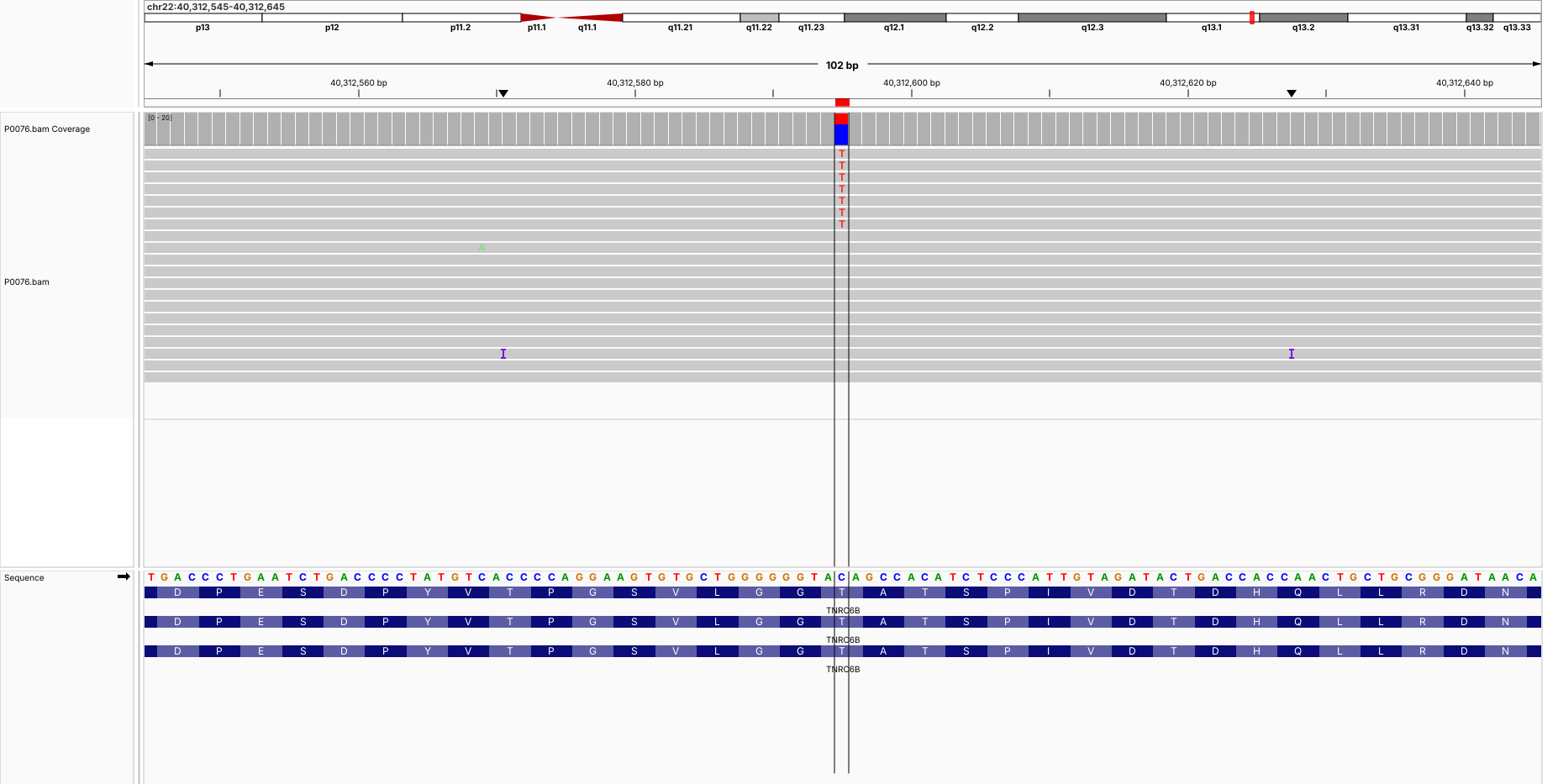

**P0078 (chr9:137,156,978-137,157,078) GRIN1**

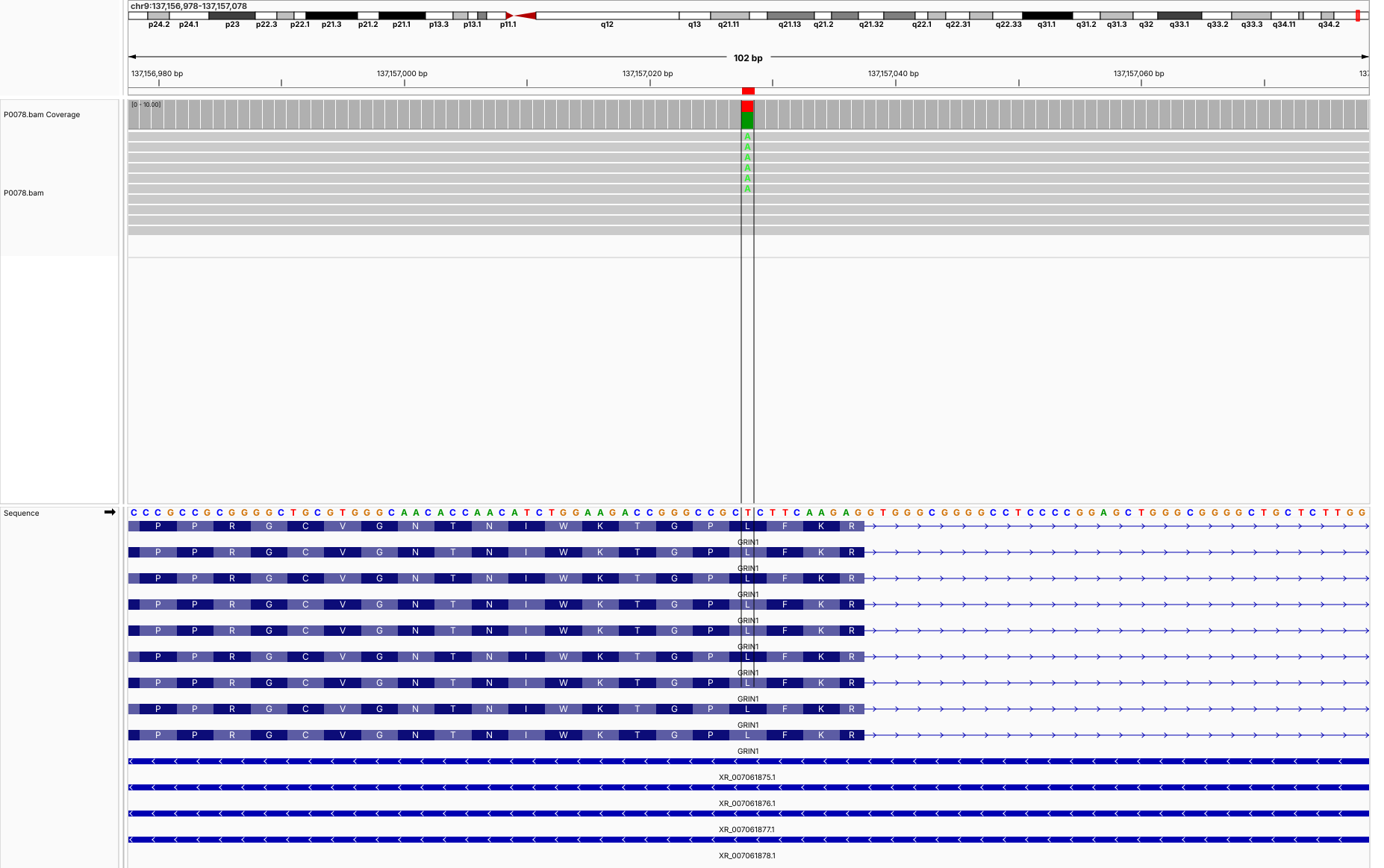

**P0078 (chr15:26,939,239-26,939,339) GABRB3**

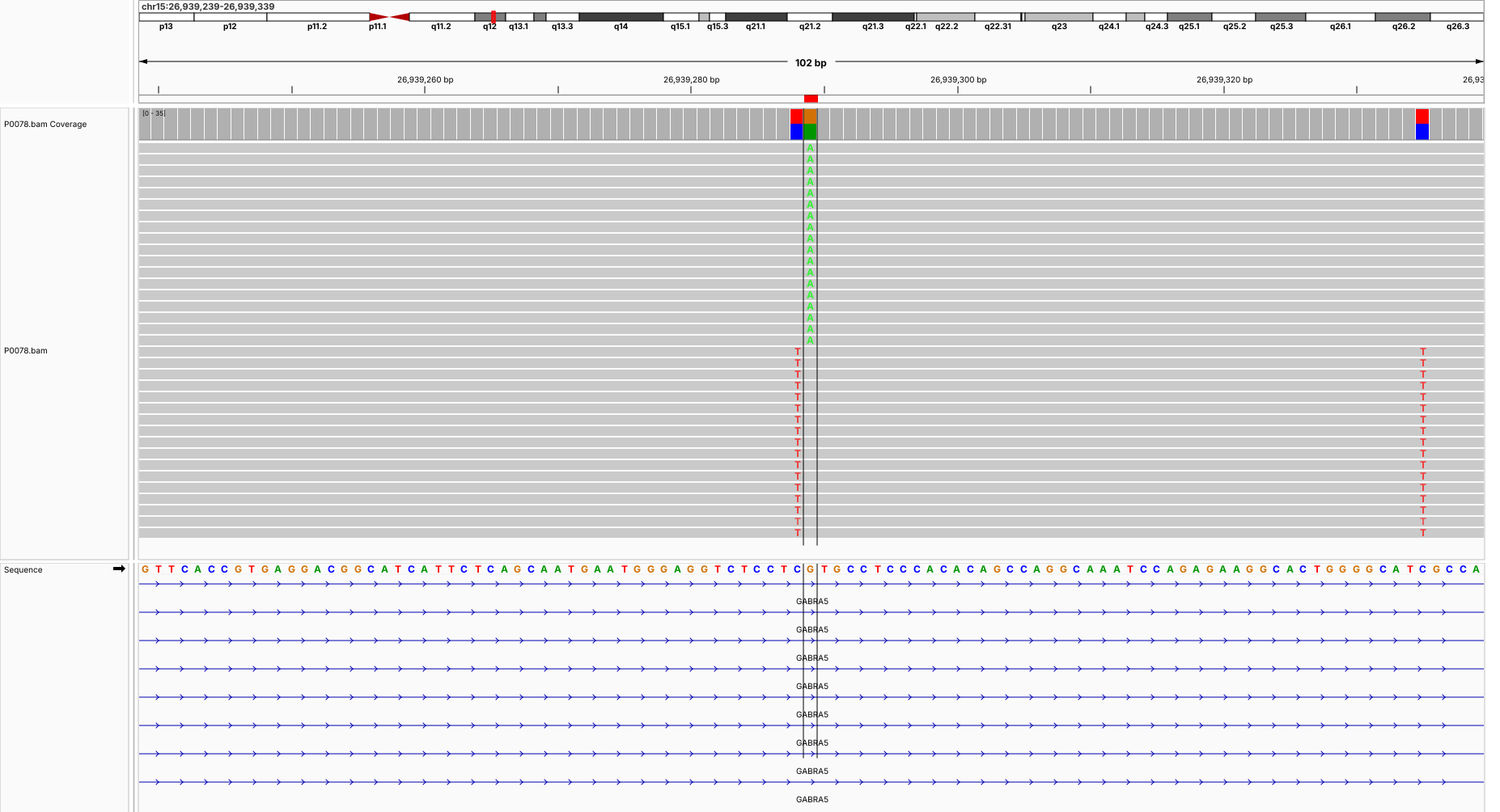

**P0079 (chr17:7,899,006-7,899,106) CHD3**

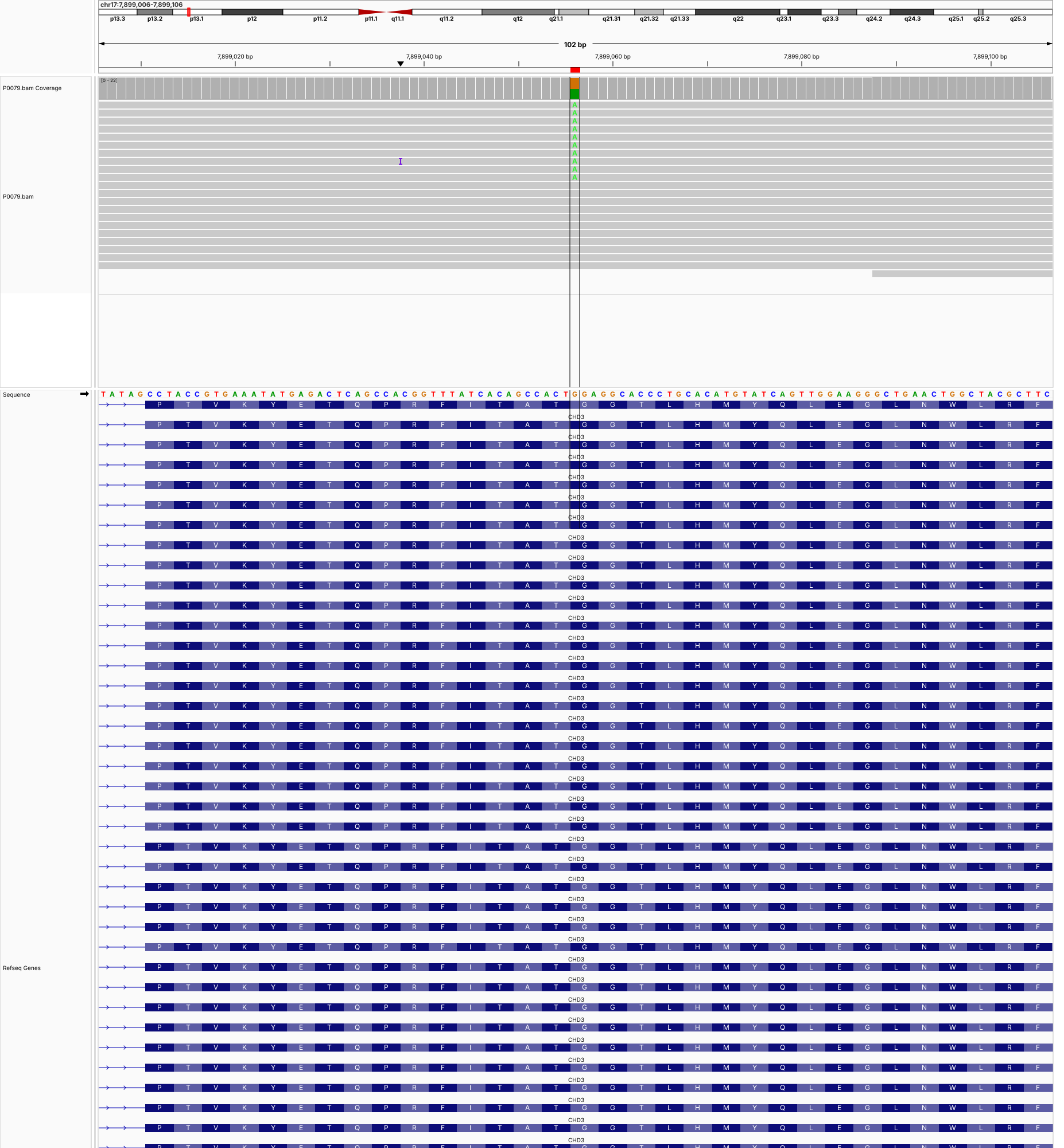

**P0079 (chr17:62,052,572-62,052,672) MED13**

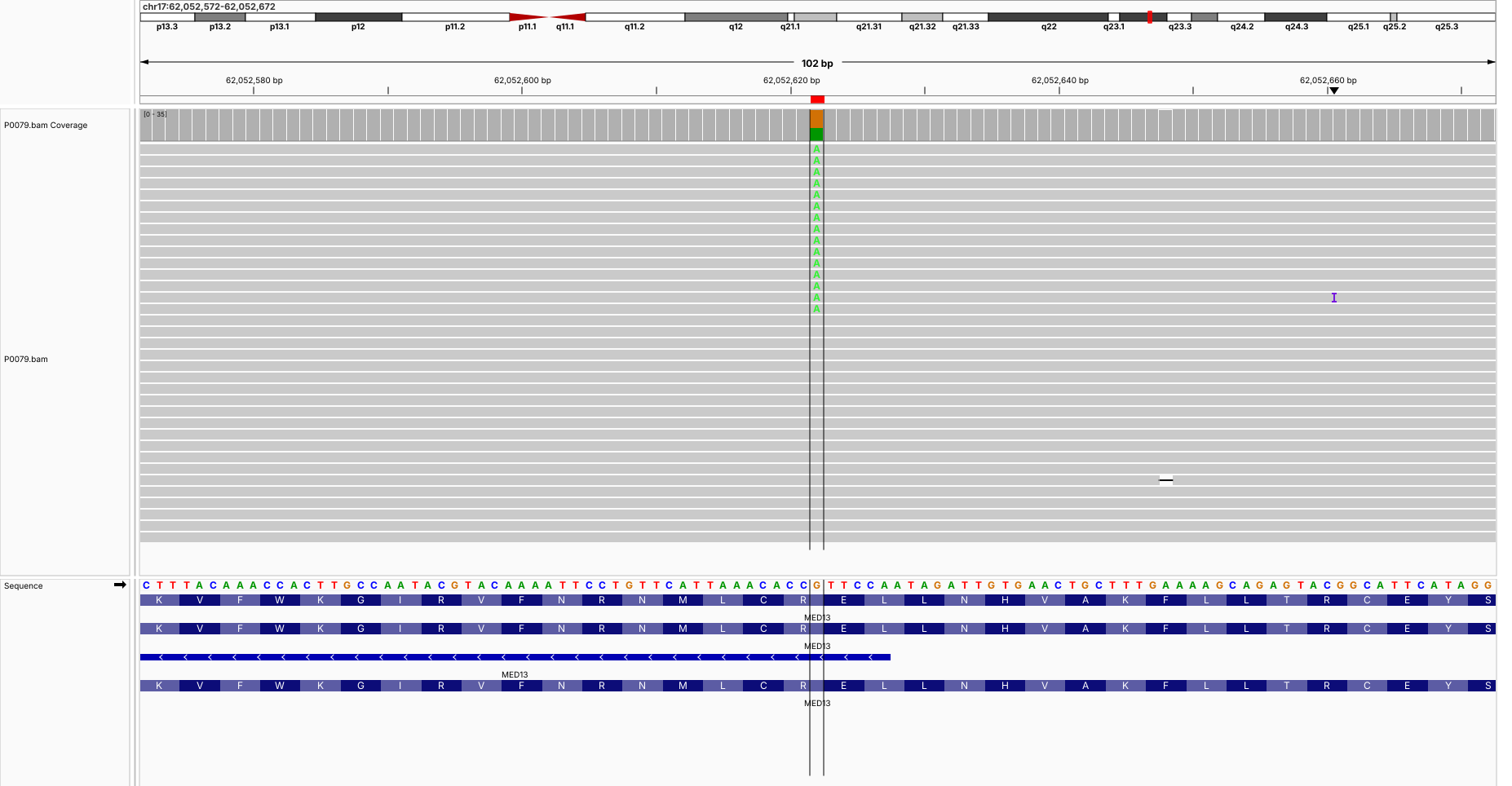

**P0081 (chr9:137,139,104-137,139,204) GRIN1**

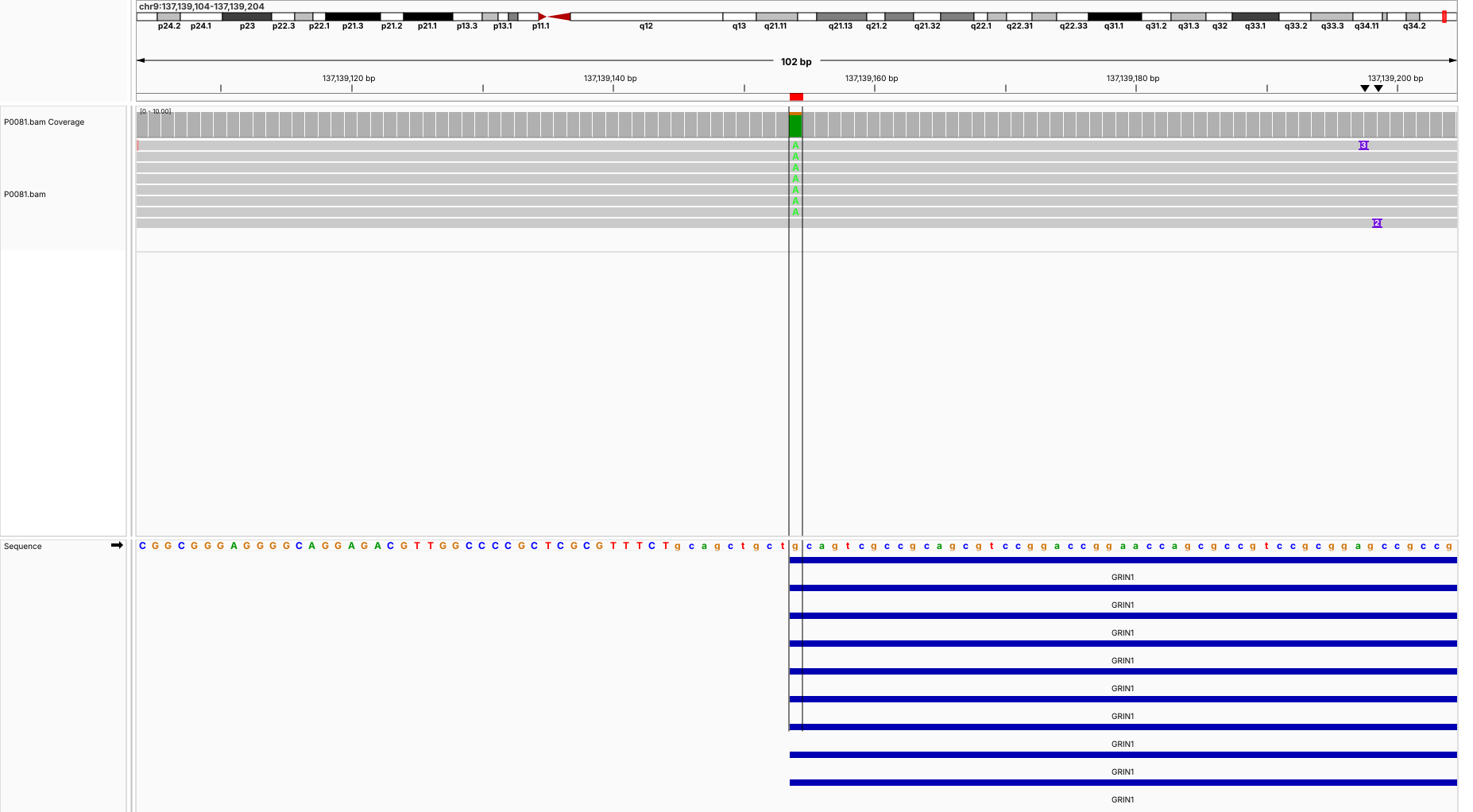

**P0081 (chr15:43,874,199-43,874,299) FRMD5**

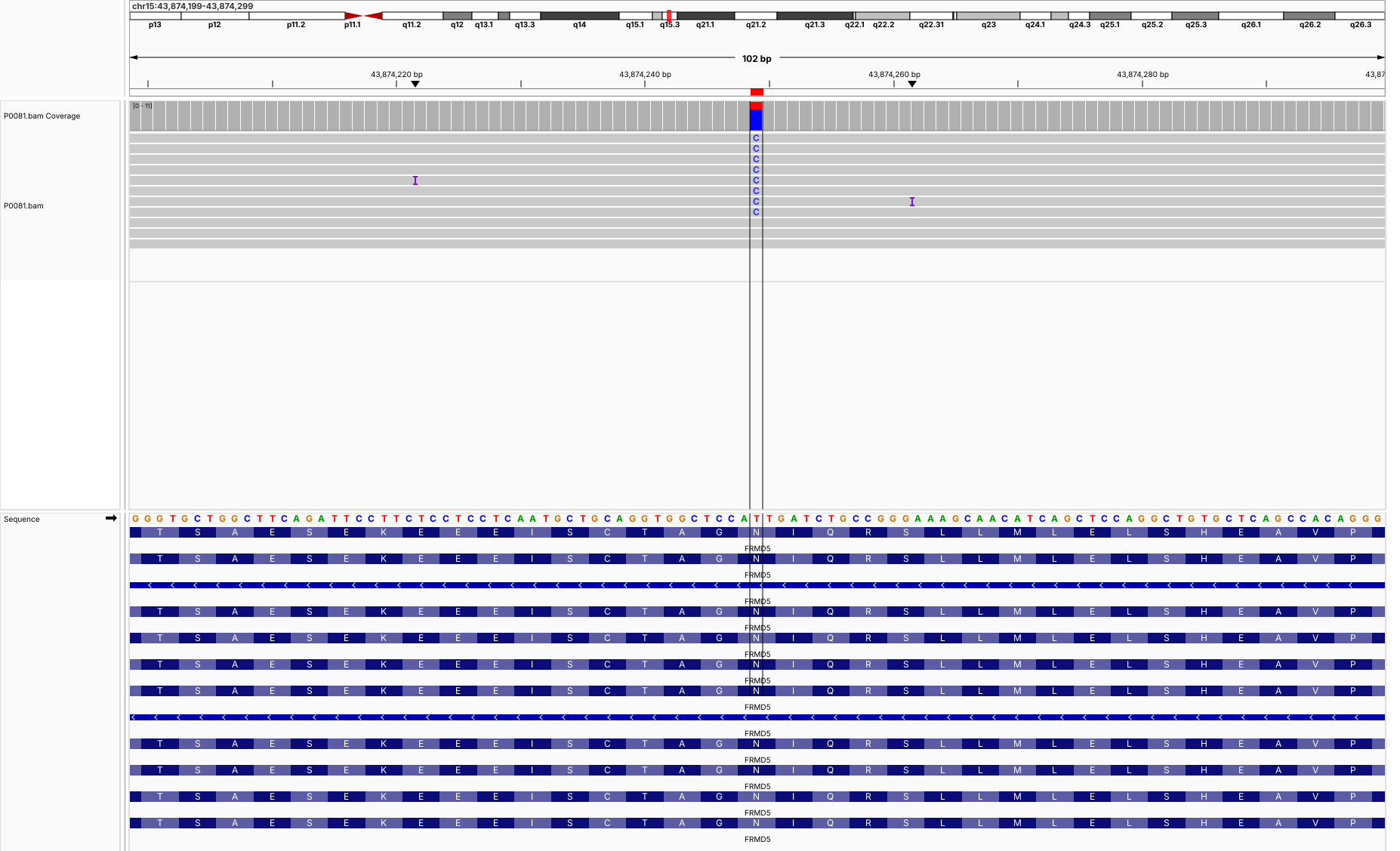

**P0081 (chr18:57,606,636-57,606,736) NARS1**

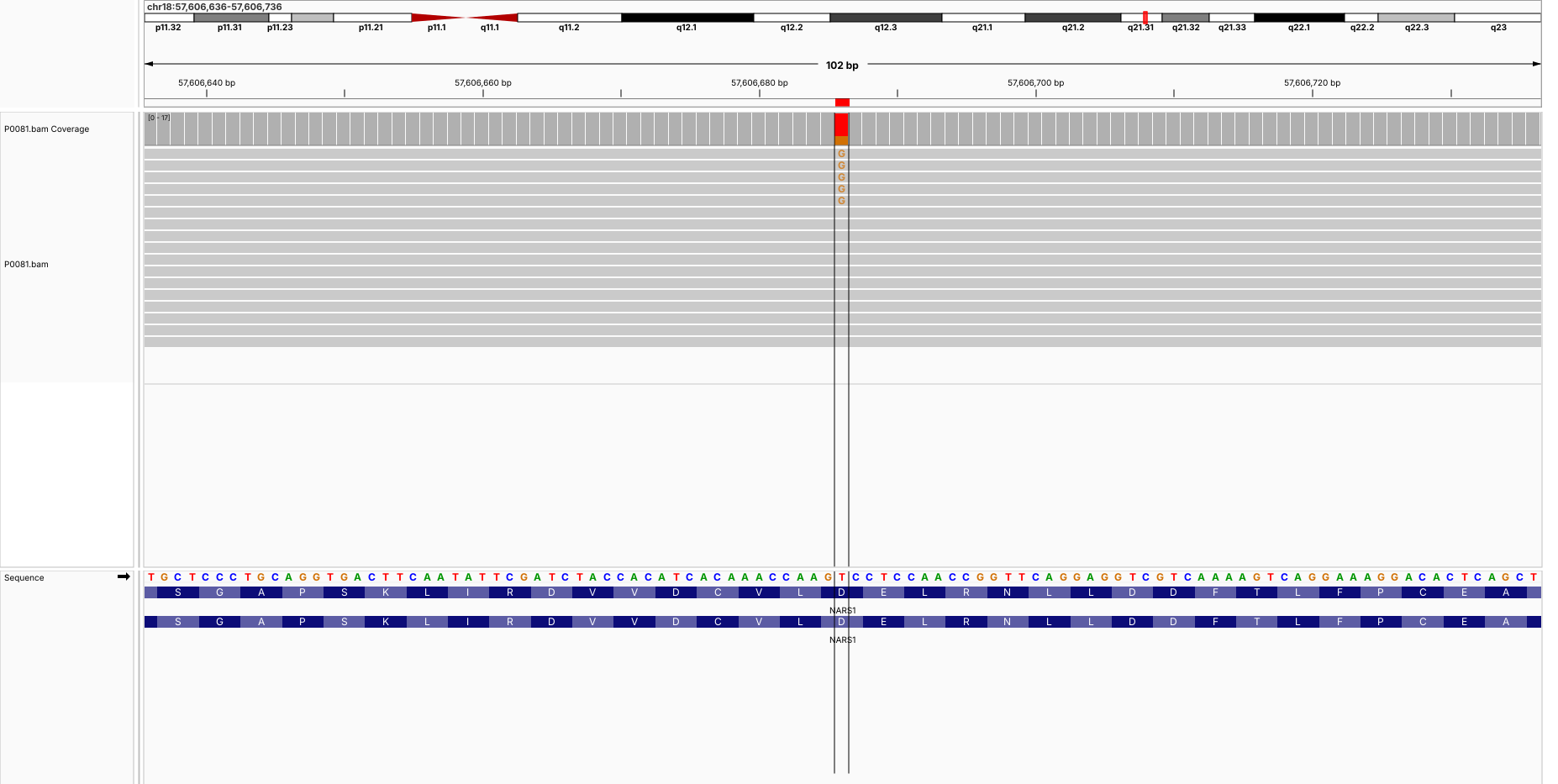

**P0084 (chr1:30,936,631-30,936,731) PUM1**

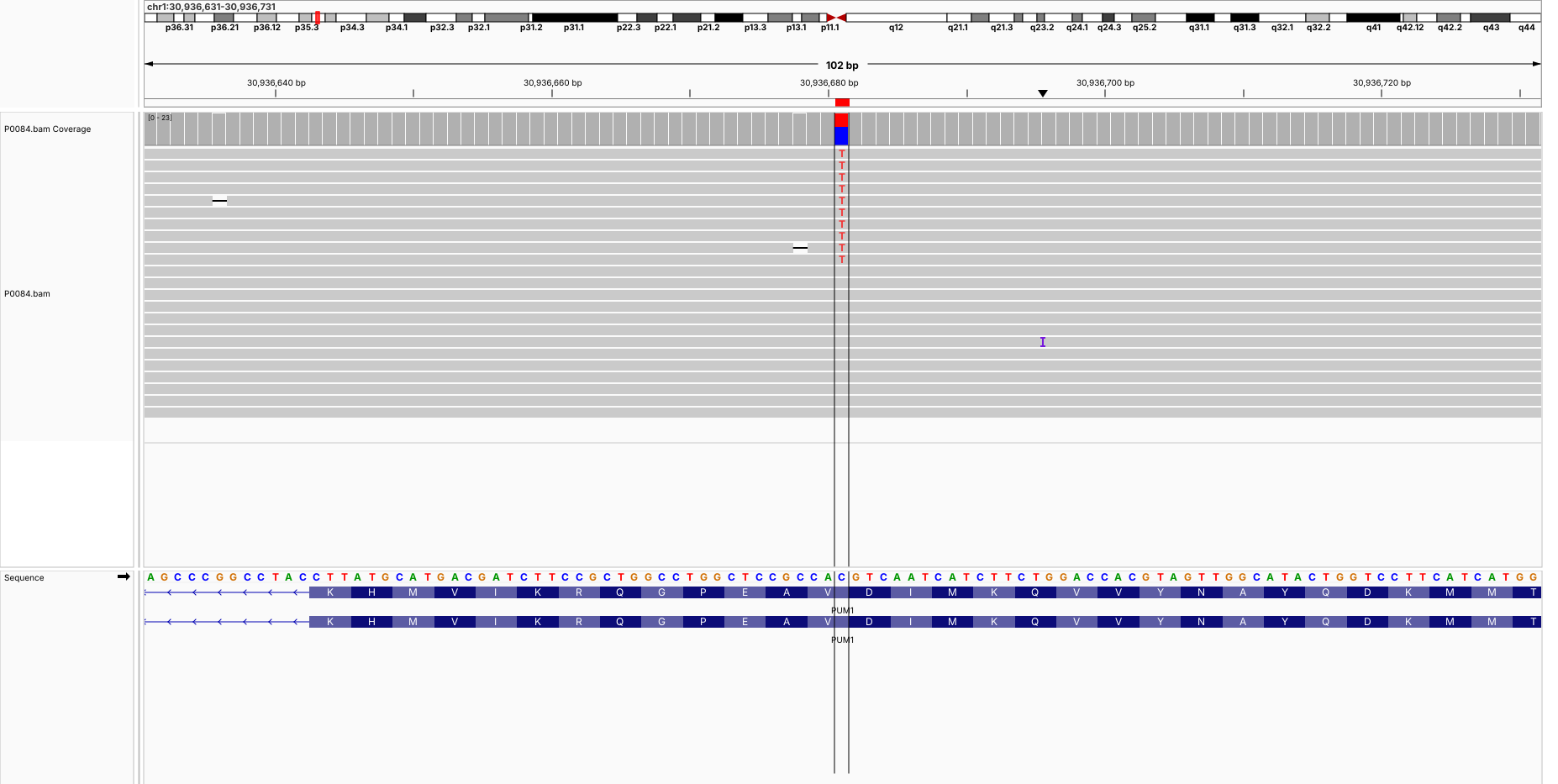

**P0084 (chr7:93,103,578-93,103,678) SAMD9**

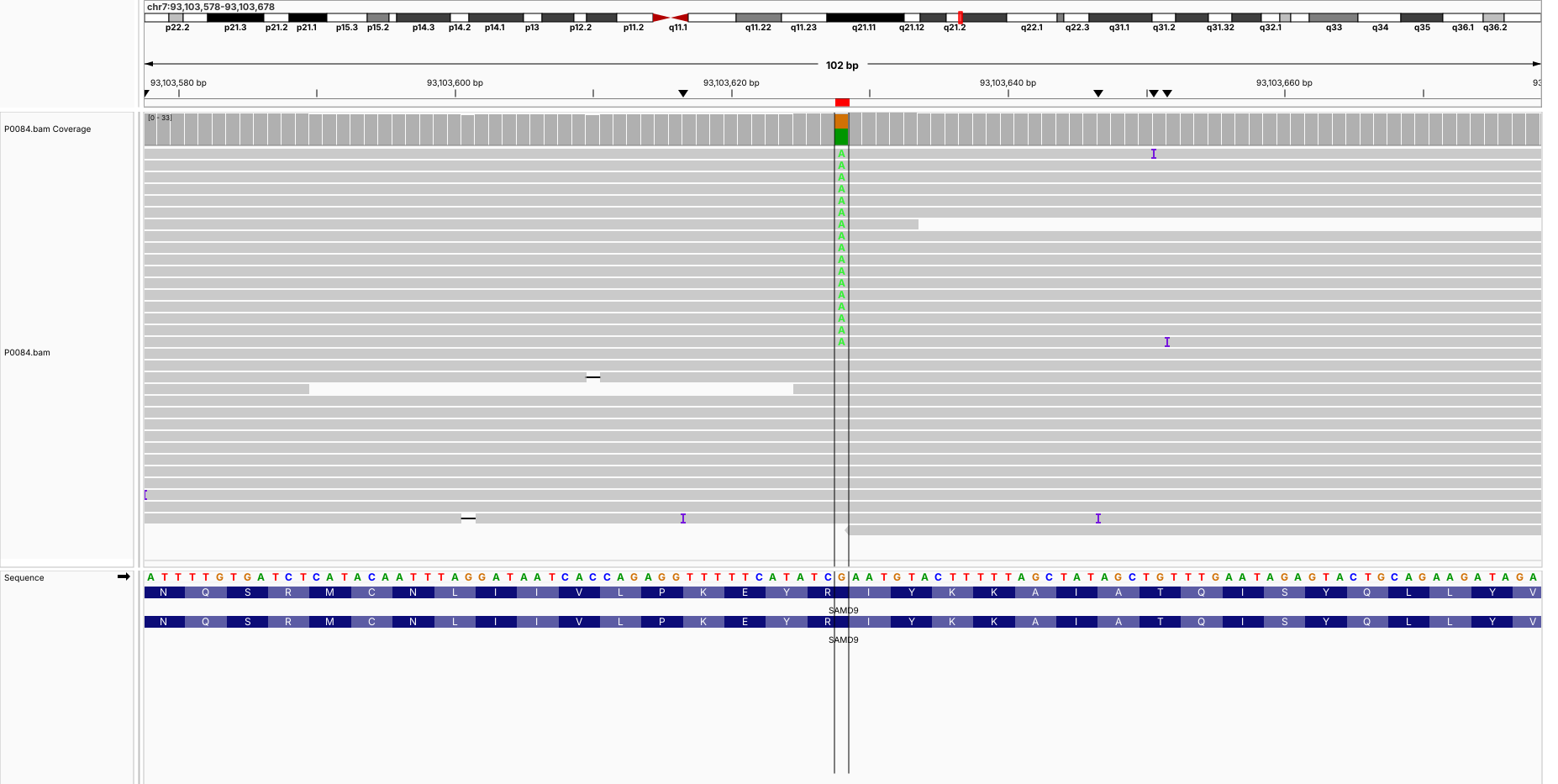

**P0084 (chr7:105,107,170-105,107,270) KMT2E**

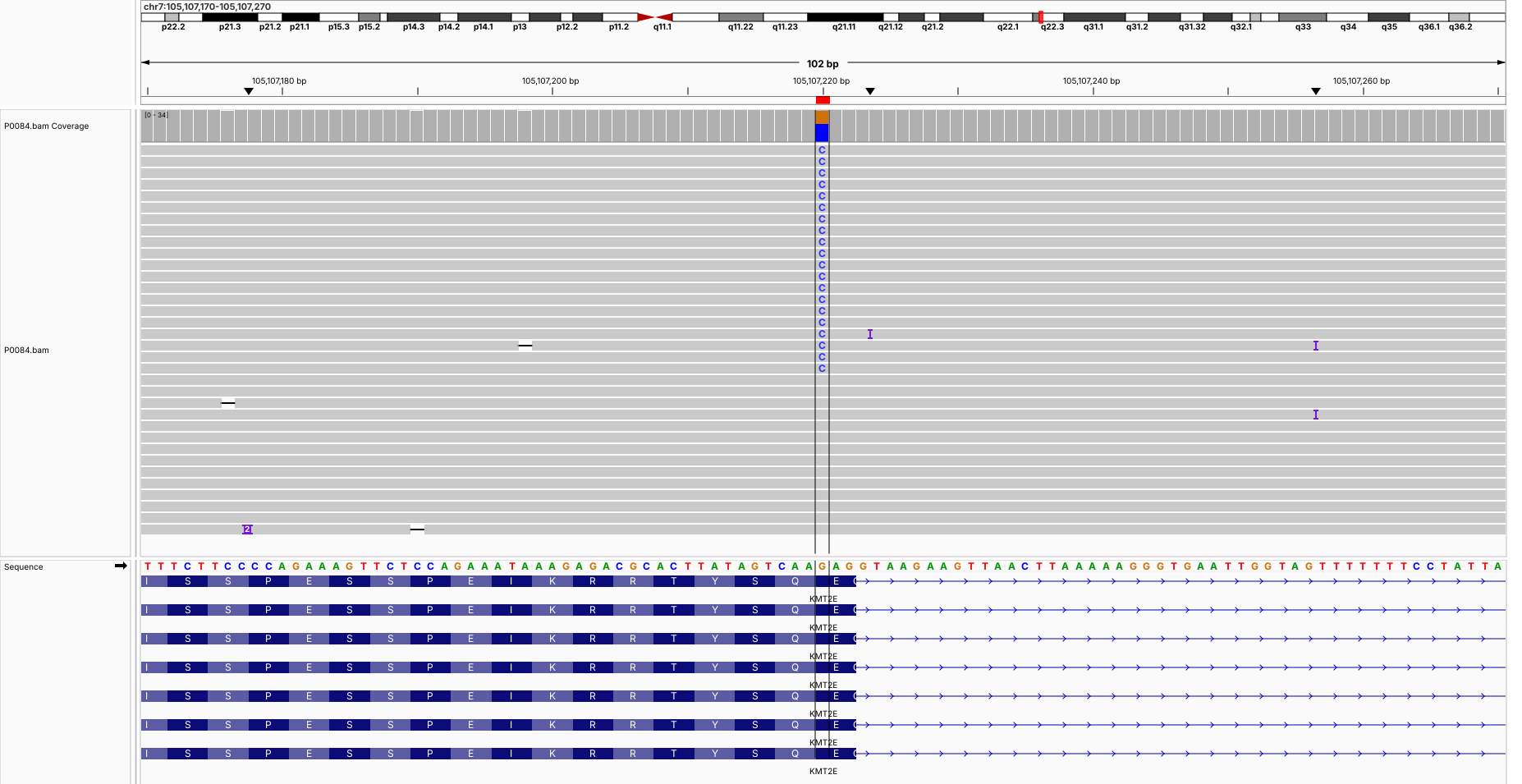

**P0085 (chr3:174,280,472-174,280,572) NLGN1**

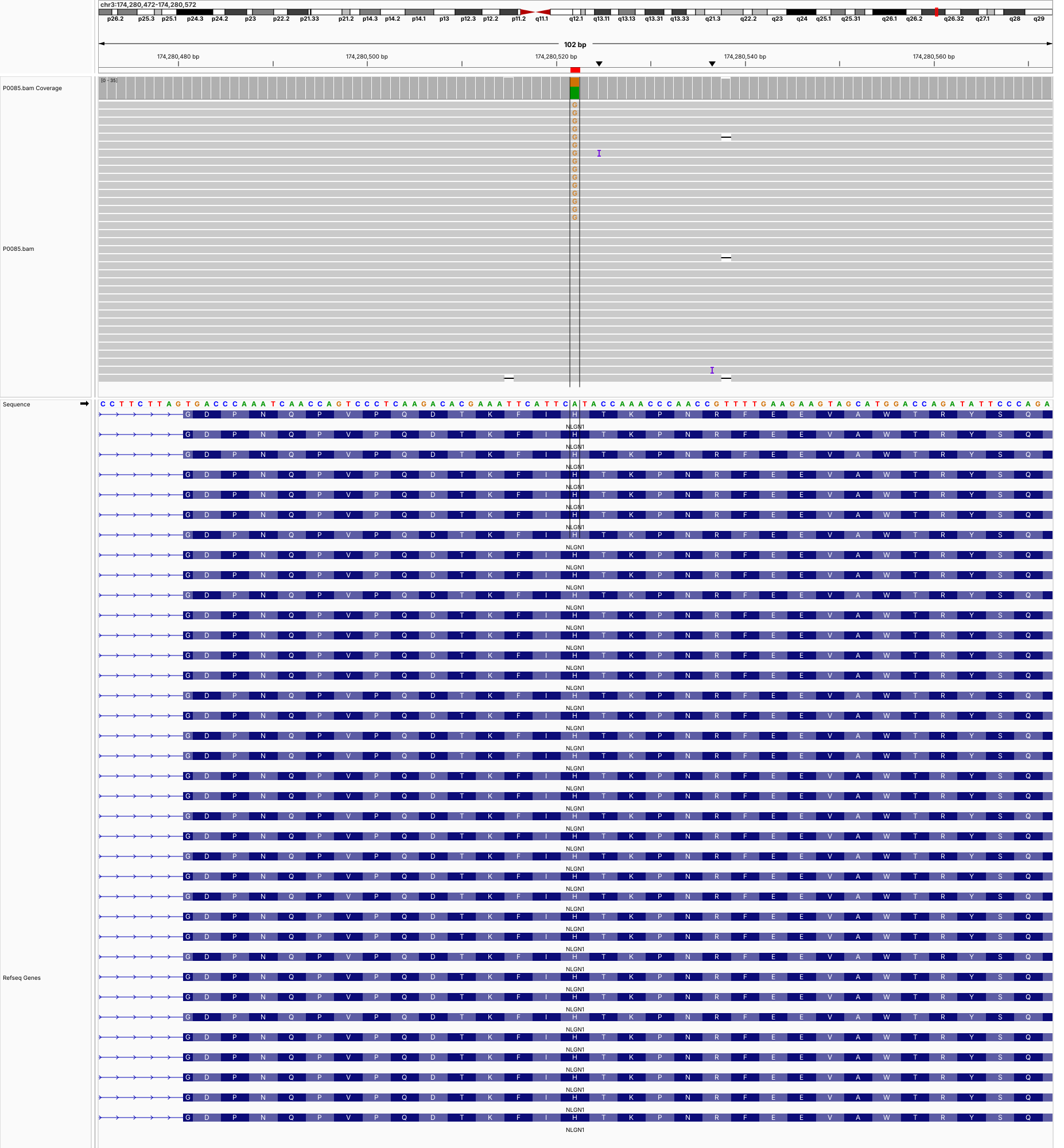

**P0085 (chr5:14,280,317-14,280,417) TRIO**

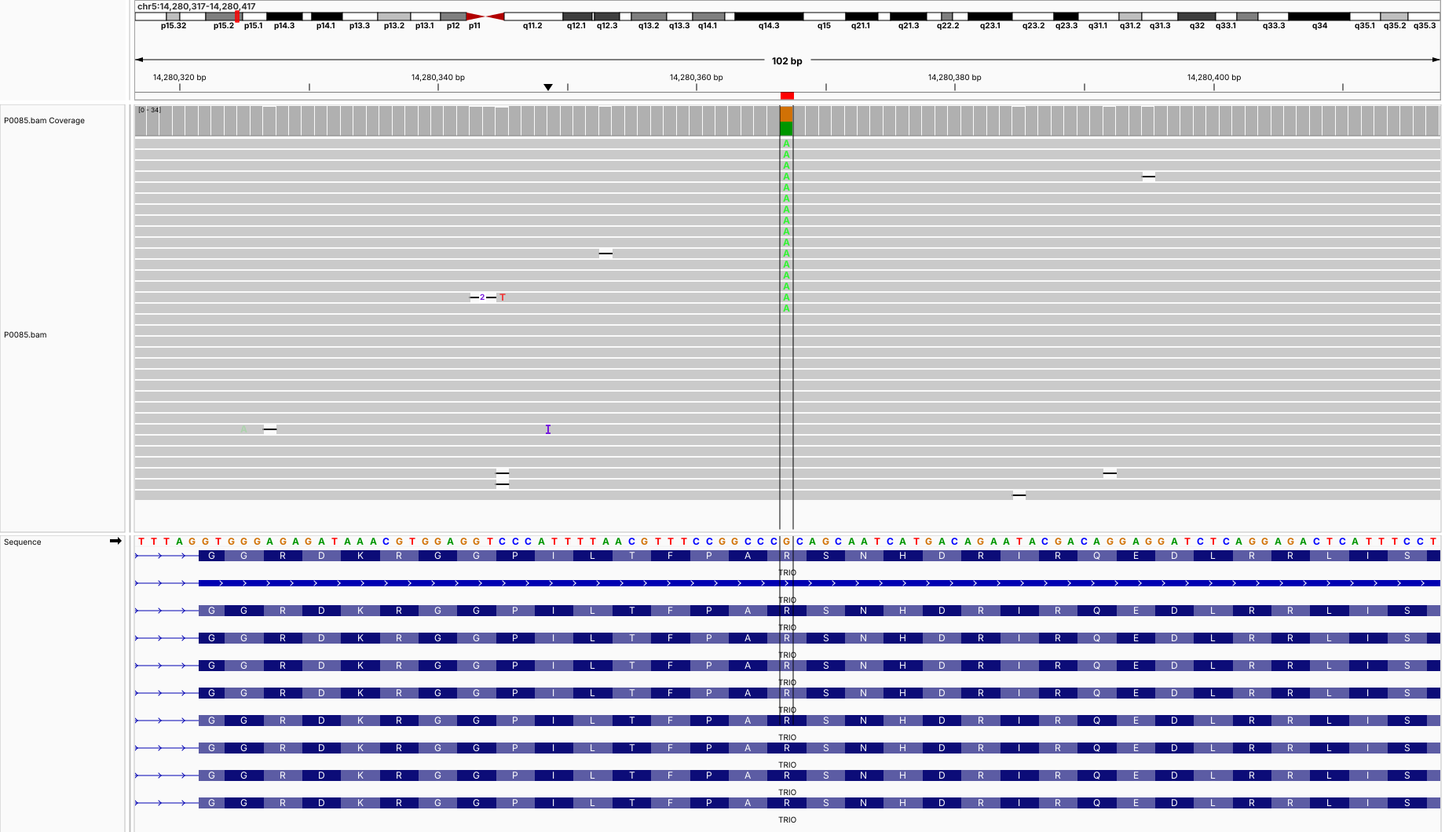

**P0086 (chr12:49,034,118-49,034,218) KMT2D**

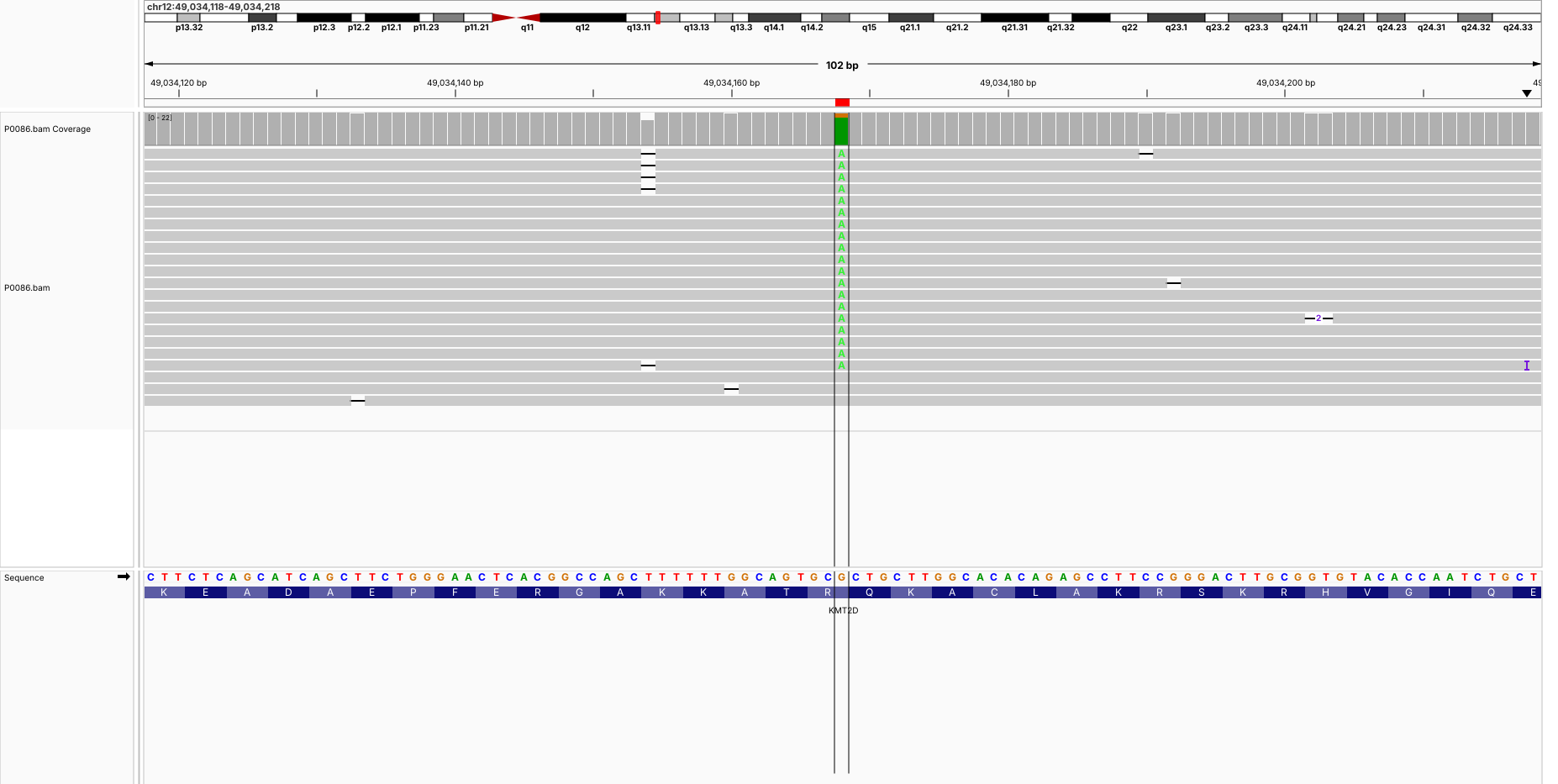

**P0087 (chr3:174,280,472-174,280,572) NLGN1**

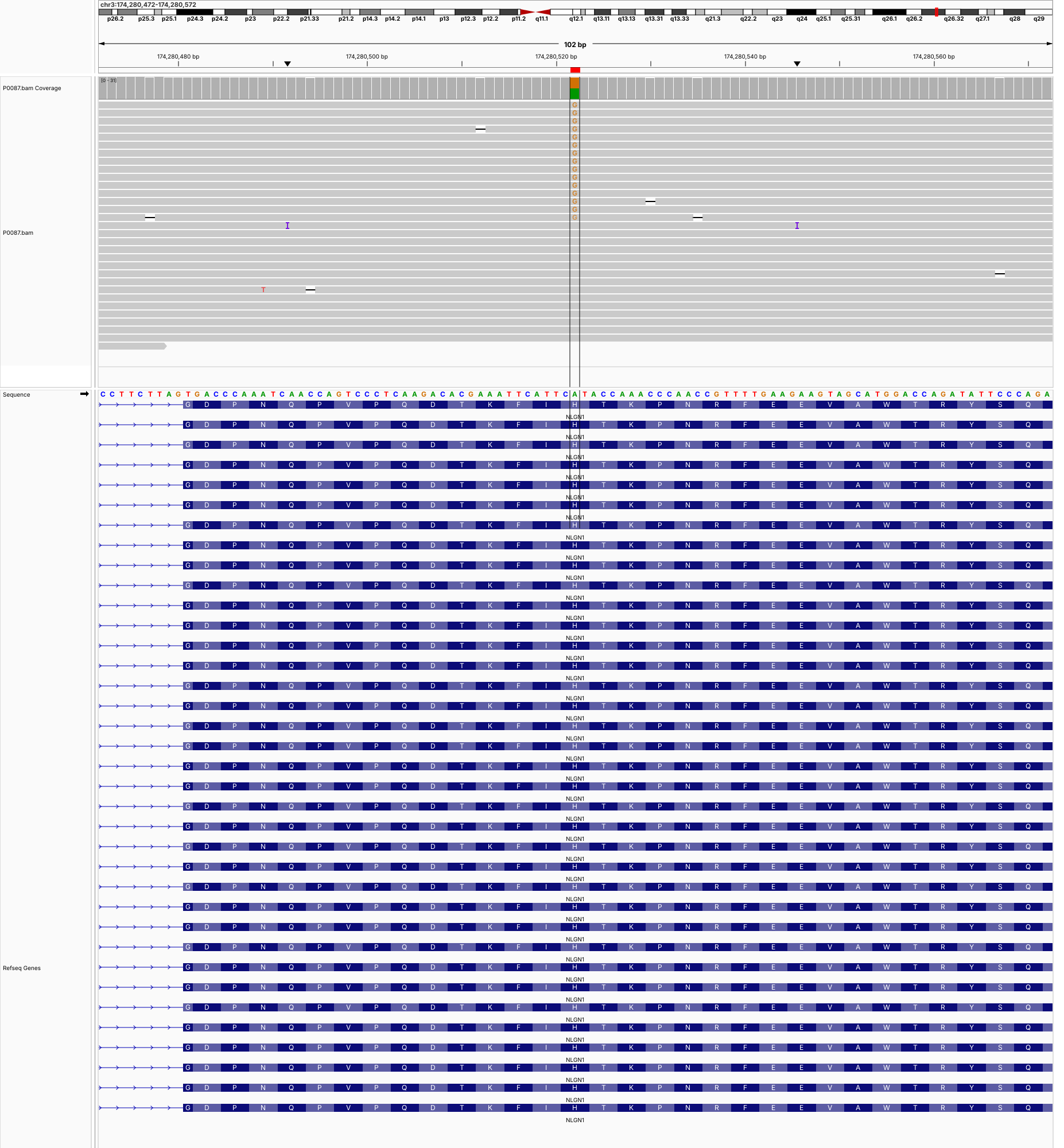

**P0087 (chr19:42,272,014-42,272,114) CIC**

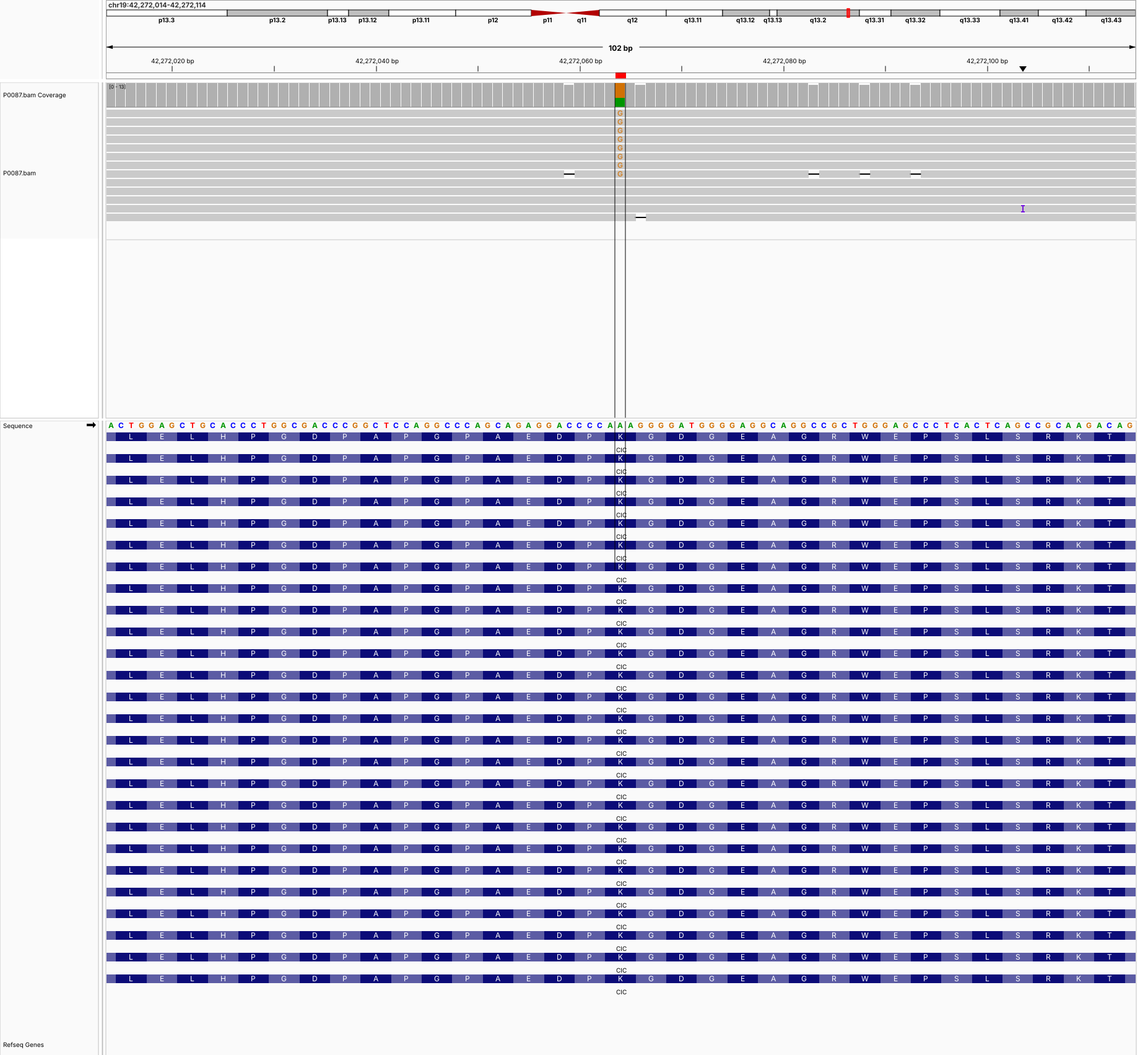

**P0088 (chr4:139,378,804-139,378,904) NAA15**

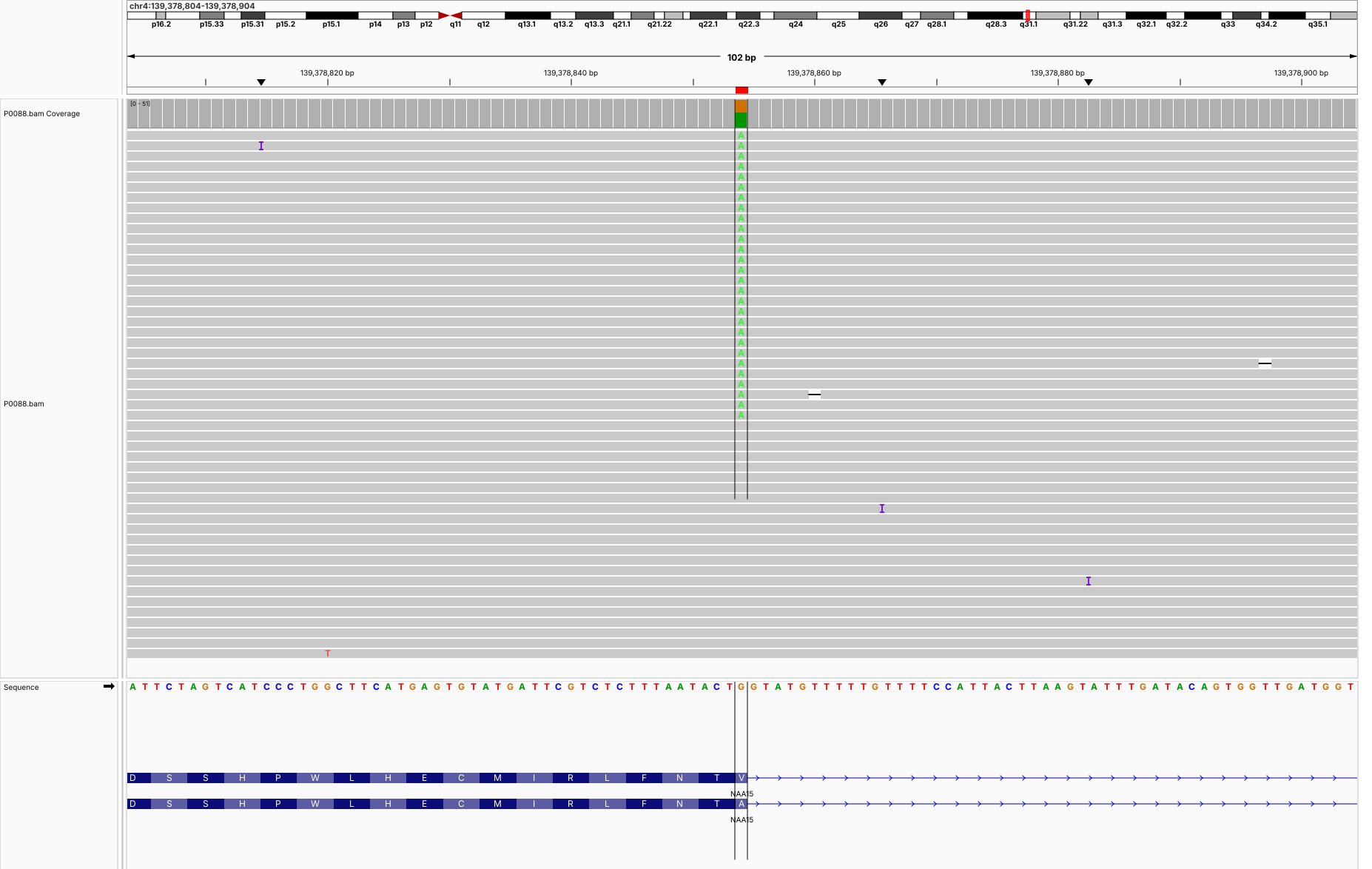

**P0090 (chr2:47,808,233-47,808,333) FBXO11**

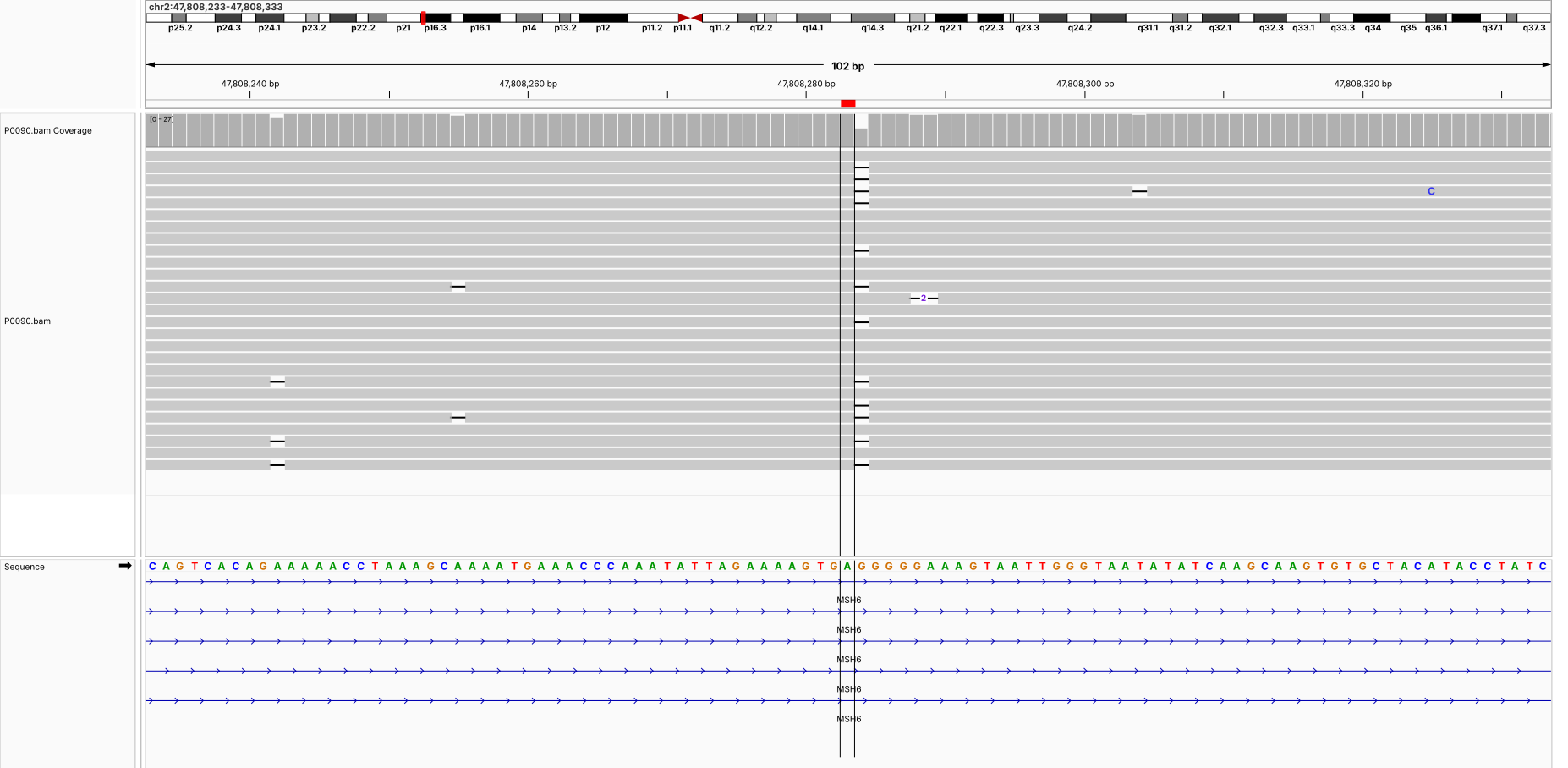

**P0090 (chr19:42,287,278-42,287,378) CIC**

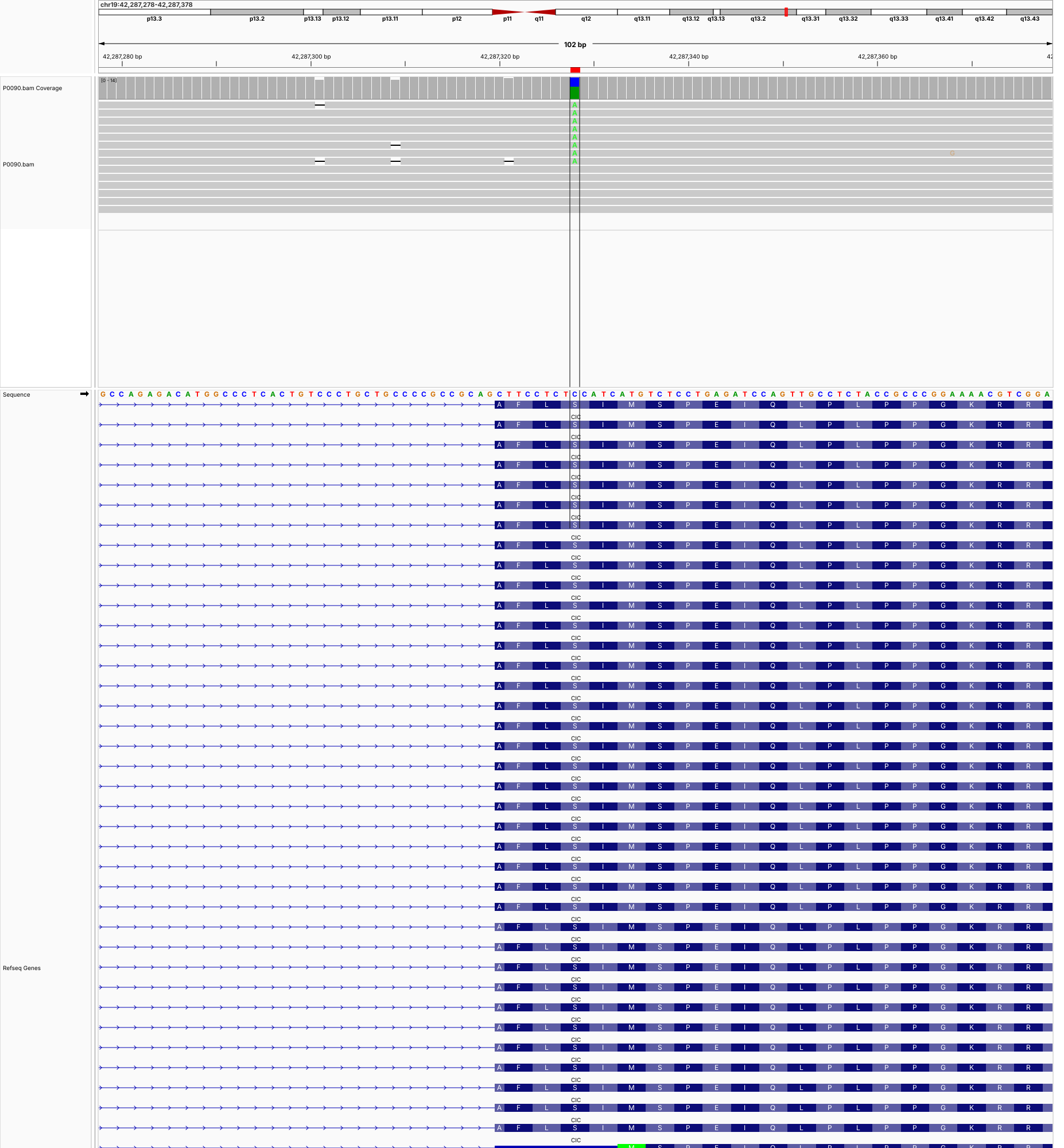

**P0091 (chr1:27,550,307-27,550,407) AHDC1**

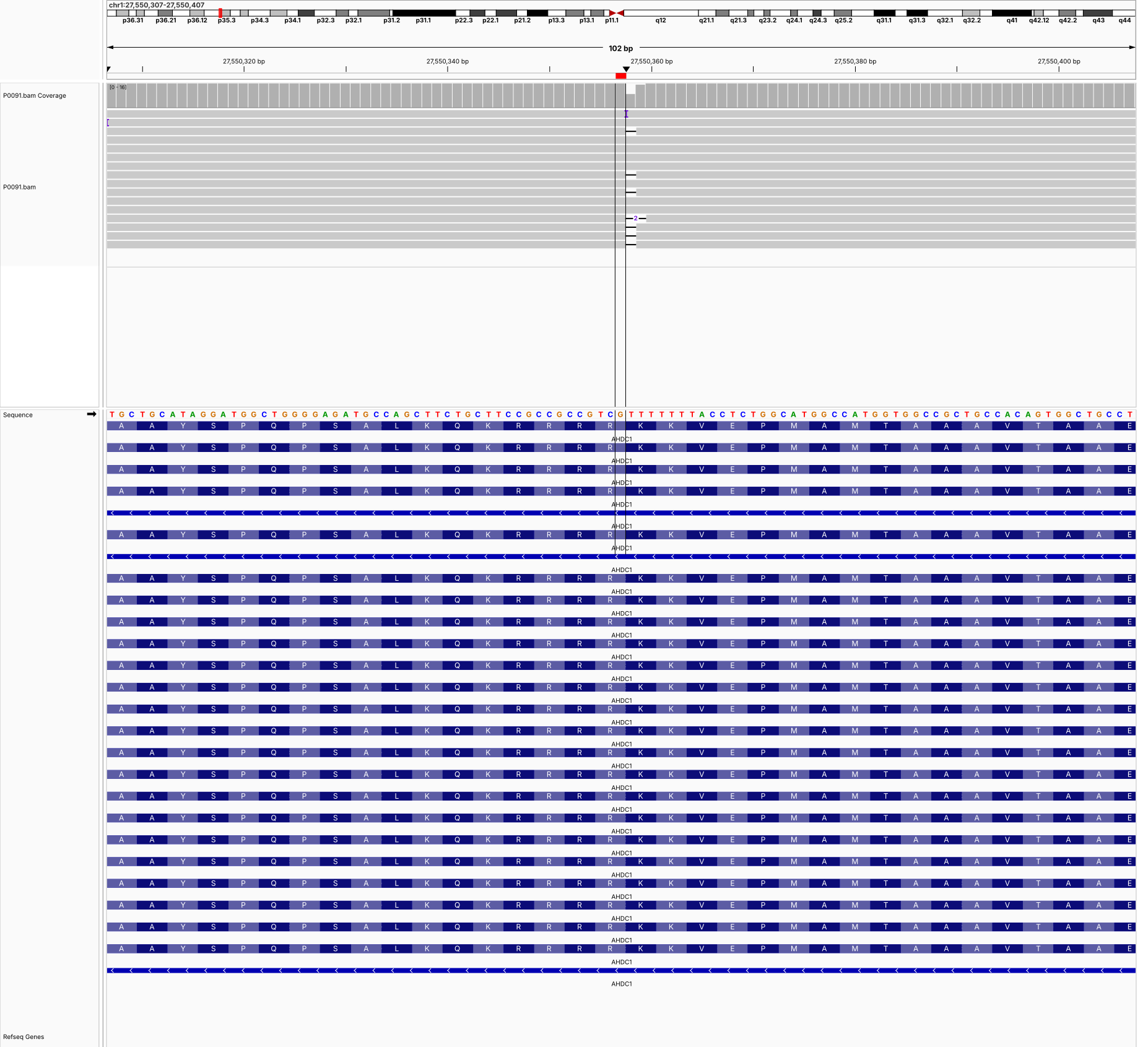

**P0091 (chr4:73,124,872-73,124,972) ANKRD17**

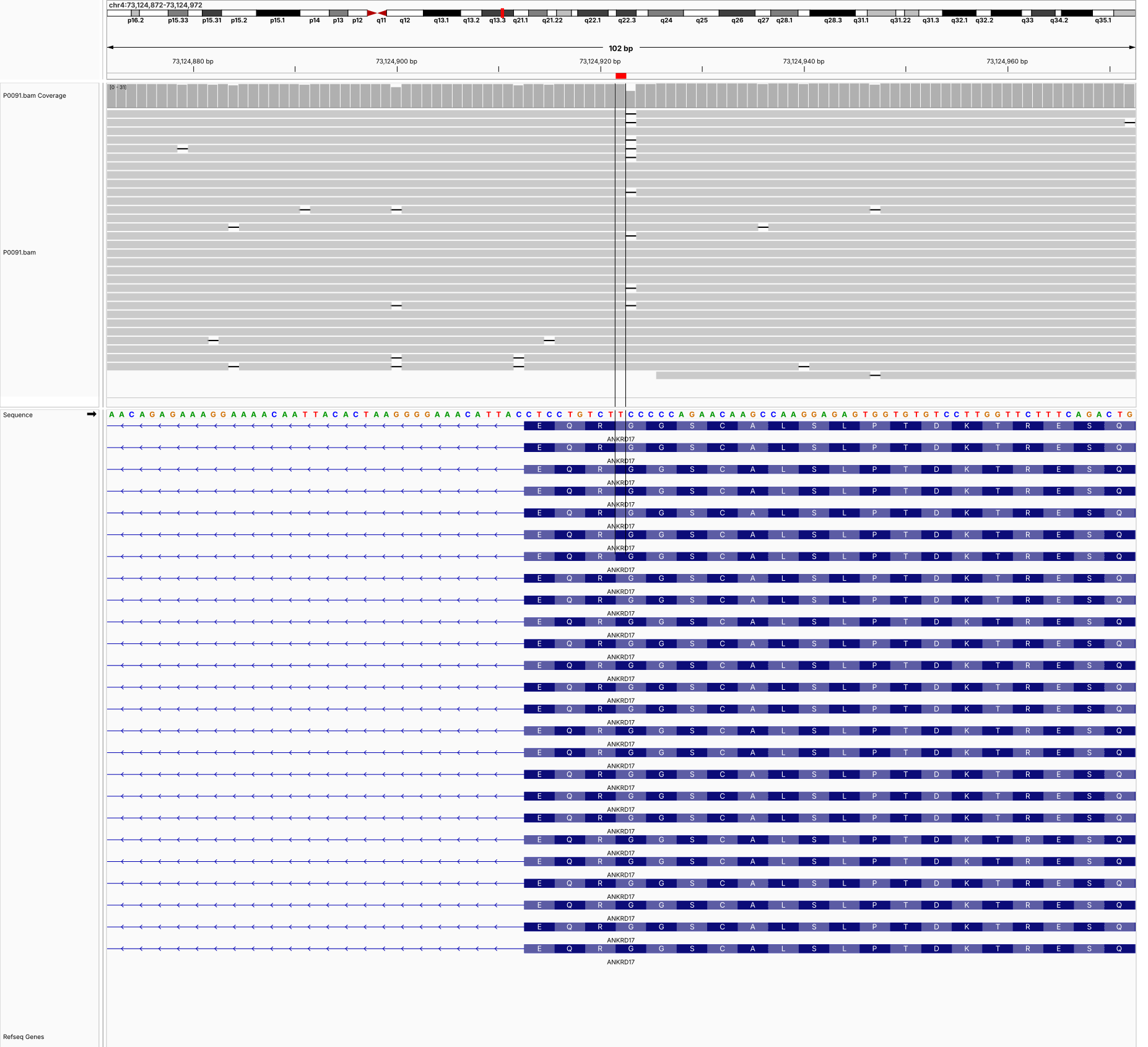

**P0091 (chrX:53,546,466-53,546,566) HUWE1**

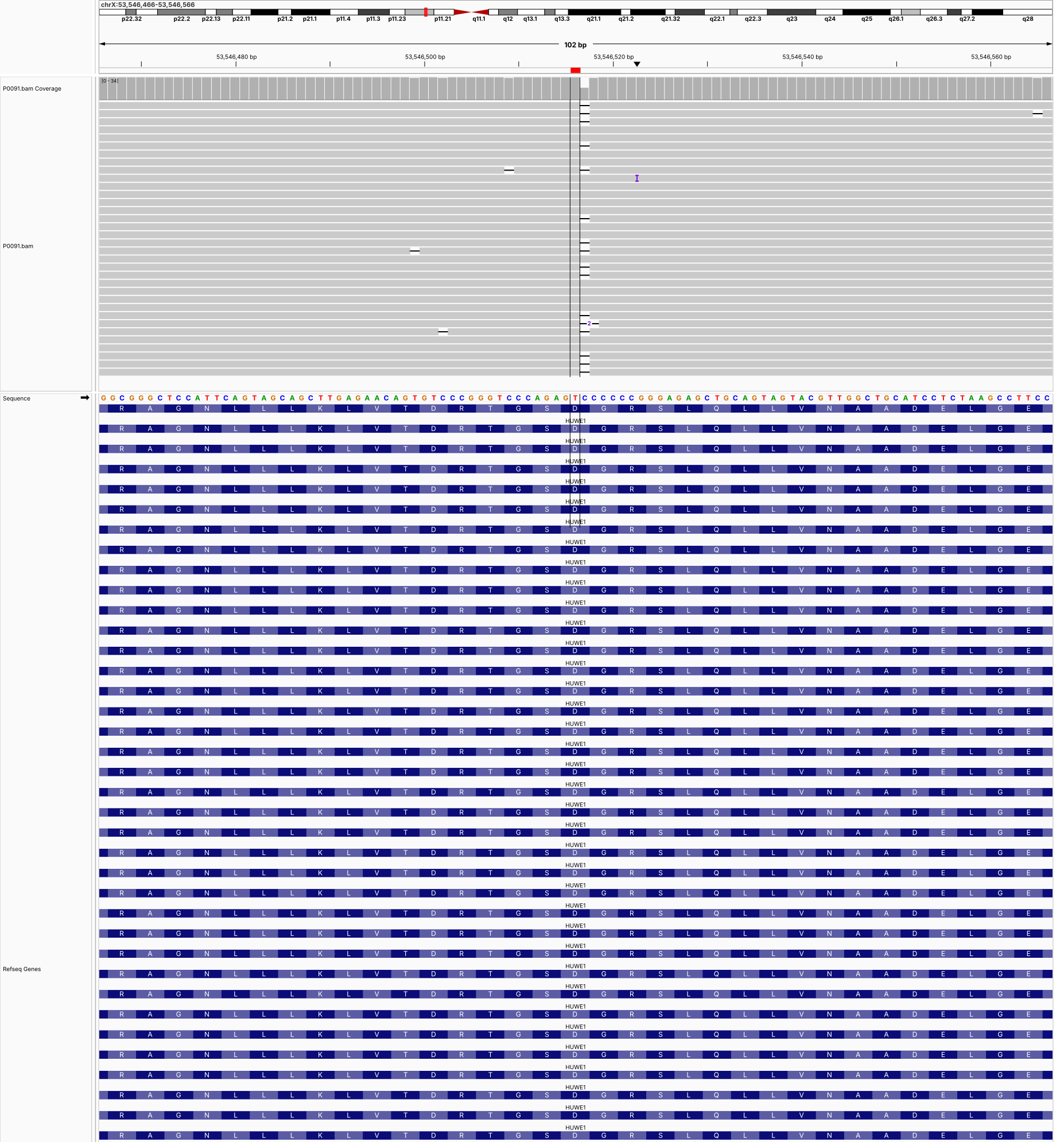

**P0092 (chr2:148,469,275-148,469,375) MBD5**

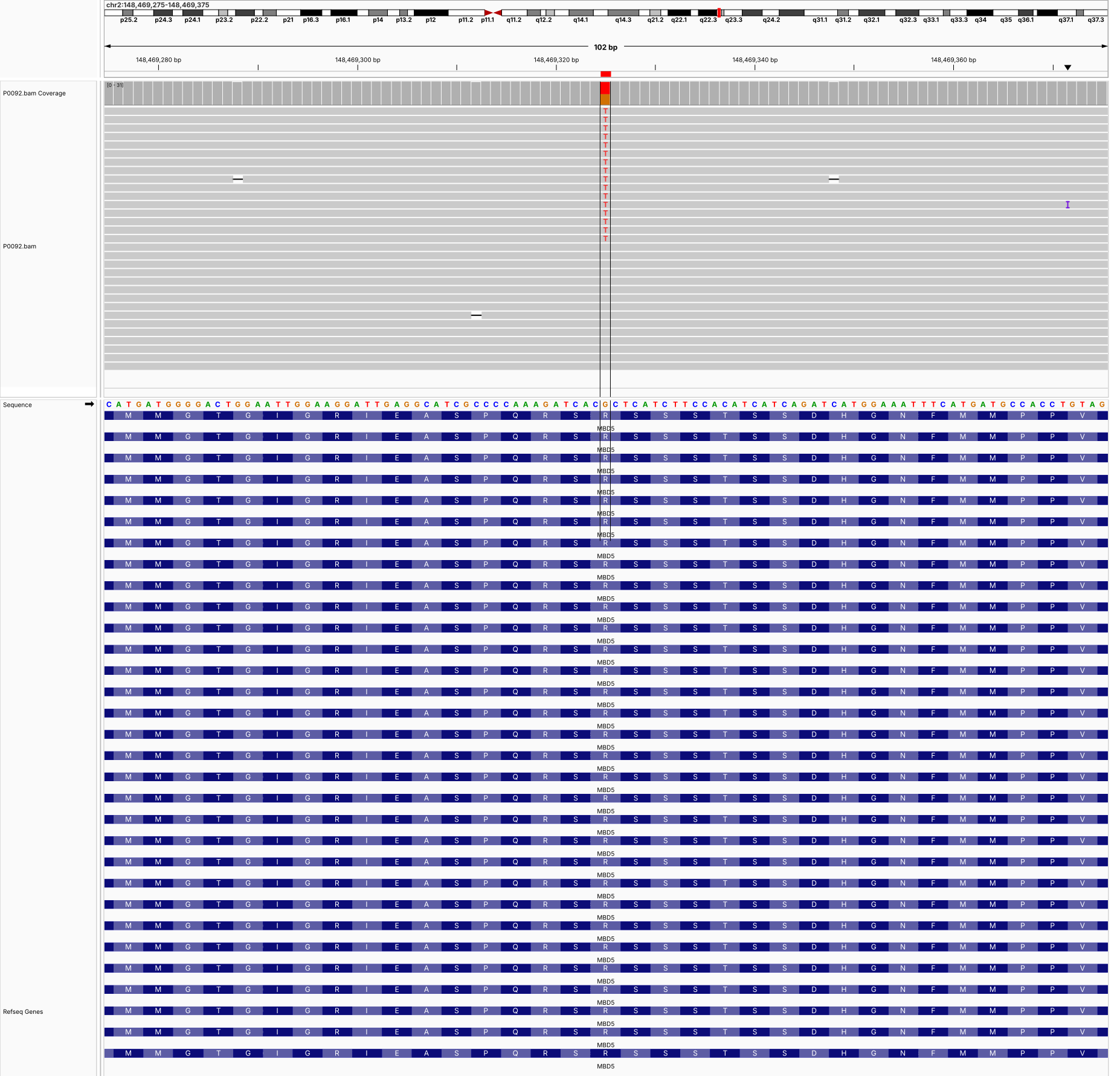

**P0092 (chr3:38,908,973-38,909,073) SCN11A**

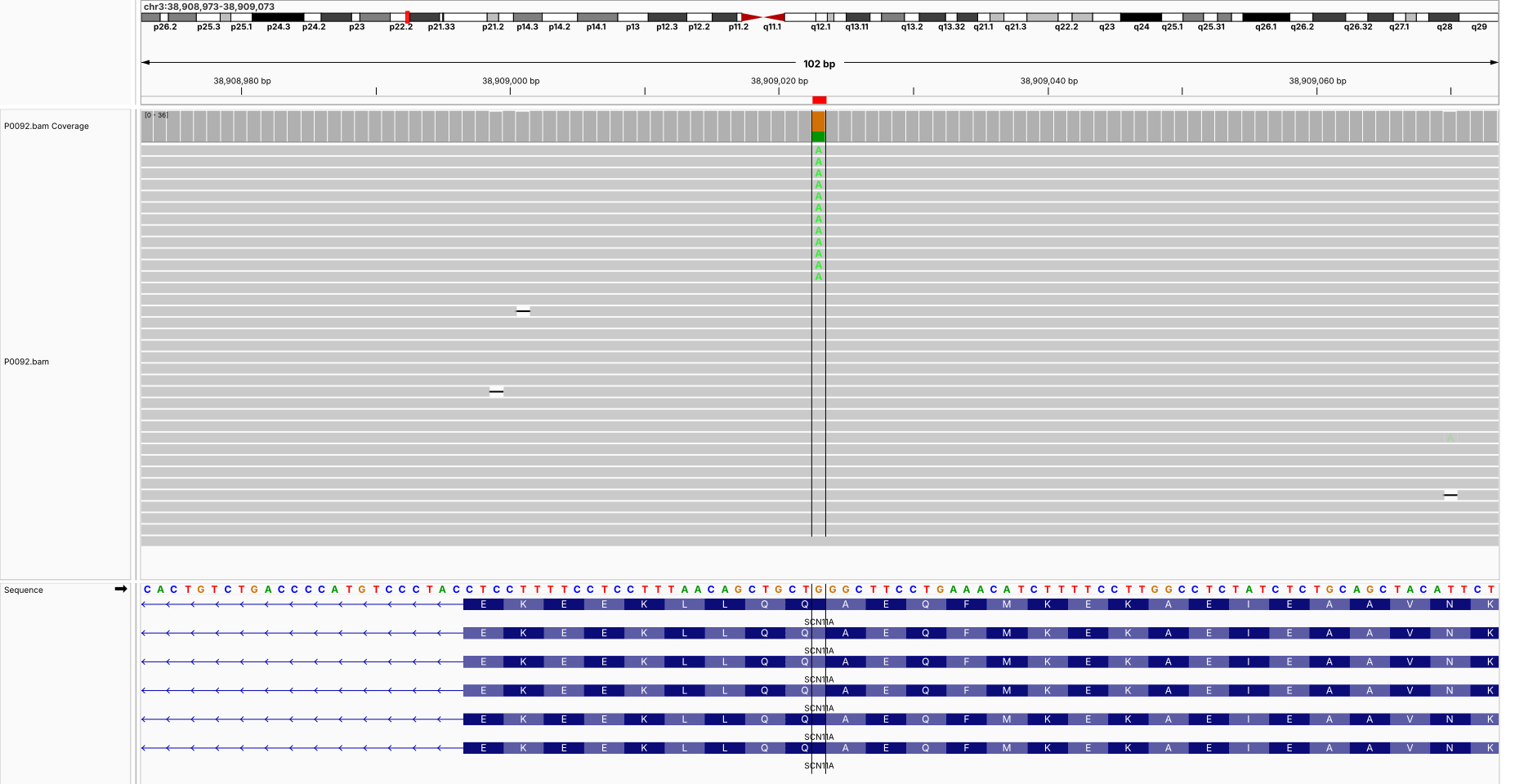

**P0092 (chr9:135,765,101-135,765,201) KCNT1**

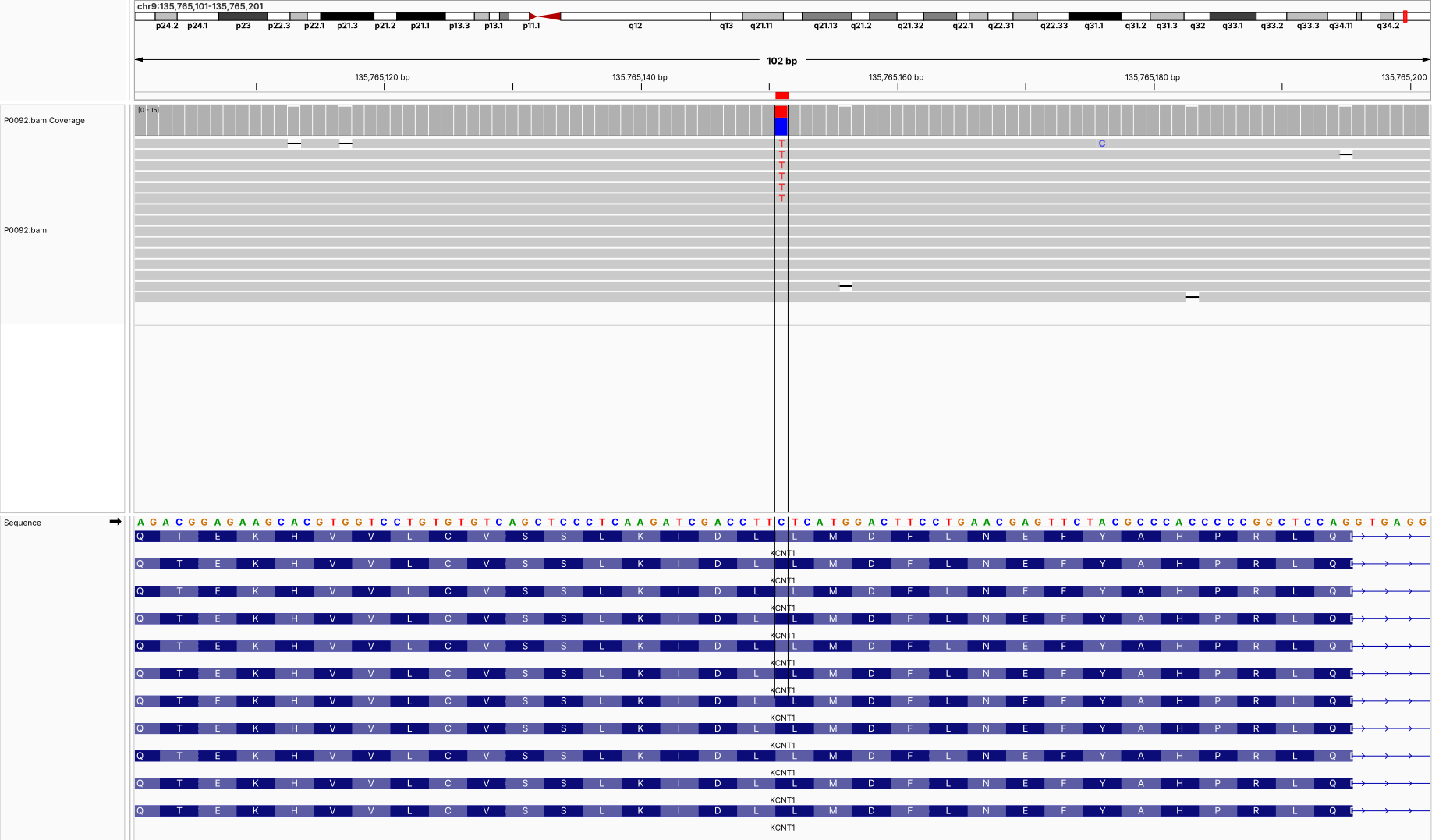

**P0092 (chrX:145,823,915-145,824,015) SLITRK2**

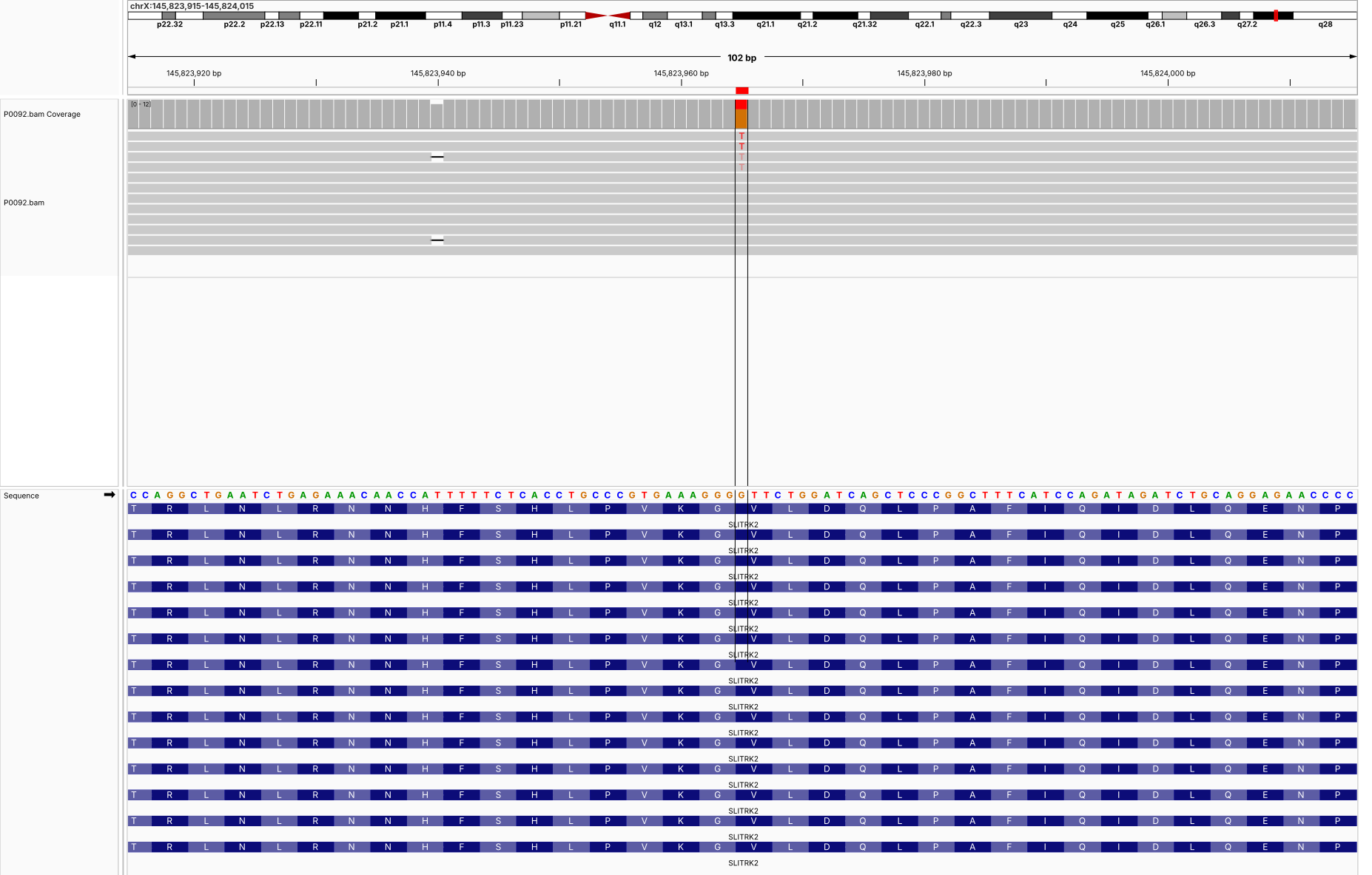

**P0094 (chr11:637,358-637,458) DRD4**

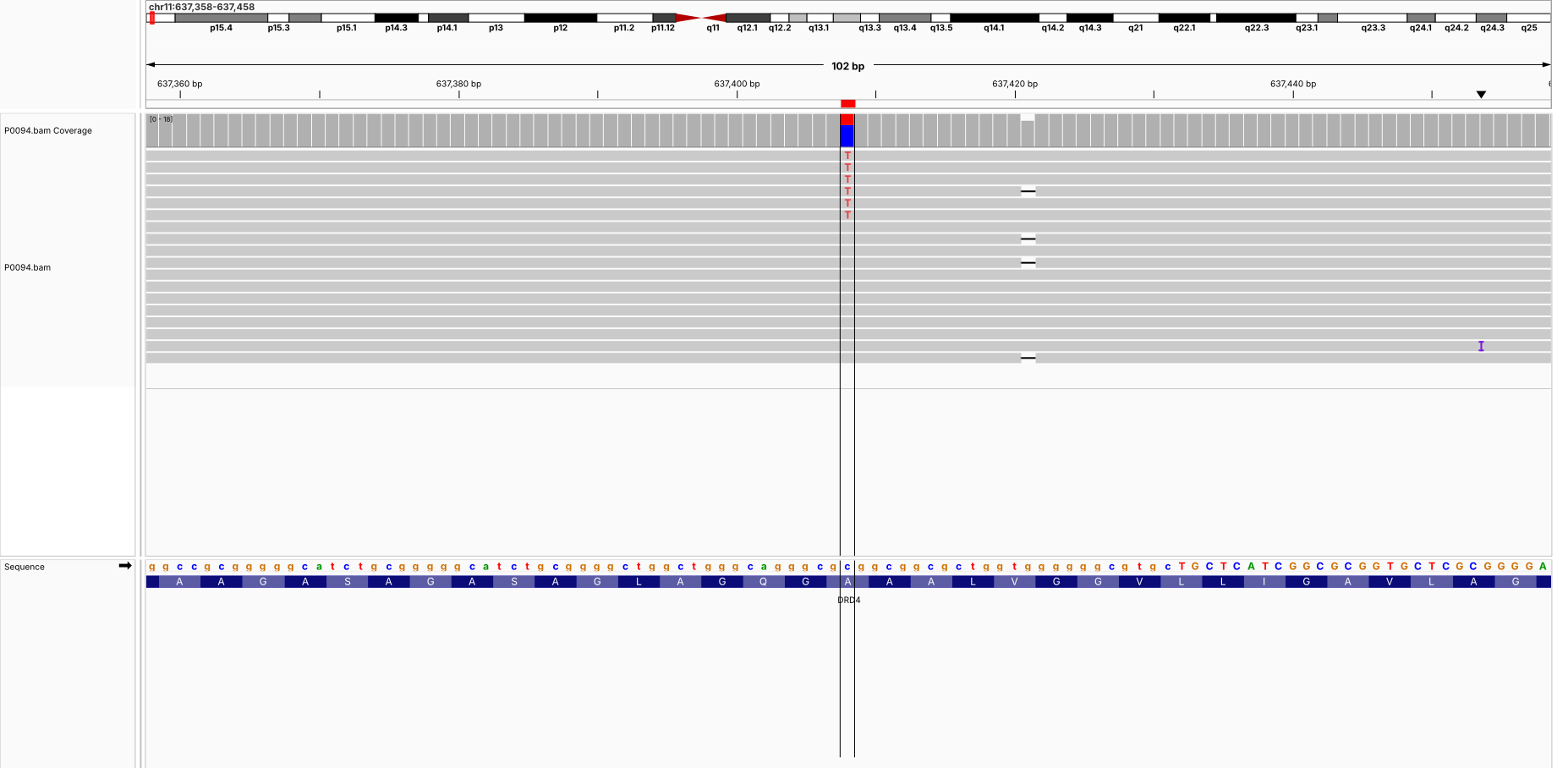

**P0094 (chr17:7,908,397-7,908,497) CHD3**

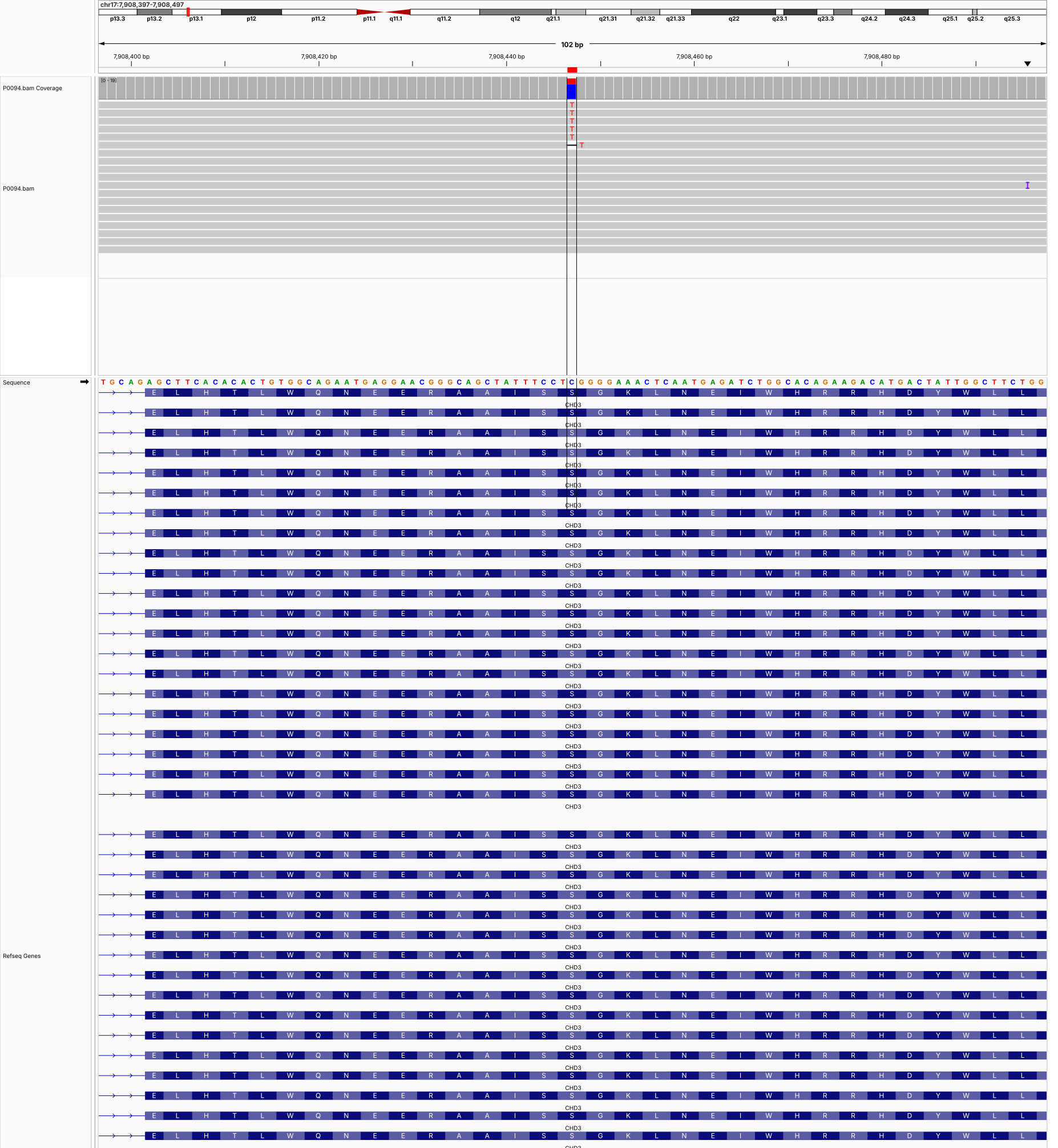

**P0095 (chr19:5,131,488-5,131,588) KDM4B**

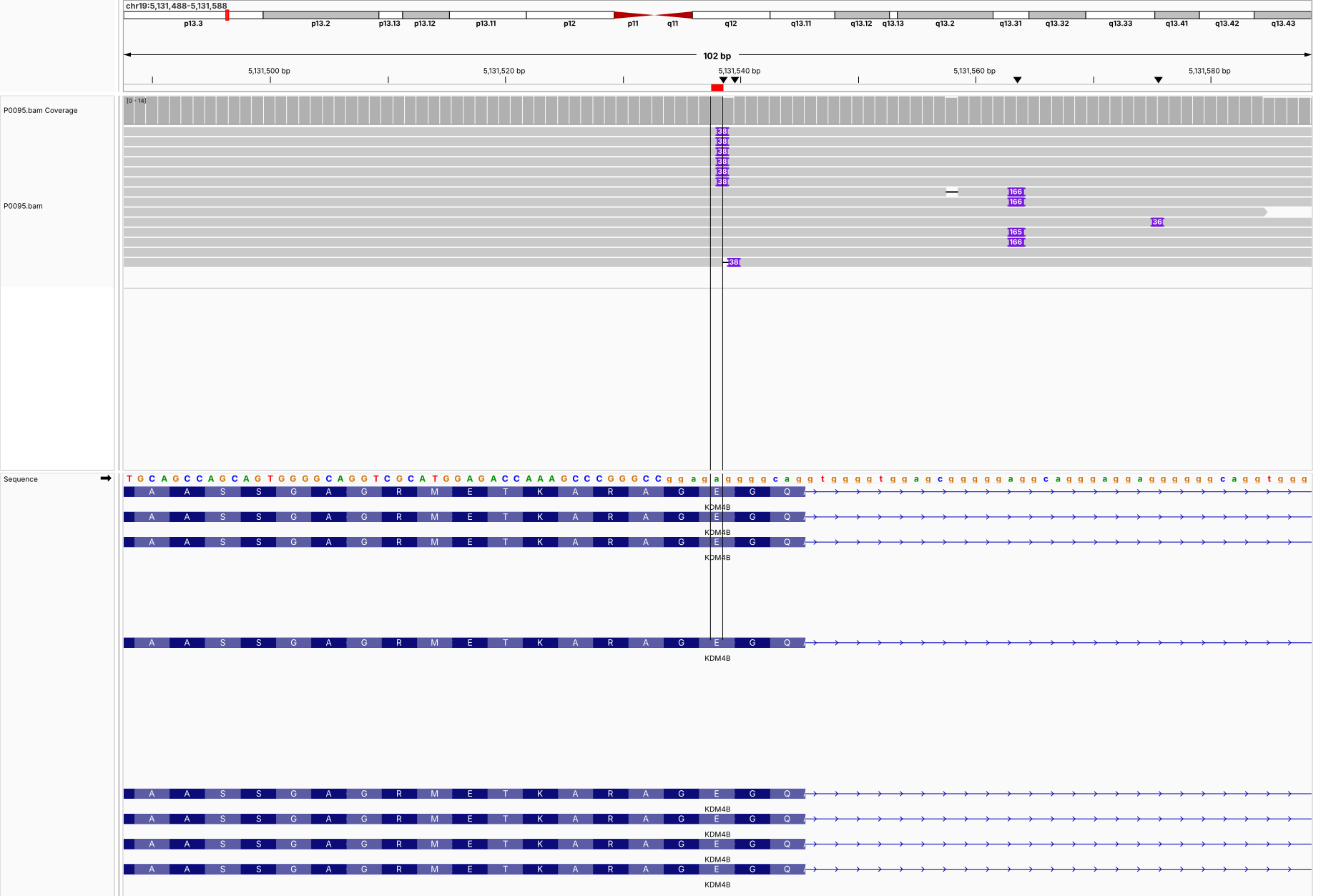

**P0097 (chr19:14,161,391-14,161,491) ADGRL1**

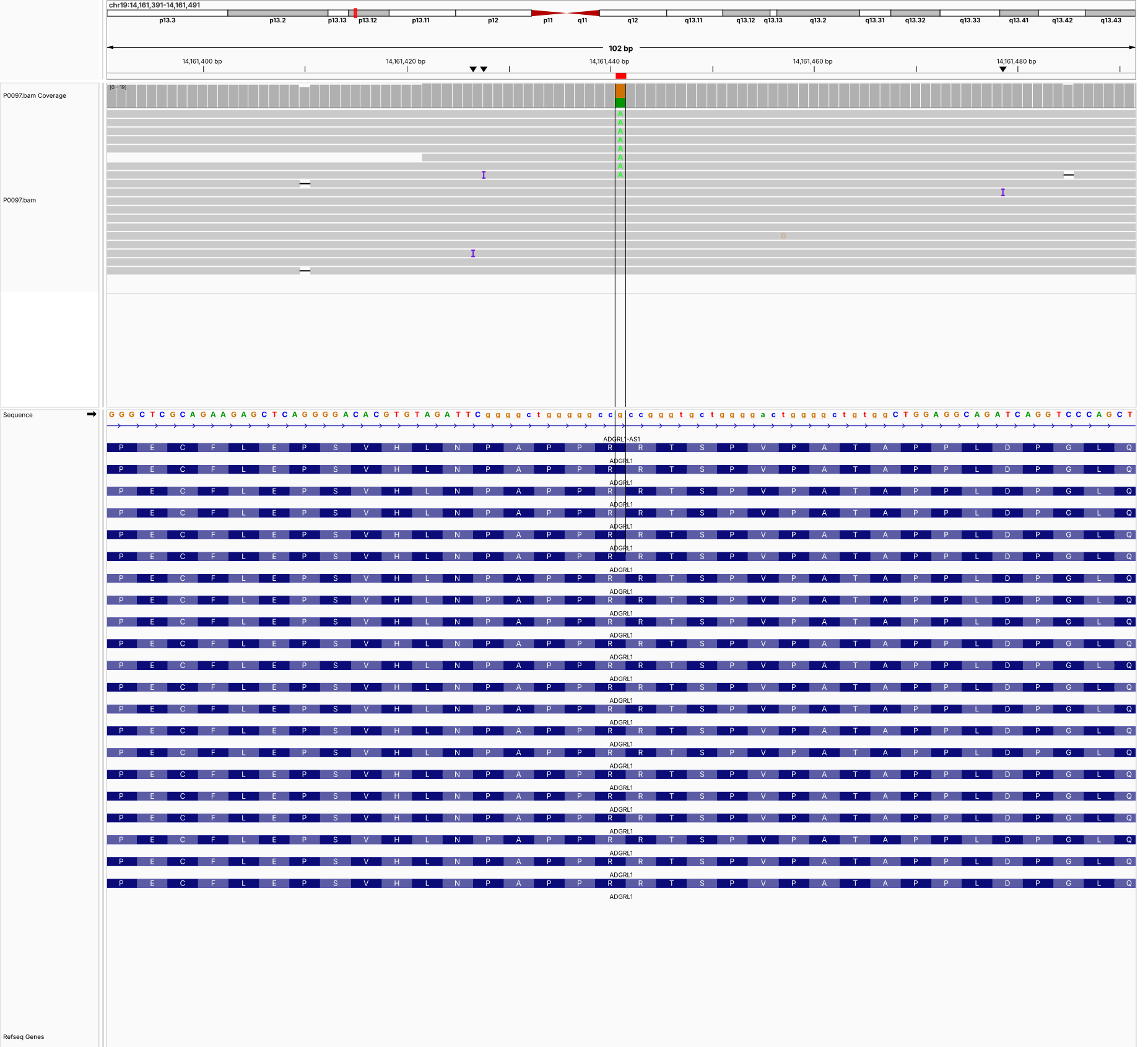

**P0102 (chr9:84,702,182-84,702,282) NTRK2**

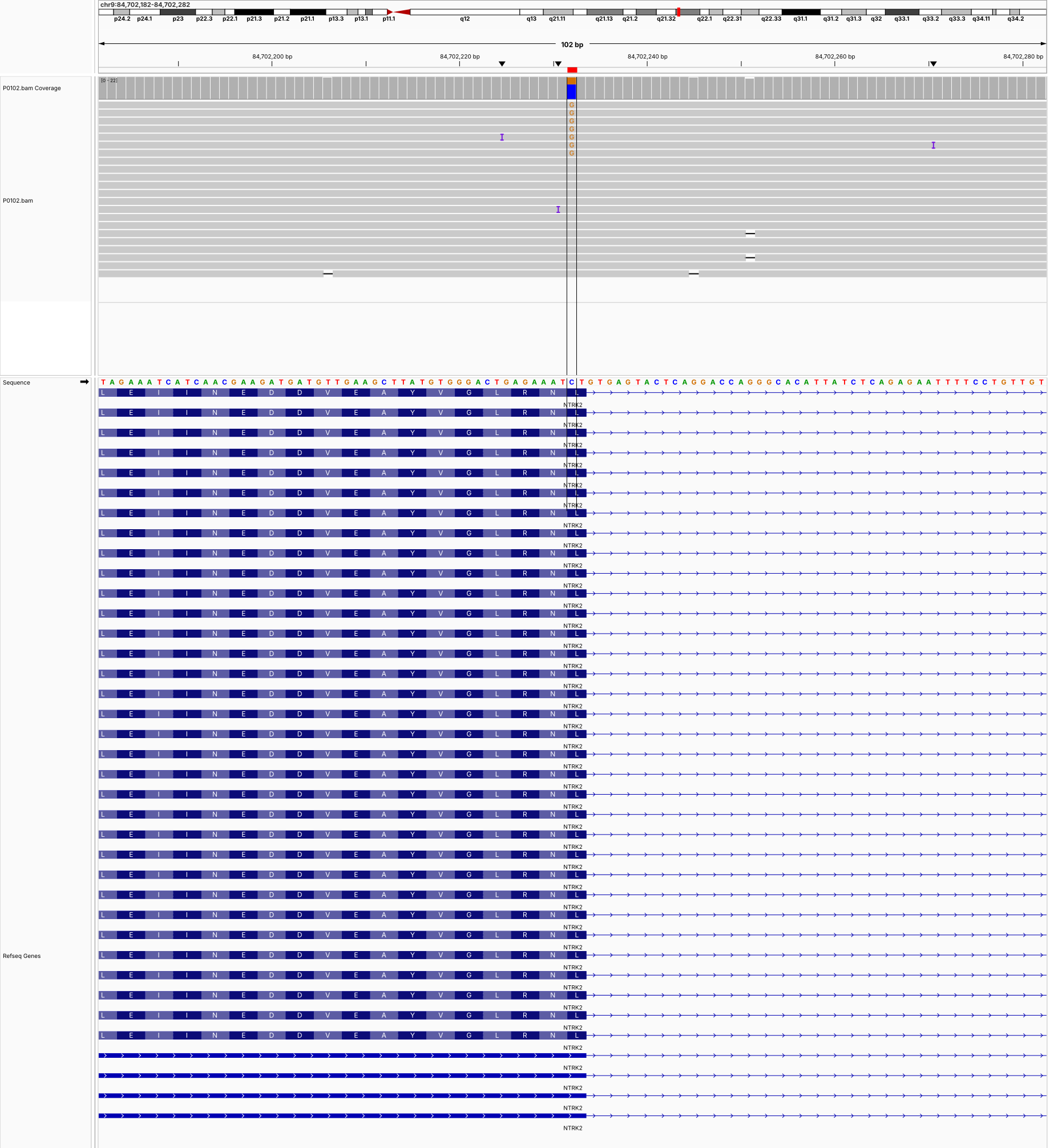

**P0102 (chr16:2,765,916-2,766,016) SRRM2**

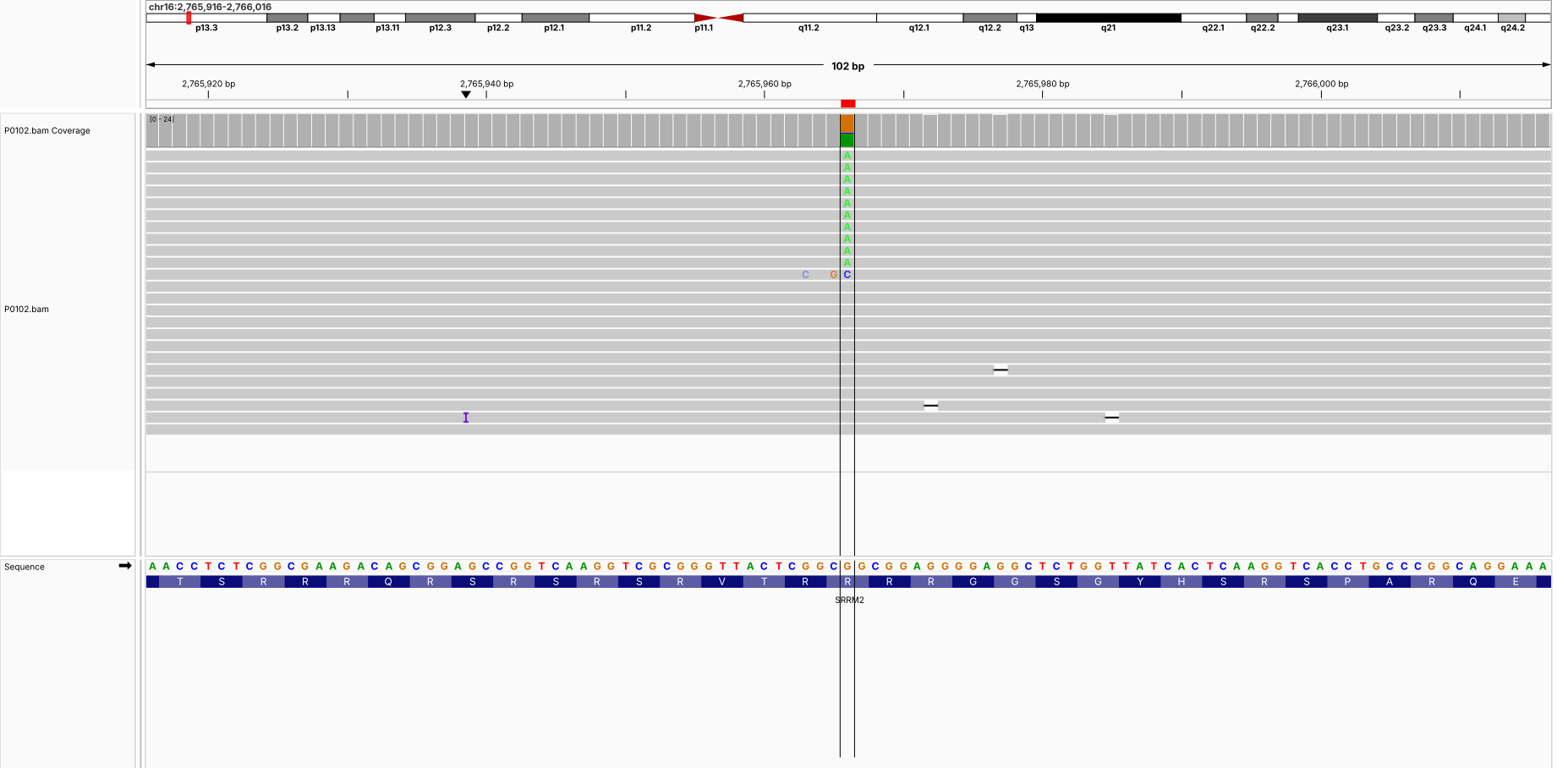

**P0103 (chr4:139,384,930-139,385,030) NAA15**

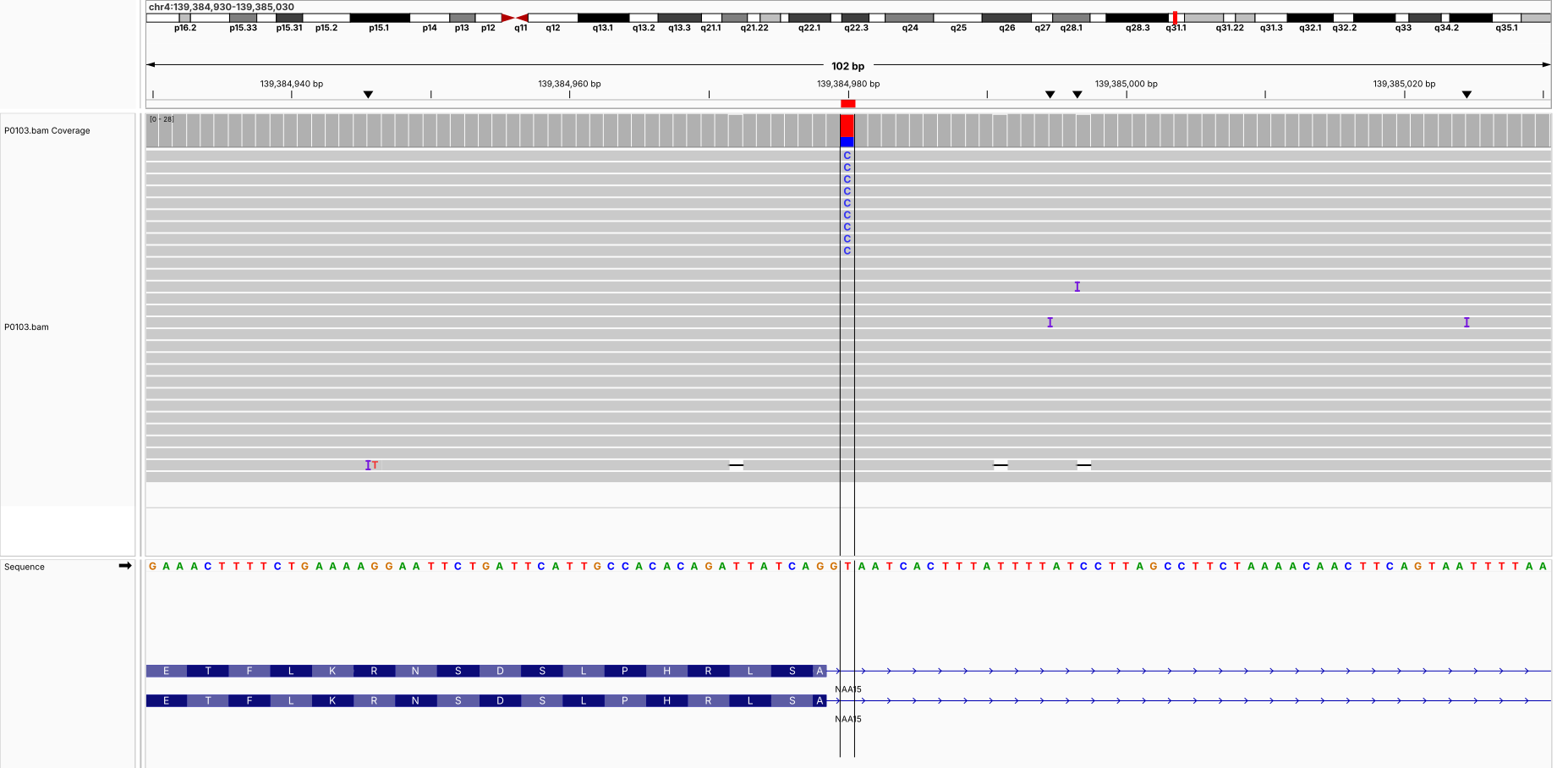

**P0109 (chr19:14,151,502-14,151,602) ADGRL1**

**P0110 (chrX:148,956,423-148,956,523) AFF2**

**P0114 (chr5:14,498,590-14,498,690) TRIO**

**P0114 (chr7:105,090,050-105,090,150) KMT2E**

**P0114 (chr19:14,150,980-14,151,080) ADGRL1**

**P0115 (chr2:1,943,255-1,943,355) MYT1L**

**P0115 (chr20:62,064,975-62,065,075) TAF4**

**P0118 (chr19:15,187,265-15,187,365) NOTCH3**

**P0118 (chr20:62,065,741-62,065,841) TAF4**

**P0120 (chr2:134,456,688-134,456,788) TMEM163**

**P0120 (chr9:70,535,937-70,536,037) TRPM3**

**P0123 (chr21:37,512,420-37,512,520) DYRK1A**

**P0130 (chr1:27,549,647-27,549,747) AHDC1**

**P0134 (chr17:31,299,486-31,299,586) NF1**

**P0137 (chr16:2,085,211-2,085,311) TSC2**

**P0147 (chr8:67,434,109-67,434,209) CPA6**

**P0147 (chr8:67,434,145-67,434,245) CPA6**

**P0147 (chr17:2,667,039-2,667,139) PAFAH1B1**

**P0156 (chr6:43,007,221-43,007,321) PPP2R5D**

**P0167 (chr4:183,680,170-183,680,270) TRAPPC11**

**P0168 (chr12:13,567,103-13,567,203) GRIN2B**

**P0189 (chr2:196,222,259-196,222,359) HECW2**

**P0201 (chr3:193,643,328-193,643,428) OPA1**

**P0201 (chr9:74,634,684-74,634,784) RORB**

**P0201 (chr11:72,579,324-72,579,424) PDE2A**

**P0201 (chrX:118,542,960-118,543,060) DOCK11**

**P0207 (chr7:45,064,597-45,064,697) CCM2**

**P0207 (chrX:154,437,117-154,437,217) GDI1**

**P0216 (chr14:102,956,189-102,956,289) CDC42BPB**

**P0217 (chr1:7,670,999-7,671,099) CAMTA1**

**P0218 (chr10:87,960,998-87,961,098) PTEN**

**P0218 (chr16:24,797,520-24,797,620) TNRC6A**

**P0221 (chr2:218,809,680-218,809,780) CYP27A1**

**P0221 (chr2:218,814,138-218,814,238) CYP27A1**

**P0221 (chr14:21,394,092-21,394,192) CHD8**

**P0222 (chr5:14,363,680-14,363,780) TRIO**

**P0223 (chrX:123,404,668-123,404,768) GRIA3**

**P0255 (chr4:15,574,145-15,574,245) CC2D2A**

**P0255 (chr4:15,601,298-15,601,398) CC2D2A**

**P0263 (chr19:35,726,300-35,726,400) KMT2B**

**P0269 (chr7:105,081,642-105,081,742) KMT2E**
