## Supplementary File 3 for "Pangenome Discoveries from Clinical Long-Read Genomes in Underrepresented Populations with Neurodevelopmental Disorders"

**STR IGV Snapshots**

**P0088 (chr19:45,770,154-45,770,317) DMPK**

DM1_DMPK_pathogenic

| **Sample** | **Motif** | **GT** | **Repeats (MC)** | **Allele bp (AL)** | **Depth** | **Purity** | **Methyl** |
| --- | --- | --- | --- | --- | --- | --- | --- |
| P0088 | CAG | 1/2 | 5 / 440 | 17 / 1327 | 15 / 9 | 0.944 / 0.998 | – / 0.70 |

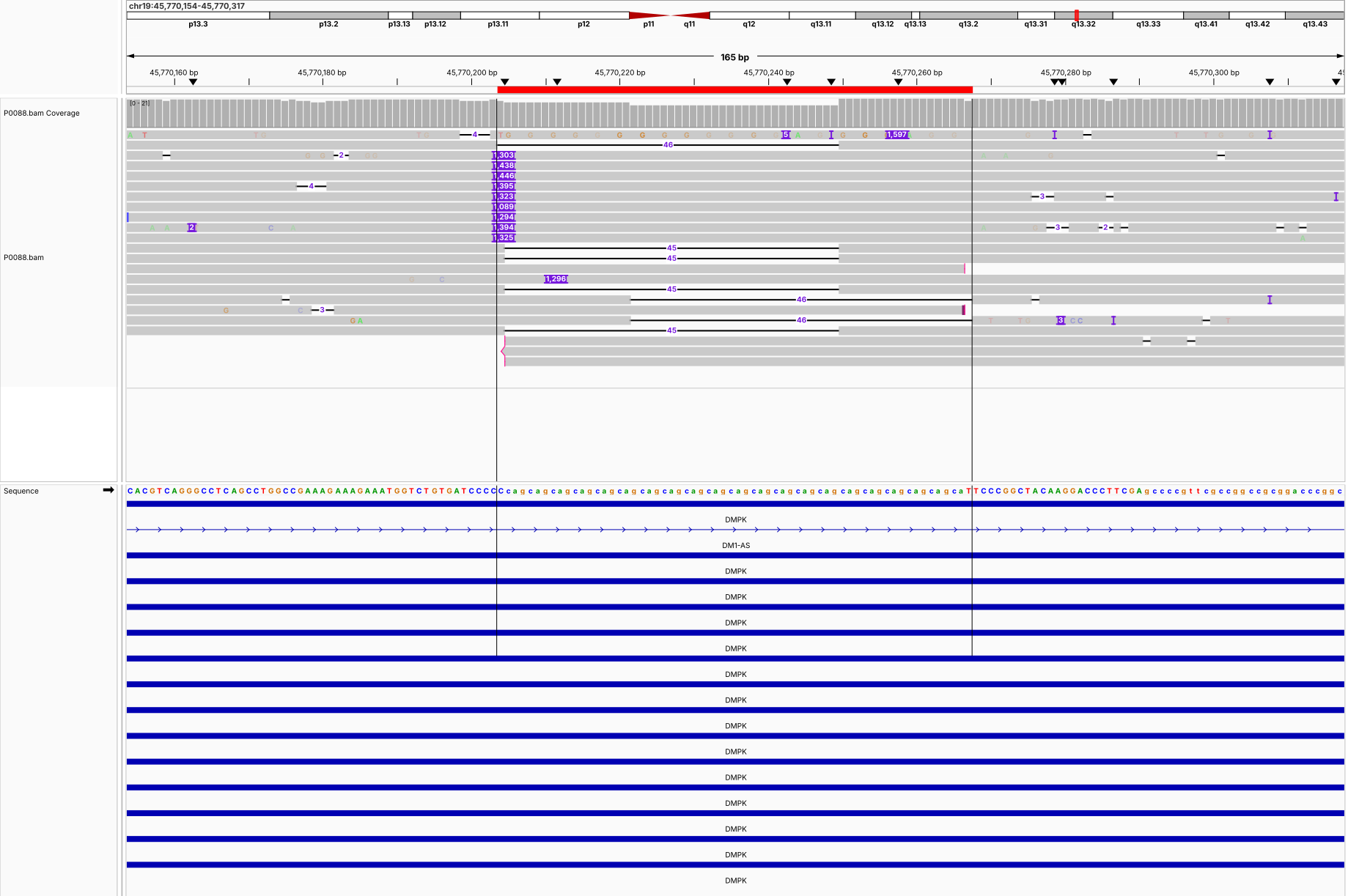

**P0077 (chr21:43,776,393-43,776,530) CSTB**

EPM1_CSTB_pathogenic

| **Sample** | **Motif** | **GT** | **Repeats (MC)** | **Allele bp (AL)** | **Depth** | **Purity** | **Methyl** |
| --- | --- | --- | --- | --- | --- | --- | --- |
| P0077 | CGCGGGGCGGGG | 1/2 | 38 / 40 | 456 / 478 | 11 / 5 | 1.000 / 0.996 | 0.29 / 0.08 |

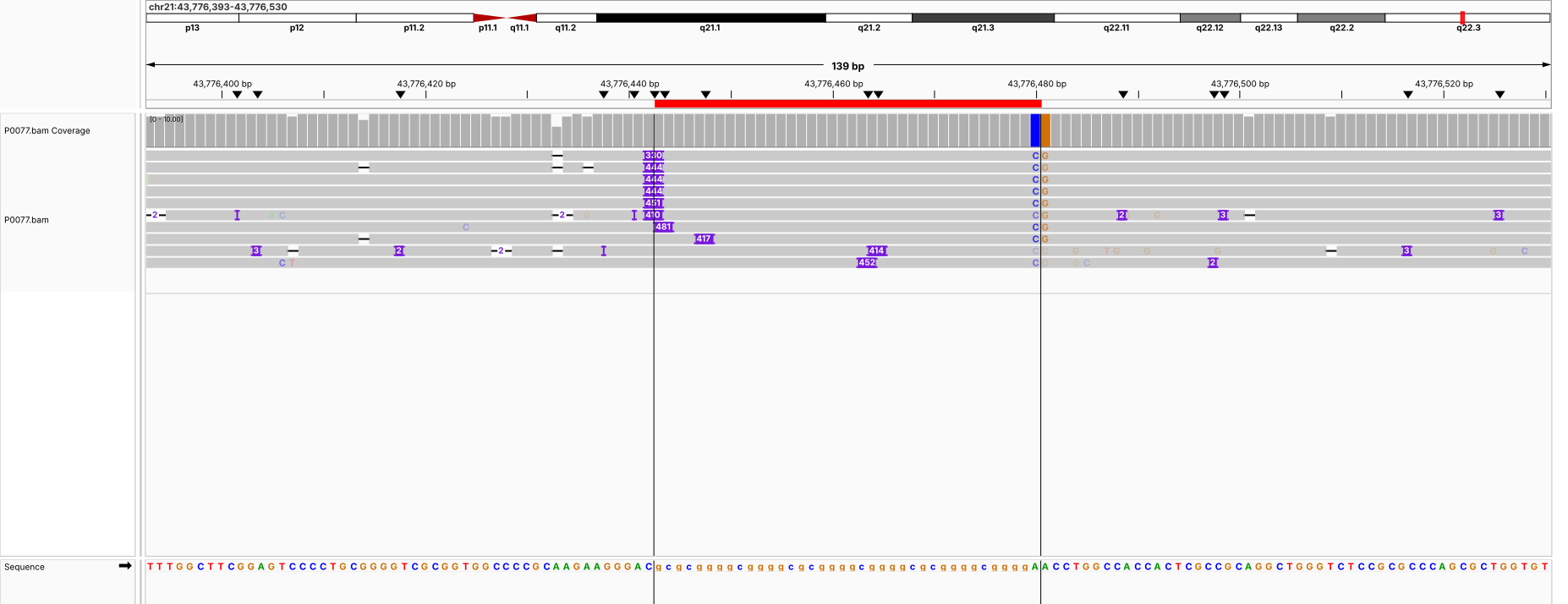

**P0195 (chr18:55,586,105-55,586,278) TCF4**

FECD3_TCF4_pathogenic

| **Sample** | **Motif** | **GT** | **Repeats (MC)** | **Allele bp (AL)** | **Depth** | **Purity** | **Methyl** |
| --- | --- | --- | --- | --- | --- | --- | --- |
| P0195 | CAG | 1/2 | 23 / 82 | 69 / 249 | 20 / 18 | 1.000 / 0.996 | – / 0.46 |

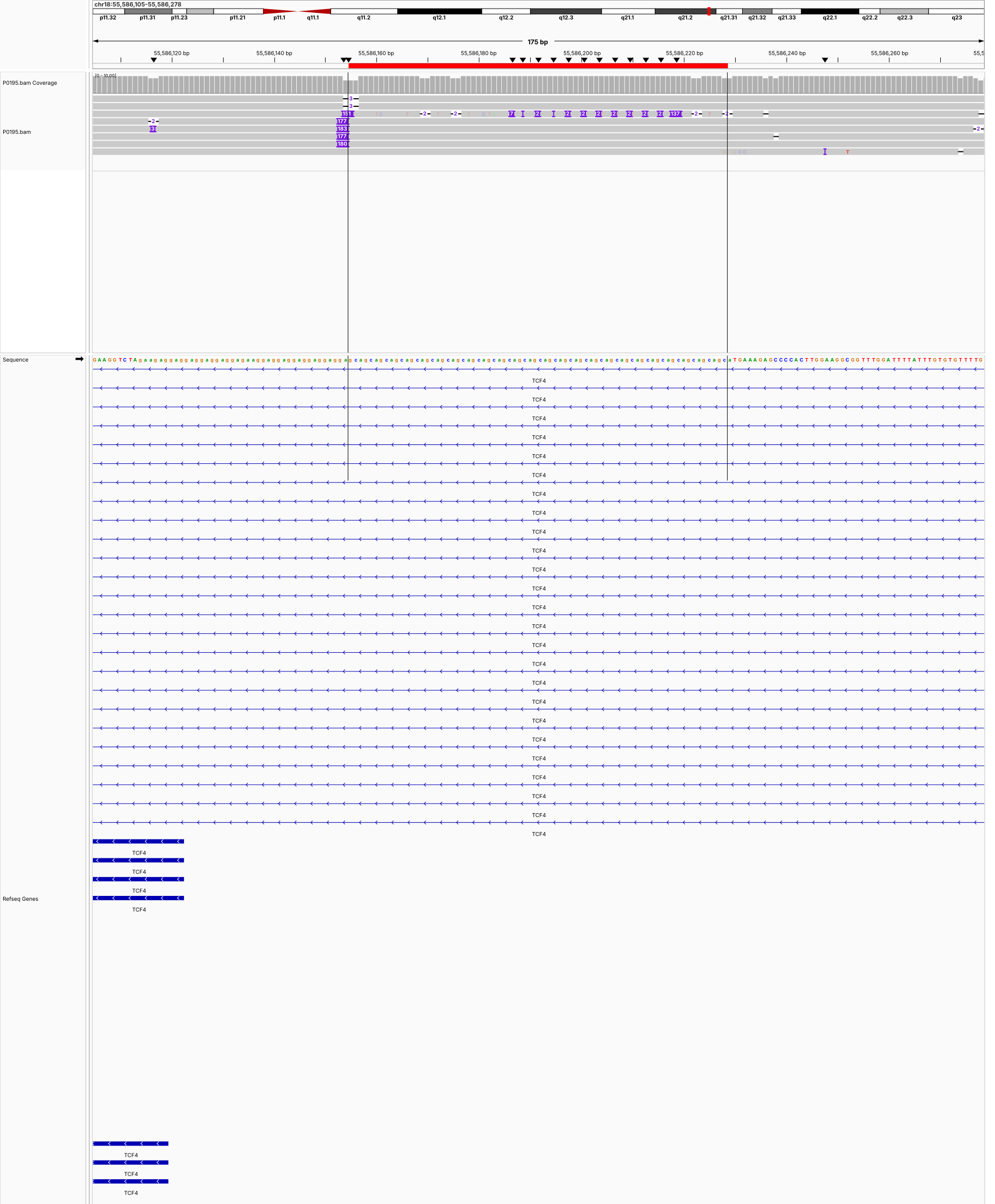

**P0241 (chr18:55,586,105-55,586,278) TCF4**

FECD3_TCF4_pathogenic

| **Sample** | **Motif** | **GT** | **Repeats (MC)** | **Allele bp (AL)** | **Depth** | **Purity** | **Methyl** |
| --- | --- | --- | --- | --- | --- | --- | --- |
| P0241 | CAG | 1/2 | 11 / 91 | 33 / 273 | 10 / 11 | 1.000 / 1.000 | – / – |

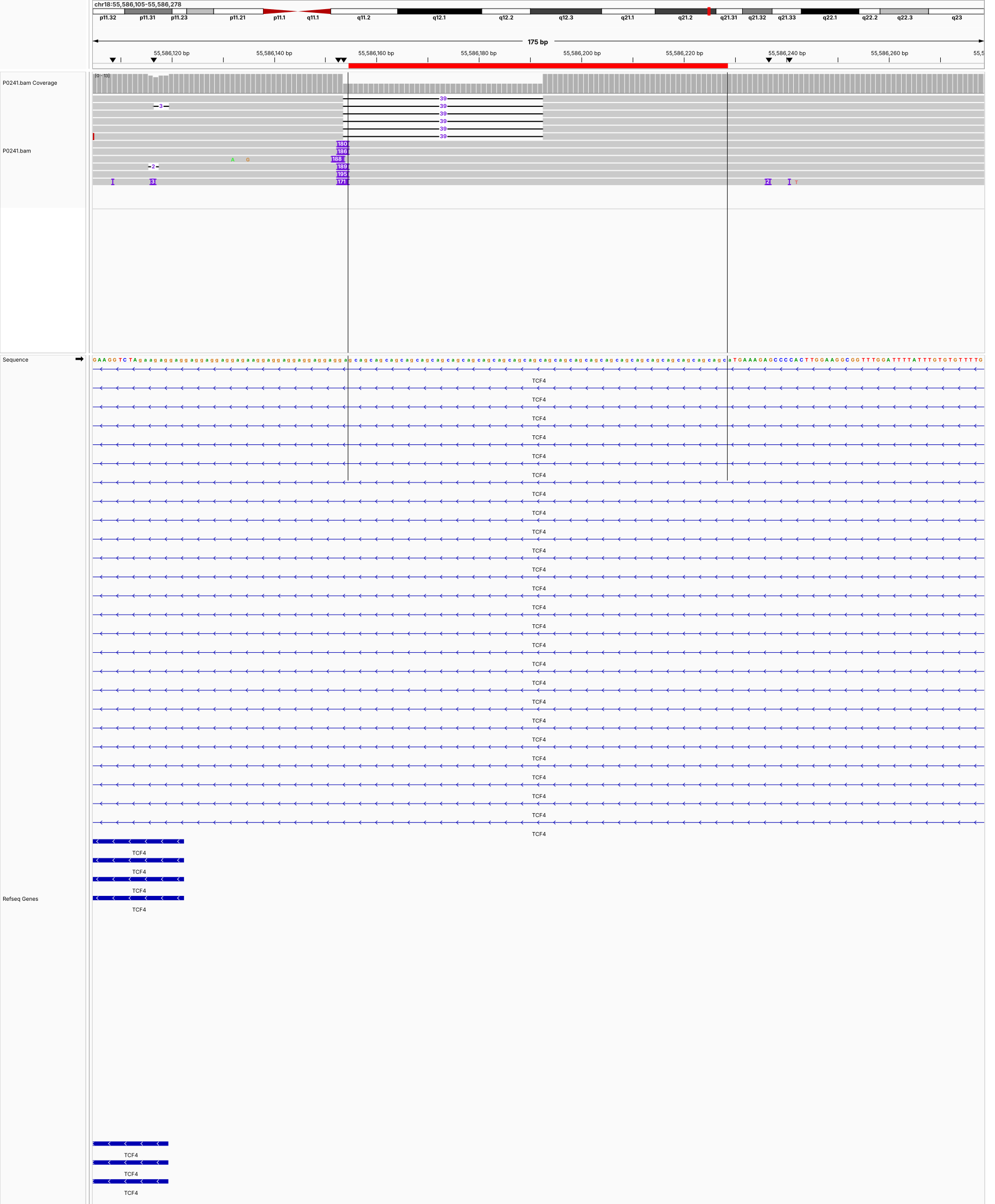

**P0226 (chr12:50,504,952-50,505,073) DIP2B**

FRA12A_DIP2B_pathogenic

| **Sample** | **Motif** | **GT** | **Repeats (MC)** | **Allele bp (AL)** | **Depth** | **Purity** | **Methyl** |
| --- | --- | --- | --- | --- | --- | --- | --- |
| P0226 | GGC | 0/1 | 6 / 447 | 20 / 1365 | 16 / 7 | 0.952 / 0.991 | 0.33 / 0.27 |

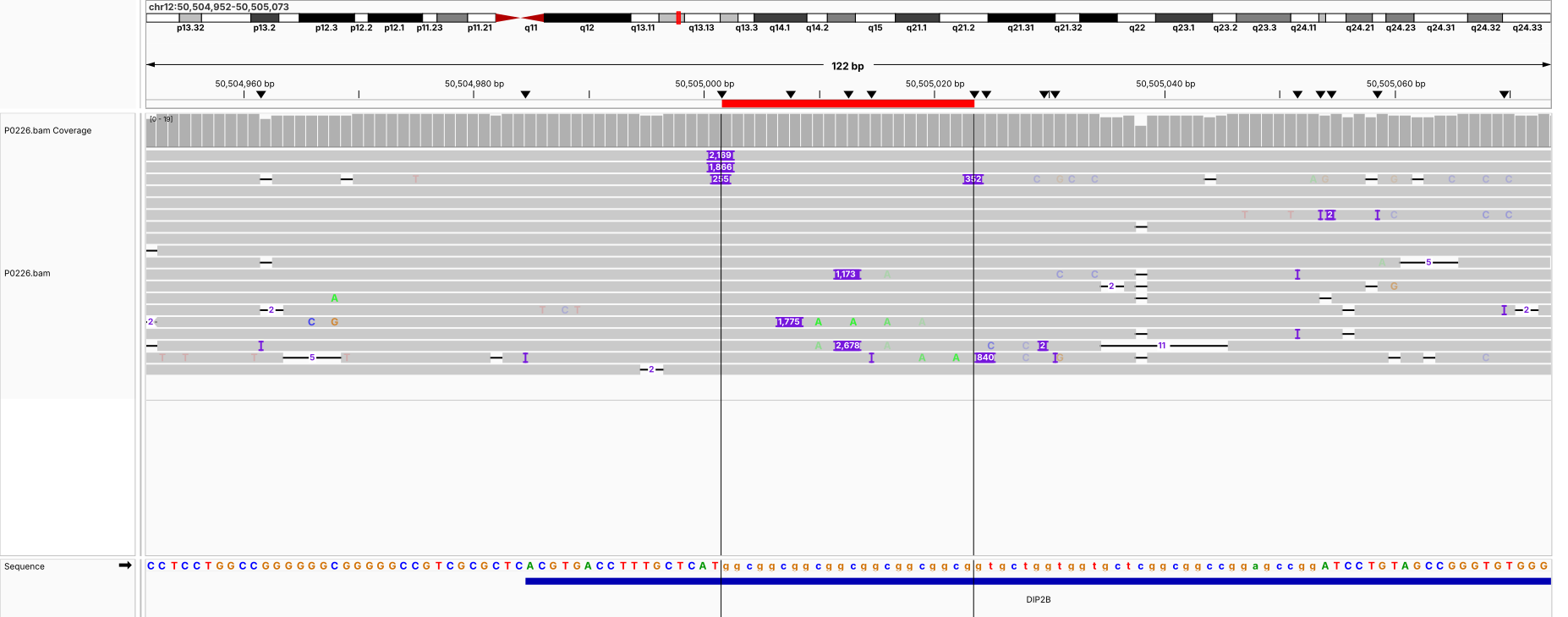

**P0212 (chrX:147,911,987-147,912,162) FMR1**

FXS_FMR1_pathogenic

| **Sample** | **Motif** | **GT** | **Repeats (MC)** | **Allele bp (AL)** | **Depth** | **Purity** | **Methyl** |
| --- | --- | --- | --- | --- | --- | --- | --- |
| P0212 | CGG | 1 | 605 | 1844 | 6 | 0.992 | 0.88 |

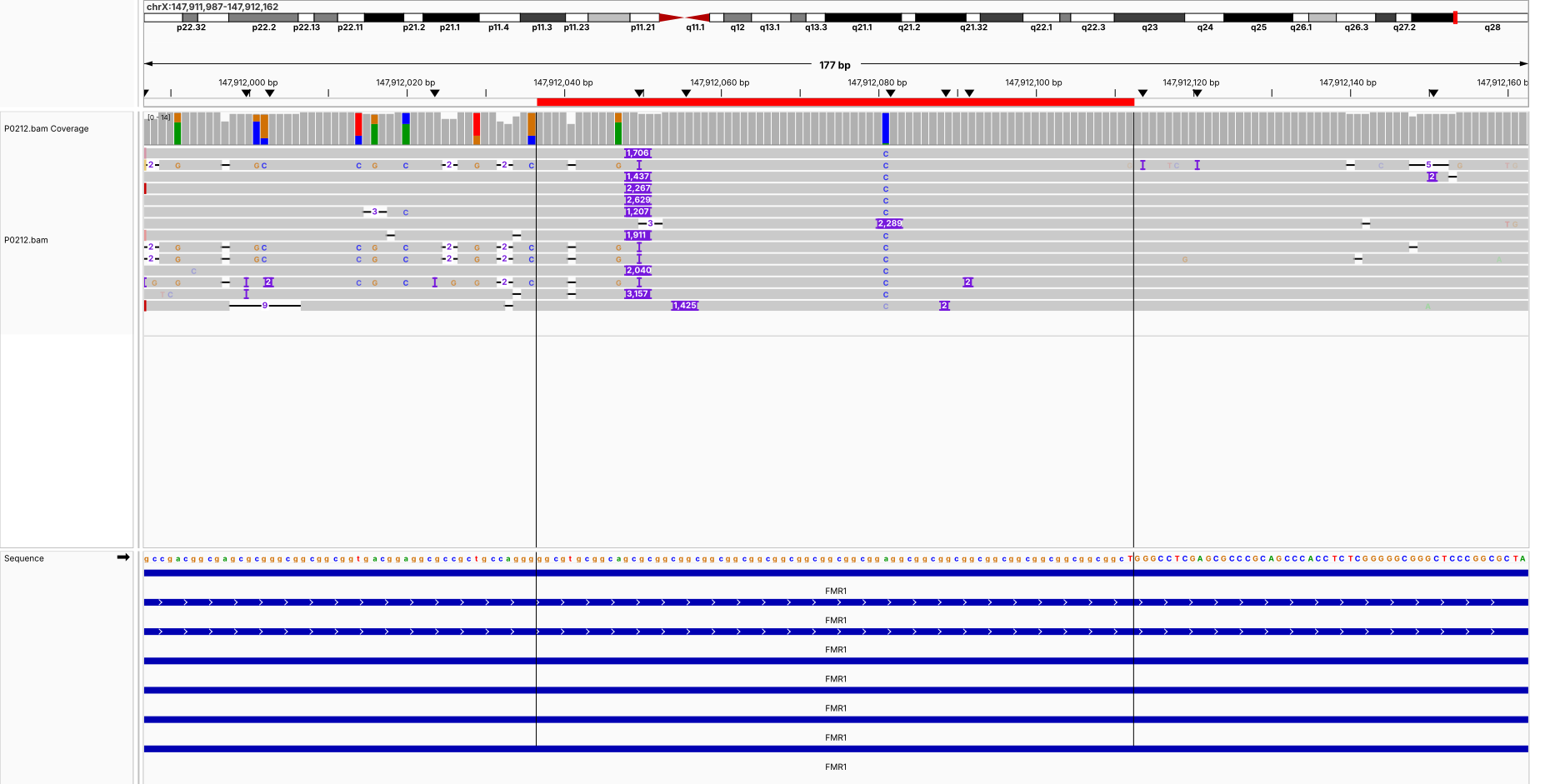

**P0213 (chrX:147,911,987-147,912,162) FMR1**

FXS_FMR1_pathogenic

| **Sample** | **Motif** | **GT** | **Repeats (MC)** | **Allele bp (AL)** | **Depth** | **Purity** | **Methyl** |
| --- | --- | --- | --- | --- | --- | --- | --- |
| P0213 | CGG | 1 | 641 | 1950 | 6 | 0.993 | 0.90 |

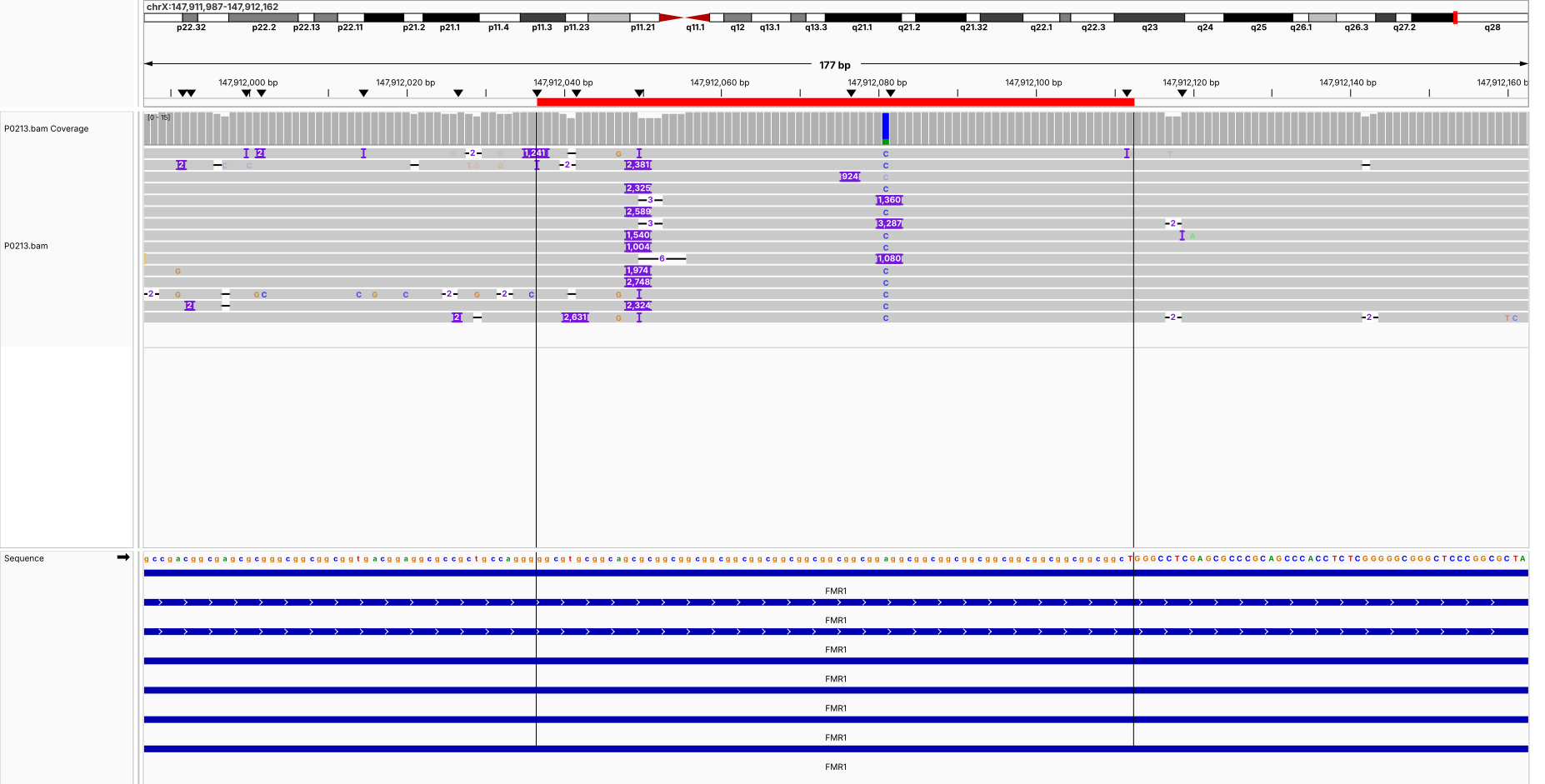

**P0242 (chrX:147,911,987-147,912,162) FMR1**

FXS_FMR1_pathogenic

| **Sample** | **Motif** | **GT** | **Repeats (MC)** | **Allele bp (AL)** | **Depth** | **Purity** | **Methyl** |
| --- | --- | --- | --- | --- | --- | --- | --- |
| P0242 | CGG | 1 | 844 | 2565 | 7 | 0.993 | 0.91 |

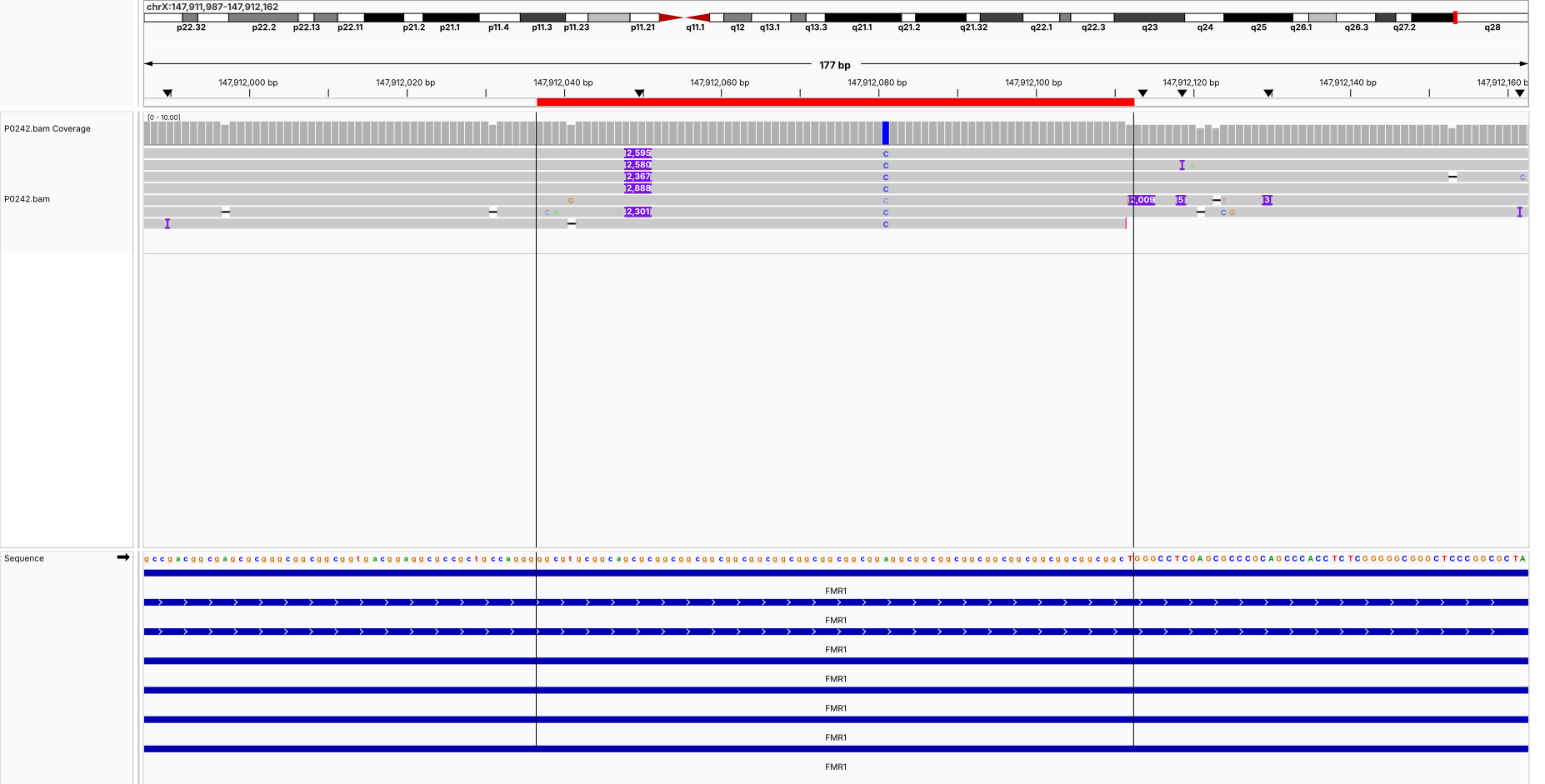

**P0243 (chrX:147,911,987-147,912,162) FMR1**

FXS_FMR1_pathogenic

| **Sample** | **Motif** | **GT** | **Repeats (MC)** | **Allele bp (AL)** | **Depth** | **Purity** | **Methyl** |
| --- | --- | --- | --- | --- | --- | --- | --- |
| P0243 | CGG | 1 | 28 | 101 | 20 | 0.931 | 0.07 |

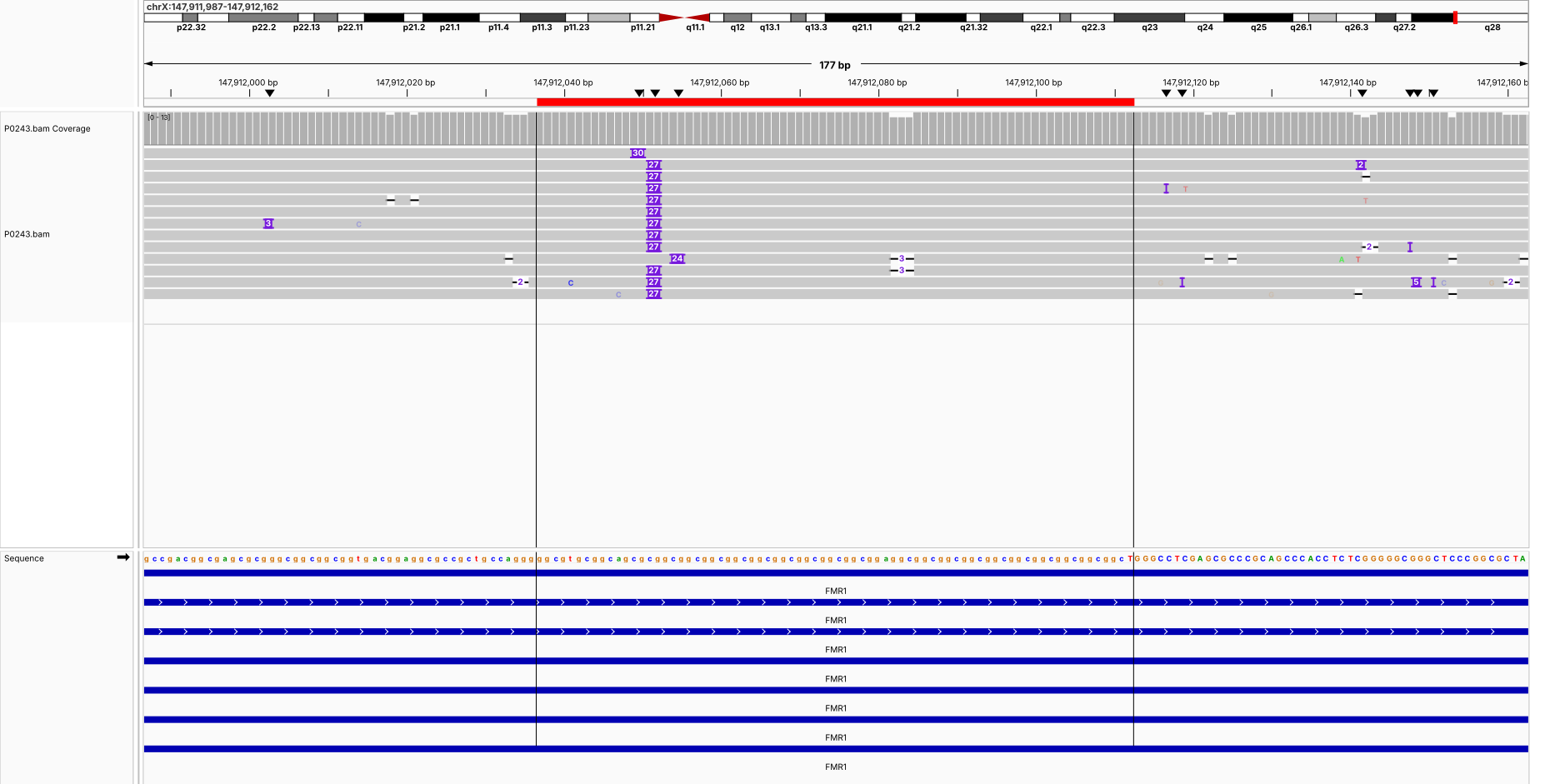

**P0244 (chrX:147,911,987-147,912,162) FMR1**

FXS_FMR1_pathogenic

| **Sample** | **Motif** | **GT** | **Repeats (MC)** | **Allele bp (AL)** | **Depth** | **Purity** | **Methyl** |
| --- | --- | --- | --- | --- | --- | --- | --- |
| P0244 | CGG | 1/2 | 30 / 113 | 107 / 350 | 21 / 10 | 0.935 / 0.986 | 0.33 / 0.84 |

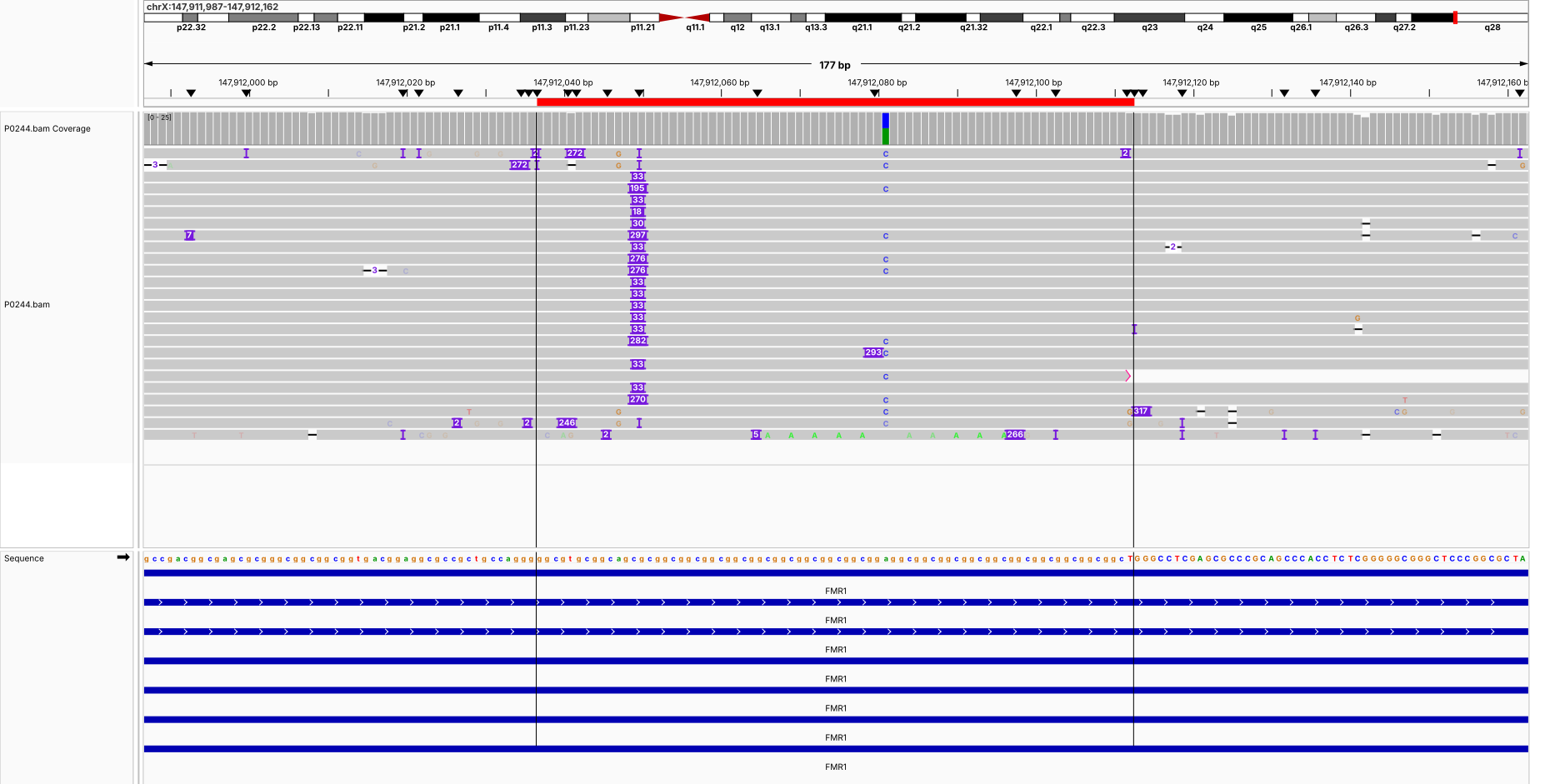

**P0242 / P0243 / P0244 (chrX:147,911,899-147,912,251) FMR1 trio**

FXS_FMR1_pathogenic

| **Sample** | **Motif** | **GT** | **Repeats (MC)** | **Allele bp (AL)** | **Depth** | **Purity** | **Methyl** |
| --- | --- | --- | --- | --- | --- | --- | --- |
| P0242 | CGG | 1 | 844 | 2565 | 7 | 0.993 | 0.91 |
| P0243 | CGG | 1 | 28 | 101 | 20 | 0.931 | 0.07 |
| P0244 | CGG | 1/2 | 30 / 113 | 107 / 350 | 21 / 10 | 0.935 / 0.986 | 0.33 / 0.84 |

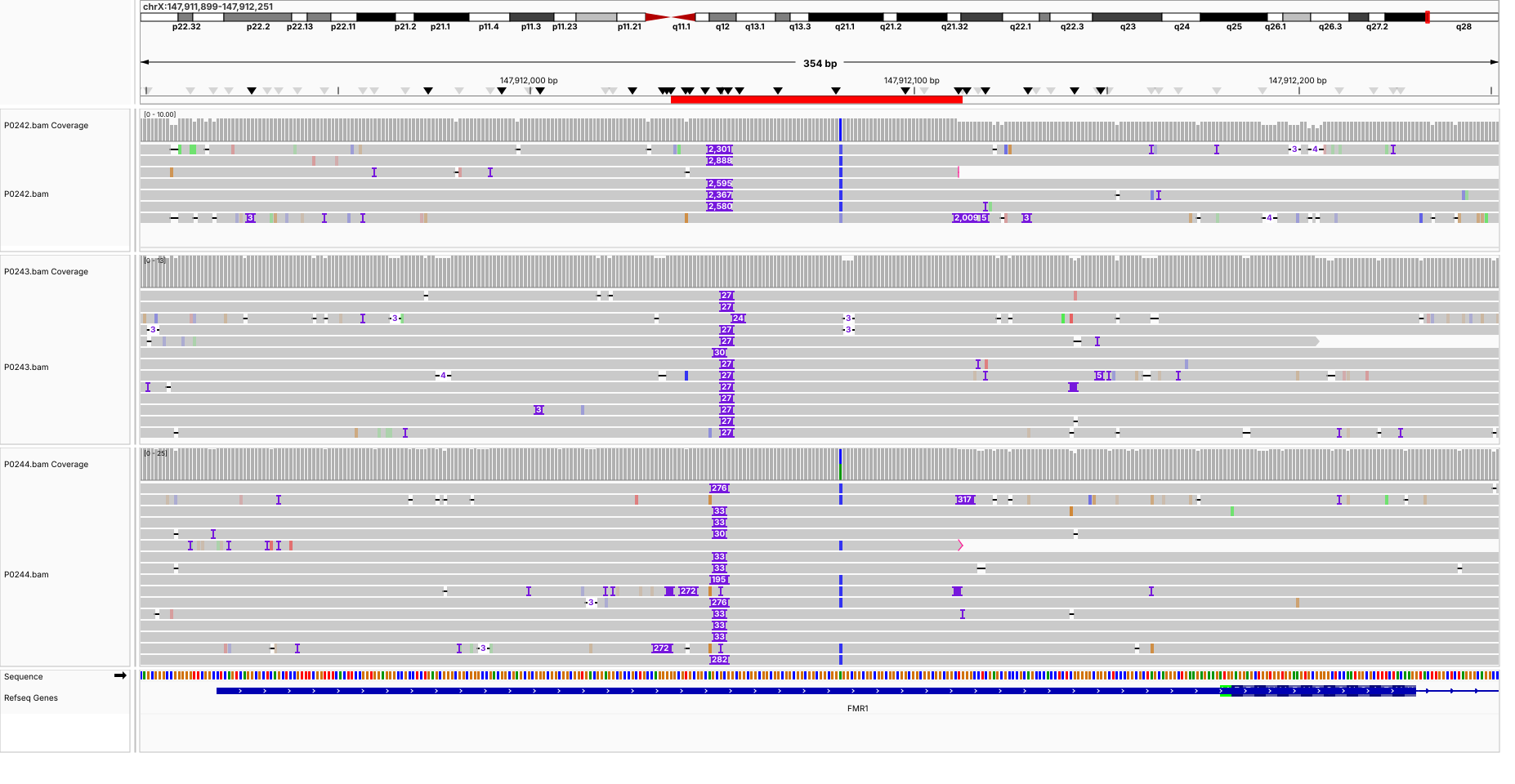
